# Unveiling the Origins and Genetic Makeup of the ‘Forgotten People’: A Study of the Sarmatian-Period Population in the Carpathian Basin

**DOI:** 10.1101/2024.10.04.616652

**Authors:** Oszkár Schütz, Zoltán Maróti, Balázs Tihanyi, Attila P. Kiss, Emil Nyerki, Alexandra Gînguță, Petra Kiss, Gergely I. B. Varga, Bence Kovács, Kitti Maár, Bernadett Ny. Kovacsóczy, Nikoletta Lukács, István Major, Antónia Marcsik, Eszter Patyi, Anna Szigeti, Zoltán Tóth, Dorottya Walter, Gábor Wilhelm, Réka Cs. Andrási, Zsolt Bernert, Luca Kis, Liana Oța, György Pálfi, Gábor Pintye, Dániel Pópity, Angela Simalcsik, Andrei Dorian Soficaru, Olga Spekker, Sándor Varga, Endre Neparáczki, Tibor Török

**Author notes:** These authors contributed to this work equally (Oszkár Schütz, Zoltán Maróti).

## Abstract

The nomadic Sarmatians dominated the Pontic Steppe from 3rd century BCE and the Great Hungarian Plain from 50 CE until the Huns’ 4th-century expansion. In this study, we present the first large-scale genetic analysis of 156 genomes from 1st- to 5th-century Hungary and the Carpathian foothills. Our findings reveal minor East Asian ancestry in the Carpathian Basin (CB) Sarmatians, distinguishing them from other regional populations. Using F4-statistics, qpAdm, and IBD analysis, we show that CB Sarmatians descended from Steppe Sarmatians originating in the Ural and Kazakhstan regions, with Romanian Sarmatians serving as a genetic bridge between the two groups. We also identify two previously unknown migration waves during the Sarmatian era and a notable continuity of the Sarmatian population into the Hunnic period, despite a smaller influx of Asian-origin individuals. These results shed new light on Sarmatian migrations and the genetic history of a key population neighbouring the Roman Empire.

## Introduction

The Sarmatians were a group of nomadic people who likely originated from the southern Ural region during the 4th and 2nd centuries BCE, identified archaeologically with the Prokhorovka culture^1^. In the subsequent centuries, they gradually expanded into the Pontic Steppe territories, displacing the culturally related Scythians^2,3^. During the Iron Age, they established the first significant political formations in the area between the Don, Volga, North Caucasus, and Ural Mountains. Based on names preserved in ancient sources, they are believed to have been part of a group of northern Iranian-speaking people.

By the first century CE, Sarmatian groups had settled in the area between the eastern foothills of the Carpathians and the Lower Danube region (modern Romania)^4^. In the early decades CE, the first Sarmatian tribes, known as the Iazyges, entered the Carpathian Basin, occupying the northern and central areas of the Danube-Tisza interfluve. They then gradually expanded into the Trans-Tisza region, eventually occupying the entire Great Hungarian Plain, and likely extending their rule over the local Celtic and Scythian groups.

By the end of the 1st and the beginning of the 2nd century CE, despite initially good relations, they gradually became a formidable enemy of the Roman Empire in the Danube region^5^. After the Marcomannic-Sarmatian wars (166-180 AD), their material culture became a peripheral extension of the Roman Empire^6–10^. The dense settlement network in the Carpathian Basin within a century of their arrival indicates that the nomadic herders also adopted farming and achieved a large population size^11–13^. Despite this, many steppe traditions persisted in daily life, culture, and warfare. Sarmatians in the eastern Carpathian Basin maintained close contact with other Sarmatian groups in the steppe, and archaeological finds show several instances of eastern groups moving in during the late 2nd and 4th centuries^5,14,15^. They also formed close contacts and military alliances, with Germanic tribes (Quadi, Marcomanni, Vandals), which significantly influenced their material culture, particularly in the surrounding border areas^16–18^.

Previous archaeological research has classified Sarmatian archaeological remains in the Carpathian Basin into three chronological periods, based on the main historical events of Roman-Sarmatian relations and changes in material culture^9,19^. These periods are: 1. Early Sarmatian Period: From the arrival of the Sarmatians in the Great Hungarian Plain (~50 CE) to the 2nd half of the 2nd century CE. 2. Middle Phase: From the period of the Marcomannic wars to the end of the 3^rd^ century CE abandonment of Dacia. 3. Late Period: From the end of the 3rd century CE to the last third of the 5th century CE. Unfortunately, the Sarmatian archaeological chronology of the Carpathian Basin is not fully aligned with the chronological systems used for the Central European Germanic and Eastern European regions^20^.

Between the late 4th and mid-5th centuries, a major migration initiated by the Huns brought diverse communities, including Eastern Germanic tribes, Huns, and other eastern Sarmatian groups, into the Great Hungarian Plain^21,22^. After the Hunnic empire moved its center to the Carpathian Basin, many Sarmatians remained in their original homeland. Their cemeteries were used until the early 5th century, and their settlements continued until the mid-5th century collapse of the Hun Empire^12,22,23^. According to written sources, the Sarmatians may have maintained an independent political organization until the 470s^4^. After the fall of the Hun Empire, they were assimilated into the population of the Gepid Kingdom^24,25^

To date, 45 published Sarmatian genomes, are available across 7 different studies, from the Ural region and the Central Steppe^26–32^. Among these studies, only two articles provide a detailed discussion of the Sarmatians in context^28,32^, while the others address the topic only marginally. Key characteristics identified in the Uralic Sarmatians and Easten Steppe Sarmatians (collectively referred to as Steppe Sarmatians) include: a) Their genomes exhibit the admixture of three main ancestral components: 70% Steppe Middle-Late Bronze Age (steppe_MLBA), 18% Bactria–Margiana Archaeological Complex (BMAC)-related, and 12% Baikal Early Bronze Age (Baikal_EBA)-Khovsgol-related. b) Their lower Khovsgol-related East Asian component compared to the Eastern Scythians suggests they might have originated from distinct, independent late Bronze Age populations in the Ural area. c) Despite their extensive geographical distribution and relatively high genetic diversity, they remained genetically very homogeneous for over 500 years.

From the Carpathian Basin 17 Sarmatian-Period individuals were published in^33^. While detailed analyses were not provided, these genomes display a distinct shift towards European genetic profiles compared to Steppe Sarmatians, raising questions about the potential relationships between these populations.

To clarify the origins and genetic relationships of the Carpathian Basin Sarmatians (CB Sarmatians) and to explore their connections to other populations from the Eurasian Steppe, as well as to local groups from preceding and succeeding periods, we sequenced 156 genomes from the Carpathian Basin and surrounding regions, spanning the Sarmatian and Hun periods (Figure 1A). We have shown that the CB Sarmatians are descendants of the Sarmatians from the Ural and Kazakhstan regions, who migrated from the Carpathian foothills in present-day Romania. The descendants of the substantial CB Sarmatian population formed a significant portion of the population during the subsequent Hun era.

**Figure 1:**
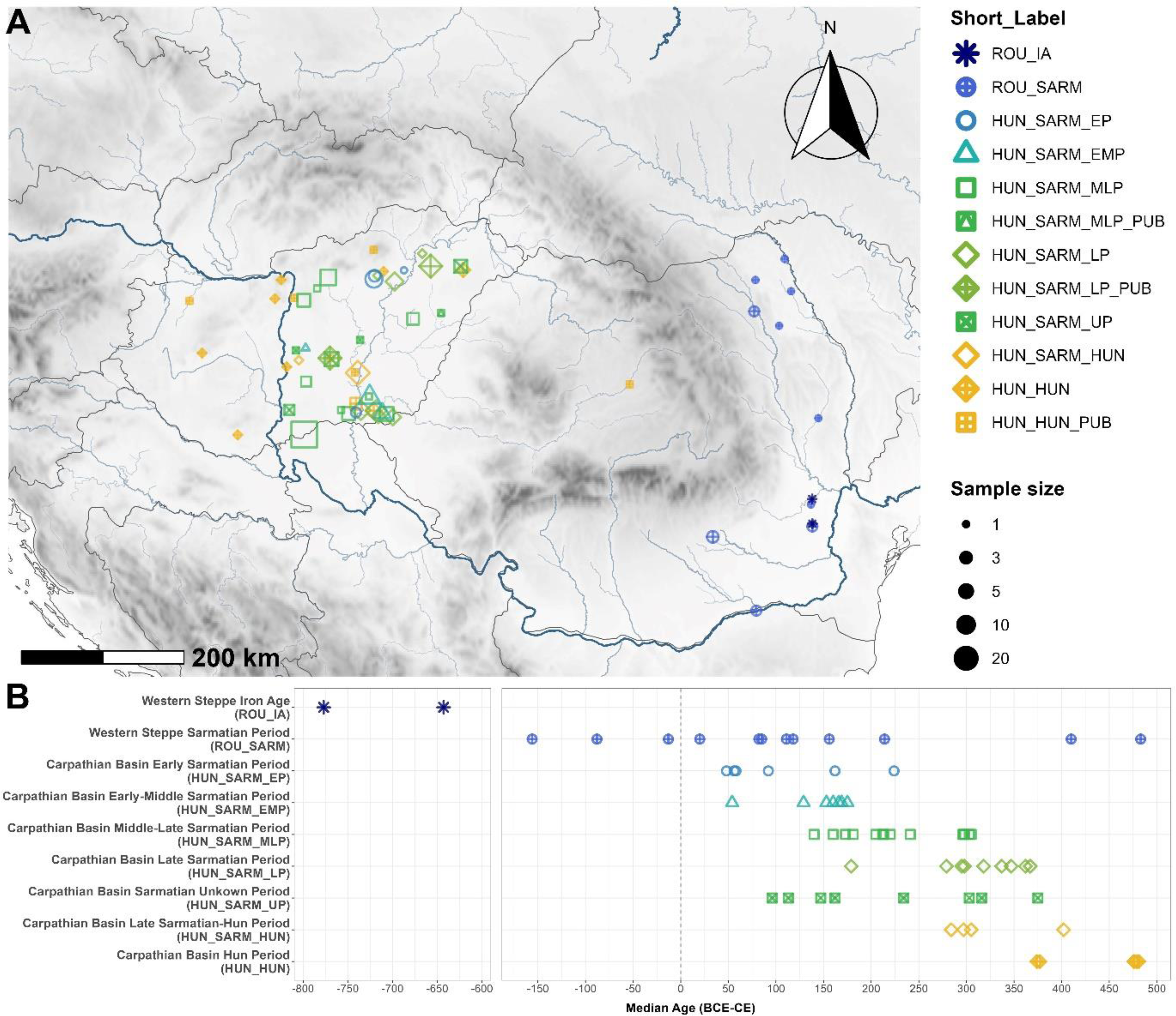
Archaeological sites and chronology of the studied samples. A) Map showing the locations of the individuals analysed in this study. Reanalysed published samples are marked with the suffix “PUB” and are labelled as HUN_SARM_MLP_PUB (n=1), HUN_SARM_LP_PUB (n=16) and HUN_HUN_PUB (n=9). B) Chart depicting the median age of the samples analysed by radiocarbon dating. A gap is included to account for the significantly earlier radiocarbon dates of the two Iron Age individuals.

## Results

### Samples

Out of the 156 samples, 118 were collected from the Great Hungarian Plain, spanning the 1st to 4th centuries, representing the Sarmatian Period population of the Carpathian Basin Barbaricum, termed HUN_SARM. Two cemeteries used during the Sarmatian period clearly extended into the Hun period, leading to their classification as HUN_SARM_HUN. To gain insights into the arrival of the Sarmatians, we also sampled 17 individuals from the Romanian Plains, likely representing the incoming Sarmatian population (ROU_SARM). Additionally, we generated 21 whole genome sequences from the 4th to 5th centuries (HUN_HUN) to assess the potential long-term impact of the Sarmatian population on the region. The 156 shotgun-sequenced whole genomes have a mean coverage of 1.42x (ranging from 0.24x to 3.75x) with negligible contamination (Table S1a).

The newly sequenced genomes were co-analyzed with 17 Sarmatian Period individuals published in Gnecchi-Ruscone et al. (2022)^33^ and 9 Hun Period individuals from Maróti et al. (2022)^34^, creating the most comprehensive genomic database of the region for these periods (Figure 1A; Table S1b).

The samples underwent a thorough review for accurate archaeological classification (for further details, see Data S1). We also conducted radiocarbon measurements on 68 samples to anchor and validate the archaeological dating approach (Figure 1B, Table S1c). The two approaches yielded concordant dates in most cases; however, some samples showed indications of a possible reservoir effect (see Supplemental Data). For this reason, we did not rely solely on radiocarbon results for sample grouping but instead integrated verified archaeological data with radiocarbon dating. This led to the creation of a separate group for individuals with uncertain periodization, termed HUN_SARM_UP. Radiocarbon dating of two Romanian samples, LMO-8 and RAK-7, revealed that these individuals were significantly older, dating to the Early Iron Age, aligning with their sparse archaeological descriptions. We include these individuals in the publication as Romania Iron Age (ROU_IA).

Based on the integrated dating procedure, we classified our samples into nine groups representing progressive archaeological periods: ROU_IA, ROU_SARM, HUN_SARM_EP, HUN_SARM_EMP, HUN_SARM_MLP, HUN_SARM_LP, HUN_SARM_UP, HUN_SARM_HUN, and HUN_HUN (Short Labels are referred to in Table S1a-b and other supplementary tables and figures).

Genetic kinship was determined using correctKin^35^. We identified 15 kin groups some of which connects the Sarmatian and Hun Periods directly to the Avar Period (Table S1e).

### Genetic diversity of the Sarmatian and Hun period samples

To determine the underlying genetic structure among our samples we first applied Principal Component Analysis (PCA) and ADMIXTURE^36,37^. We calculated primary PC axes from a present-day Eurasian population set (Table S2a) and projected the studied genomes onto these axes (Figure 2A and Table S2b). In these analyses we also included the published 45 Steppe Sarmatians grouped by their geographic origin and age.

**Figure 2:**
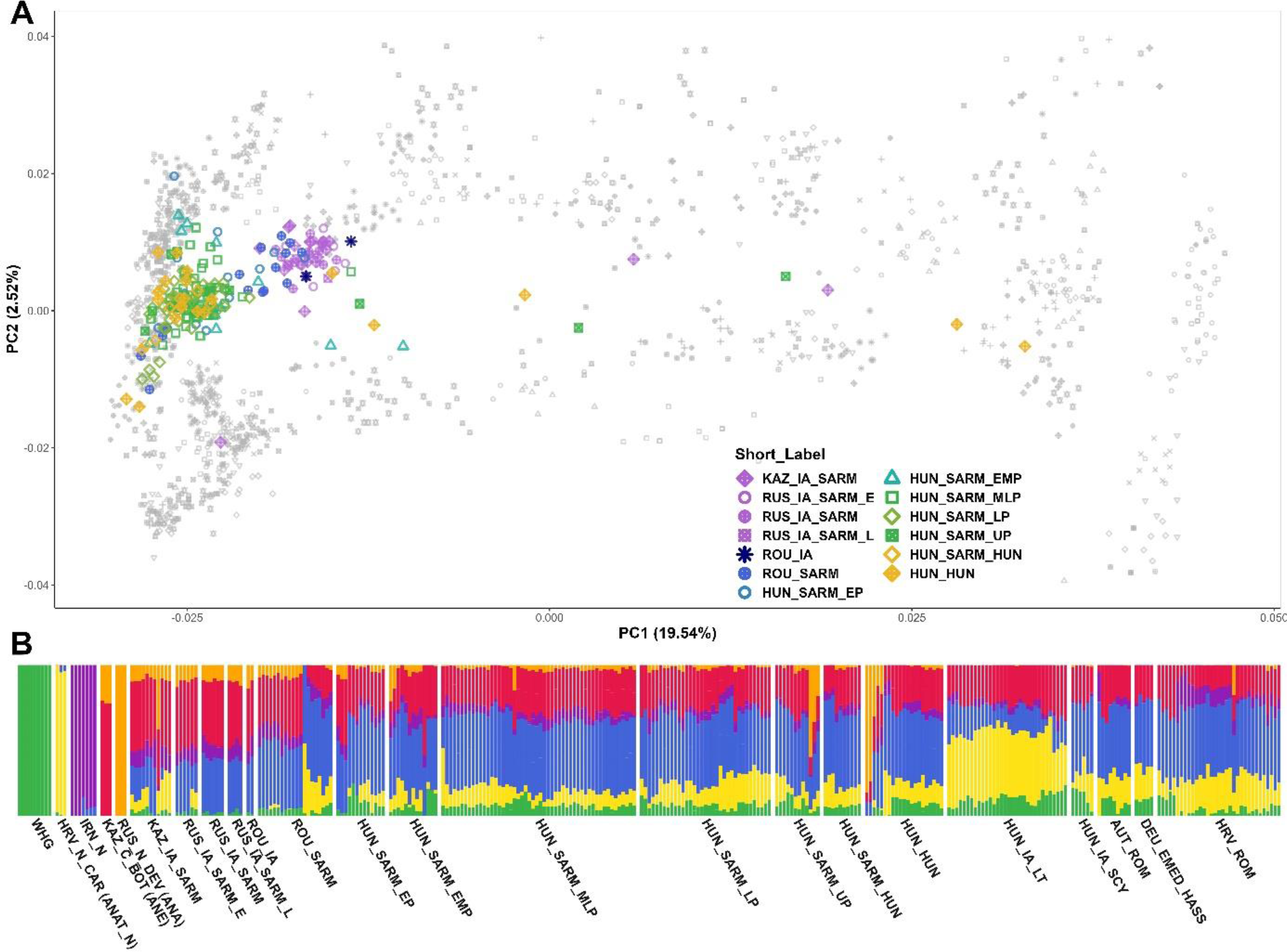
PCA and ADMIXTURE. A) PCA of the studied individuals and all available Sarmatians from the literature projected onto a modern Eurasian background (gray symbols). B) Unsupervised ADMIXTURE analysis results (K=6) of the same samples arranged in chronological order. On the right we also show selected contemporary populations from the Carpathian Basin for comparison. Groups best representing the main ancestral components are shown at the left side. Abbreviations are listed in Table S3a.

In line with previous findings, the Steppe Sarmatians are closely clustered together on the PCA, with a few outliers, and are distinctly separated from European populations. By contrast, most CB Sarmatians cluster near present-day Central European populations, only a few individuals, mainly from the early and early-middle periods, show a strong affinity towards the Steppe Sarmatian cluster. Within the European cluster, some individuals map more closely to modern Northern European populations, while others align more with Southern Europeans. Additionally, several genetic outliers are shifted towards Asian populations, suggesting connections to other nomadic groups beyond the steppe Sarmatians. This ancestry, however is also seen among some Steppe Sarmatian outliers, raising the possibility that these genetic outliers might have arrived alongside Steppe Sarmatians. Notably, a significant portion of the Romanian Sarmatians is positioned among the Steppe Sarmatians, while others form a cline between the Steppe and Carpathian Basin Sarmatians.

The unsupervised ADMIXTURE results clearly indicate the genomic components responsible for the differing PCA positions of the individuals (Table S3a). At K=6, we identified the typical macroregional ancestry components: Western Hunter-Gatherer (WHG), Anatolia Neolithic (ANAT_N), Iran Neolithic (IRAN_N), Ancient North Eurasian (ANE), and Ancient Northeast Asian (ANA) (Figure 2B). Additionally, a distinctive component (blue in Figure 2B) appeared, maximized in Carpathian Basin individuals, likely due to the overrepresentation of samples from this region in the analysis.

Most Sarmatian and Hun period individuals from the Carpathian Basin closely resemble contemporaneous populations from the region (e.g., Austria and Croatia Roman period - AUT_ROM, HRV_ROM) or populations from the immediately preceding period (e.g., Hungary Scythians - HUN_IA_SCY and Hungary Celtic - HUN_IA_LT). However, a distinctive feature of our studied groups is the presence of a small but significant fraction of the ANA component. This component is highest in the earliest groups (HUN_SARM_EP and EMP) and appears to decline over successive periods. The ANA component is significantly more pronounced in the Steppe Sarmatians and Romanian Sarmatians, who have very similar genome compositions, corresponding to their clustering on PCA and their significant shift from European groups. The PCA Asian outlier individuals also stand out in the ADMIXTURE analysis due to their significant ANA component, which is much larger than that of the Steppe Sarmatians. The two Iron Age individuals from Romania show very surprising ADMIXTURE patterns, displaying identical component ratios with the RUS_IA_SARM group (Table S3a), which is also reflected by their close PCA positions. This sharply distinguishes them from the contemporaneous Scythian and Celtic populations (HUN_IA_SCY or HUN_IA_LT).

### Carpathian Basin Sarmatians show genetic affinity with Steppe Sarmatians

The East Asian ancestry found in the CB Sarmatians sets them apart from their contemporary neighbours and suggests that they may have descended from Steppe Sarmatians, as supported by historical and archaeological sources. To test the potential genetic affinity between the two Sarmatian populations, we first reanalysed the published Steppe Sarmatian individuals with sufficient coverage (Supplemental Data). We were able to cluster them into two homogeneous yet similar groups, termed STEPPE_IA_SARM_URAL, STEPPE_IA_SARM_STEPPE, which we used as proxies in our ancestry analysis.

We applied F4-statistics in which we co-analysed the Sarmatian and Hun period samples with the populations that previously inhabited the Carpathian Basin. First, we measured the direct affinity of the samples towards the Steppe Sarmatians against a Late Neolithic Carpathian Basin population (Lengyel culture) possibly representing local elements, with the statistics: F4(Ethiopia_4500BP, Test; HUN_LN_LGY, STEPPE_IA_SARM_URAL). Positive values in this statistic indicate a major affinity towards the Steppe Sarmatian proxy, while negative values show more shared drift towards the local proxy. This was plotted together with another combination: F4(Ethiopia_4500BP, STEPPE_IA_SARM_URAL; HUN_LN_LGY, Test), which uses the same references, but actually measures the samples’ affinity towards the Sarmatian proxy if we exclude their shared drift with the local proxy (Figure 3A, Table S4a).

**Figure 3:**
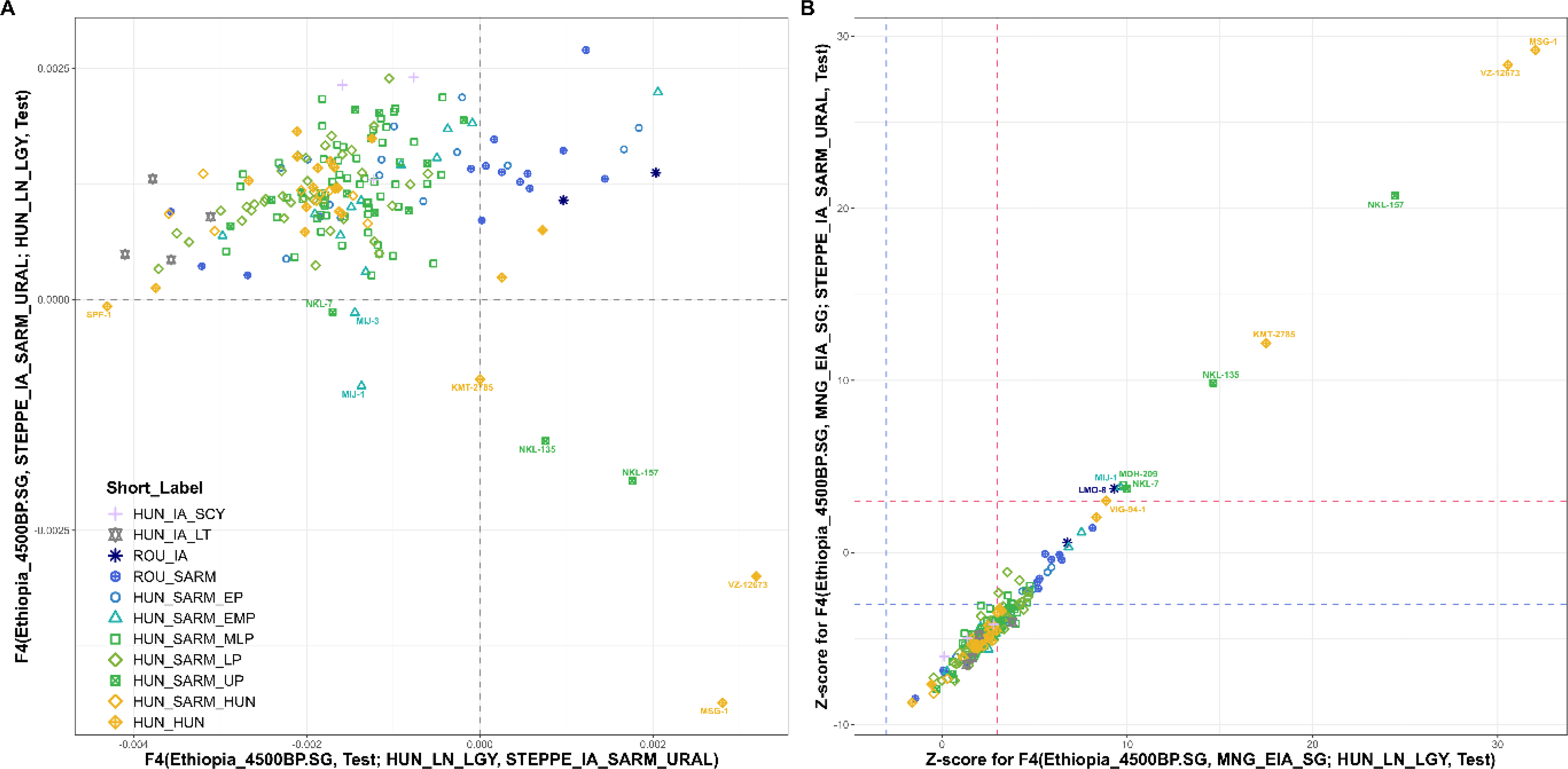
F4-statistic results from multiple F4 operational frameworks. A) F-scores of Steppe Sarmatian affinity measured in our samples and other Iron Age groups against local Carpathian Basin ancestry. The X axis shows the two-way affinity of the samples to either the local or the Sarmatian proxy, while the Y axis measures the affinity toward the Steppe Sarmatian proxy after excluding their local marker set. B) Z-scores of East Asian affinity of the samples measured by excluding their local marker set (X axis) or their marker set potentially shared with the Steppe Sarmatians (Y axis). Blue segmented lines indicate the negative significance threshold (−3), red lines indicate the positive significance threshold (3). Samples with exceptionally high East Asian affinity are labelled.

Figure 3A illustrates that most samples, including those from the Scythian and Celtic groups in the Iron Age Carpathian Basin, share the majority of their markers with the local proxy. However, a few individuals, particularly from the ROU_SARM and HUN_SARM_EP groups, are positioned on the positive side of the X axis. This trend is further supported by the Y axis, where nearly all individuals appear on the positive side, indicating at least a limited affinity toward the Steppe Sarmatian proxy in addition to their local affinity. The genomes located in the upper right quadrant of Figure 3A suggest a strong Steppe Sarmatian affinity. Notably, these samples also exhibit overlapping PCA positions and similar ADMIXTURE patterns to those of Steppe Sarmatians.

A few individuals in the lower right quadrant show strong negative F-scores on the Y axis but retain Steppe Sarmatian affinity on the X axis. They also have an elevated East Asian genomic component, as seen in PCA and ADMIXTURE, distinguishing them from the Steppe Sarmatian proxy. On the other hand, these samples share significant drift with Steppe Sarmatians on the X axis, likely due to their East Asian ancestry. This raises the possibility that other samples with Steppe Sarmatian affinity on the Y axis might show similar patterns because of their elevated East Asian ancestry.

To address this uncertainty, we conducted two additional F4 analyses: 1. F4(Ethiopia_4500BP, MNG_EIA_SG; HUN_LN_LGY, Test) that measures the East Asian affinity of the samples while excluding markers shared with the local proxy. 2. F4(Ethiopia_4500BP, MNG_EIA_SG; STEPPE_IA_SARM_URAL, Test) that assesses the same East Asian affinity but excludes markers shared with the Steppe Sarmatian proxy. In Figure 3B, we plotted Z-scores instead of F-values, as the significance level of the statistics provides a more precise answer to our question.

The X axis of Figure 3B demonstrates that all individuals on the right side of Figure 3A show significant shared drift with the East Asian proxy beyond their local ancestry. However, the Y axis reveals that the East Eurasian affinity of the samples in the upper right quadrant of Figure 3A matches that of the Uralic Sarmatians, as this affinity becomes non-significant when Uralic Sarmatian markers are excluded. In contrast, individuals in the lower right quadrant of Figure 3A exhibit significant Z-scores on both axes, indicating that they possess a higher level of East Asian genetic ancestry than can be explained by the Steppe Sarmatian proxy alone. The presence of these genetic types suggests a population independent of the Sarmatians.

While the two Iron Age groups from the Carpathian Basin, HUN_IA_SCY and HUN_IA_LT, show some affinity toward the Steppe Sarmatian proxy in Figure 3A (especially HUN_IA_SCY), this affinity likely stems from a different component. This is supported by Figure 3B, where these groups fall well below any significance line, indicating no detectable East Asian genetic affinity.

Next, we devised a qpAdm analysis framework to determine whether Steppe Sarmatians are essential for modelling the CB Sarmatians. Based on preliminary qpAdm runs, we assembled a comprehensive set of 15 source populations to best represent the potential local Carpathian Basin elements in most individuals. The two Steppe Sarmatian groups were used as representative sources for the proposed Sarmatian immigrants. Additionally, we included 8 other sources, consisting of Central and Inner Asian populations with known connections to the Carpathian Basin or those that could represent further Asian immigrants independent of the Sarmatians. For further details on the analysis framework and the exact composition of the LEFT and RIGHT populations see STAR Methods and Table S5.

Two individuals, DZS-41 and FKD-150, who were among the earliest Sarmatians in the Carpathian Basin according to archaeological data and radiocarbon dating, along with one Romanian Sarmatian (OSU-1), produced unambiguous models, in which they formed a genetic clade with the Steppe Sarmatians. We successfully obtained appropriate 2-source or 3-source models for all other individuals using the local European and Steppe Sarmatian sources (see Table S5b-c), except for 12 samples. To model these remaining samples, we incorporated additional source candidates from a broader spatio-temporal range and successfully developed valid models for all but one individual, HVF-10 (Table S5d).

Among the Romanian Sarmatians, ~2/3 (9 out of 15) had over 50% Steppe Sarmatian ancestry, while only 12% of CB Sarmatians (14 of 118) showed this level.

We need to note that nearly half of the samples had very low East Eurasian components, making it impossible for qpAdm to identify the exact source. These samples included both Sarmatian and other Central-East Asian sources in their feasible models, raising doubts about the precise origin of the East Asian ancestry. Nevertheless, many explicit models clarify some doubts. For instance, several models clearly show Steppe Sarmatian ancestry as a minor component (e.g., MDH-265, A181015). These often appear in the same cemetery as uncertain models, suggesting that Steppe Sarmatian ancestry is the most plausible source.

In summary, the hypothesis tests confirmed the F4 results, showing that most CB Sarmatians are best modelled with Steppe Sarmatian sources. Furthermore, the qpAdm models align with the periodization of our samples. Romanian Sarmatians have the highest Steppe Sarmatian component, consistent with their PCA positions. Among the CB Sarmatians, those with a predominant Steppe Sarmatian ancestry are primarily from the early and middle periods (HUN_SARM_EP and MLP groups). Only one individual from the subsequent Hun period (ASZK-1) shows a majority Steppe Sarmatian ancestry. However, this individual is most likely a Hun period immigrant from the same Steppe Sarmatian population, with a minor East Asian genetic component. Overall, all analyses suggest that Romanian Sarmatians represent a genetic link between the Steppe and CB Sarmatians.

We obtained unambiguous models for the PCA Asian outlier Sarmatians (e.g., MIJ-1, MIJ-3, MDH-209, NKL-135). Their eastern ancestry was modelled from Hun, Xiongnu, or Avar elite sources, but never from Steppe Sarmatians, consistent with their substantial ANA ancestry components. However, most individuals with majority East Asian ancestry are found in the HUN_HUN group (NKL-157, KMT-2785, MSG-1, VZ-12673), indicating 4th-century population changes reported in historical sources. Despite the appearance of new eastern immigrants, most HUN_SARM_LP, HUN_SARM_HUN, and HUN_HUN individuals still exhibit at least marginal Steppe Sarmatian ancestry, highlighting substantial population continuity.

### IBD analysis links all Sarmatian groups

In order to explore the genealogical links across different geographic regions and time periods in Central Europe and the Central Steppe, we conducted Identity-by-Descent (IBD) analyses. For this purpose, we selected and imputed 504 individuals spanning from the Iron Age to the early Middle Ages additionally to the 158 genomes presented in this study.

The main criteria for selecting individuals were their spatio-temporal origin and the library preparation method, excluding capture-enriched genomes to avoid erroneous genotype inferences during imputation. We applied a stringent entry threshold for imputation recommended by^38^, requiring a minimum of 0.5-fold coverage and low contamination, resulting in the exclusion of 7 new samples. Additionally, we excluded the 17 Sarmatian individuals published in Gnecchi-Ruscone et al. 2022^33^ due to their capture sequencing method. The final list of samples used for IBD analyses is provided in Table S6b.

We identified IBD genomic segments of at least 8 centimorgans (cM) in length using the ancIBD software^39^ with optimizations described in the Methods section. IBD connection networks were visualized as graphs using the Fruchterman-Reingold (FR) weight-directed algorithm^40^. In these graphs, individual samples are represented as points (vertices), and IBD connections between points are shown as lines (edges). The algorithm operates iteratively, calculating attractive forces for vertices connected by edges and repulsive forces for points not connected by edges in each cycle. We used the total length of IBDs shared between individuals as weights for calculating the attractive force, making the distance between connected vertices roughly proportional to their genealogical distance.

First, we investigated the genealogical links between different populations across different time periods. For this reason, we grouped the samples by archaeological period and culture. This allowed us to examine intergroup connections among various European and Carpathian Basin groups, including relevant groups from the Central Steppe and Asia. In these analyses the clouds of groups were handled as single points and a group relation layout was calculated with the FR algorithm, where the weights were the number of IBD connections between each group (see details in STAR Methods). This way, the distribution of the clouds themselves actually reflects the connectedness between the groups (Figure 4).

**Figure 4:**
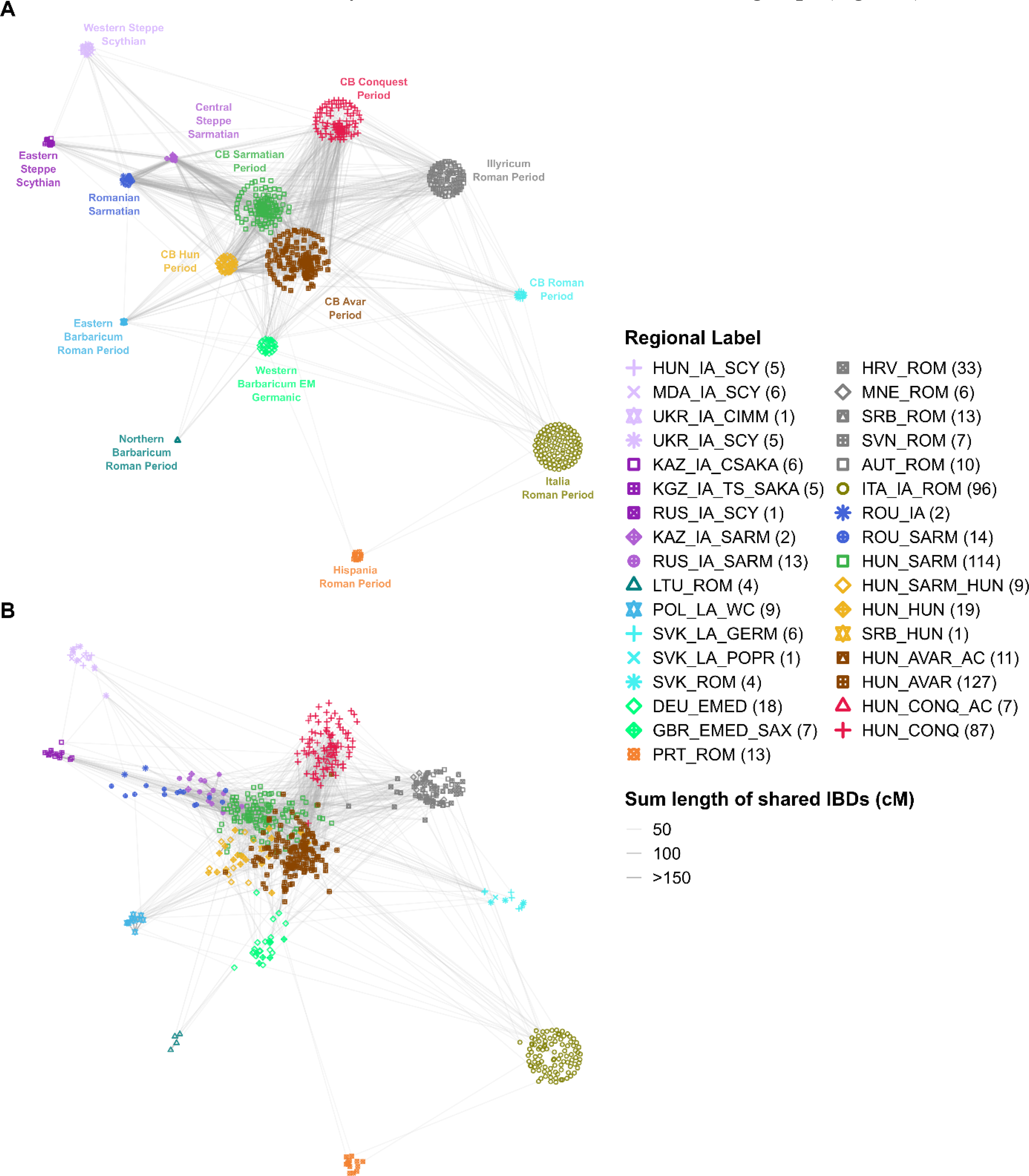
IBD sharing of groups potentially related to Sarmatians. A) Intergroup IBD sharing graph of 662 ancient shotgun genomes including the samples presented in this publication. Individuals were grouped according to geographical region and archaeological period (for further data see Table S6b). B) Vertices were allowed to reposition driven by the FR weight directed algorithm for 100 iterations. The maximum displacement of the points was reciprocally toned down by the number of outgoing connections (edges pointing outside the respective groups). Only edges representing intergroup connections are plotted.

The CB Sarmatians occupy a central position in this plot, alongside the Hun Period and Avar Period individuals from the Carpathian basin published in Maróti et al. 2022^34^. This central position is not surprising, as these groups are populous and occupy a central temporal location, providing them with the greatest opportunities to produce detectable genealogies that connect earlier and later populations. However, this does not undermine the importance of their numerous connections with seemingly distant groups and especially with each other.

Interestingly, steppe-related groups - Scythians, Steppe Sarmatians, and Romanian Sarmatians - all cluster near the CB Sarmatians. These groups share the most IBD segments with the CB Sarmatians and with each other. It is especially notable that within these groups, the Western Scythians (HUN_IA_SCY, MDA_IA_SCY, UKR_IA_CIMM, UKR_IA_SCY) exhibit the fewest connections to any other group, including the nearby and contemporary CB Sarmatians, who have significantly more connections with the geographically more distant Steppe Sarmatians.

Also interesting is the Hun Period group, which barely shows any pattern of intragroup relatedness and instead has a disproportionately high ratio of intergroup connections, especially with individuals from the preceding Sarmatian Period and the subsequent Avar Period. This suggests that the Hun Period group may not represent a distinctly separate population.

These observations are further illustrated in Figure 4B, where we used another FR algorithm with a 100-iteration limit, allowing points to shift between groups based on their primary attraction forces. As can be seen, the Roman provincial samples along the borders of the plot barely moved, reflecting their sparse connectedness to the Carpathian Basin individuals. In contrast, the Sarmatians excavated in Romania and on the Central Steppe moved rapidly inward, clustering with the CB Sarmatians. As expected, the Hun Period individuals also quickly lost their group coherence and shifted towards the Sarmatian and Avar Period individuals.

To better illustrate the IBD sharing patterns on an individual level, we prepared a graph with no grouping (Figure 5). Here we included all studied groups from the Carpathian Basin (HUN_SARM, HUN_SARM_HUN, HUN_HUN, HUN_AVAR, HUN_CONQ) and its vicinity (ROU_IA, ROU_SARM) as well as Steppe Sarmatians (KAZ_IA_SARM, RUS_IA_SARM). From the other groups shown in Figure 4A, we included only 17 samples that had at least 5 IBD connections to the studied individuals. This threshold was chosen to reduce the number of additional individuals to ≤10%, thereby reducing the clutter of the figure and improving its clarity. This resulted in a collection of 423 individuals, who were plotted with the same original coordinate positions as seen in Figure 4A, but the FR algorithm was allowed to freely reposition the points for 1000 iterations. Individual IBD sharing data can be inspected in Table S6c.

**Figure 5:**
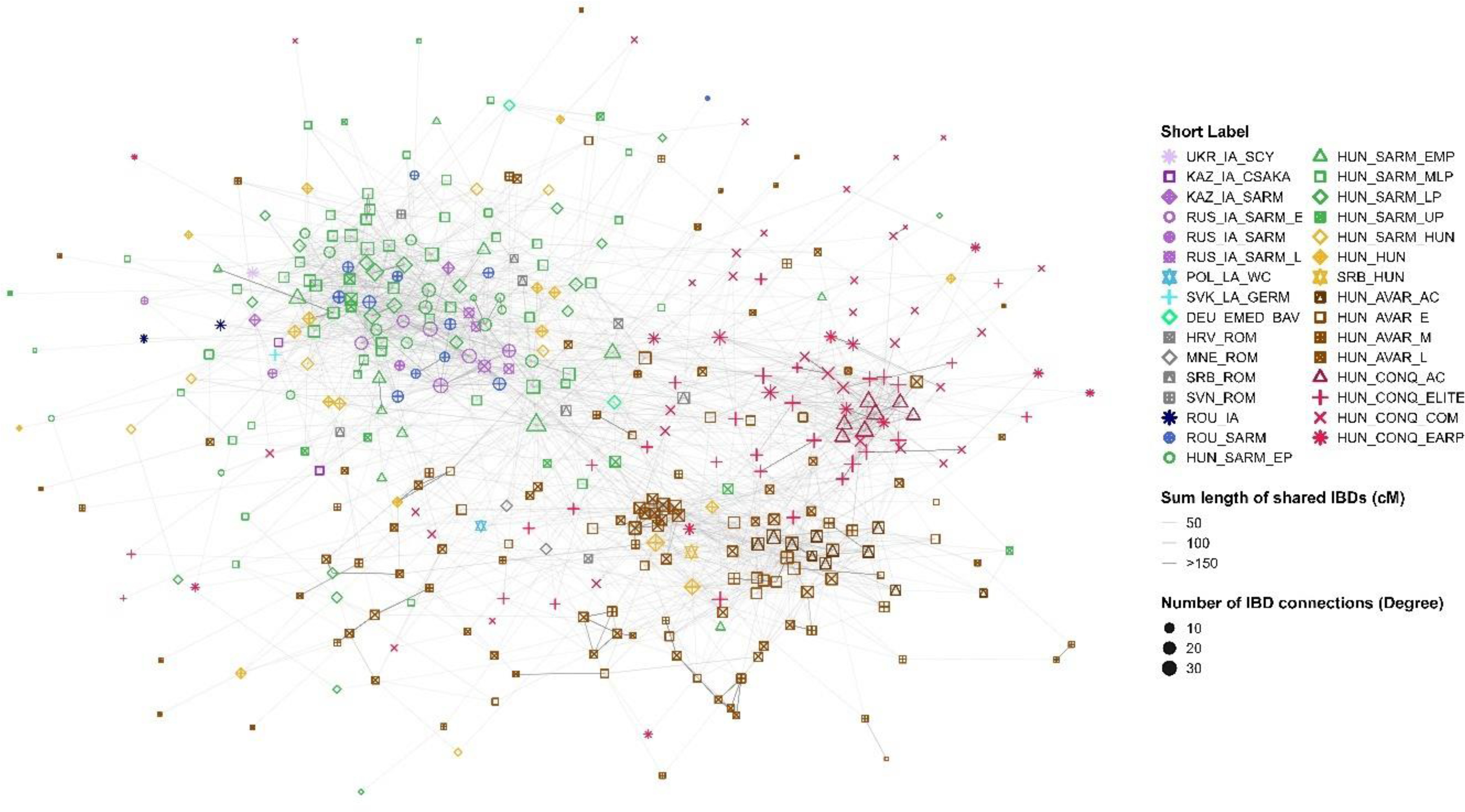
IBD sharing. IBD sharing graph of 423 selected individuals. Points represent individual samples. The size of the points is proportional to the number of connections (degree) a point has. Edges are shaded according to the total lengths of IBDs shared along them in cM. The individuals included in this plot are highlighted in Table S6b.

In Figure 5, the three main groups, Sarmatian, Avar and Conquest period samples from the Carpathian Basin, form distinct clusters, reflecting high intraperiod connectivity. The first-generation immigrant “core” individuals of each medieval group appear to occupy central positions, particularly during the Conquest Period, where they exhibit very high levels of both intra- and intergroup connectedness. During the Sarmatian Period, this central position is seemingly delegated to the Steppe Sarmatians, within the cluster of ROU_SARM and HUN_SARM_EP individuals. Conversely, the Hun Period individuals do not form isolated groupings but instead seamlessly blend into the “Sarmatian cloud.” Notably, some HUN_SARM_UP (e.g., NKL-157) and HUN_HUN individuals (e.g., MSG-1, VZ-12673, KMT-2785) have the majority of their shared IBDs with later Avar Period individuals (Figure S1). This pattern provides further evidence that eastern immigrants distinct from the Steppe Sarmatians also appeared during these periods.

### IBD connections between the periods suggest new migrations

Next, we analysed pairwise combinations of group connectedness across subsequent time frames (Figure 6). We carefully normalized the degree centrality data by dividing the detected connections by the total number of possible connections, resulting in the ratio of fulfilled connections as described in the Methods section. The X-axis in Figure 6 displays the short label of each group (see Table S6b), while the columns represent the ratio of fulfilled IBD connections between the indicated group and every other group.

**Figure 6:**
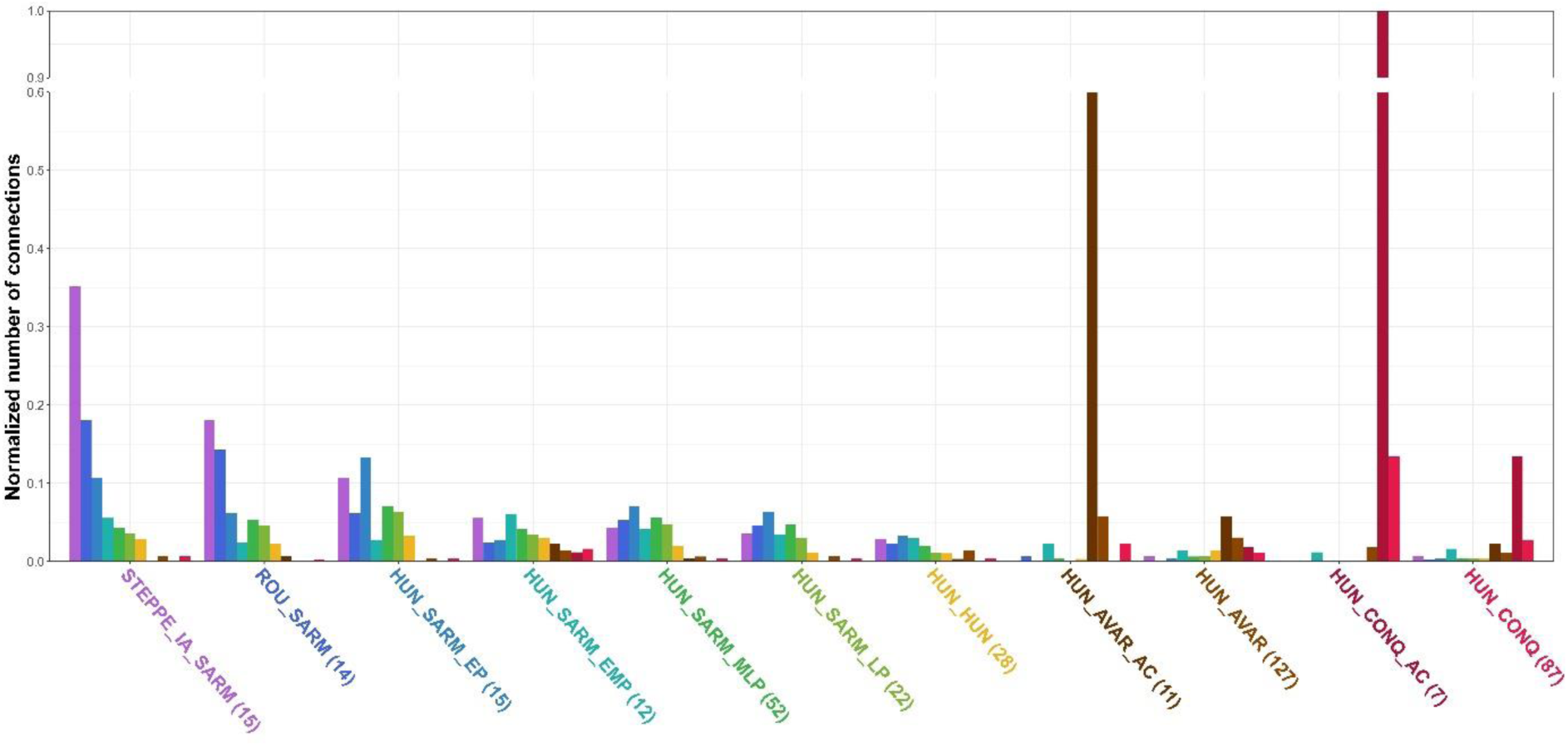
Normalized number of IBD connections across the different archaeological periods of the Carpathian Basin and its vicinity. The colors of the columns correspond to the colors of the group labels, with the height of each column representing the strength of the IBD connections for the group labelled with the letter code. The groups are arranged from left to right in chronological order. The plot has been truncated above the 0.6 line to accommodate the unusually high (100%) intragroup sharing of the HUN_CONQ_AC group. Normalized values were calculated as described in STAR Methods.

Figure 6 illustrates that the Steppe Sarmatians (STEPPE_IA_SARM) exhibit a consistently decreasing sharing pattern across the progressive time periods of the Carpathian Basin. This trend is consistent with a possible founding effect, where the genomic contribution of the earliest group naturally diminishes over subsequent generations. Additionally, the steadily declining pattern of intergroup connections suggests a continuous chain of generational transmission, without any abrupt population turnovers throughout the successive archaeological periods.

A similar trend is observed in the connectedness of the ROU_SARM and HUN_SARM_EP groups with subsequent periods. However, there is a notable sharp decline in their connectedness with the HUN_SARM_EMP group, indicating a significant gap in IBD transmission during the Early-Middle Sarmatian period. The HUN_SARM_EMP group again shows a declining pattern of IBD connections with subsequent periods but also reveals extensive connections with the post-Sarmatian Avar and Conquest period groups (HUN_AVAR, HUN_CONQ), which were negligible in the earlier Sarmatians.

This phenomenon is likely attributed to a second wave of immigration during the EMP period, involving new groups. The HUN_SARM_EMP group is represented by two large cemeteries, Makó-Igási Járandó (MIJ) and Hódmezővásárhely-Fehértó (HVF). F4 and qpAdm analyses identified at least two individuals from MIJ (MIJ-1, MIJ-3) as potential migrants from Eastern or Central Asia. In contrast, four individuals from HVF (HVF-4, HVF-8, HVF-10, HVF-21) showed significant Northern European-related ancestry, with 3 of these individuals being genetic outliers, and we were unable to model HVF-10 accurately (see Table S5). This suggests that the HVF population likely represents new migration from Northern Europe. Nevertheless, the MIJ and HVF cemeteries do not fully represent the entire population of the HUN_SARM_EMP period, as the populations from the HUN_SARM_MLP and LP periods show a much stronger connection with the HUN_SARM_EP group.

The HUN_HUN group has the lowest IBD sharing within itself but shows high connections to Sarmatian and Avar periods, bridging the HUN_SARM_LP and HUN_AVAR groups. However, many individuals from this group were from solitary graves or single representatives, which may underestimate their true intragroup connectedness.

The so-called “immigrant cores” of the later Avar and Conquest Periods (HUN_AVAR_AC and HUN_CONQ_AC) described by Maróti et al. (2022)^34^, exhibit distinctive IBD sharing patterns compared to other groups, with the exception of the STEPPE_IA_SARM. Their prominent intragroup IBD sharing and relatively low sharing with contemporary neighbours indicate a distinct, endogamous population. In contrast, the majority of sequenced individuals from the Avar and Conquest Periods (HUN_AVAR, HUN_CONQ) exhibit a more regular sharing pattern, suggesting they likely represent a broader segment of the population from that era. Despite a reduction in connections, links to Sarmatian Period individuals are still observable in these later groups, indicating that at least a portion of the pre-Hun Period population persisted in the Carpathian Basin. We also visualized the normalised intragroup and intergroup connectivity of each individual from each period in Figure S2, which further highlights that individuals from the three steppe groups, STEPPE_IA_SARM, HUN_AVAR_AC, and HUN_CONQ_AC, display an unusually high ratio of intragroup IBD sharing compared to the other groups.

Plotting intergroup IBD sharing by cemetery (Figure S3) reveals a significant depletion of IBD connections in about half of the Late Sarmatian (HUN_SARM_LP) and Hun period (HUN_HUN) cemeteries compared to earlier periods. This phenomenon can be well explained for the Hun period, where significant new immigrants with elevated East Asian genomic components were detected. Most of their IBD connections are with later Avar and Conquest-period groups (Figure S1), rather than with preceding Sarmatians.

A similar pattern observed in the Late Sarmatian period is also likely due to new immigration rather than sampling error, as it appears in half of the cemeteries from this period. This suggests that these cemeteries (OFU, TD, CSO, SPT) likely belong to a new wave of immigrants.

### Male lineage turnover in the Sarmatian Period

When examining the distribution of Y-chromosome haplogroups (Y-Hg) across different periods in the Carpathian Basin (Figure 7), we observe significant turnover in male lineages over time. The Neolithic turnover linked to the migration of Anatolian farmers^41–43^ and the Bronze Age turnover associated with the Yamnaya migrations^41,43,44^ are well-documented. However, during the Sarmatian period, we observe a new and previously unreported shift: the R1a~ haplogroups R-Z283 and R-Z93 appear in abundance and remain prevalent into the Hun period. Particularly notable is the subclade R1a1a1b2~ (R-Z93), which is characteristic of Middle-Late Bronze Age Steppe populations such as the Sintashta and Srubnaya cultures^31^, and their Iron Age descendants, the Eastern Scythians^27,32,45^. This haplogroup is also the most prevalent among the Steppe Sarmatians and Romanian Sarmatians, highlighting a direct link between all Sarmatian groups and reinforcing the conclusions drawn from the autosomal data.

**Figure 7:**
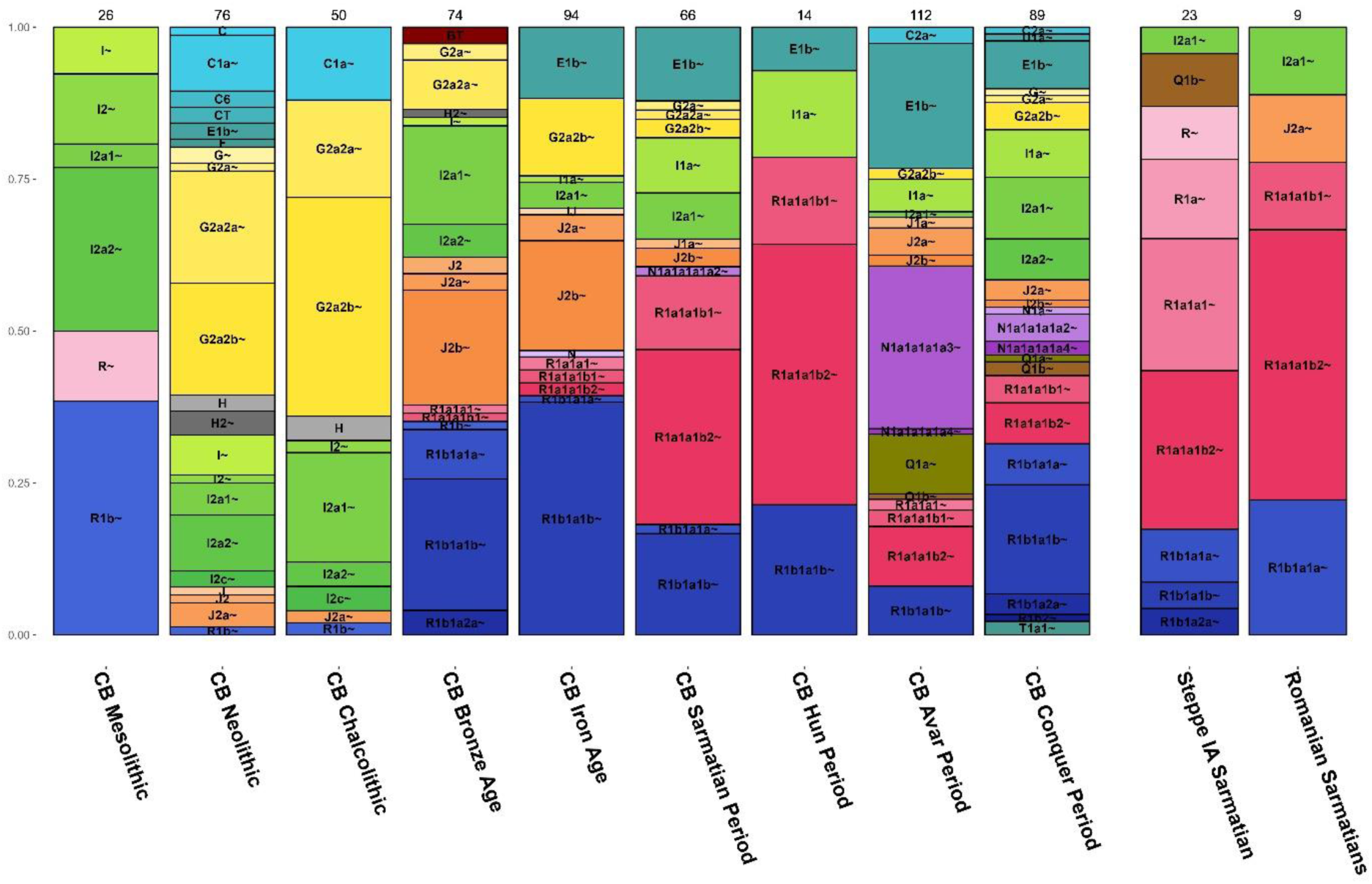
Y-haplogroup distribution of the progressive Carpathian Basin populations. Data were collected as described in Methods, further details are provided in Table S7.

Finally, the influx of Far Eastern Y-Hgs is observed during the Avar Period, with similar changes noted during the Conquest period as previously reported^46,47^.

## Discussion

### Relation to Steppe Sarmatians

To study the origin and genetic relations of CB Sarmatians, we have compiled the most representative database of this era and region to date. We have shown that the CB Sarmatians differed significantly from Steppe Sarmatians and more closely resembled the earlier local populations, except for their small but significant ANA component. In contrast, most Sarmatians from outside the Carpathians in Romania were more similar to the Steppe Sarmatians and appeared to form a genetic link between the two groups.

It is quite striking that the Romanian and CB Sarmatians shared negligible IBDs with the geographically closest and immediately preceding Western Scythians from Hungary, Moldova, and Ukraine. Instead, their immediate source populations, the Steppe Sarmatians, evidently derived from the more remote Ural and Kazakh regions.

We demonstrated that most studied Sarmatians require a Steppe Sarmatian source in their genome modelling, with this component gradually diminishing over time. This pattern suggests a founder effect from a single migration, indicating that Romanian Sarmatians are close descendants of Steppe Sarmatians who mixed with the local population after migrating to the Carpathian Basin. IBD data supports this, showing strong connections between Steppe, Romanian, and CB Sarmatians. Notably, the FKD-150 CB Sarmatian female shares 4 IBD segments totalling 48.5 cM with the DA139 Steppe Sarmatian female from the Pontic Steppe. DA139, in turn, shares 88 cM IBD with the chy001 Uralic Sarmatian and 64 cM with the POG-10 Romanian Sarmatian.

During the Sarmatian period, the male lineage composition in the Carpathian Basin underwent significant changes, highlighted by the rapid spread of R1a lineages. Notably, the Asian R1a-Z93 subclade gained prominence, clearly originating from the Steppe and Romanian Sarmatians, where this lineage is particularly common. It is worth noting that the available steppe Sarmatian genomes generally have low coverage, so a large proportion of those classified under the broader R1a1a1 haplogroup likely belong to the R1a1a1b2-Z93 subclade. In contrast, the maternal lineages did not undergo significant changes (Figure S4), which suggests that the westward migration of the Sarmatians may have been primarily driven by male participants. Nevertheless, it is noteworthy that in early Sarmatian period cemeteries, female burials, such as those in the ‘Golden Horizon’ graves, were especially prominent.

These findings align with historical accounts suggesting that the migrating Sarmatians initially targeted the Roman Empire along its northern borders at the Lower Danube.

The pronounced intragroup connectedness observed in the Steppe Sarmatians and other recently arrived groups from the steppe (HUN_AVAR_AC and HUN_CONQ_AC) suggests that the nomadic lifestyle is likely the primary factor responsible for the exceptionally high levels of intragroup IBD sharing. As the Sarmatians settled in the Carpathian Basin they transitioned to a more sedentary lifestyle, with agriculture becoming predominant. The subsequent decrease in intragroup connectedness in later Sarmatian groups is most likely attributed to this lifestyle shift and the increasing population size.

### Multiple new migration waves

Based on changes observed in Sarmatian archaeological material, archaeologists hypothesize multiple waves of Sarmatian migration, suggesting the arrival of new populations during the late 2nd and 4th centuries. Our findings support this, indicating that new groups likely arrived during both the early-middle and late Sarmatian periods, roughly aligning with the archaeological timeline.

In the EM period, the populations of the HVF and MIJ cemeteries show significant differences from both the Sarmatians and earlier local populations, and they share substantial IBD connections with Avar and Conqueror populations.

The HVF individuals show a genetic shift towards northern European populations, with qpAdm analysis confirming the presence of northern European genometypes in this cemetery. For example, HVF-4 and HVF-21 can be exclusively modelled from Scandinavian genomes, while HVF-8 forms a clade with the Poland Wielbark population. The local component of the remaining HVF individuals was typically modelled from Germany_EMedieval_Alemanic_SEurope^48^.

The MIJ individuals, on the other hand, appear to carry northern European genomes admixed with East Asian ones, with most of them significantly shifted towards Asia in PCA. Their local components were typically modeled from Germany_EMedieval_Alemanic_SEurope, while MIJ-1 and MIJ-3 also carry 25% and 20% Xiongnu/Hun-Elite ancestry, respectively. Additionally, MIJ-7 and HVF-2, women with Sarmatian genetic affinity, are closely related, sharing 6 IBD segments with a total length of 142 cM.

These findings suggest that during the Sarmatian_EM period, there may have been two distinct migration waves: one from northwestern Europe, possibly related to the Germanic tribes of the Marcomannic Wars, and another from the Eastern Steppe, consisting of a population of East Asian origin distinct from the Sarmatians.

A likely second wave of migration detected in the Late Sarmatian period is evident from individuals in the OFU, TD, CSO, and SPT cemeteries (Figure S3). These individuals have sparse IBD connections with the Sarmatians but show significant ties to the Avar period. On PCA, they align with the local European population, with three individuals showing a clear shift towards modern Southern Europeans. In qpAdm analysis, all these individuals exhibited the most significant affinity to sources related to the Roman Empire (e.g., Germany_Roman.SG, Italy_IA_Republic.SG, Austria_Ovilava_Roman.SG, Italy_Imperial.SG), with some local and Sarmatian admixture (Table S5e). Therefore, in the Late Sarmatian period, the new migration likely came from neighbouring Roman provinces rather than from the steppes.

It is important to note that, genetically, we cannot detect the possible new influx of groups with a similar composition to the first Sarmatian wave, especially if these migrations followed a stepping-stone pattern. For instance, while archaeological data suggests possible elite migrations from the East between the late 2nd century and early 3rd century^15^, these have not been detected genetically.

All analyses identified 5 outlier individuals with elevated ANA ancestry from the Sarmatian period, which cannot be attributed to Steppe Sarmatians. These individuals were excavated from the MIJ, MDH and NKL cemeteries. The two individuals from the MIJ site have already been discussed above. MDH-209, dating to the Middle-Late Sarmatian period, shares IBD segments with multiple Avar period samples, as well as with individuals from the Early Sarmatian, Hun, and Roman periods.

The NKL cemetery is divided archaeologically into two sections, one from the Sarmatian period and the other from the Hun period. However, all four Sarmatian NKL samples were grouped into the uncertain category (HUN_SARM_UP) because their radiocarbon dates were spread across an unrealistically wide time range. Despite this, the eastern components of the NKL-7, NKL-135, and NKL-157 outliers are consistently modeled from Xiongnu/Hun elite ancestry, and they share IBD segments almost exclusively with Avar samples, including Avar elites (Table S6c). These findings strongly suggest that these individuals are more likely associated with migrations during the Hun period, though they may have arrived somewhat earlier.

The Hun period samples are distinctly separated into two IBD-sharing clusters (see Figure S1). Nearly all of the newly sequenced HUN_HUN genomes carry local genome types and are associated with the Sarmatian cluster, including Steppe Sarmatians, with only marginal connections to Roman, Avar, and Conquest period individuals. In contrast, the previously published Hun era genomes^34^ contain significant Asian components and align with the Avar cluster, including a few Conqueror elites. These results clearly indicate that during the Hun period, most of the Carpathian Basin population represented the existing local population, which survived well into the Conquest Period, while new Hun-era immigrants with Asian roots were in the minority. This is entirely consistent with historical data, which indicates that Sarmatian cemeteries were used until the early 5th century, and settlements until the mid-5th century.

Particularly noteworthy is the genome of the ASZK-1 individual from the Hun Period, which forms a clade with Steppe Sarmatians in most qpAdm models, while also exhibiting some East Asian admixture in other valid models^34^. This solitary and rich Hun burial is well-dated and bears numerous parallels to similar finds in the Kazakh Steppe. The burial customs and the entirety of the findings suggest an Eastern individual from an Eastern environment, making it likely that this individual arrived with the Huns^49^. This genome suggests that the descendants of the Steppe Sarmatians were also present among the incoming Huns.

The IBD connections separate the populations of the three successive migration waves (Sarmatian, Avar and Hungarian Conquest) into three distinct clusters - in Figure 5, suggesting that the three migration waves are largely associated with different populations.

### Iron Age samples

The two Early Iron Age samples LMO-8 and RAM-7 from the Carpathian foothills were contemporary neighbours of the European Scythians, radiocarbon dated to 2,6-2,7 kyears BP and predating the first Steppe Sarmatians (Table S1c). The LMO-8 male had a Scythian-style bronze arrowhead within his ribcage, which may have caused his death, or perhaps he wore it as a necklace (Supplemental Data). Surprisingly, despite the large temporal and geographical distances, the ADMIXTURE patterns of LMO-8 and RAM-7 are identical to those of the Steppe Sarmatians. In qpAdm models, RAM-7 nearly forms a clade with the Steppe Sarmatians, while LMO-8 appears to be approximately 75-90% Steppe Sarmatian with about 10-25% East or Central Asian admixture. Additionally, the R1a1a1b2a2a~ (R1a-Z2124) Y-chromosomal haplogroup of LMO-8 aligns with the typical haplogroups found among the Steppe Sarmatians (RAM-7 being female).

Their potential connection to the Steppe Sarmatians is further supported by IBD data. LMO-8 shares IBDs with three Steppe Sarmatians, including one segment that is 22.5 cM long. Additionally, LMO-8 shares IBDs with a Central Saka and a Tien Shan Saka, as well as with FKD-150, an early CB Sarmatian. Similarly, RAM-7 shares IBD with two early Sarmatians from the Ural region and with a Scythian from Moldova.

These data demonstrate that the two Iron Age individuals can indeed be considered genetically Sarmatian. This suggests that migrations from the Ural region westward may have already occurred during the Early Iron Age, at least sporadically. It is worth noting that the age of these two individuals is much closer to the European appearance of the Cimmerians, and one well-covered genome identified as Cimmerian (MDA_IA_CIMM, Table S3a) does indeed show a similar ADMIXTURE pattern. Thus, it cannot be ruled out that their appearance may be connected to the “Cimmerian” migrations.

## Supporting information

Supplemental informations

## ACKNOWLEDGMENTS

We are grateful to our archaeologist colleagues Csilla Balogh, János Dani, Csilla Farkas, Péter Gróf, Gyöngyi Gulyás, Eszter Istvánovits, Mónika Merczi, Boglárka Mészáros, Margit Nagy, Katalin Ottományi, Ágota S. Perémi, Enea Sergiu, Kornél Sóskúti and Csaba Szalontai for providing archaological material. We are thankful to all the anthropologists who provided bone material for this study Ágota Buzár, Sándor Évinger, Ana Ștefan and János Rovó. This research was funded by grants from the National Research, Development and Innovation Office (TUDFO/5157-629 1/2019-ITM and TKP2020-NKA-23) to E.N. and grant from the Ministry of Culture and Innovation (MCI-670-19/2023/FÁFIN) to T.T. and E.N. This research was partially funded by the Competence Centre of the Life Sciences Cluster of the Centre of Excellence for Interdisciplinary Research, Development and Innovation of the University of Szeged to E.N., Z.M., and T.T. (the authors are members of the ‘Ancient and modern human genomics competence center’ research group). I.M. was supported by the János Bolyai Research Scholarship of the Hungarian Academy of Sciences (BO/00710/23/10).

## AUTHOR CONTRIBUTIONS

Conceptualization: T.T., O. S.^1^ (Oszkár Schütz), A.P.K., and B.T. Supervision: T.T. Project administration and funding acquisition: T.T and E.N. Data curation, software: Z.M., O. S.^1^ and E.NY. Formal analysis, validation, methodology: Z.M., O.S.^1^. Investigation: O.S.^1^, Z.M., A.G., P.K., B.K., K.M., G.I.V. Resources: B.NY.K., N.L., I.M., A.M., E.P., A.SZ., Z.T., D.W., G.W., R.CS.A., ZS.B., L.K., L.L.O., GY.P., G.P., D.P., A.S., A.D.S., O.S.^2^ (Olga Spekker), S.V. Visualization: O.S.^1^and Z.M. Writing – Original Draft: O. S.^1^, T.T., B.T., A.P.K., Z.M., B.NY.K., N.L., I.M., A.M., E.P., A.SZ., Z.T., D.W., G.W. Writing – Review & Editing: all authors contributed.

## SUPPLEMENTAL INFORMATION TITLES AND LEGENDS

### Supplemental Data

Investigating the reservoir effect

Reassessment of Steppe Sarmatians

Individual-level analysis of IBD sharing patterns

Intergroup IBD sharing patterns between cemeteries

IBD connection network of individual cemeteries

Uniparental data

Novel approach for IBD segment filtration of raw ancIBD output

Archaeological Background

**Figure S1:**
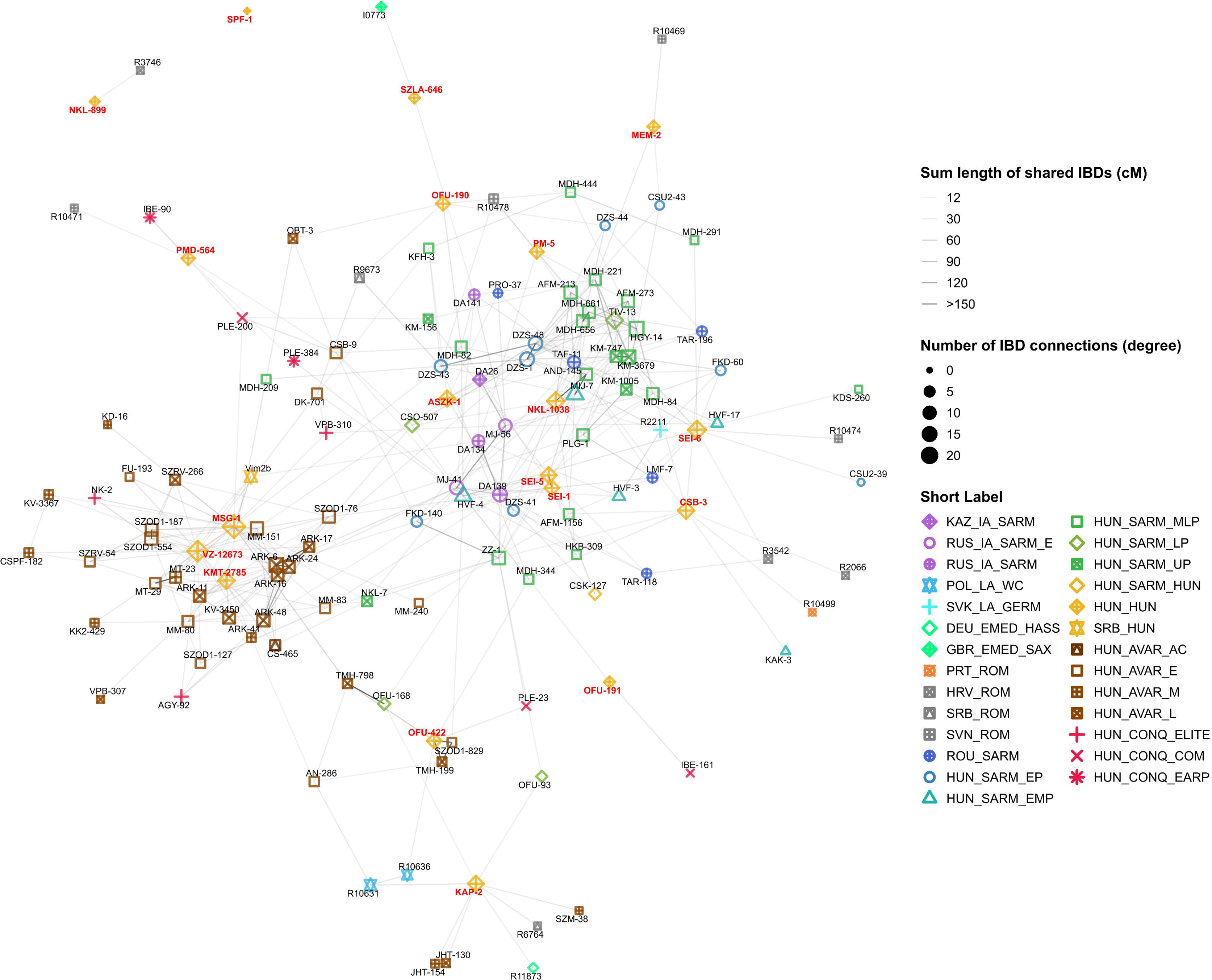
IBD sharing graph of the HUN_HUN individuals. All individuals from the HUN_HUN group are plotted alongside others with a minimum of 12cM sum IBD shared with any member of the group. The samples from the HUN_HUN group are highlighted in yellow, with red labels indicating their sample names. This plot was prepared in the same manner as the separate cemetery plots (see STAR Methods).

**Figure S2:**
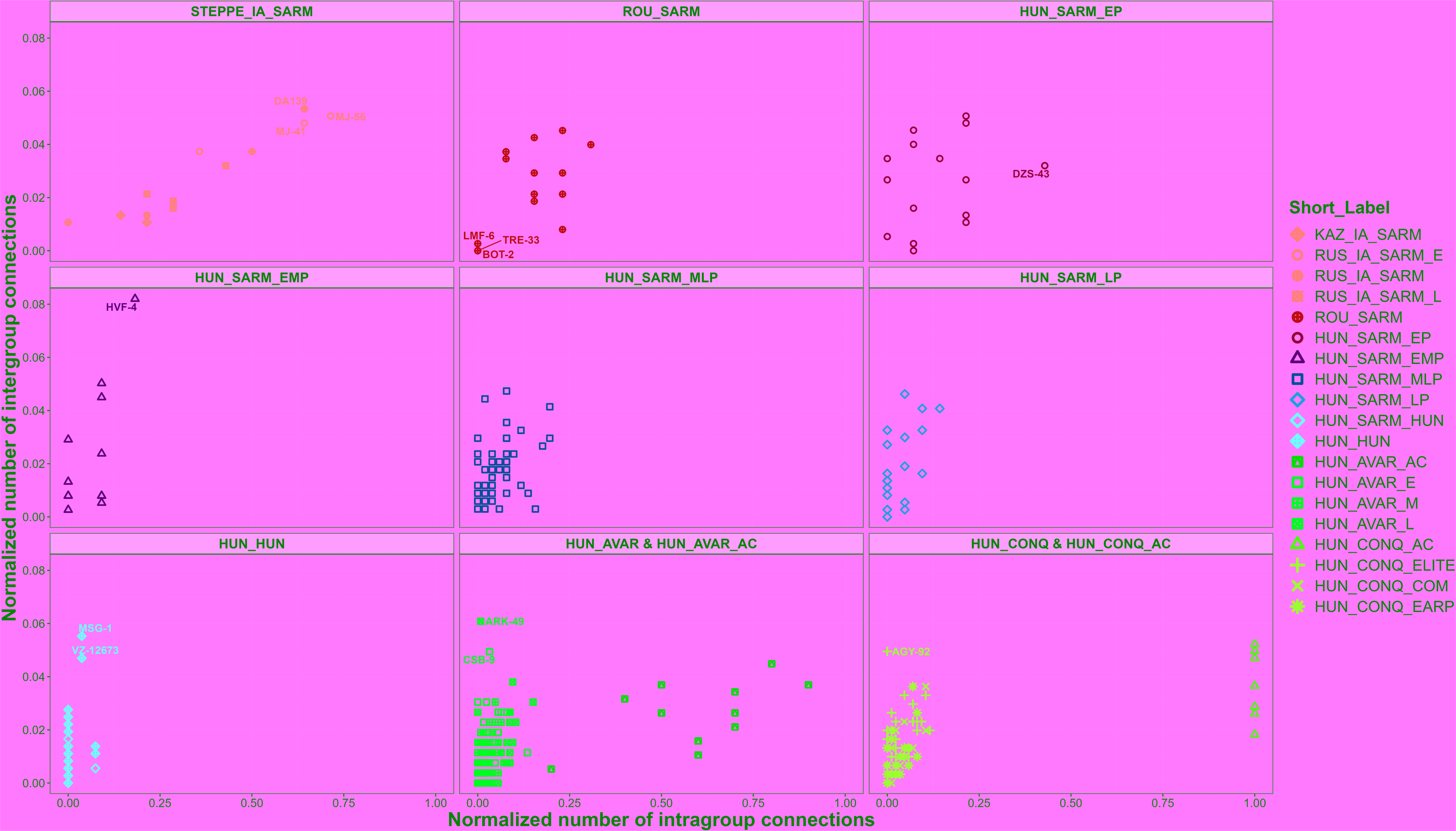
Normalized counts of intragroup and intergroup connections of individuals across various archaeological periods in the Carpathian Basin and its vicinity. Group names are written in the strip text above each subplot. For simplicity, the HUN_AVAR-HUN_AVAR_AC and HUN_CONQ-HUN_CONQ_AC groups are plotted together, although calculations were performed using the original groupings (see Column Label in Table S6b). The Y-axis represents the normalized number of intergroup connections (number of detected outgroup connections/total number of all individuals - group size]). The X-axis represents the normalized number of intragroup connections (number of detected intragroup connections/[group size – 1]). Each connection was considered equal, independent of IBD number and size.

**Figure S3:**
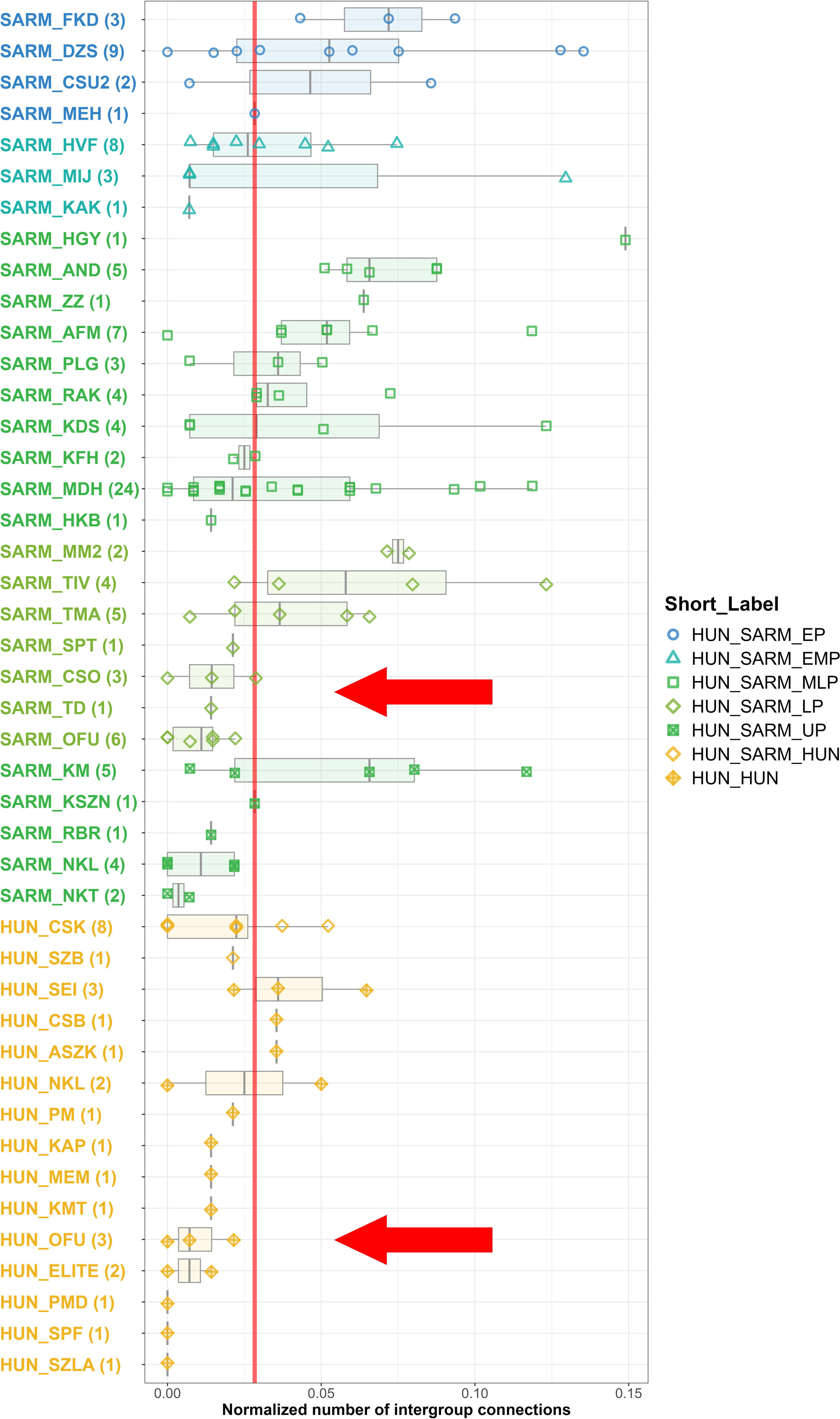
Intergroup sharing among cemeteries of the Sarmatian and Hun Periods of the Carpathian Basin. Box plots display the normalized number of intergroup IBD sharing among Sarmatian and Hun period cemeteries. Abbreviations for the cemeteries are listed in Table S6b. The red line indicates the median of the intergroup IBD connections across the plotted individuals.

**Figure S4:**
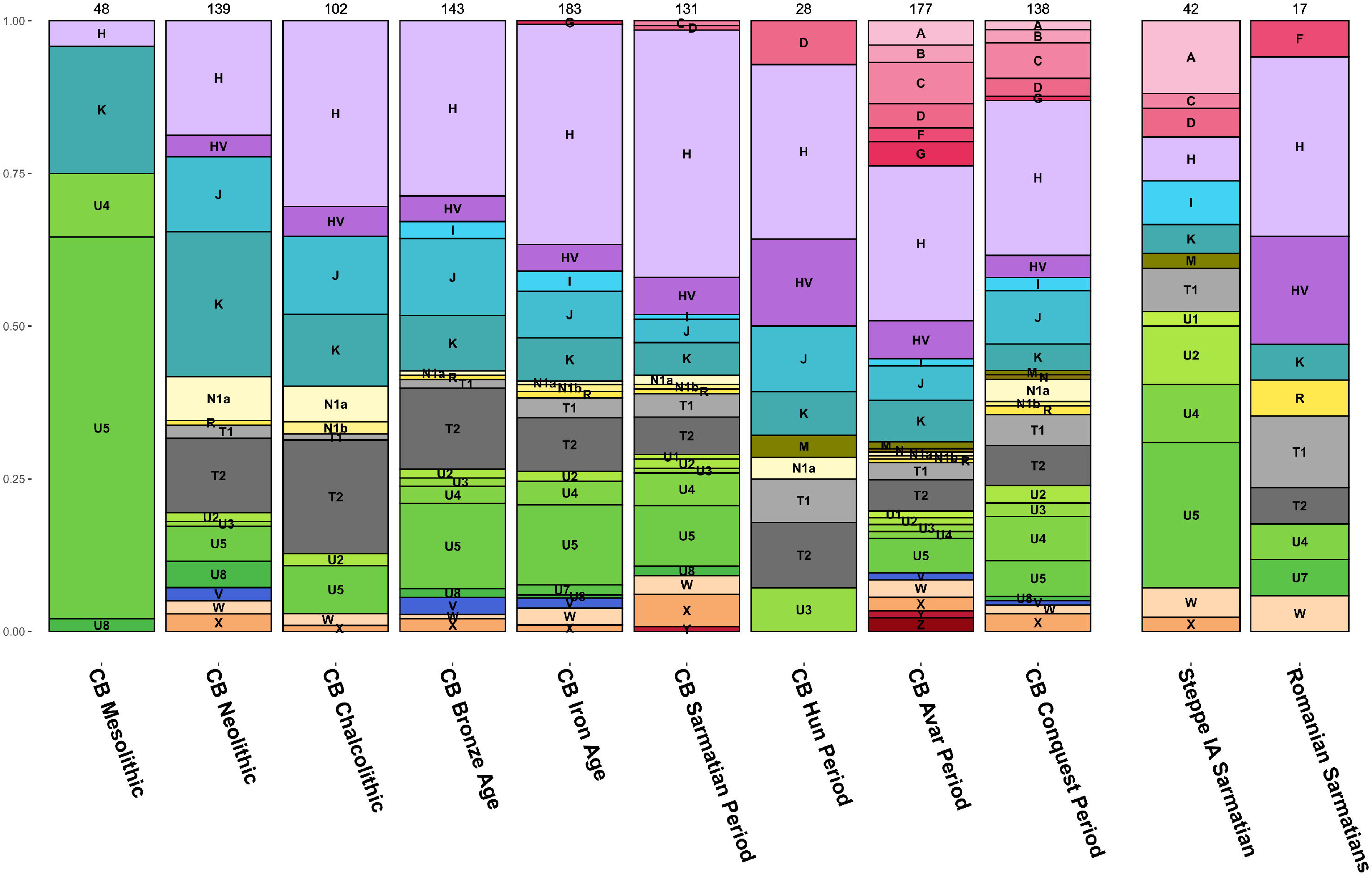
Mitochondrial haplogroup distribution across successive periods in Carpathian Basin populations.

### Table S1: Sample data

**Table_S1a:** Archaeological and genetic data of the newly sequenced samples from Romania and Hungary.

**Table_S1b:** Archaeological and genetic data of the previously published samples analysed in this study.

**Table_S1c:** Calibrated radiocarbon dates of newly sequenced samples.

**Table_S1d:** Kinship groups of the studied individuals identified by correctKin (Nyerki et al. 2023).

**Table_S1e:** Meta data of the unsuccessfully sampled individuals.

**Table_S1f:** Postmortem damage (PMD) patterns of the newly sequenced genomes.

### Table S2: PCA data

**Table_S2a:** Meta data and PCA coordinates of the modern individuals used as background in the PCA analysis.

**Table_S2b:** Meta data and PCA coordinates of the ancient individuals projected onto the axes defined by the modern background.

### Table S3: ADMIXTURE data

**Table_S3a:** Results of the ADMIXTURE analysis (K = 6). Ancestral components are named after representative sources. Group Order represents the hierarchical clustering results of centroids created from grouping the samples by the Short Label column. Individual Order represents a simple hierarchical clustering order if each individual.

**Table_S3b:** Cross-validation plot of different K values assembled from 30 parallel repetitions.

**Table_S3c:** ADMIXTURE plot of each group defined by Short Label. The order of the groups corresponds to Group Order. The groups studied in this article are indicated with a red X.

### Table S4: F4-statistics data

**Table_S4a:** Results of the F4 analysis in the form F4(POP1, POP2; POP3, POP4).

**Table_S4b:** Metadata of the individuals used in the F4 analysis.

### Table S5: qpAdm analyses data

**Table_S5a:** Summary table of the qpAdm analyses results. The p-value for the best model and the rank of that model is also included here for clarity. The data from Maróti et al. 2022 are not reanalyzed here, as they were thoroughly examined in that article using a very similar qpAdm framework.

**Table_S5b:** 2-Source qpAdm modelling of the studied individuals.

**Table_S5c:** 3-Source qpAdm modelling of the studied individuals.

**Table_S5d:** 2-Source qpAdm modelling of outlier individuals, who did not produce feasible models in the previous frameworks.

**Table_S5e:** Summary table of individuals and populations used as LEFT (Source) and RIGHT (Reference) populations.

### Table S6: IBD analyses data

**Table_S6a:** Results of the ancIBD software analysis (Ringbauer et al. 2024).

**Table_S6b:** Meta data of the individuals used in the IBD analyses.

**Table_S6c:** Adjacency matrix containing the pairewise total length of shared IBDs among each individual analysed. Empty cells represent no connections.

### Table S7: Y-chromosomal and mitogenome data

**Table_S7a:** Haploid information and meta data of the individuals studied in this article.

**Table_S7b:** Haploid information and meta data of ancient individuals from the Carpathian Basin based on the AADR repository (Mallick et al. 2024).

## STAR Methods

### Sample collection and DNA preparation

To uncover the genetic composition of the Sarmatian population living in the Carpathian Basin Barbaricum we collected bone samples from a wide spatio-temporal range (from the I-Vth c. CE). From the sampled 244 individuals we successfully generated whole genome shotgun sequences in 156 cases (Table S1a). The unsuccessful attempts were mainly the cause of low endogenous DNA content. To ensure transparency we publish the Master ID and cemetery label of the unsuccessfully sampled individuals in Table S1e.

All steps of sampling, DNA extraction and library preparation were carried out as described in^50^, in the dedicated ancient DNA laboratory of the Department of Genetics, University of Szeged. We used the minimally invasive extraction method described in ^51^, thus we primarily collected teeth samples, where it was feasible and their connection to other bones of the studied individual could be securely determined. In every other case we sampled the petrous bone. During the library preparation step, we used partial-UDG treatment to counteract extensive post-mortem damage. The double stranded libraries were than shallow sequenced on Illumina iSeq platform to monitor their human DNA content. Selected libraries were deep sequenced on Illumina NovaSeq platform to an average genome coverage of 1.42-fold (0.24x-3.75x, Table S1a).

### Data processing and quality control

The adapters of paired-end reads were trimmed with the Cutadapt software^52^, and sequences shorter than 25 nucleotides were removed. Read quality was assessed with FastQC^53^. The raw reads were aligned to GRCh37 (hs37d5) reference genome using the Burrows-Wheeler-Aligner (v 0.7.17) software, with the MEM command in paired mode, with default parameters and disabled reseeding^54^. Only properly paired primary alignments with ≥ 90% identity to reference were considered in all downstream analyses to remove high mapping quality exogenous DNA containing non-aligned overhangs. Samtools v1.1 was used for merging the sequences from different lanes and also for sorting, and indexing binary alignment map (BAM) files^55^. PCR duplicates were marked using Picard Tools MarkDuplicates v 2.21.3^56^. To randomly exclude overlapping portions of paired-end reads and to mitigate potential random pseudo haploidization bias, we applied the mergeReads task with the options “updateQuality mergingMethod=keepRandomRead” from the ATLAS package^57^. Single nucleotide polymorphisms (SNPs) were called using the ANGSD software package (version: 0.931-10-g09a0fc5)^58^ with the “-doHaploCall 1 -doCounts 1” options and restricting the genotyping with the “-sites” option to the genomic positions of the 1240K panel.

Ancient DNA damage patterns were assessed using MapDamage 2.0^59^ and read quality scores were modified with the Rescale option to account for post-mortem damage (Table S1f). Mitochondrial genome contamination was estimated using the Schmutzi algorithm^60^. Contamination for the male samples was also assessed by the ANGSD X chromosome contamination estimation method^61^, with the “-r X:5000000-154900000 -doCounts 1 -iCounts 1 -minMapQ 30 -minQ 20 -setMinDepth 2” options (Table S1a).

The raw nucleotide sequence data of the samples were deposited to the European Nucleotide Archive (http://www.ebi.ac.uk/ena) under accession number: PRJEB80732.

### Sex determination

Biological sex was determined with the method described in ^62^. Fragment length of paired-end data and average genome coverages (all, X, Y, mitochondrial) were assessed by the ATLAS software package^57^ using the BAMDiagnostics task. Detailed coverage distribution of autosomal, X, Y, mitochondrial chromosomes was calculated by the mosdepth software^63^ (Table S1a).

### Radiocarbon dating

Radiocarbon analysis was mainly performed on skeletal bone fragments of the sampled individuals to confirm the archaeological dating of the remains. When it was viable we instead used part of the remaining petrous powder to conduct the analysis, thus minimising the destruction of the human remains (Table S1a and c). The measurements were done by accelerator mass spectrometry (AMS) in the AMS laboratory of the Institute for Nuclear Research, Hungarian Academy of Sciences, Debrecen, Hungary (AMS Lab ID: DeA-37107; technical details concerning the sample preparation and measurement in ^64,65^). The conventional radiocarbon date was calibrated with the OxCal 4.4.4 software (https://c14.arch.ox.ac.uk/oxcal/OxCal.html, date of calibration: 20.02.2024) with IntCal 20 settings^66^.

### Haplogroup assignment and uniparental analysis

Mitochondrial haplogroups (Mt Hg) were determined using the HaploGrep 2 (version 2.1.25) software^67^, using the consensus endogen fasta files resulting from the Schmutzi Bayesian algorithm. The Y Hg assessment was performed with the Yleaf software tool^68^, updated with the ISOGG2020 Y tree dataset (Table S1a). In one case (MDH-405) the Y Hg could not be determined due to low coverage, this was marked as “inconclusive” in Table S1a.

To shed light on the most feasible origin of the uniparental lineages of our samples, we assembled a comprehensive uniparental database of the Carpathian Basin. We have chosen samples from the Allen Ancien DNA Resource (AADR)^69^ database based on their country of origin to cover this region. This included samples from Hungary, Slovakia, Romania, Serbia, Croatia, Slovenia and Austria. The haplogroups were collected from the original publications and cross-referenced with the publicly available databases published in ^70,71^ as well as our own classifications in the cases where the genomes were already downloaded for other analyses.

### Map

Maps were created in R 4.1.0^72^ with the help of “ggplot2”, “sf”, “ggspatial”, “rnaturalearth” and “elevatr” packages^73–77^. First, a data table was called containing the spatial coordinates for the selected geographical area using “rnaturalearth”. Waterways were added as a separate layer using the spatial data published in ^78^. Finally, we obtained elevation data with “elevatr” than superimposed it on the spatial coordinates as a “geom_tile” using “ggplot2”.

### Unsupervised ADMIXTURE

We used ADMIXTURE analysis to model our genomes as compositions of hypothetical ancestral populations^36^.

We performed unsupervised ADMIXTURE analysis on 2578 ancient individuals using the 1240K SNP set^69^. This included **149** newly sequenced individuals and **2429** ancient individuals from the AADR^69^ (Table S3a). As we did not include modern individuals from the HO dataset we obtained a significantly higher number (391,178) of overlapping SNP sights. We excluded individuals with less than 200K SNPs covered and samples with >4% contamination, we also taken out close relatives to avoid the appearance of undesired ancestral components. Cross-validation error calculation showed that modeling K-6 ancestral components yielded the most consistent results (Table S3b).

### Principal Component Analysis (PCA)

To uncover the underlying structure of our studied individuals in a hypothesis independent manner, we conducted PCA analysis. We used the same modern Eurasian genome dataset as our previous publication^34^ confined to the HO SNP set, to draw a modern PCA background on which ancient samples could be projected. This consisted of a generalized set of 1397 modern individuals from 179 modern Eurasian populations (Table S2a). PCA Eigen vectors were calculated from these pseudo-haploidized modern genomes with smartpca (EIGENSOFT version 7.2.1)^37^.

All ancient genomes were projected on the modern background with the ‘‘lsqproject: YES and inbreed: YES’’ options. Since the ancient samples were projected, we used a more relaxed genotying threshold (>50k genotyped markers) to exclude samples only where the results could be questionable due to the low coverage.

### qpAdm analysis

We used qpAdm^79^ from the ADMIXTOOLS software package^80^ for modelling our genomes as admixtures of two or three source populations and estimating ancestry proportions. The qpAdm analysis was done with the HO dataset, as in many cases suitable RIGHT or LEFT populations were only available in this dataset.

Our main goal of the study was to investigate the relationship between the Sarmatian individuals found in the Carpathian Basin and Sarmatians of the Central Steppe. Thus, in our qpAdm analysis framework we wanted to evaluate whether the available steppe Sarmatian individuals are a necessary source population for modelling the studied individuals. We considered three types of possible modelling sources (LEFT populations); a) a population set representing the supposed local inhabitants of the region, b) a population set representing the proposed Sarmatian ancestry, and c) a population set representing other possible Central and East Asian sources as our ADMIXTURE analyses indicated at least a marginal appearance of these.

The set of 15 local sources were selected from an extensive preliminary qpAdm run, among 162 possible candidates from the AADR^69^. The selection was based on population size, number of markers, PCA and ADMIXTURE clustering and goodness of fit in the acquired models. We assembled this optimal source population subset for the explicit purpose to model the highest number of our test subjects in a single qpAdm run and not to acquire their true ancestral composition. To represent the arriving Sarmatian population we assembled two genetically homogeneous source population from the available Steppe Sarmatians published from Russia and Kazakhstan^69^ (see Supplemental Data). The remaining sources represent other possible Central or Eastern Asian immigrants. We also used populations from our previously published article^34^, which had been shown to have extensive connections to the Carpathian Basin.

The reference population set (RIGHT populations) contained Ethiopia_4500BP (fixed), Iran_GanjDareh_N, Turkey_N, Latvia_HG, Baikal_EN (Russia_Shamanka_Eneolithic.SG and Russia_Lokomotiv_Eneolithic.SG), WSHG (Russia_Tyumen_HG and Russia_Sosnoviy_HG), Russia_Steppe_Maikop, Karitiana and Poland_Koszyce_GlobularAmphora.SG. For detailed list of LEFT and RIGHT populations see Table S5e.

During the runs we set the details:YES parameter to evaluate Z-scores for the goodness of the fit of the model (estimated with a Block Jackknife). As qpWave is integrated in qpAdm, the nested p values in the log files indicate the optimal rank of the model. This means that if p value for the nested model is above 0.05, the Rank-1 model should be considered^79^.

As our extensive source population set (LEFT populations) portended a great number of alternate models we applied the model competition framework explicitly discussed in ^31^ and ^34^. In this setup we test each resulting qpAdm model with a feasible p-value, by iteratively rerunning it with moving each of the LEFT populations in the RIGHT population set. This results in a bilateral improvement. On one hand the true source population – when included in the RIGHT population set – should consistently exclude any suboptimal models, as the test will (by design) have their highest shared drift with their true sources. On the other hand we will have a distribution of p-values for each resulting model which enables us to better quantify their goodness of fit, as demonstrated in ^79^. As we run each model multiple times, we can obtain further useful information concerning the feasibility of the individual models. Thus, based on the output of the qpAdm algorithm we included further quality measurements into our analysis framework. The meaning of the columns in Table S5b-d is as follows. **Test**: the name of the individual/population modelled. **SourceX**: the name of the designated sources in descending order of contribution. **SourceX ratio**: the obtained mixture coefficient for the source population/individual in question. **Valid models**: the number of cases a given model passed the quality criteria. **Excluded models**: the number of cases a model has been excluded by one of the reference populations. **BadFit models**: quality criterion, number of cases the obtained mixture coefficient differs from the jackknife calculated mixture coefficient by at least 10%. **Negative models**: quality criterion, number of cases where one of the source coefficients is lower than 0. **Non-significant nested p-value models**: the number of models where the obtained nested p-value is higher than 0.05. This indicates that the k-1 model is more plausible. **Average nested p-value**: the average of the obtained p-values for the k-1 model. **Minimum p-value**: the lowest p-value obtained from the model competitions. **Maximum p-value**: the highest p-value obtained from the model competitions. **Average p-value**: the average of the obtained p-values of the valid model repetitions (passing the quality criteria). **Average p-value summary**: either the average p-value or the average nested p-value if the number of non-significant nested p-value models is higher than the half of the valid models. This was used to order the qpAdm results. **Minimum p-value reference**: the name of the LEFT population/individual which was moved to the RIGHT population set when the lowest p-value was obtained for the model in question. **Maximum p-value reference**: the name of the LEFT population/individual which was moved to the RIGHT population set when the highest p-value was obtained for the model in question. **Excluding reference**: the name of the LEFT population/individual which was moved to the RIGHT population set when the model was excluded.

We ran comprehensive 2-way modelling runs for all studied TEST individuals with the above-described RIGHT population set and freely combined LEFT population set (Table S5b). Subsequent 3-way modelling was only conducted on a selected subset of the TEST individuals with unsatisfactory 2-way models (Table S5c). We selected these individuals based on preliminary qpAdm analyses where a sufficient increase in their p-value was reasonably expected. After 3-way modeling, 12 individuals still remained with no feasible models (p-value <0.05 or all models excluded by the model-competition). As the ADMXITURE and PCA profile of these individuals showed a very similar composition as outlier individuals in our previous article^34^, we included some further sources representing possible Northern and Southern European populations of the time as some of the outliers seemed clearly deriving a portion of their ancestry from these regions. We swapped our Steppe Sarmatians sources for some of our already modelled individuals and individual Russian and Kazakh Sarmatians, as they may containe some minor components that were not sufficiently represented in the grouped Sarmatian reference populations (Table S5d). In the end we successfully modelled all of our studied individuals except for a single individual (HVF-10) with high average p-values (Table S5a). The single unmodeled individual seems to be an outlier which has no sufficient source in the database yet.

### Imputation

We imputed our studied genomes together with other shotgun sequenced ancient genomes from a similar spatio-temporal distribution (Table S6) with the GLIMPSE2 framework (version 2.0.0)^38^ according to the recommendations of ^81^. Approximately 78 million biallelic common markers from the 1KG dataset were imputed with GLIMPSE2, utilizing the 1KG phase III data as a reference. The reference dataset was normalized, and multi-allelic sites were split using bcftools (version 1.16-63-gc021478 with htslib 1.16-24-ge88e343), applying the “norm –m –any” subcommand. Biallelic SNPs were filtered using the “view –m 2 –M 2 –v snps” subcommand. The autosomal chromosomes of the human reference genome were divided into 580 genomic chunks using the GLIMPSE2_chunk tool with the “-sequential” option. Following the GLIMPSE2 guidelines, we generated the binary reference data using the GLIMPSE2_split_reference tool, based on the 580 genomic regions and 1KG biallelic SNP variants. For the imputation process, we included only samples with shotgun WGS data exceeding 0.25x mean genome coverage and a contamination level below 0.04, as recommended in the GLIMPSE2 manuscript. Due to these criteria, we excluded 17 Sarmatian samples published in ^33^, as they were obtained through capture enrichment sequencing, along with 7 of our newly sequenced genomes that exceeded the contamination threshold.

### Kinship estimation and IBD sharing analysis

Kinship analysis was performed with correctKin^35^ (Table S1d). As reference population we applied the same database as in ^50^.

The shared IBD segments were identified using the ancIBD framework (version 0.5)^39^ according to the recommended workflow. Phased and imputed variants were post-filtered to include only the positions of the 1240K AADR marker set and lifted to the hdf5 data format. IBD fragments were identified applying the standard five-state HMM with the haploid_gl2 emission model, which is appropriate for the GLIMPSE2 posterior likelihoods, to ascertain the raw IBD segments >=8cM.

During the subsequent filtration of raw IBD segments, we deviated from the marker density threshold (≥ 220 SNPs/cM) used in the original ancIBD framework^39^. We found that applying this global threshold led to a significant number of false positive and false negative IBD segment identification. Instead, we implemented a novel method that uses marker informativity scores to dynamically mask and exclude genomic regions lacking sufficient power to detect true IBD segments. A detailed description of our algorithm, along with a comparison of the two methods, can be found in Supplemental Data. We developed this filtration method in Python for use with the raw IBD output generated by ancIBD. Our tool integrates seamlessly into the ancIBD framework and is available on GitHub [www.github.com/zmaroti/scoreFilterIBD].

Shared IBD network was generated in R 4.1.0^72^, with the application of packages ggplot2 3.4.2^77^, igraph^82^ and qgraph^83^.

### Plotting

To generate Figure 4A, we partitioned our dataset into wider spatio-temporal groups based on the archaeological and geographical data (Regional Group in Table S6b). We first contracted our whole IBD network into single points (vertices) representing each group, then calculated the distribution of these points using the Fruchterman-Reingold weight directed algorithm implemented in qgraph, where weights were the number of connections (edges) between each group. We centralized and expanded this distribution to create sufficient space, then we calculated graph distributions for each groups separately with the same weight directed algorithm, but now the weights were given as the sum total length of shared IBDs between each individual. We added the coordinates obtained from the single vertex calculation to each corresponding groups’ coordinates to arrange the separate group distributions according to the single vertex distribution. Finally we ran a new weight directed algorithm, with the recalculated coordinates as initial coordinates and with *niter = 1* and *max.delta = 0* parameters to not allow points deviating from their initial coordinates. We only plotted edges corresponding to intergroup connections to make the plot more straightforward. Intragroup connections are represented in this cause by the distance of the points from each other.

For Figure 4B we allowed the algorithm to run for 100 iterations and we used *max.delta = (0.1 + % between-group connection)* which allows for points to move along their edges but proportional to the number of intergroup connections. This approach emphasizes the net direction of intergroup attractions.

For Figure 5 we defined a core set of populations that we were primarily interested in. This included individuals with the Regional Group labels: Western_Steppe_Sarmatian, Carpathian_Basin_Sarmatian_Period and Carpathian_Basin_Hun_Period, which included the individuals published in this article, and Central_Steppe_Sarmatian, Carpathian_Basin_Avar_Period, Carpathian_Basin_Conquer_Period as main candidates for references (Table S6b). We included a further 17 individuals from other Regional Groups that had at least five connections to any of the core populations as to not inflate the plot with non-informative data. This partition was made to find an equilibrium between a strict constrain that also preserves only the most informative individuals. After defining the desired individual set, we ran a simple Fruchterman-Reingold weight directed algorithm with no constrains for 1000 iterations, where weights were given as the sum total length of IBD fragments shared between individuals.

An important metric for graph-based calculations is the number of connections an individual has (degree or degree centrality, *d*). When comparing degree centrality among groups or individuals it’s important to consider that a connection always exists between two endpoints, thus we carefully avoided counting individual connections multiple times. Another pitfall to consider is comparing groups of different sizes, where the sample size disproportionately affects the chance of finding IBD connections (linearly increasing sample sizes quadratically increase the possibility of uncovering connections) thus, to properly compare groups we must normalize the obtained degree centrality values by dividing them with the number of possibly available connection thus producing the ratio of fulfilled connections. For Figure 6 and Figure S2 the number of connections (degree centrality, *d*) was normalized by the product of the sizes of the two groups in case of intergroup connections (*d_i_’ = d_i_ / [n_i_ x n_j_]*), while intragroup connections were normalized by the equation: *d_i_’ = d_i_ / ([n_i_ x {n_i_-1}]/2)*, where *n*symbolizes the group size. To produce Figure S3, we normalized the degree centrality of the cemetery groups by dividing it with the product of the size of the cemetery groups (*n_i_*) and the number of all remaining individuals (*n-n_i_*): *d_i_’ = d_i_ / (n_i_ x [n-n_i_])*.

