## Supplemental informations for "Unveiling the Origins and Genetic Makeup of the ‘Forgotten People’: A Study of the Sarmatian-Period Population in the Carpathian Basin"

### SUPPLEMENTAL DATA

This document contains the following sections:


Investigating the reservoir effect
Reassessment of Steppe Sarmatians
Individual-level analysis of IBD sharing patterns
Intergroup IBD sharing patterns between cemeteries
IBD connection network of individual cemeteries
Uniparental data
Novel approach for IBD segment filtration of raw ancIBD output
Archaeological Background


#### INVESTIGATING THE RESERVOIR EFFECT

An extensive radiocarbon investigation was conducted on 68 samples to ensure accurate periodization, which was a key parameter for grouping samples in the subsequent genetic analysis. In most cases, the radiocarbon and traditional archaeological dates aligned, with only minor differences in their upper or lower range values. These cases were then assigned to specific phases such as early, early-middle, middle-late, and late Sarmatian phases, as well as the Hun period. However, five samples showed major discrepancies between the radiocarbon and archaeological dates, with the radiocarbon results suggesting significantly earlier dates (by more than one century). After ruling out issues with the archaeological dating and the possibility of sample misidentification, we suspected methodological issues with the radiocarbon dating. Specifically, the much earlier dates could be due to reservoir effects, such as the freshwater reservoir effect linked to a fish-based diet 1,2. Unfortunately, no background data—such as from domesticated animal bones—was available to help filter or quantify the impact of the reservoir effect. To address this, simulations were performed to estimate the potential impact of varying levels of freshwater-related diets (Extended Figure 1).

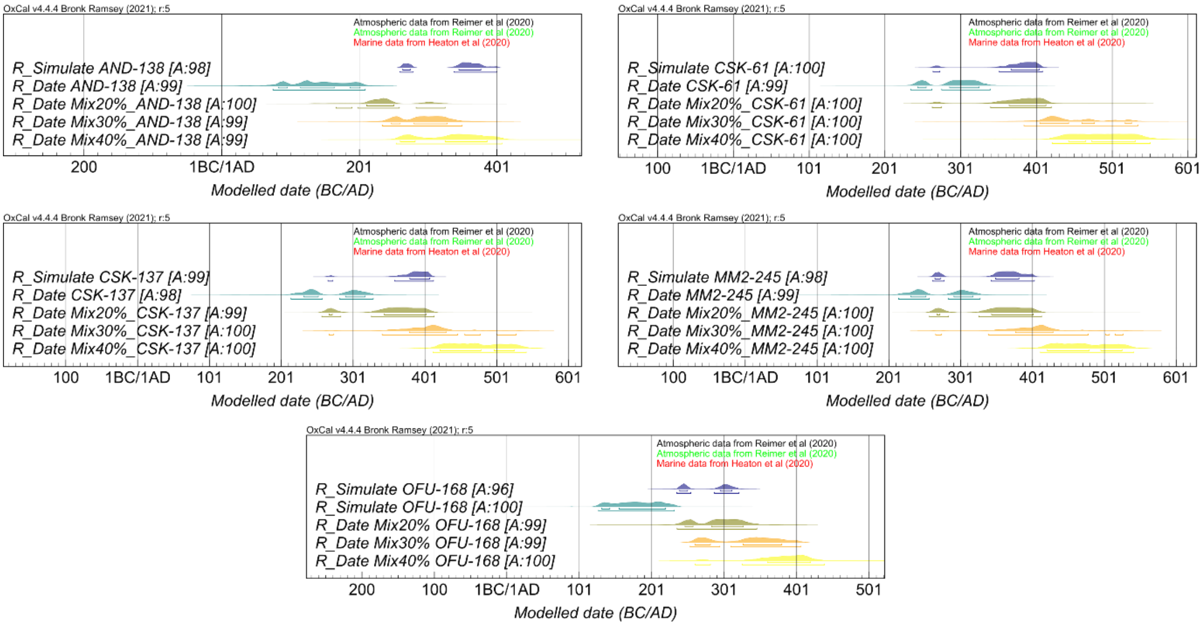


**Extended Figure 1: Simulations for the possible impact of reservoir effect on individuals with uncertain radiocarbon dating.**

These simulations indicated that fish consumption levels of 20–30% could account for the older radiocarbon dates, which is plausible given that all the samples came from the southern Great Hungarian Plain, near the Tisza and Maros rivers—key economic resources for local populations since ancient times. Consequently, in these cases, we prioritized the traditional archaeological dates over the radiocarbon dates. As a precaution, samples with radiocarbon results but lacking precise archaeological dating within the Sarmatian period were labeled as ‘unknown’ (HUN_SARM_UP) to avoid misclassification in the genetic analyses.

Reservoir effect was particularly evident in one individual (NKL-7), whose archaeological age was ambiguous. Radiocarbon dating placed NKL-7 in the early 1st to 2nd century CE (84-95 (3.3%); 116-216 (92.2%) calCE), however, this early date is strongly contradicted by genetic evidence. NKL-7 shares multiple 4th-degree relations with unpublished Early Avar Period individuals from Tiszavasvári-Kashalom dűlő (TKD) and Üllő (ULL), based on kinship coefficient estimates (Table S1d) and identity by descent (IBD) analysis, resulting in a discrepancy of approximately 300 years between the genetic and radiocarbon data. Therefore, the early radiocarbon date is likely due to a reservoir effect.

The same applies to two samples from the Óföldeák - Ürmös (OFU) cemetery (OFU-168, OFU-422). The radiocarbon date of OFU-168 was 127-232 CE and that of OFU-422 was 261-415 CE. Nevertheless, these individuals had multiple close familiar connections to Late Avars from the unpublished Tiszafüred – Majoroshalom (TMH) site (Table S1d). Therefore, if reservoir effect is considered, these individuals can be rather dated to the early Avar period.

#### REASSESSMENT OF STEPPE SARMATIANS

The Sarmatians were a nomadic people, who likely originated from the Central Steppe region (possibly from the Southern Urals) and during the 4th-2nd centuries BC they gradually encroached upon the territories formerly held by the Scythians [1].

Ancient authors designated multiple tribes under their name which is also corroborated by numerous archaeological findings all around the North-Caspian and Central Steppe regions. Thus, they likely exerted a strong cultural influence over vast geographic distances with an abundant populace (Extended Figure 2). Despite this significance, there are only a handful of articles in the literature referencing these people. The Allen Ancient DNA Resource [2] (v54.1) – which is quite up-to-date in this regard – only contains 45 definitive Sarmatian individuals from 7 different articles [3–9] Among these only two articles discuss the Sarmatians in context [4,6], while the others only marginally.

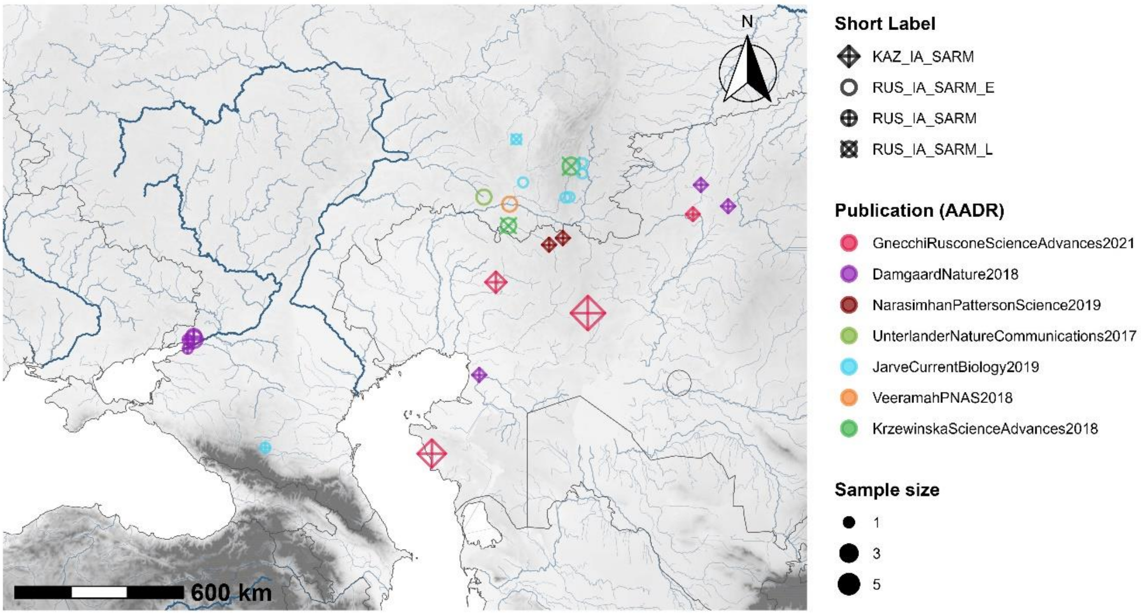

**Extended Figure 2: Geographical distribution of the Sarmatian samples excavated on the Steppes.**
Point colors represent the parent articles. A minor jitter effect was added to show off cemeteries with very close geographical locations. The point sizes represent sample size.

Since according to historical sources the Sarmatians of the Carpathian Basin are thought to be descendants of those from the eastern steppes, we aimed to reanalyze the published Steppe Sarmatian data, to create as homogeneous reference sets as possible for further analyses. We ran PCA, ADMIXTURE and qpAdm analyses to detect any previously unnoticed subtle differences in the published genomes. While the Sarmatian individuals from the steppe regions show strong clustering in the PCA, that persists across time, the ADMIXTURE analysis identified at least two marginally distinct clusters (Extended Figure 3, Table S3a). These findings were also supported by the qpAdm analysis (data not shown).

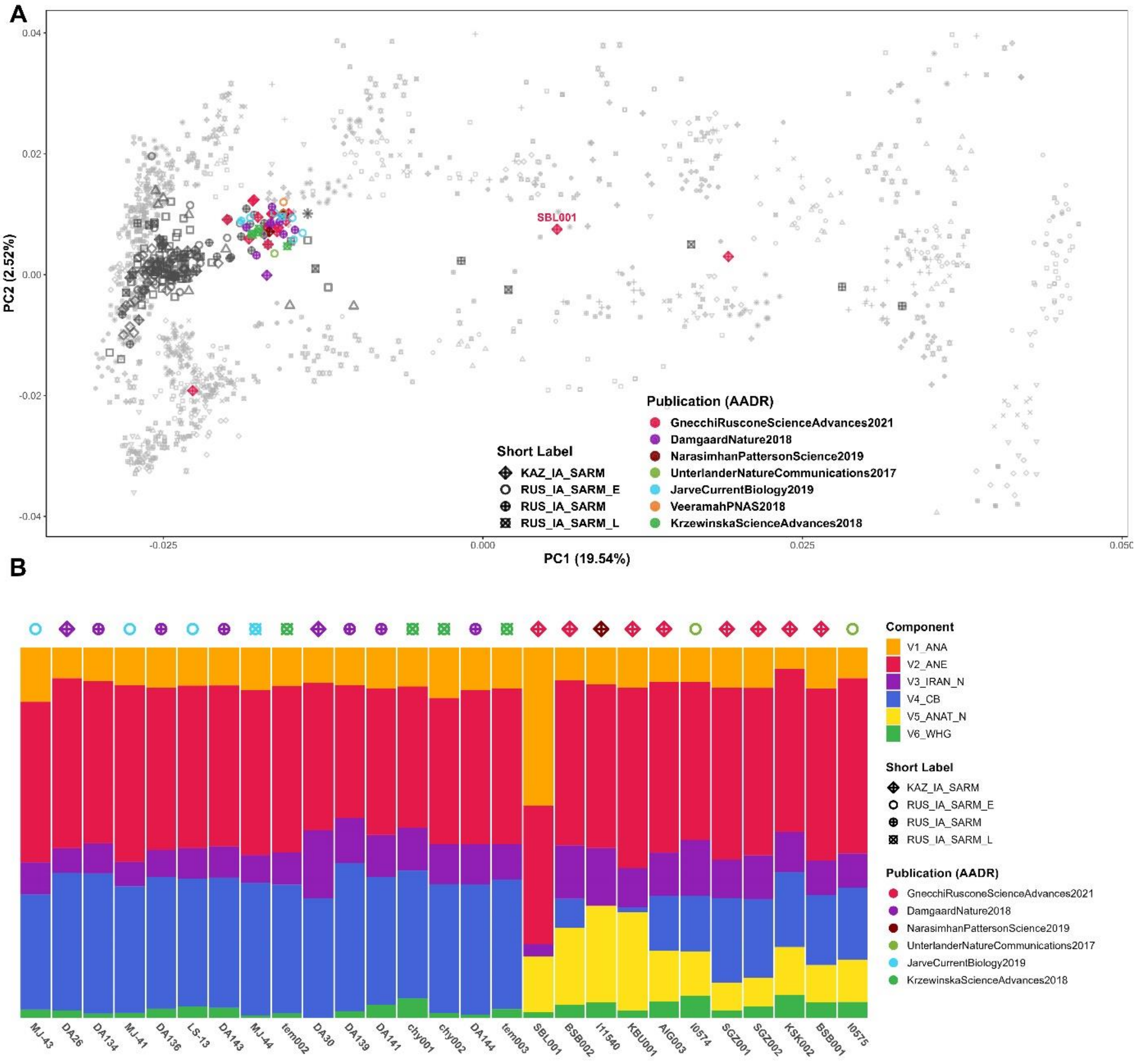


**Extended Figure 3: PCA and ADMIXTURE results of the Steppe Sarmatians.**
A) PCA analysis of the available Steppe Sarmatian genomes projected onto the same modern Eurasian population set as in Figure 2. Points are colored according to the parent publication. B) ADMIXTURE results of the Sarmatian samples with appropriate genome coverage distinguish two groups.

Unfortunately, many published individuals did not possess sufficient coverage to pass our stringent inclusion criteria for ADMIXTURE but they could be analysed with qpAdm (results not shown).

We finally constructed two genetically homogeneous groups formally titled STEPPE_IA_SARM_URAL (including chy001, DA134, DA139, DA141, DA143, DA144, DA145, DA26, DA30, MJ-41, MJ-44, tem001, tem002) and STEPPE_IA_SARM_STEPPE (including AIG001, AIG002, AIG006, BSB001, BSB003, CLK001, I0247, I11537, I11540, KBU001, KBU002, KSK002, MJ-39, Pr4). These groups were used as source populations in subsequent qpAdm modelling frameworks.

The PCA reveals several outlier individuals who are shifted toward Asia. Unfortunately, most of these had coverage too low for ADMIXTURE and further analyses, with the exception of one individual, SBL001.

Nevertheless, it is remarkable that, despite the broad region and timeframe considered, the overwhelming majority of the individuals cluster within a highly homogeneous genetic group. Moreover, the presence of outliers with presumably elevated Inner Asian ancestry aligns with our findings of similar outliers in the HUN_SARM_EMP and later groups. This suggests the possibility that distinct steppe groups may have accompanied the Sarmatian migrants into the Carpathian Basin.

We encourage further research on this topic, as the Sarmatians likely left a lasting genetic legacy among the later populations of the region.

### IBD ANALYSIS

#### Individual-level analysis of IBD sharing patterns

To better understand the underlying genealogical patterns producing Figure 6, we also visualized the normalized intragroup and intergroup sharing patterns on an individual scale (Figure S2). The first phenomenon that is readily apparent is the unusually high intragroup connectedness of the STEPPE_IA_SARM, HUN_AVAR_AC and HUN_CONQ_AC groups.

This pattern can also be observed in the ROU_SARM and HUN_SARM_EP groups for some extent, but it steadily declines over the subsequent periods. A common feature of these three particular groups is their nomadic lifestyle, which may explain this phenomenon. Nomadism increases mobility across large geographical distances, while an agricultural lifestyle generally localizes people, resulting in much sparser connections over relatively close distances. While this pattern could also be due to increasing population size or sampling bias, the elevated intragroup connectedness as a sign of nomadism must be considered, at least in the case of the STEPPE_IA_SARM group. These individuals were sampled from a very wide spatio-temporal range, yet still exhibit this pattern.

An interesting individual is HVF-4, a Northern European genetic outlier from the HUN_SARM_EMP period, who has the most connections among all individuals in the dataset. This person alone harbours genealogical connections to 36 different individuals (~1.74% of all possible connections among everyone), including multiple Avar and Conquest Period individuals, even some elite individuals from the Conquest Period. This prolific person alone well exemplifies the survival of descendants from the Sarmatian Period population up until the Conquest Period.

Once again, the HUN_HUN group shows minimal intragroup connections, while their intergroup connectivity is relatively high. This further suggests that this group is not genetically distinct, but rather separated primarily by time period. Particularly intriguing are two individuals with the highest intergroup connectedness, MSG-1 and VZ-12673, who are clear genetic outliers from Inner Asia and were buried during the Hunnic occupation with typical Hun period findings. They project numerous long IBD connections to later Avar elite individuals and some elite individuals from the Conquest period (AGY-92, NK-2, VPB-310, PLE-200), all of whom share an Inner Asian origin (see also Figure S1). The relational network of these two individuals strongly indicates the presence of Inner Asian migrants arriving in the Carpathian Basin during the Hunnic period.

#### Intergroup IBD sharing patterns between cemeteries

Plotting intergroup IBD sharing by cemetery (Figure S3) reveals a significant depletion of IBD connections in about half of the Late Sarmatian (HUN_SARM_LP) and Hun period (HUN_HUN) cemeteries (red arrows), compared to earlier periods. Several possible explanations could account for these results. The first is sampling bias, as many of the individuals with low connections are from solitary finds or represent their cemetery alone, suggesting that more connections might emerge with more comprehensive sampling. The second explanation could be an increasing population size, combined with sampling bias. A third possibility is that these cemeteries primarily represent new migrants with limited connections to the preceding local population.

In the case of the Hun Period, we have documented migrants with very strong Inner Asian genetic connections, suggesting that migration is the most plausible explanation, even if sampling bias is a factor. Besides, archaeologists detect considerable population decline in the Hun Period, so an increasing population size is an unlikely factor.

Migration is also the most likely explanation for the same phenomenon in the Late Sarmatian period, especially when considering the genomic composition of individuals with low IBD connections. These individuals display sparse IBD ties to the Sarmatians, but show significant connections to the Avar period, which was not considered in Figure S5. This is particularly evident in the Óföldeák–Ürmös (OFU) cemetery from the HUN_SARM_LP group, where three individuals (OFU-168, OFU-190, OFU-422) have close kinship relations with both published and unpublished Avar-period individuals (see Table S1d). Interestingly, these Avars were not Asian immigrants but originated from Europe, as most individuals from the HUN_SARM_LP group with low connections show a clear PCA shift toward modern Southern Europeans. In qpAdm analysis, these individuals exhibit the strongest affinities with sources related to the Roman Empire (e.g., Germany_Roman.SG, Italy_IA_Republic.SG, Austria_Ovilava_Roman.SG, Italy_Imperial.SG), along with some local and Sarmatian admixture (Table S5e). Therefore, in the Late Sarmatian period, the new migration likely originated from neighboring Roman provinces rather than the steppes.

The Hun period samples are distinctly separated into two IBD-sharing clusters in Figure S1. One cluster primarily consists of Avar-period samples (brown), including a few Conqueror elites. while the other is dominated by Sarmatian-period samples (green), including several Romanian (blue) and Steppe Sarmatians (purple), with only marginal connections to Roman, Avar, and Conquest period individuals. The Hun-period samples in the Avar cluster are all dominated by Asian genomic components, indicating they are recent immigrants from Asia during the Hun period. In contrast, the Hun-period samples in the Sarmatian cluster clearly descend from the local population of the Sarmatian period.

#### IBD connection network of individual cemeteries

Below, we present the full IBD connection network for one or two large, representative cemeteries from each period, offering insights into the broader local networks of each era. These graphs were generated by including individuals from the specific cemetery and supplementing them with all other individuals who share a minimum total IBD length of 12 cM with at least one cemetery member (with a single IBD length constrained to 8 cM). This approach was primarily used to create clear, interpretable plots, as plots of larger cemeteries could otherwise become overcrowded. This approach emphasizes closer genealogical connections, providing a more detailed understanding of the deeper structures within the cemeteries, which may complement the archaeological descriptions.

By examining specific cemeteries, we can gain a better understanding of the demographic landscape that the Carpathian Basin Sarmatians experienced over the centuries of their occupation.

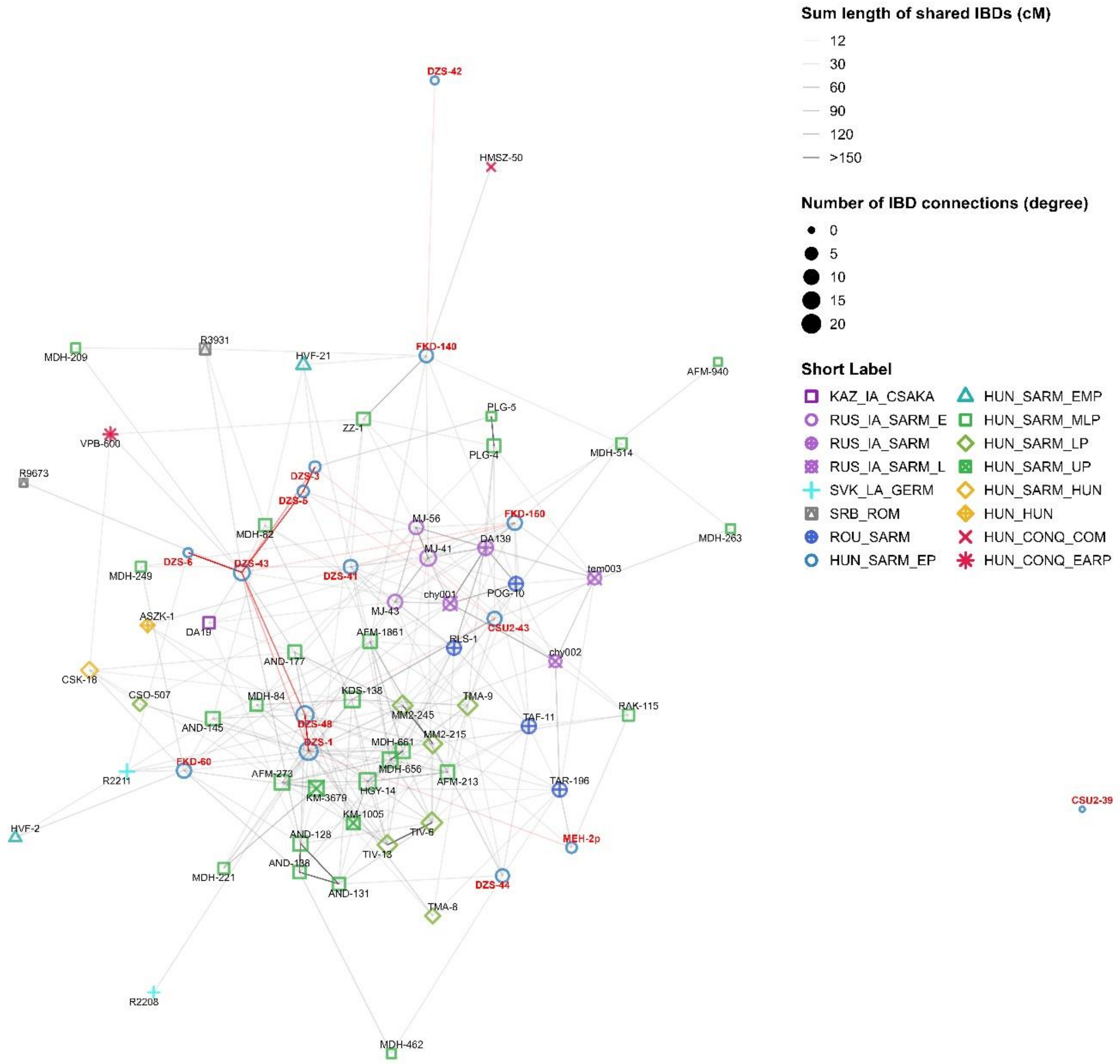


**Extended Figure 4: IBD graph of all individuals from the Early Period**

All HUN_SARM_EP individuals were plotted alongside those who shared a minimum total of 12 cM IBD with them. The HUN_SARM_EP individuals are highlighted with red labels, and their intragroup connections are indicated by red edges. Short labels in the legend are explained in Table S6b.

Extended Figure 4 displays the IBD connection network of the HUN_SARM_EP individuals. This period is mainly represented by two large cemeteries: Dormánd – Zsidótemető (DZS) and Füzesabony – Kastélydűlő (FKD). As shown in Figure 5, these cemeteries exhibit signs of a potential founder effect. It can be indeed observed that they share many connections with both Steppe Sarmatians and individuals from later periods. The DZS cemetery appears to represent a close-knit group, with nearly all sequenced individuals belonging to an extended family. In contrast, the three individuals from FKD display less internal connectivity, each showing distinct patterns. FKD-60 has numerous short connections with individuals from the HUN_SARM_EMP and HUN_SARM_MLP groups, and even some with a HUN_SARM_HUN individual from three centuries later. FKD-150, a genetically female individual, primarily shares IBD connections with Steppe Sarmatians from the Southern Ural region and individuals from the ROU_SARM group. Especially interesting is her connection with DA139 (another Steppe Sarmatian female from the Caspian Steppe), with whom she shares a total lengths of 48 cM IBDs in 4 fragments, indicating a closer than 10-degree genealogical connection. FKD-140 occupies an intermediate position, showing shared IBDs with both later HUN_SARM_MLP and Steppe Sarmatian individuals. These connections complement the qpAdm analysis results, suggesting that the HUN_SARM_EP individuals indeed descended from Steppe Sarmatians who arrived in the Carpathian Basin. Additionally, their numerous connections to individuals from subsequent periods support their role as a plausible source of Steppe Sarmatian ancestry detected in many of the studied individuals.

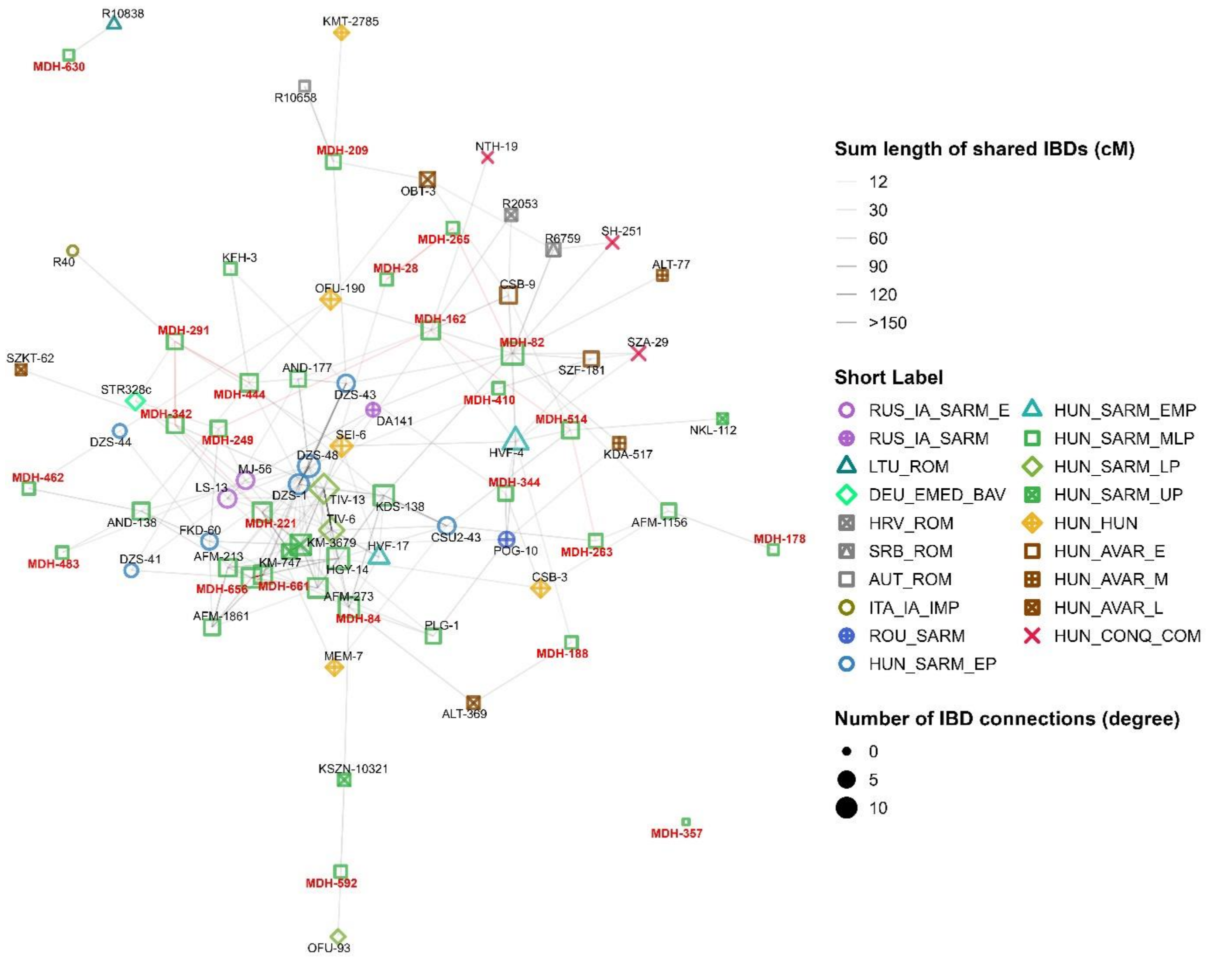


**Extended Figure 5: IBD graph of the individuals from the Madaras – Halmok (MDH) cemetery belonging to the Middle-Late period**

All SARM_MDH individuals were plotted (short labels in the legend are explained in Table S6b), highlighted with red labels indicating their identity and their intragroup connections are signified by red edges.

The largest cemetery we were able to sample is Madaras – Halmok (MDH), dating to the Middle-Late Sarmatian Period. The connections from this cemetery (Extended Figure 5) provide deeper insight into the population of that era. A key difference from the HUN_SARM_EP period is the sparse intra-cemetery connections. Out of a possible 276 connections, only 9 are present within the cemetery group. This could suggest that the cemetery was part of a larger metropolitan population center. Alternatively, sampling bias may have influenced this result, as only a small fraction of the individuals excavated were sampled due to poor preservation conditions.

Despite this, most outgoing IBD connections position these individuals among other Sarmatians in the Carpathian Basin, highlighting the interconnectedness of the population. Numerous genealogical links connect this cemetery to Dormánd – Zsidótemető (DZS) from the Early Period (14 connections) and to Apc – Farkas-major (AFM) and Tiszavalk (TIV) from the Middle and Late Periods (9 and 8 connections, respectively), although these sites are hundreds of kilometers to the north, near the foothills of the Northern Mountains in Hungary. These findings suggest that the community remained somewhat mobile across the Great Hungarian Plains, although their overall connectedness had significantly decreased compared to the previous period, as shown in Figure S1.

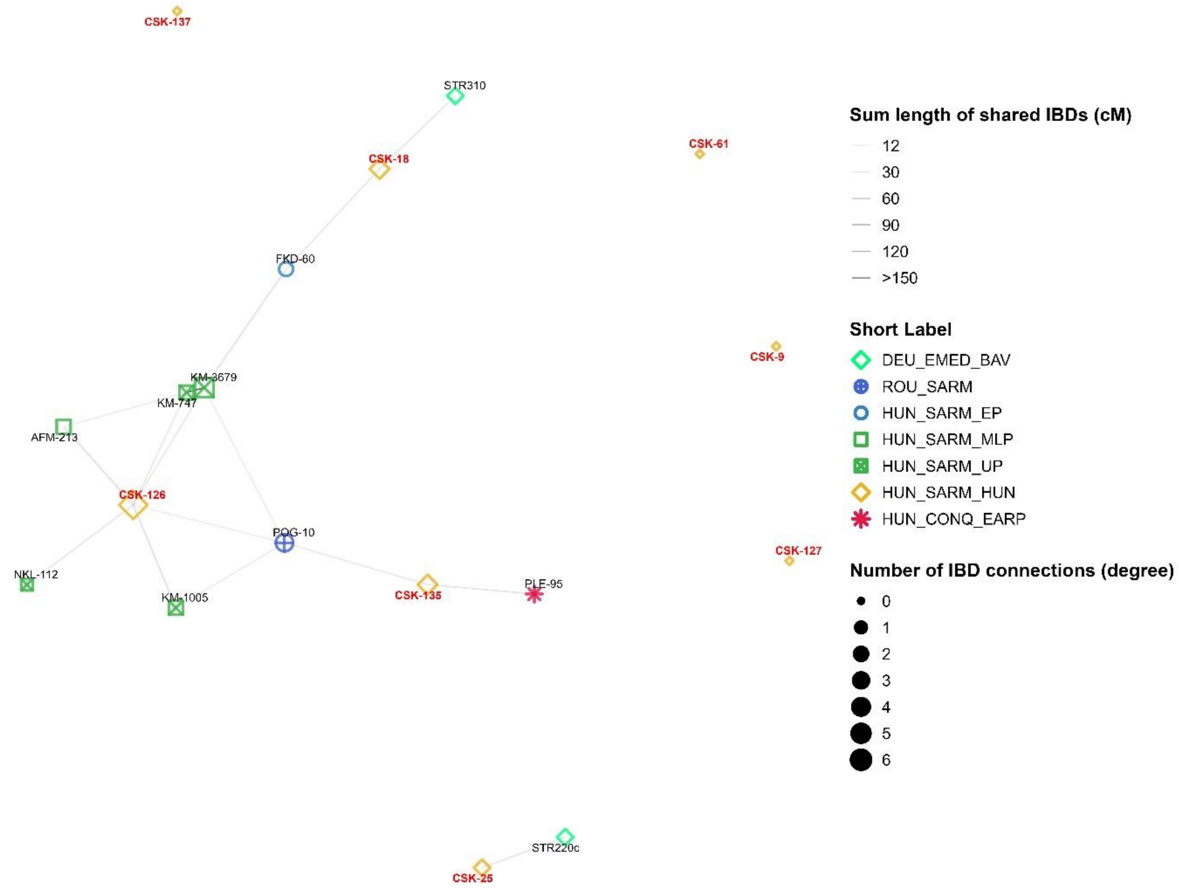


**Extended Figure 6: IBD graph of the individuals from Csongrád – Kenderföldek (CSK) from the Late Period**

HUN_CSK individuals are all plotted, with red labels indicating their identity, and their intragroup connections are represented by red edges. Short labels in the legend are explained in Table S6b.

Finally, we examined the transition between the Late Sarmatian and Hun periods at the turn of the 4th century through the cemetery of Csongrád – Kenderföldek (CSK, Extended Figure 6). This cemetery is unique in our dataset as its use began during the Late Sarmatian Period and continued through the Hun Period. Interestingly, we did not identify any IBD connections among the individuals in this cemetery. We observe that the individuals can be divided into two categories: some (such as CSK-18, CSK-126, and CSK-135) show clear connections to earlier Sarmatian periods, while others (like CSK-61 and CSK-137) lack any embedded relationships in the region.

This could indicate that a mixed community used this site, with some individuals having deep roots in the region and others newly arriving in the Carpathian Basin. Notably, individuals from Early Medieval Bavaria (STR220c and STR310) appear among the connected individuals, which is intriguing as their cemetery, Straubing – Bajuwarenstraße, has been linked to possible Sarmatian migrants (Source ??) during the Great Migration Period.

### UNIPARENTAL DATA

#### Uniparental database

Uniparental data, a specific form of IBD information, can provide valuable insights into population movements and familial relations. To leverage this, we assembled a comprehensive database of ancient individuals from the Carpathian Basin. The database was based on the AADR and included samples from countries within the Carpathian range, as well as Steppe Sarmatian samples for reference. These countries include Austria, Slovenia, Croatia, Serbia, Romania, Slovakia, Hungary, and Russia and Kazakhstan for the Steppe Sarmatians.

The uniparental data was collected from original publications, although some samples were excluded due to insufficient data. For the final haplogroup classifications, we consulted databases published in references [10,11], as well as our own classifications for genomes previously downloaded for IBD analysis (references [3,9,12]). We further excluded first-degree relatives, samples with a contamination rate higher than 10%, and low-coverage samples where mitochondrial haplogroups (Mt-Hg) could not be determined.

The final table includes 1,148 individuals, encompassing newly reported samples, spanning from the Mesolithic Period of the Carpathian Basin to the arrival of the Conquering Hungarians (Table S7). These individuals are divided into 512 females and 636 males.

#### Y-chromosome data

Y-chromosome data (Figure 7) indicates that subclades of R1a~ were only marginally present in the Carpathian Basin before the arrival of the Sarmatians. This haplogroup was detected in one Hungary_IA_Scythian (DA197), one Hungary_IA_LaTene_o3 (I25524), and four individuals from the Late Antiquity period (R2211, POP23, R6759, R9673), who were contemporaneous with the Sarmatians. The prevalence of R1a~ was significantly higher among Western Scythians from Ukraine, suggesting that this haplogroup was already present in the region around the Carpathian Basin during the Iron Age.

Notably, several Ukrainian Western Scythian individuals (MJ-14, MJ-15, MJ-33, MJ-34) with similar R1a~ subhaplogroups to the Steppe Sarmatians were identified as genetic outliers and exhibit very late radiocarbon dates (later than ~300 calBCE). This timing may suggest their direct connections to the Sarmatians, who were already present west of the Don River around this period (Instvánovics & Kulcsár 2017). Additionally, the very early Iron Age individual LMO-8 shares the R-Z93 haplogroup, further linking him to the Steppe Sarmatians, alongside IBD connections. Another Early Iron Age individual, MJ-31 (Ukraine_Cimmerians_o1, 1284-1055 calBCE), exhibits a similar haplogroup, R-Z645, which is on the same branch as R-Z93 but is a higher-level subclade.

While the most prevalent Y-haplogroup among the Steppe Sarmatians is R1a~ subclade R-Z93, there are many higher branches of the R1 haplotype reported among them. It is important to note that many of these haplotypes may indeed belong to R-Z93. This is due to the poor coverage of many available genomes from capture data, which often prevents precise terminal classification of their haplotypes.

The most prevalent haplogroup among our samples was the R1a~ subclade R1a1a1b2a~ (R-Z94), which was found in 4/9 Romanian Sarmatian males, 17/67 Sarmatian Period males, and 6/14 Hun Period males. This haplogroup can be further divided into subbranches R1a1a1b2a2a~ (R-Z2125) and R1a1a1b2a2b~ (R-Z2122). The subclade R-Z2125 is predominantly found among Romanian Sarmatians and Hun Period individuals, and is also present in several Steppe Sarmatians from Kazakhstan (I11540) and Russia (DA144), as well as in the Early Iron Age individual from Romania (LMO-8), the East Asian immigrant Hun MSG-1, several Avar Period individuals from Szarvas – Grexatéglagyár, and the Avar Period elite outlier DK-701.

The subclade R-Z2122 is more prevalent among the Carpathian Basin Sarmatians, appearing in Middle and Late Sarmatian Period cemeteries such as Madaras – Halmok (MDH), Apátfalva - Nagyút dűlő (AND), Apc - Farkas-major (AFM), and Kiskundorozsma – Subasa (KDS). These sites are also related according to the IBD analysis. Additionally, R-Z2122 is found in multiple Steppe Sarmatians from Russia (DA134, DA145, and Pr4), the Conquer Period elite, East Asian outlier K1-3286, a single Conquer Period individual from Árkus (ARK-14), and VPB-279, classified as Carolingian based on archaeological findings.

#### Mitochondrial data

The mitochondrial data (Figure S4) reveals that after the Neolithic transition, the mitochondrial haplogroup (Mt-Hg) composition in the Carpathian Basin remained relatively stable. This contrasts sharply with Y-chromosome frequency patterns, which indicate major population turnover events at the onset of the Bronze Age, the Sarmatian Period, and the Migration Period, linked to the Avar and Conquering Hungarian migrations.

During the Sarmatian Period, Y-chromosome data shows a significant turnover with the sudden appearance and expansion of the Asian R1a subclade R1a-Z93. However, this emergence of Asian Y-chromosomes is not mirrored in the mitochondrial haplogroup record, which closely resembles the preceding Iron Age pattern. A major influx of Asian Mt-haplogroups appears only in the later Avar Period and begins to decline during the Conquest Period. These results suggest that the westward migration of the Sarmatians was primarily driven by male participants. This aligns with the observation that Asian maternal haplogroups are present among Steppe Sarmatians at a moderate ratio but significantly drop among Romanian Sarmatians and become negligible among Carpathian Basin Sarmatians.

Nevertheless, most Steppe Sarmatians also carried mitochondrial haplogroups that were already present in Europe before their arrival (e.g., H, K, T1, U4, U5, W, and X). This allows for the possibility that both males and females might have migrated.

Comparing the maternal haplogroup distributions of Steppe Sarmatians and Romanian Sarmatians reveals substantial differences, more pronounced than their Y-chromosome distributions, while Romanian and Carpathian Basin Sarmatian maternal lineages do not differ radically. Therefore, a more plausible hypothesis is a male-driven spread of Sarmatians across the Pontic Steppe. However, we cannot definitively determine if this was also the case during their arrival in the Carpathian Basin.

### NOVEL APPROACH FOR IBD SEGMENT FILTRATION OF RAW ANCIBD OUTPUT

#### Background

The current implementation of SNP density filtration in the ancIBD framework is a solid general approach to eliminate the majority of false positive identity-by-descent (IBD) segments and retain most of the true positives from raw IBDs. On the one hand, the single global threshold allows easy control of the sensitivity and specificity; on the other hand, it does not account for the potential differences in the marker composition within and between different chromosomal regions and marker sets. Accordingly, for each marker set, it has to be fine-tuned and validated on synthetic data to achieve an optimal balance on sensitivity and specificity. In the ancIBD manuscript, the 220 SNP/cM threshold was validated for the 1240K Allen Ancient DNA Resource (AADR) marker set for the >8cM IBD segments, showing a good compromise for sensitivity and specificity. This can be also seen if we compare the number of raw IBD and the filtered IBD segments that meet the density criterion in a large cohort of samples (Extended Table 1).

| chr | length (cM) | marker count | raw IBD count | IBD count (density) |
|---|---|---|---|---|
| **1** | 284,26 | 89079 | 13950 | 1214 |
| **2** | 268,82 | 94167 | 1503 | 1166 |
| **3** | 223,26 | 77600 | 1040 | 952 |
| **4** | 214,2 | 68714 | 1131 | 896 |
| **5** | 204,05 | 69350 | 1048 | 932 |
| **6** | 191,72 | 75811 | 976 | 975 |
| **7** | 187,15 | 59833 | 908 | 746 |
| **8** | 168 | 61091 | 1156 | 836 |
| **9** | 166,14 | 50628 | 922 | 783 |
| **10** | 180,91 | 58760 | 1205 | 975 |
| **11** | 158,22 | 54795 | 797 | 781 |
| **12** | 174,59 | 53881 | 998 | 763 |
| **13** | 125,51 | 38991 | 79427 | 781 |
| **14** | 118,6 | 36279 | 782573 | 441 |
| **15** | 141,34 | 34407 | 29958 | 450 |
| **16** | 134,03 | 34372 | 758 | 530 |
| **17** | 128,5 | 29289 | 758 | 436 |
| **18** | 117,55 | 33899 | 631 | 468 |
| **19** | 107,73 | 18440 | 631 | 113 |
| **20** | 108,21 | 29053 | 678 | 463 |
| **21** | 63,64 | 16031 | 9950 | 286 |
| **22** | 72,44 | 15792 | 669362 | 159 |

**Extended Table 1: Per chromosome raw and filtered counts of > 8cM IBD segments in a large cohort of mixed Carpathian Basin and Sarmatian period individuals.**

In our study we co-analyzed approximatively 1400 individuals from numerous cemeteries where we had a considerable number of close relatives indicated by kinship analysis. Thus, it is expected that in such a large cohort, where true relatives and also ~1 million pairwise relations exist between mostly unrelated individuals, we will have a considerable number of true IBD segments and also a considerable number of false hits in the raw unfiltered data. As shown in Extended Table 1 the number of raw IBD in a large cohort of individuals has very large variations between the chromosomes, while the filtered IBD counts have much less variations, showing that the density-based raw IBD filtration overall has good specificity. The uneven raw IBD distribution indicates that the local SNP density must also greatly vary within and between chromosomes in the 1240K marker set.

However, the original manuscript also emphasizes that, in the case of smaller than 8 cM long IBD segments, in addition to the density criterion, it is recommended to apply masking of specific genomic locations to avoid an excessive number of false positive IBD segments. This suggests that while the density criterion is a good general approach for longer IBD segments there are other factors that also influence the accuracy of the IBD sharing analysis.

Despite the good specificity of true IBD filtration with the applied length and density criteria, the density criterion can also lead to unexpected behavior. For example, an IBD segment of 8cM with sufficient SNP density will be identified as a true IBD, however, the same IBD segment extending into a less densely (marker density) represented genome region could lead to it being filtered out, as the larger

IBD segment's mean SNP density could fall below the 220 SNP/cM threshold. This contradicts our expectation, since in case we have statistically enough markers to prove that two individuals share IBD, then having more markers that also comply with the shared IBD state, this should not be an exclusion criterion.

An extremity of the 1240K marker set is chromosome 19 where the overall SNP density is only ~171 SNP/cM for the whole chromosome, therefore it falls below the recommended 220 SNP/cM threshold. Accordingly, for all parents and offspring who by definition share their entire chr19 as a single IBD segment, this IBD segment will be removed by the density criterion. On the other hand, smaller IBD segments that fall into denser marker regions of the same chr19 where the local SNP density exceeds the 220 SNP/cM threshold can still be identified. Consequently, in this case, although the density method controls the specificity very well, it lacks sensitivity for genome regions with lower SNP density.

In theory, IBD segments are randomly distributed across the genome; therefore, the number of IBDs should correlate well with the genetic map length of the chromosomes. The considerable number of true IBD segments indicated by the density criterion in our experiment is sufficiently large that small stochastic variations should not significantly influence the expected distribution. To test and visualize our null hypothesis we plotted the number of filtered (true) IBD segments and the length of chromosomes (Extended Figure 7).

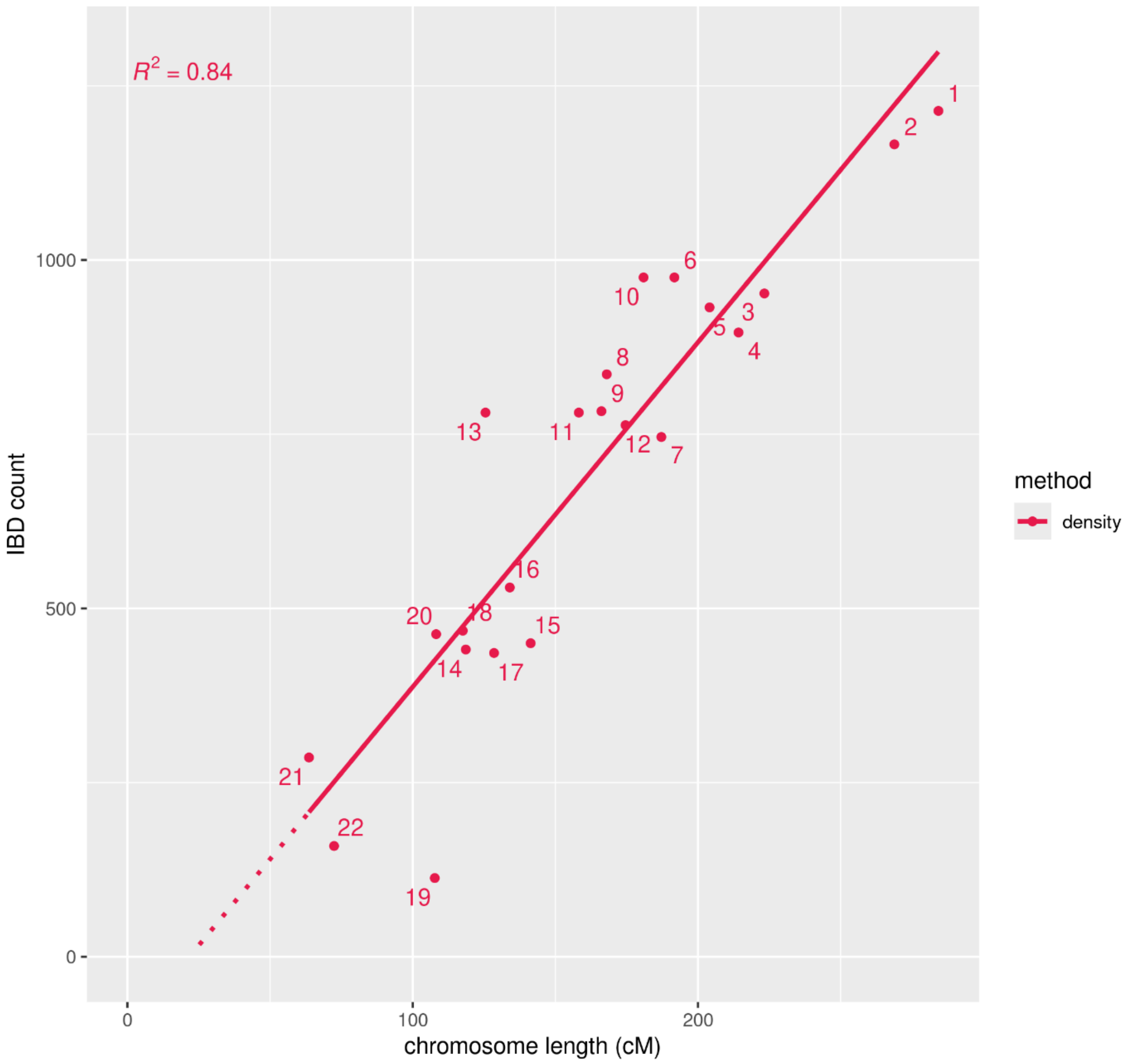

**Extended Figure 7: IBD count per chromosome indicated by the SNP density method in a large cohort of samples.**

Consistent with our null hypothesis, there is a significant correlation (R=0.84) between the observed count of indicated true IBD segments and chromosome length, suggesting that the density criterion is effective in the majority of cases. While the density filtration method and the 8cM length criterion removes majority of false positive IBD segments, the experimental IBD counts of the density-based filtration also illustrates some anomalies compared to the null hypothesis. Notably chromosomes such as chr13, chr10 and chr6 exhibits significantly higher than expected IBD counts likely due to excess of false positive IBD segments that were not excluded by the density criterion. In contrast there are also chromosomes that have less than expected IBD counts (chr19 is a prominent example), indicating a potential lack of sensitivity. The positive intercept of the linear fit along the x-axis further corroborates the suboptimal sensitivity, indicating that in some chromosomes an excess of true positive IBD segments were excluded.

#### Experimental distribution of the raw IBD segments within the genome

Assuming the null hypothesis that the true IBD segments are randomly dispersed within the genome, the IBD coverage defined as the number of true IBD segments crossing any genome position should follow a Poisson distribution. In practice, we only have sparse genotype information at the positions of the markers, and the IBD segments are defined as a subset of sequential markers with their unique genotypes. Accordingly, the experimental IBD count (coverage) at any marker position can be defined as the number of IBD segments that includes the particular marker. Due to the false positive IBD segments in the raw data the distribution of the experimental IBD count of the markers deviates from the Poisson distribution. However, according to our null hypothesis, the subset of markers present in the true IBD segments should exhibit a Poisson distribution for this metric. Our results show, that the majority of markers follow the expected Poisson distribution for IBD count (Extended Figure 8).

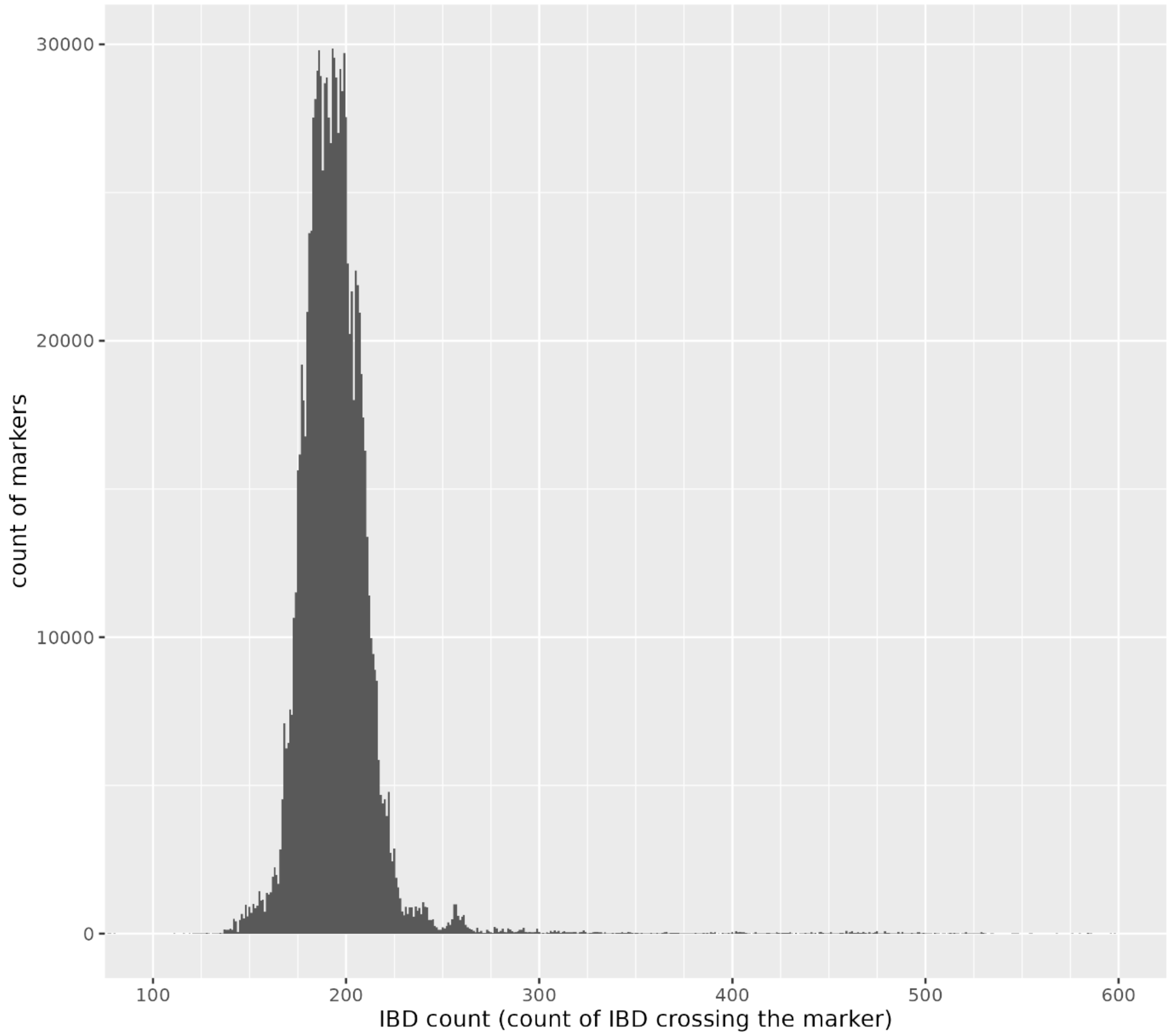


**Extended Figure 8: Histogram of the number of 1240K AADR markers included in a particular number of IBD segments (IBD count) based on experimental raw IBD data.**

A small fraction of the markers has higher than expected IBD counts. Since the very long right tail of the distribution is hard to visualize in Extended Figure 8 we also present the percentile table of IBD counts of the 1240K markers (Extended Table 2). The percentile data indicates, that less than ~3% of the 1240K marker set has IBD counts that are much higher than those expected from the Poisson distribution with a median of 193 IBD count. Only about 0.3% to 0.5% of the markers have several magnitudes higher IBD counts, ranging from about 100,000 to 800,000. These counts correspond to the case that 10% to 80% of all sample pair combinations share raw IBD segments at these particular locations, thereby accounting for the majority of the false positive raw IBD segments.

| percentile | IBD count |
|---:|---:|
| **1,0%** | 155 |
| **5,0%** | 171 |
| **10,0%** | 176 |
| **25,0%** | 184 |
| **50,0%** | 193 |
| **75,0%** | 204 |

| | |
|---|---|
| **90,0%** | 213 |
| **95,0%** | 222 |
| **96,0%** | 227 |
| **97,0%** | 240 |
| **98,0%** | 263 |
| **99,0%** | 1091 |
| **99,1%** | 2627 |
| **99,2%** | 6180 |
| **99,3%** | 9742 |
| **99,4%** | 11441 |
| **99,5%** | 29309 |
| **99,6%** | 79017 |
| **99,7%** | 79018 |
| **99,8%** | 660260 |
| **99,9%** | 777376 |

**Extended Table 2: The distribution of markers in the 1240K AADR marker set based on the metric of the count of IBDs that include the particular marker.**

The proportion and the distribution of the affected markers indicates that the problematic regions are confined to a small fraction of the genome where large fraction of unrelated samples seemingly share IBD with each other. These genomic regions mainly overlap with the mask regions described in the ancIBD manuscript (Extended Table 3-4).

| CHR | start (cM) | end (cM) | length (cM) |
|---|---|---|---|
| **1** | 147,421 | 163,079 | 15,658 |
| **2** | 0,014 | 26,182 | 26,168 |
| **4** | 0,341 | 11,535 | 11,194 |
| **8** | 0,0004 | 31,511 | 31,5106 |
| **10** | 62,682 | 77,491 | 14,809 |
| **14** | 111,711 | 120,2 | 8,489 |
| **15** | 0,005 | 43,484 | 43,479 |
| **17** | 54,307 | 63,702 | 9,395 |
| **18** | 0,1604 | 25,465 | 25,3046 |
| **19** | 0,002 | 29,543 | 29,541 |
| **19** | 74,533 | 107,7316 | 33,1986 |
| **21** | 0,861 | 21,041 | 20,18 |
| **22** | 1,723 | 23,842 | 22,119 |

**Extended Table 3: The "mask" regions according to the ancIBD manuscript**

| CHR | start (cM) | end (cM) | length (cM) | mean IBD count | SNP/cM |
|---|---|---|---|---|---|
| **1** | 147,119 | 164,242 | 17,123 | 3927,4 | 192,8 |
| **2** | 2,479 | 10,178 | 7,699 | 357,5 | 145,6 |
| **8** | 21,257 | 30,052 | 8,795 | 443,6 | 171,6 |
| **10** | 63,722 | 74,055 | 10,333 | 410,7 | 127,8 |
| **13** | 0,614 | 10,978 | 10,364 | 58760,7 | 224,3 |
| **14** | 2,477 | 16,408 | 13,931 | 470279,9 | 179,0 |
| **15** | 14,099 | 25,337 | 11,238 | 18001,4 | 103,8 |
| **21** | 1,896 | 12,078 | 10,182 | 7852,9 | 181,2 |
| **22** | 2,492 | 20,169 | 17,677 | 398523,2 | 135,5 |

**Extended Table 4: The “mask” regions based on the experimental IBD count (genome locations exceeding the median IBD count with greater than 6SD).**

According to experimental data, the majority of mask regions have low marker density in the 1240K marker set. In general, both the ancIBD mask track and our experimental data suggest that the problematic regions correspond mainly with the telomeric and centromeric regions of chromosomes. However, not all chromosomes are equally affected, and these regions also exhibit large variations in the extent of false positive IBDs and the spatial distribution of markers (Extended Figure 9).

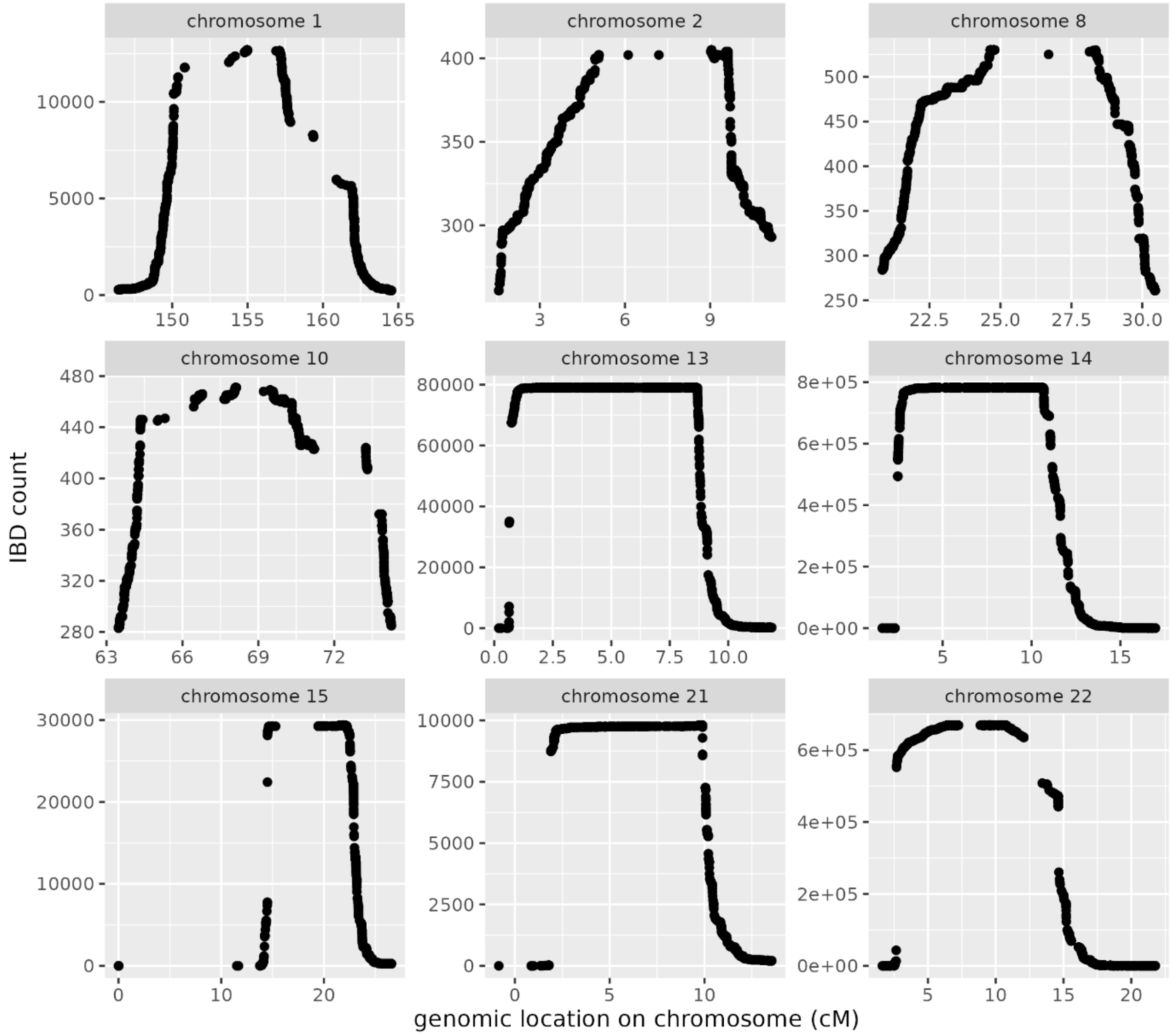


**Extended Figure 9: The experimental IBD count distributions at the low confidence genomic regions (mask regions).**

The higher incidence of low-confidence regions near the telomeric and centromeric regions is likely the result of multiple factors. In the telomeric and centromeric regions, the density of unique ancestry-informative markers is generally lower because of the higher fraction of repetitive sequences. In case of low-coverage ancient sequences, the genotypes that are the basis of IBD analysis are inferred by imputation. Consequently, the accuracy of the imputation also plays an important role. Since imputation uses the spatial genotype context to infer the most likely haplotype(s), and in these regions the sufficient length of flanking context is partially missing, it could also attribute to the lower accuracy in these regions. The observed patterns suggest that the differences between the distribution and the density of markers in the 1240K marker set, and also the imputation accuracy are mainly responsible for the observed differences between the accuracy of IBD analysis within and between the chromosomes.

Although the low-confidence regions suggested by our analysis overlap with the mask track provided in ancIBD, we also see some notable differences. According to the experimental IBD count data, approximately 80000 sample pair combinations share IBD on the telomere of chromosome 13 which is several magnitudes higher than the expected 193 IBD count. The excess of IBD count indicates a low confidence region despite the greater than 220 SNP/cM density that is not included in the ancIBD mask track. An additional difference is that the mask track of the ancIBD manuscript includes the centromere of chromosome 14 while the experimental data suggest that only in the telomere do we have much higher than expected IBD sharing compared to the null hypothesis (Extended Figure 9).

We have to note that in the ancIBD manuscript the locations of the mask regions were determined using the experimental data of a large cohort of Eurasian Ancestry individuals, while our data set included mainly Carpathian basin individuals and Sarmatian related individuals out of the Carpathian basin. In theory masking is only recommended below 8cM IBD length, thus we expect that the ancIBD mask track would include additional low-confidence regions where the specificity of density filtration drops. However, the additional low confidence region even at 8cM and the observed differences between the optimal mask areas of the two sample cohorts suggest that the uncertainty of IBD detection could also depend on the genome structure of the analyzed samples.

Although the global marker density in the panel largely correlates with the density of the actual markers informative for the analyzed cohort, in a smaller fraction of genomic regions where markers do not represent all populations equally, differences in the analyzed populations could influence the distinguishing power of the density-based single global threshold filtration method. Consequently, the optimal mask track could be somewhat different based on the populations being analyzed, as we expect that these false positive IBD areas correlate with the actual marker informativity representing the analyzed populations in the given region. For example, if our marker set in a specific genomic region mainly contains markers that have genotype, and haplotype variability in only African people, then this region will be appropriate for analyzing individuals with African ancestry. However, despite of the overall SNP density in this region, individuals of non-African ancestry will have a smaller genotype and haplotype variability, leading to a greater chance of sharing an identity-by-state (IBS) with other individuals of non-AFR ancestry. This is especially true as we decrease the length of the analyzed IBD segments. Furthermore, this also implies that for different marker sets (with their unique marker composition and distribution), the problematic regions could be different and the mask track has to be optimized individually for the optimal sensitivity and specificity.

#### Marker informativity based filtration

To address the aforementioned issues, we propose a new approach for filtration based on the experimental distribution of raw IBD segments. According to our null hypothesis, the IBD segments confined to these problematic genomic regions with unexpectedly high raw IBD counts are likely false. Consequently, the distribution of the experimental raw IBD segments can be used to indicate the 'mask areas' specific for the analyzed cohort. We defined IBD count of a marker as the number of IBD segments a particular marker is included. As we expect Poisson distribution for the true IBD segments, the ratio of the expected (true) and observed raw IBD counts (true + false) of each marker reflects the likelihood that the marker indicates true IBD segments. Based on this, we can calculate a marker informativity for each marker by the following formula:

$$M_I(i) = \frac{\boldsymbol{med}(I_c)}{I_c\,(i)}$$

Where $M_I(i)$ is the marker informativity of marker *i*, $\boldsymbol{med}(I_c)$ is the median of all marker IBD counts and $I_c(i)$ is the IBD count of marker *i*.

Since each IBD segment includes multiple markers, we can also calculate the sum of marker informativity a metric for each IBD segments that correlates with likelihood that the particular IBD segment represents a true IBD by the following formula:

$$I_s(j) = \sum_{i \in M(j)} M_I(i)$$

Where $I_s(j)$ is the IBD informativity score of IBD segment $j$ and $M(j)$ denotes all markers included in IBD segment $j$. Similarly, to the density method, a single threshold can be used to test against the calculated probability of all IBD segment to assess whether the raw IBD segment is likely false or true positive. However, unlike the global density threshold, the proposed metric is based on empirical probabilities that indicate whether the underlying markers within the IBD segment represent a true IBD segment. Consequently, this approach is less susceptible to the limitations posed by specific marker sets or population-specific regions that exhibit low accuracy in imputation and/or IBD analysis, leading to better sensitivity and specificity of IBD filtration.

The global 220 SNP/cM density threshold validated on the 1240K AADR marker set for IBD segments larger than 8cM corresponds to the IBD informativity score of $8 \times 220$. Therefore, the evaluation of raw IBD segments with the score criterion in the 1240K marker set, could be done with the following test.

$$I_s(j) = \begin{cases} > 8 \times 220, & true\ IBD \\ \leq 8 \times 220, & false\ IBD \end{cases}$$

This threshold would result in a very similar performance to density-based method for the majority of genome regions (approximatively 97% of the markers) where the raw IBD counts follow the Poisson distribution. This is due to the fact that in case of a greater than 8cM IBD segment with density of greater than 220 SNP/cM, when the underlying markers are within a genomic region with a median IBD coverage it is granted that the IBD informativity score will be greater than 8×220 and therefore the IBD will be classified true just like with the density model.

As highlighted by the ancIBD manuscript, the specificity and accurate measurement of the lengths of the IBD segments are vital in IBD analysis to mitigate the risk of drawing incorrect conclusions based on invalid IBD segments or misinterpreted close relationships. Accordingly, the implementation of our method consists of two consecutive steps.

The first step involves calculating the experimental IBD informativity scores from the raw IBD distribution. Our formula effectively reduces the scores of IBD segments that fall within low-confidence areas and improves the exclusion of false positive raw IBDs leading to better specificity. In the subsequent step, we also test whether the ends of the identified IBDs extend into a low marker-informativity "mask" area. In such a case, we truncate the end of the IBD extending into this low-confidence region where the markers are likely inaccurate, and only include the IBD in the final results in case the remaining high-confidence part of the IBD is still longer than applied length threshold. This second step ensures that IBD lengths are not over-estimated at low confidence genomic regions.

The notable differences against the density models are the following:

- If a large portion of the markers falls within a region where the empiric IBD count is high indicating a likely false positive "mask region", then these markers will have <<1 marker informativity and the IBD informativity score can go below the 1760 (8×220) threshold signaling, that even in case the global marker density in the region is higher than 220 SNP/cM, many of these markers are likely not indicating a true IBD segment. Furthermore, the end of IBDs extending into low-confidence regions are truncated, and the IBD segment is only kept, if the remaining high-confidence part is still larger than the length threshold. Hence, no manual masking is required while we expect improved specificity and a conservative length estimate around the low-confidence regions.

- The other notable difference is that the density method may drop out IBDs spanning larger genomic segments that have a general low density of SNPs, while the IBD informativity score method can still indicate these as valid IBD if sufficiently large number of markers indicates that the IBD is shared between the two individuals and these makers do not fall within the problematic "mask" regions with low probability of true IBD. Consequently, the score method will not throw out an 8cM IBD segment if it extends into a lower density but non-problematic genomic region, and it can also identify IBDs at generally low marker density areas if the supporting marker informativity is sufficiently high. Hence, we expect an improved sensitivity.

#### Comparison of the density and marker informativity methods

To assess the feasibility of the proposed method, we used a large cohort of samples from the Carpathian basin (approximatively 1300 individuals, 50+ cemeteries) and approximately 100 publicly available samples (that are source of some migrations into the Carpathian basin from the same period but outside of the Carpathian basin). Our data contained a large number of proven true relatives including first, second, and more distant relatives, thus we expected considerable amount of true IBDs in the cohort, while also most of the sample pair combinations should be between individuals that are unrelated, providing us enough false positive raw IBD segments to filter out. We included only shotgun WGS data for all of the samples (no capture data). We excluded individuals with lower than 0.5x mean genome coverage or greater than 4% contamination suggested by ANGSD X contamination or Schmutzi mtHG contamination analysis. The common markers of 1KG Phase 3 data (~78 million positions) were imputed by GLIMPSE2 using the phased genotypes of the 1KG phase 3 individuals as reference.

We used the official ancIBD workflow, to restrict sites to the 1240K marker set with the 8cM length threshold for IBD identification. Using the *hapBLOCK_chroms()* function, ancIBD indicated a large number of raw IBD segments (~**1.6M** > 8cM raw IBD segments in all chromosomes). We used the official 220 SNP/cM density method and the proposed IBD informativity score method to filter the raw IBD segments and compare the results. As expected, both methods filtered out most of the false positive IBD segments, while retaining a very similar number of true IBDs (Extended Table 5).

| method | IBD count |
|---|---|
| density | 15146 |
| score | 16390 |

**Extended Table 5: True IBD counts filtered by density and IBD informativity score method**

Furthermore, the two methods were largely concordant as ~92% of the indicated true IBD segments were the same. The score method rejected 662 IBD segments indicated by the density method while it also identified 1906 additional IBDs (Extended Table 6).

| type of IBD | IBD count |
|---|---|
| concordant | 14484 |
| exclusive to density | 662 |
| exclusive to score | 1906 |

**Extended Table 6: Concordance between density and IBD informativity score methods.**

To test against the random distribution null hypothesis, we also collected the number of true IBDs per chromosome indicated by the two methods (Extended Table 7).

| CHR | length (cM) | marker count | IBD count (score) | IBD count (density) |
|---|---|---|---|---|
| **1** | 284.26 | 89079 | 1247 | 1214 |
| **2** | 268.82 | 94167 | 1201 | 1166 |
| **3** | 223.26 | 77600 | 1023 | 952 |
| **4** | 214.20 | 68714 | 1010 | 896 |
| **5** | 204.05 | 69350 | 1011 | 932 |
| **6** | 191.72 | 75811 | 975 | 975 |
| **7** | 187.15 | 59833 | 866 | 746 |
| **8** | 168.00 | 61091 | 780 | 836 |
| **9** | 166.14 | 50628 | 836 | 783 |
| **10** | 180.91 | 58760 | 879 | 975 |
| **11** | 158.22 | 54795 | 792 | 781 |
| **12** | 174.59 | 53881 | 917 | 763 |
| **13** | 125.51 | 38991 | 545 | 781 |
| **14** | 118.60 | 36279 | 468 | 441 |
| **15** | 141.34 | 34407 | 514 | 450 |
| **16** | 134.03 | 34372 | 634 | 530 |
| **17** | 128.50 | 29289 | 611 | 436 |
| **18** | 117.55 | 33899 | 589 | 468 |
| **19** | 107.73 | 18440 | 402 | 113 |
| **20** | 108.21 | 29053 | 572 | 463 |
| **21** | 63.64 | 16031 | 282 | 286 |
| **22** | 72.44 | 15792 | 336 | 159 |

**Extended Table 7: Per chromosome stats on the true IBD counts by the density and IBD informativity score methods**

We plotted the number of true IBD segments and the length of the chromosomes to visualize the correlation expected from our null hypothesis (Extended Figure 10). Our results show that the IBD informativity score method has a better correlation (R=0.95) between chromosome length and IBD counts, suggesting that the distribution of indicated IBD segments is closer to the random distribution as assumed from our null hypothesis. Furthermore, the intercept on the x-axis is closer to the expected 0, suggesting that the score method also has slightly improved sensitivity, without gross error on the specificity.

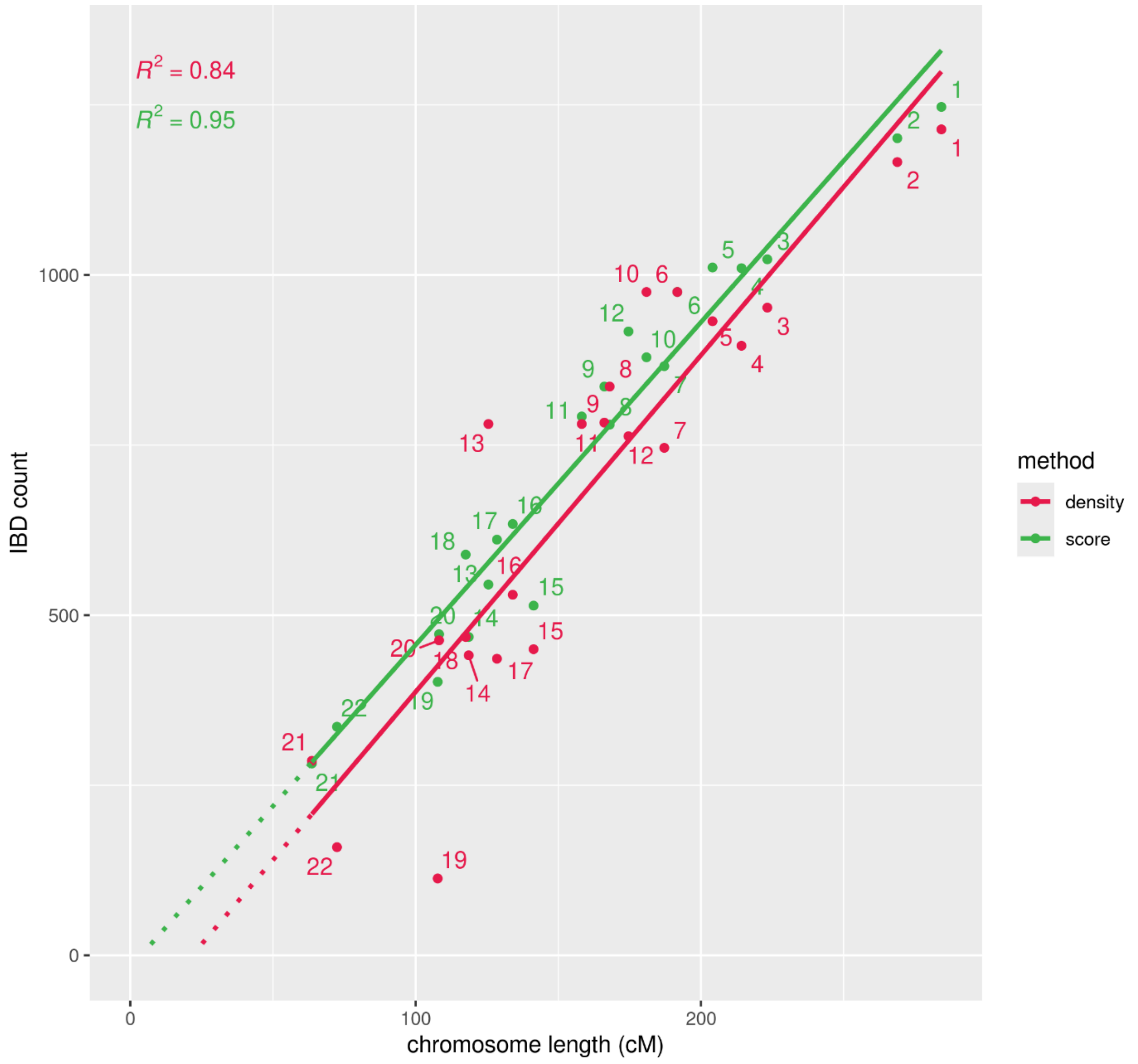


**Extended Figure 10: IBD count per chromosome indicated by the SNP density and the IBD informativity score methods in a large cohort of samples.**

Unsurprisingly, most of the extra IBD segments indicated by the density method and excluded by the score method was on chromosomes (13, 10) where we had an excess of IBD compared to the expectation based on the chromosome length as seen in Extended Figure 10. These excluded IBDs were concentrated around mask areas, where a large number of unrelated individuals shared IBD and had marker densities slightly above the 220 SNP/cM density threshold (Extended Table 8).

| start | end | start (M) | end (M) | length (marker) | length (M) | CHR | ID1 | ID2 | score | density (SNP/cM) |
|---|---|---|---|---|---|---|---|---|---|---|
| **78** | 2345 | 0,0074 | 0,1075 | 2267 | 0,100111 | 13 | RUN2_ind1 | RUN12_ind1 | 136,73 | 226,45 |
| **78** | 2345 | 0,0074 | 0,1075 | 2267 | 0,100111 | 13 | RUN2_ind1 | RUN15_ind1 | 136,73 | 226,45 |
| **73** | 2377 | 0,0064 | 0,1096 | 2304 | 0,103246 | 13 | RUN2_ind2 | RUN22_ind1 | 157,55 | 223,16 |
| **78** | 2383 | 0,0074 | 0,1098 | 2305 | 0,102427 | 13 | RUN2_ind3 | RUN22_ind2 | 161,83 | 225,04 |
| **78** | 2507 | 0,0074 | 0,1147 | 2429 | 0,107308 | 13 | RUN2_ind4 | RUN4_ind1 | 272,81 | 226,36 |
| **78** | 2343 | 0,0074 | 0,1074 | 2265 | 0,100095 | 13 | RUN2_ind5 | RUN19_ind1 | 135,61 | 226,29 |
| **176** | 2598 | 0,0092 | 0,1188 | 2422 | 0,109577 | 13 | RUN2_ind5 | R11118.SG | 369,5 | 221,03 |
| **73** | 2446 | 0,0064 | 0,1127 | 2373 | 0,106332 | 13 | RUN2_ind6 | RUN3_ind1 | 213,51 | 223,17 |
| **73** | 2376 | 0,0064 | 0,1095 | 2303 | 0,103196 | 13 | RUN2_ind6 | RUN18_ind1 | 156,85 | 223,17 |
| **78** | 2629 | 0,0074 | 0,1226 | 2551 | 0,115238 | 13 | RUN2_ind6 | RUN18_ind1 | 404,21 | 221,37 |
| **74** | 2376 | 0,0064 | 0,1095 | 2302 | 0,103148 | 13 | RUN2_ind6 | RUN20_ind1 | 156,85 | 223,17 |

| | | | | | | | | | | |
|---|---|---|---|---|---|---|---|---|---|---|
| **78** | 2458 | 0,0074 | 0,1128 | 2380 | 0,105427 | 13 | RUN2_ind7 | RUN13B_ind1 | 224,44 | 225,75 |
| **78** | 2457 | 0,0074 | 0,1127 | 2379 | 0,105385 | 13 | RUN2_ind7 | RUN19_ind1 | 223,49 | 225,74 |
| **78** | 2420 | 0,0074 | 0,1119 | 2342 | 0,104523 | 13 | RUN2_ind8 | RUN13A_ind1 | 190,76 | 224,07 |
| **78** | 2420 | 0,0074 | 0,1119 | 2342 | 0,104523 | 13 | RUN2_ind8 | RUN22_ind1 | 190,76 | 224,07 |

**Extended Table 8: Example IBDs at chromosome 13 excluded by the IBD informativity score method.**

Since this region had much higher number of IBD segments than expected compared to our null hypothesis, marker informativity was low. Consequently, their IBD informativity score was only between in the range of 136-404 significantly below the 1760 applied threshold, although the SNP density within these IBD segments was slightly above the overall 220 SNP/cM density threshold. Despite the sufficient marker density, the low IBD informativity scores mean that this area is a "mask" area, as also shown in Extended Figure 9.

The other example where the density method significantly deviated from the null hypothesis was chromosome 10 (Extended Figure 10). In Extended Table 9 we present a few examples of the IBD segments excluded by the IBD score method, all found within the low confidence centromeric region of chromosome 10 (Extended Figure 9).

| start | end | start (M) | end (M) | length (marker) | length (M) | CHR | ID1 | ID2 | score | density (SNP/cM) |
|---|---|---|---|---|---|---|---|---|---|---|
| **18598** | 20717 | 0.6194 | 0.7002 | 2119 | 0.0808 | 10 | RUN12_ind1 | RUN10_ind1 | 1359.5 | 262.2 |
| **18526** | 20848 | 0.6180 | 0.7049 | 2322 | 0.0869 | 10 | RUN2_ind1 | RUN2_ind1 | 1477.5 | 267.1 |
| **18670** | 20944 | 0.6208 | 0.7063 | 2274 | 0.0855 | 10 | RUN13B_ind1 | R10620.SG | 1396.8 | 265.9 |
| **18702** | 20945 | 0.6216 | 0.7063 | 2243 | 0.0847 | 10 | RUN13B_ind1 | R10631.SG | 1371.7 | 264.8 |
| **20454** | 23051 | 0.6765 | 0.7907 | 2597 | 0.1142 | 10 | RUN12_ind1 | RUN18_ind3 | 1821.9 | 227.5 |
| **18670** | 22109 | 0.6208 | 0.7616 | 3439 | 0.1408 | 10 | RUN12_ind2 | RUN18_ind4 | 2188.2 | 244.2 |
| **17380** | 22173 | 0.5844 | 0.7643 | 4793 | 0.1799 | 10 | RUN12_ind2 | RUN22_ind1 | 3419.7 | 266.4 |
| **17309** | 20419 | 0.5822 | 0.6650 | 3110 | 0.0827 | 10 | RUN14_ind1 | RUN8_ind1 | 2420.6 | 375.8 |
| **17691** | 20684 | 0.5982 | 0.6969 | 2993 | 0.0987 | 10 | RUN15_ind2 | RUN5_ind1 | 2164.3 | 303.2 |
| **21073** | 24566 | 0.7109 | 0.8225 | 3493 | 0.1116 | 10 | RUN15_ind2 | RUN21_ind1 | 2889.8 | 313.1 |
| **21127** | 24767 | 0.7324 | 0.8267 | 3640 | 0.0943 | 10 | RUN17_ind1 | RUN1_ind1 | 3040.3 | 386.1 |
| **20963** | 24253 | 0.7070 | 0.8157 | 3290 | 0.1087 | 10 | RUN18_ind1 | RUN6_ind1 | 2664.8 | 302.7 |
| **18670** | 22119 | 0.6208 | 0.7631 | 3449 | 0.1423 | 10 | RUN18_ind1 | RUN22_ind1 | 2197.0 | 242.4 |
| **17681** | 23343 | 0.5964 | 0.7982 | 5662 | 0.2019 | 10 | RUN19_ind1 | RUN21_ind1 | 4158.1 | 280.5 |
| **18528** | 21422 | 0.6181 | 0.7412 | 2894 | 0.1231 | 10 | RUN19_ind2 | MJ-43_noUDG.SG | 1773.3 | 235.2 |

**Extended Table 9: Example IBDs from chromosome 10 excluded by the IBD informativity score method.**

In case of chromosome 10, the core of the low confidence area between ~61-70 cM has a general high mean marker density, with some larger gaps between markers within the region and also around 71 cM (Extended Figure 9). All the indicated IBD segments within this region have greater than 220 SNP/cM density; therefore, they were kept by the density filtration method. On the contrary, shorter segments of IBD that only span the low confidence area were excluded due to low IBD informativity scores. However, in most cases, after truncating the low-confidence regions, the length of the high-confidence part of the IBD fell below the threshold, leading to exclusion by the score method.

In summary, our result suggests that the differences between the two methods are mainly due to that fact that a small fraction of genomic regions has excessive raw experimental IBD counts, signaling that the region has low confidence in IBD analysis. While the majority of such regions also fall within the low SNP density areas, a tiny fraction of these region had above than 220 SNP/cM density leading to significant number false positive IBD segments with the density approach.

In contrast to the suboptimal specificity observed in certain chromosomes, Extended Figure 10 also illustrates that the density method has a lack of sensitivity in a few chromosomes, with chromosome 19 being the most notable example. As described previously, chromosome 19 has a low overall marker density (~171 SNP/cM). Unsurprisingly, a large portion of newly indicated IBDs are on chromosome 19 where the density model excluded many IBDs due to low global SNP density. Since we had many first relatives and even sample duplicates in our cohort, we could verify that known IBDs that were excluded due to the global low SNP density of chr19 were still identified with the probability method (Extended Table 10).

| start | end | start (M) | end (M) | length (marker) | length (M) | CHR | ID1 | ID2 | score | density (SNP/cM) |
|---|---|---|---|---|---|---|---|---|---|---|
| **11** | 18440 | 0,0008 | 1,0773 | 18429 | 1,0766 | 19 | RUN10_ind10 | RUN13A_ind1 | 16081,7 | 171,18 |
| **16** | 18440 | 0,0012 | 1,0773 | 18424 | 1,0762 | 19 | RUN10_ind7A | RUN10_ind7B | 16068,5 | 171,2 |
| **8403** | 18440 | 0,5250 | 1,0773 | 10037 | 0,5523 | 19 | RUN10_ind7A | RUN18_ind15 | 9042 | 181,73 |
| **188** | 5902 | 0,0303 | 0,4536 | 5714 | 0,4233 | 19 | RUN10_ind7A | RUN8_ind19 | 4462,82 | 135 |
| **13732** | 18440 | 0,7514 | 1,0773 | 4708 | 0,3260 | 19 | RUN10_ind7A | RUN8_ind19 | 4229,24 | 144,43 |
| **8403** | 18440 | 0,5250 | 1,0773 | 10037 | 0,5523 | 19 | RUN10_ind7B | RUN18_ind15 | 9042 | 181,73 |
| **178** | 5875 | 0,0281 | 0,4514 | 5697 | 0,4233 | 19 | RUN10_ind7B | RUN8_ind119 | 4444,11 | 134,59 |
| **13732** | 18440 | 0,7514 | 1,0773 | 4708 | 0,3260 | 19 | RUN10_ind7B | RUN8_ind119 | 4229,24 | 144,43 |

**Extended Table 10: Example IBDs from chromosome 19 between known relatives and sample duplicates that were indicated by score but excluded by the density criterion.**

To assess the validity of the IBDs that are unique to a specific filtration method, we also compared the IBD segments suggested exclusively by either method with the consensus IBD segments, which were suggested by both methods. We calculated the number of IBDs that were between sample pairs already indicated by the consensus IBD segments. We also calculated the number of IBDs between newly indicated sample pairs that were not indicated by any IBD within the consensus IBD segments (Extended Table 11).

| IBD suggested by only | IBD between already indicated sample pairs | IBD between newly indicated sample pairs | percent of already indicated | percent of newly indicated |
|---|---|---|---|---|
| density | 74 | 588 | 11,18% | 88,82% |
| score | 1186 | 720 | 62,22% | 37,77% |

**Extended Table 11: Concordance of the sample pairs indicated by the method exclusive IBDs with the sample pairs indicated by the consensus IBD segments (indicated by both methods).**

Our analysis shows that a large portion (~90%) of the IBD segments indicated by the density method only and excluded by the score method indicate a connection between the sample pairs that did not share IBD within the consensus IBD segments. While 62% of IBDs proposed by only the score method indicate IBD sharing between sample pairs that already share IBD within the consensus IBD segments. The ratio also suggests that the score method excludes likely false negative IBD segments, as most excluded IBDs are between random unrelated people indicated in genome regions with unexpectedly high IBD counts where the local SNP density is slightly above the 220 SNP/cM threshold. On the other hand, majority of the IBD segments indicated exclusively by the score method are between sample pairs that also share IBD according to the consensus of the two methods, suggesting that these IBD segments are likely true positives that only fell out due to the SNP density criterion in the genome regions with lower local SNP density.

Lastly, we also plotted the inferred IBD count and cumulative IBD length for the greater than 12 cM long IBD segments indicated by the score method (Extended Figure 11) to compare the distribution with the empirical distribution published in the original ancIBD manuscript based on 4248 Eurasian individuals [13].

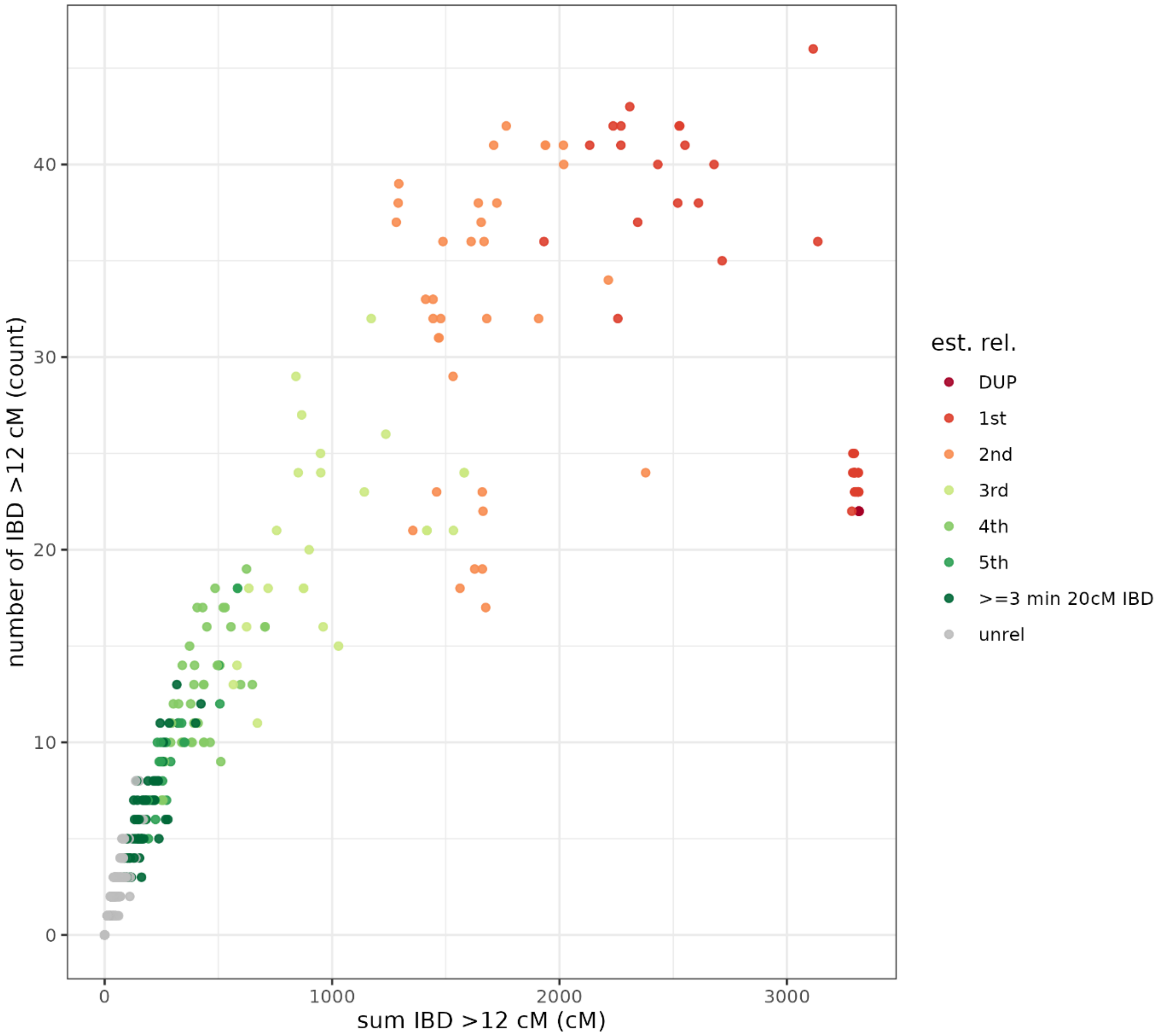


**Extended Figure 11: Inferred IBD among pairs of 1,388 ancient Carpathian basin and Sarmatian ancestry related individuals.**
The plot visualizes both the count (y axis) as well as the summed length (x axis) of all IBD >12cM long. Kinship relations were estimated with the correctKin tool using the same 1240K AADR marker data used in the IBD analysis.

We also applied the five-state HMM for haploid IBD sharing analysis where the length and count distribution of the parent-offspring and full siblings are markedly different. The plot shows that both distribution based on imputed experimental low coverage ancient data are very similar. However, the IBD count of the 39 known sample duplicates and 60 parent-offspring indicated by the score method are within a much closer range (22-25 respectively) to the expected IBD count of 22 (sharing all haploid autosomal chromosome) than the distribution of IBD count of parent-offspring relations indicated by the density-based method (20-35 respectively, as seen in Figure 3 of the ancIBD manuscript). As the data points representing these relations are very closely aligned on Extended Figure 11, it cannot be seen that in the majority of cases the score method indicated the expected 22 IBD count, and only a smaller fraction of cases this value deviated from this (Extended Table 12).

| indicated chromosome count | number of cases |
|---|---|
| 22 | 72 |
| 23 | 15 |
| 24 | 9 |
| 25 | 3 |

**Extended Table 12: The count of >12 cM IBD shared between 39 sample duplicates and 60 parent-offspring, where the expected number of shared IBD is the 22 autosomal chromosome.**

Furthermore, the range of cumulative IBD lengths of the parent-offspring is in a narrower range (3300.7-3356.6 cM) compared to the density-based plot (~3200-3400cM respectively) presented in the ancIBD manuscript. These figures also indicate that the score method offers improved sensitivity and specificity compared to the density approach.

#### Visualization of the TP and FP IBD segments based on the two metrics

To compare the distribution of consensus and method exclusive data, we plotted the distribution of the true positive IBD segments (TP consensus, density, score method), the suggested false positives (FP consensus, density, score method), the true positives suggested by only the density method, the true positives suggested by only the score method, and finally the IBDs that were excluded in the score method after truncating the low-confidence part of the IBD extending into a mask area. We present both the distribution of SNP density and IBD informativity scores versus the length distribution of the IBD segments in the 9 IBD groups (Extended Figure 12-13).

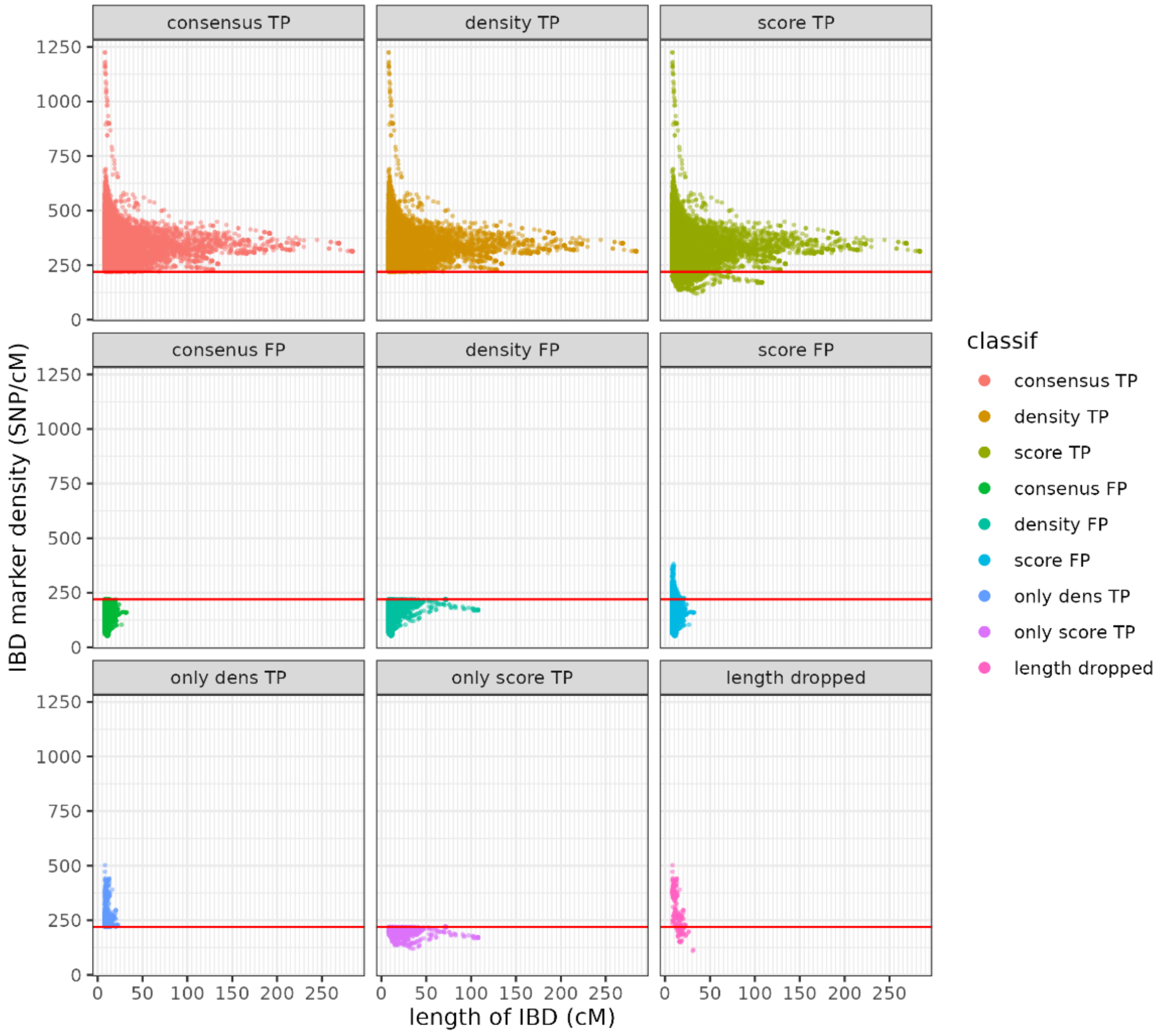

**Extended Figure 12: The SNP density/length distribution of IBDs classified as: true positive (TP) or false positive (FP) by both method (consensus), density method, score method; the TPs of only the density method, only the score method; and the SNP density/length distribution of the length dropped IBDs of the score method.**

We must note that in our cohort there are ~1.6M raw IBDs. Compared to this, the true positive IBDs indicated by either method are only a tiny fraction (15-16K), while the method exclusive IBDs are even smaller fraction (662 and 1906). Consequently, although it cannot be properly visualized in Extended Figure 12, almost all of the 1.6 million raw IBDs are in the consensus FP category as excluded by both methods. In the case of the indicated false positive IBD segments, Extended Figure 12 shows that compared to the density method distribution, the consensus distribution is much more similar to the score method distribution. We also see that the score method FPs include some very short IBDs with high SNP density. According to the IBD informativity formula, this can only happen in the case of IBD segments that consists largely of markers with low marker informativity. Therefore, these short IBDs must fall into genomic regions where a much higher number of individuals share IBDs than expected by the random distribution of true IBD segments. It is important to note that the TPs exclusive to the density method indicate small IBD segments only, with most segments marginally above the 220 SNP/cM density threshold. In contrast, the true positives identified exclusively by the score method are generally longer IBD segments, some even exceeding 100 cM length.

The IBD informativity score versus the length distribution of the IBD segments for the same 9 IBD groups is shown in Extended Figure 13.

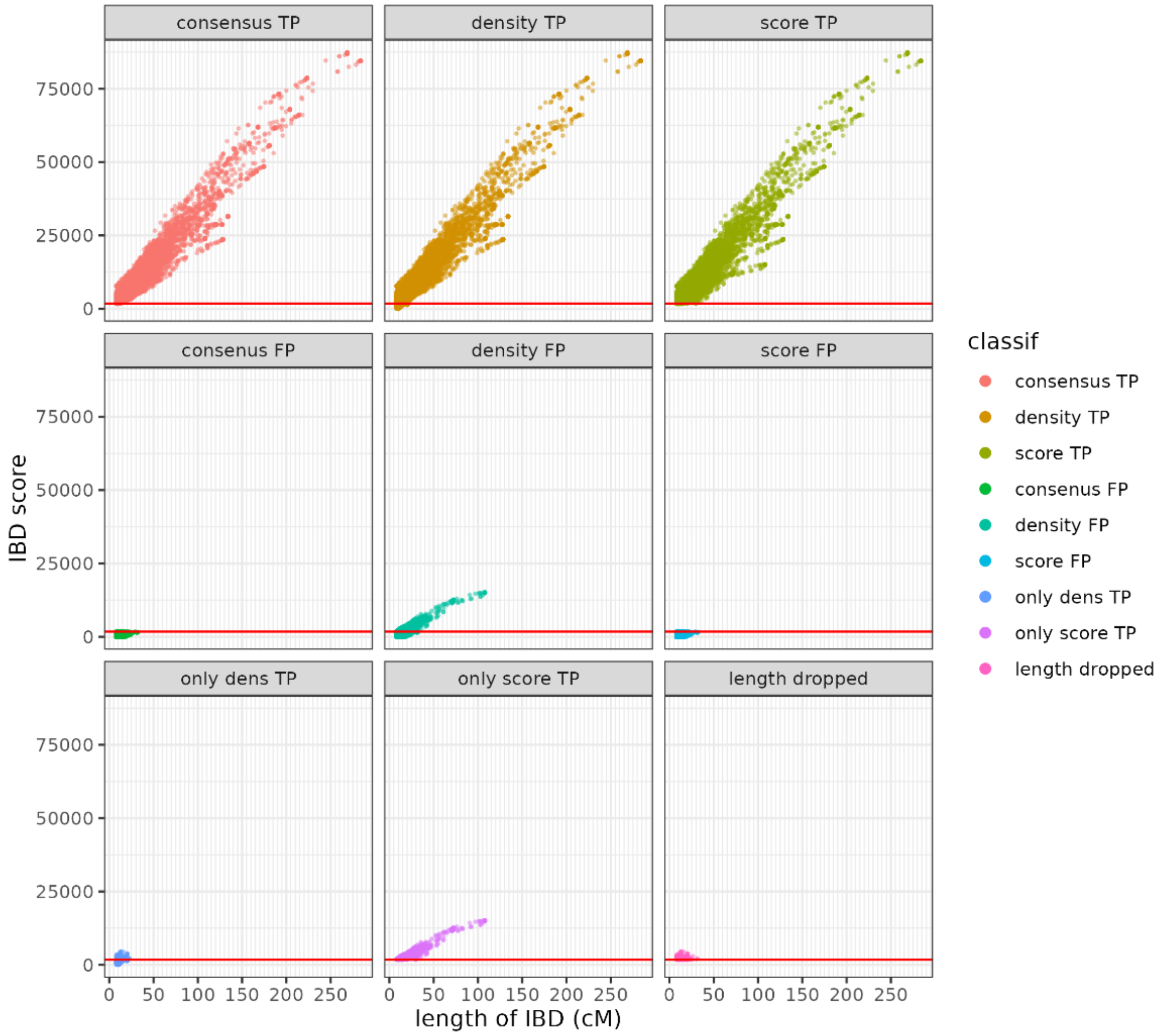


**Extended Figure 13: The IBD informativity score/length distribution of IBDs classified as: true positive (TP) or false positive (FP) by both method (consensus), density method, score method; the TPs of only the density method, only the score method; and the IBD informativity/length distribution of the length dropped IBDs of the score method.**

As we can see in Extended Figure 13 the IBD informativity score differentiates TP and FP IBDs on a much larger scale. Most FPs included in the consensus FP group have <1760 IBD informativity score and are confined to a very small distinct area in the plot. On the other hand, most consensus TPs have magnitudes of higher IBD informativity scores and are more distinct from consensus FPs compared to the SNP density plot (Extended Figure 12). Similarly, as seen in Extended Figure 12 the distribution of consensus FP IBD segments is more similar to the distribution of the IBD score method compared to the SNP density method. Furthermore, the distribution of density FP is much more similar to the consensus TP compared to the consensus FP, indicating that these IBDs were falsely rejected by the global density threshold. On the contrary, the density TPs are all low-score IBDs, many of which are also included in the length dropped group, indicating that while the marker density within the IBD segment is above the 220 SNP/cM threshold, these IBD segments are mainly overlapping or extending into low-confidence mask areas and their IBD informativity score or their high-confidence IBD length is below the applied thresholds.

Our result shows that the distribution of the consensus TP and FP IBD segments aligns more closely with the distributions indicated by the IBD informativity score than with those derived from the density method. The score method displays reduced overlap between the distributions of consensus FP and consensus TP IBD segments. This suggests, that the currently used 1760 IBD informativity score threshold could be likely fine-tuned and a more stringent threshold could be applied to improve specificity without substantial compromise on the sensitivity.

##### Conclusion

Unlike the density method, which treats each marker as equally informative for identifying true IBD segments, our approach assesses the likelihood that the underlying markers indicate a true IBD segment. The formula applied for the calculation of the IBD informativity score mitigates the manual “masking” of low confidence genomic areas in an automatic manner, largely independent of the length of the IBD, the specific set of markers and the differences due to population structure of the analyzed individuals. Our analysis suggests that the IBD informativity score method improves the sensitivity and specificity of the raw IBD filtration compared to the density approach. These improvements together could potentially allow for more robust filtration of raw IBDs and analysis of shorter IBD segments.

### ARCHAEOLOGICAL BACKGROUND

#### GENERAL ARCHAEOLOGICAL DESCRIPTION OF THE 1–5TH CENTURY CE PERIOD IN THE CARPATHIAN BASIN

##### The Steppe connections and migration of the Sarmatians

By the middle of the 1st century CE, the political and ethnic landscape of the Carpathian Basin underwent significant changes. By the end of the previous century, the dominant political entity in the region had become the Dacian Kingdom, centered in Transylvania, which united primarily Dacian and Getae tribes under its rule. In addition to Dacian expansion into the territories of the late Iron Age Celtic and Scythian populations living in the eastern part of the Carpathian Basin, the new century saw the emergence of political formations that would shape the region’s history for centuries. Roman administration gradually established its system in Transdanubia (the province of Pannonia), while north of it, the first Germanic-speaking tribes (the Quadi and the Marcomanni) appeared. Meanwhile, taking advantage of the temporary weakening of the Dacian state, the first Iranian-speaking groups, the Sarmatians, moved into the northern strip of the Duna-Tisza Interfluve [14,15].

The Sarmatian tribes (the Aorsi, Siraci, Roxolani, and Iazyges) had already been influential groups in the Eastern European steppe since the 3rd-2nd centuries BCE. Culturally, in terms of their lifestyle and social structure, they were closely tied to the Scythian groups that preceded them [16–18]. The Sarmatian tribes, who migrated from the Ural and Volga regions north of the Black Sea, took over the role of the Scythians within the nomadic world after the collapse of Scythian power and brought the steppe under their control. Among the confederations of tribes collectively referred to by the ancient world as "Sarmatians," the Iazyges and Roxolani, living furthest to the west, played a prominent role in the history of the Carpathian Basin. According to written sources, the Iazyges, arriving from the lower Danube region, were the first to enter the Great Hungarian Plain, when they became mercenaries for Vannius, king of the Quadi, around 50 CE [15]. These communities exploited a temporary political situation, as neither the Roman political and administrative system had yet fully established itself following Roman expansion, nor had Dacian power fully recovered from its brief decline.

The first archaeological materials associated with these groups date from the end of the 1st century to the beginning of the 2nd century, showing a slight temporal shift compared to the written sources. The early Sarmatian communities are primarily represented by small family cemeteries and the graves of women and children, often containing gold ornaments brought from the east, forming what is referred to in the archaeological literature as the "gold horizon" of burials. Alongside these, male elite figures, associated with steppe artifacts such as tamga-marked gold fittings and ring-hilted swords, also emerge around the turn of the century (e.g., Dunaharaszti, Újszilvás) [19,20]. The material culture of the first migrating generation, showing nomadic and Black Sea regional influences, was also impacted by the local Celtic and early Roman culture, as well as the regional Dacian culture [15,21]. However, the early elite burials often contained objects reflecting international connections with more distant regions (e.g., Vistula region, Veresegyház) [21,22]. By the end of the 1st century, the Sarmatians played an increasingly important role in local political and diplomatic affairs, becoming regular participants in the anti-barbarian wars led by Emperor Domitian [1].

#### The Sarmatians' relations with neighboring peoples (Quadi, Vandals, Marcomanni, Romans)

After the fall of the Dacian Kingdom following Roman military campaigns in 106 CE, Sarmatian settlements, as indicated by archaeological findings, extended eastward beyond the Tisza River and began to populate the southern areas of the Great Hungarian Plain near the Roman frontier (Bácska and Banat, now northern Serbia) [1,23]. Even at the time of their first appearance, the Sarmatian communities in the Great Plain were part of a broader international network, as evidenced by their earliest burial finds (e.g., Dabas, Veresegyház). During the early Sarmatian period, while Eastern-style material culture and burial traditions were prevalent, the greatest influences came from Dacian and Roman provincial cultures.

The populations migrating from the steppe into the Carpathian Basin quickly adapted to local conditions, abandoning their nomadic lifestyle relatively swiftly. This transition is evidenced by the extensive settlement network that developed and the archaeological findings indicating significant agricultural and artisanal activities. [24]. Although several elements of their burial customs remained tied to steppe traditions (e.g., graves with ditch enclosures, burial mounds, and incense burners), these practices saw little change over the centuries [25]. The material culture of the Sarmatians in the Great Plain incorporated many local influences, especially in clothing, weapons, and pottery, which bore marks of Celtic, Dacian, and Roman impact [26].

In addition to elements of early Celtic-Dacian pottery traditions, the Sarmatians increasingly came under Roman influence from the 2nd century onward. As they expanded, they likely absorbed local populations throughout the century [27,28]. During the Marcomannic Wars (166–180 CE), the Sarmatian groups in the Great Plain launched intense attacks on neighboring Roman provinces (Pannonia, Dacia, Moesia). Throughout the war and its aftermath, the movement of small warrior bands outside the Roman Empire likely intensified, as they sought plunder near the empire's borders. Despite the Roman victory and the harsh peace terms (e.g., trade restrictions and mandatory conscription), by the end of the 2nd century, the Sarmatians emerged as one of the most significant groups in the region. Unlike the Germanic tribes, their territories were not occupied by the Romans during the conflicts, which contributed to their continued prominence [29].

The settlement of a significant portion of the Great Hungarian Plain was completed during this period, and a network of interconnected settlements was established. From the late 2nd to the mid-3rd century, their relations with the Romans and Germans flourished, as evidenced by the influx of imported goods into the Great Hungarian Plain [19,26,30,31]. Roman items appearing in Sarmatian territories were quickly replicated and adapted to their local culture. The 2nd and 3rd-century local material culture can be interpreted as a kind of peripheral Roman culture in the Barbaricum. These close ties with Rome laid the economic foundation for producing Roman-influenced goods locally during the 3rd and 4th centuries.

It is also likely that new Sarmatian groups migrated from the east to the Great Hungarian Plain during and after the Marcomannic Wars, as seen in the distinct archaeological finds and new Eastern burial rites (e.g., the Hévízgyörk-Vizesdpuszta group) [1,32–34]. Alongside increasing Roman contacts, a mixed and blended culture also emerged from the 2nd century onward in regions neighboring the Germanic peoples [35–37]. In regions adjacent to the Quadi and Przeworsk culture territories, objects typical of Germanic culture were found, with similar items increasingly appearing further south in the Great Hungarian Plain. This can be seen as the blending of cultures (as reflected in recent finds and scientific data from the Hódmezővásárhely region) [38].

During this period, a new neighbor appeared near the Upper Tisza region—the Przeworsk culture (the archaeological culture of the Vandals)—with whom the Sarmatians developed intensive relations in their northern periphery [39]. By the late 3rd century, pressure on the Sarmatian groups increased due to the migration of Gothic tribes into the Black Sea region. Internal tensions manifested in increased raids on Roman territories and internal conflicts. The Roman state also intervened in the affairs of the late Sarmatian period in the Great Plain, building defensive structures such as parts of the so-called Csörsz Ditch.

The 4th century saw further population growth, likely due to the arrival of new groups from the east. Alongside strong late Roman connections, the material culture of the Great Hungarian Plain was also heavily influenced by the Marosszentanna-Chernyakhov culture (an archaeological culture that emerged in Eastern Europe from the 3rd century, incorporating both earlier Sarmatian populations and newly settled Eastern Germanic groups). These influences became more prominent in the archaeological record of the Great Hungarian Plain during this period [40–42].

#### The Sarmatians' lifestyle in the Great Hungarian Plain: semi-nomadic to settled

The Sarmatian populations, who appeared in the Carpathian Basin during the 1st and 2nd centuries CE, brought with them a nomadic way of life from the steppe regions above the Black Sea and the Lower Danube. However, the newly acquired territory was not conducive to sustaining their previous lifestyle for long periods, forcing the Sarmatian groups to gradually adopt a more settled and complex way of life. Archaeological evidence of larger, permanent settlements begins to appear by the mid-2nd century, but it was during the 3rd and 4th centuries that these large, village-like settlements reached their peak across the Great Hungarian Plain [43].

Despite the shift to a settled lifestyle with the emergence of agriculture, large-scale animal husbandry continued to play a significant role in the everyday livelihood of the Sarmatians in the Carpathian Basin. Based on current botanical and archaeozoological data, the cultural foundations of steppe nomadism persisted throughout the period [44]. Archaeological findings primarily reflect the cultivation of crops typical of nomadic agriculture (barley, emmer wheat, millet), and the livestock, dominated by cattle and small ruminants like sheep, also exhibits characteristics typical of nomadic practices.

Due to their trade and cultural interactions with the Romans, certain crops and animal breeds from the provinces also appeared in regions near the frontier. Additionally, crossbreeding between Roman and local livestock is evident in some of these areas [44–46]. Nevertheless, the Sarmatians maintained strong elements of their nomadic heritage, particularly in agricultural and pastoral practices, even as they adapted to a more settled way of life on the Great Hungarian Plain.

#### Changes in the 4th–5th centuries and the possible continuity of the Sarmatians

From the latter half of the 4th century onwards, the Barbaricum (non-Roman territories) of the eastern Carpathian Basin underwent significant transformations. The increasing pressure of the Huns' advance, beginning in the 370s, triggered a large-scale migration across the former imperial regions of Central Europe. Tensions had already been building before the Huns' arrival, evident from attacks on Roman borders and changes in archaeological finds. The Hunnic migration resulted in cultural shifts, with earlier archaeological cultures showing foreign influences in their final phases, likely due to the movement of smaller groups [47].

In the Great Hungarian Plain, these changes were reflected in the appearance of new objects and customs in Sarmatian archaeological sites, indicating the presence of new populations. During this period, more weapon burials were discovered compared to previous centuries. The influence of the Marosszentanna-Chernyakhov culture (from the east) and the Przeworsk culture (associated with Vandals and other Germanic groups) also became evident. Previously, research had focused on northeastern Hungary (e.g., the Tiszadob group) to track these changes, but similar phenomena have now been found throughout the Great Hungarian Plain [36], reflecting the blending of different cultures and preparing the way for the unified Hunnic period [48,49].

By the early 5th century, written sources and archaeological evidence suggest that part of the earlier Roman-period population left the Carpathian Basin (e.g., Radagaisus' invasion, the westward migration of the Alans, Vandals, and Suebi). The late Sarmatian culture is primarily identified through settlements, which continued to be occupied into the Hunnic period [45,49,50]. Some of the larger cemeteries may have been abandoned around the early 5th century, but many settlements show continuity, even after the mid-5th century, despite new influences.

In the southern regions of the Great Hungarian Plain (around Szeged and Csongrád), archaeological sites from the Hunnic period display a combination of contemporary artifacts with Sarmatian traditions, especially in burial customs and material culture [51,52]. Written sources suggest that a significant portion of the Sarmatian population remained in the area. Even after the collapse of the Hunnic Empire, records indicate the presence of Sarmatian leaders in the region. The last known reference to an independent Sarmatian political structure dates to the 470s. After this, it is likely that the remaining Sarmatian population assimilated into the emerging Gepid Kingdom [53].

Thus, despite the dramatic upheavals of the 4th and 5th centuries, there is evidence of Sarmatian cultural continuity in parts of the Carpathian Basin, especially in their settlements, burial practices, and interactions with new groups like the Huns and Gepids.

#### Chronology of the Sarmatian Period in archaeological research

The Sarmatian period has different chronological frameworks in Eastern Europe and the Carpathian Basin, which often do not align. In the vast region between the Ural Mountains and the Carpathian passes, Russian and Ukrainian archaeological research during the 20th century considered the peak of Sarmatian culture to span from the 2nd century BCE to the end of the 4th century CE. This lengthy period is generally divided into three sub-periods, primarily defined by distinct archaeological cultures:

1. **Early Sarmatian Period** (3rd/2nd century BCE – 1st century BCE).
2. **Middle Sarmatian Period** (1st century CE – mid-2nd century CE), known as the Susli culture.
3. **Late Sarmatian Period** (second half of the 2nd century CE – 4th century CE), transitioning into the early Sipovo culture [1,54].

Efforts have been made in recent decades to align the chronology of Eastern European nomadic archaeological cultures with the periodization systems established for Central and Northern Europe.

Recently, Alexandr Symonenko has attempted to establish a new chronology for the Sarmatian archaeological horizon in Eastern Europe [55]. He fundamentally divided the previously established system, which mainly followed Russian archaeological cultures, into two major periods (Period I: the 2nd century BCE to the mid–2nd century CE , period II: the second half of the 2nd to the 4th century CE), within which he distinguished further sub-periods (Period I: stage A1 – the 2nd century BCE; stage A2 – end of the 2nd and 1st century BCE; stage A3 – 1st half of the 1st century CE; stage B (with the horizon of the remains with features specific to the new Sarmatian wave arriving from the east) – the second half of the 1st – mid–2nd century CE. Period II: stage C1 (early) – the second half of the 2nd – 1st half of the 3rd century CE; stage C2 (late): sub-stage C2a – the second half of the 3rd – mid–4th century CE; sub-stage C2b – the second half of the 4th century CE.). In doing so, he took into account objects with more reliable dating value and the shifts in archaeological horizons. In recent years, Vitalie Bârcă, in collaboration with

Simonenko, has conducted an in-depth study of the Sarmatian archaeological material from the eastern foothills of the Carpathians, the modern Romanian Plain, and the Pontic region [14,56,57].

Hungarian archaeological research first began focusing on Sarmatian groups in the early 20th century. The first attempts at distinguishing territorial and chronological groups were made by Mihály Párducz, who based his analysis on burial finds [58–60]. Párducz concentrated on the connections between the nomads of the eastern steppes and other cultures in the Carpathian Basin. He combined these perspectives with his observations on burial customs to establish regional and chronological groups, though these categorizations did not stand the test of time.

Later, in the 1980s and 1990s, more comprehensive syntheses emerged, which remain influential in Hungarian Sarmatian archaeology today. In 1989, Andrea Vaday created a chronological framework for the Sarmatian period in the Carpathian Basin by carefully analyzing the material finds and drawing on the better-researched Roman and Germanic chronologies. [61]. Vaday also incorporated data from historical sources to define broader chronological divisions, often tied to significant milestones in Sarmatian-Roman interactions. The Sarmatian chronological system of the Carpathian Basin have been further developed at the level of individual objects and phenomena in recent decades by Eszter Istvánovits and Valéria Kulcsár, who, alongside Eastern analogies, extensively utilized the Roman and Germanic artifacts that were available [1].

The Carpathian Basin's eastern half has its own distinct archaeological chronology, which poses several challenges from both Central-Northern European and steppe perspectives. For regions outside the Roman Empire, such as the Barbaricum, the chronological framework established in the second half of the 20th century for Germanic territories was typically applied. This framework, based on relative chronological observations and Roman finds, is divided as follows: **B1**: 0 – 80 CE, **B2a**: 80 – 120 CE, **B2b-C**: 120 – 160 CE, **B2/C1-C1a**: 160 – 200/230 CE, **C1b**: 200/230 – 260/270 CE, **C2**: 260/270 – 300 CE, **C3a**: 300 – 340/350 CE, **C3b**: 340/350 – 370/380 CE, **D1**: 370/380 – 410 CE [62].

Based on the main historical events of Roman-Sarmatian relations and changes in material culture, previous archaeological research has classified Sarmatian archaeological remains in the Carpathian Basin into three chronological periods. These periods are: 1. **Early Sarmatian Period**: From the arrival of the Sarmatians in the Great Hungarian Plain (second third and second half of the 1st century CE) to the 2nd half of the 2nd century CE. 2. **Middle Sarmatian Period**: From the period of the Marcomannic wars to the end of the 3rd century CE abandonment of Dacia. 3. **Late Sarmatian Period**: From the end of the 3rd century CE to the last third of the 5th century CE.

To date, only one attempt (by Margit Nagy, in the case of the Pécel cemetery) has been made to integrate the archaeological development of Sarmatian groups in the Great Hungarian Plain into this system [63]. This attempt was motivated by the strong Germanic influences at the site. Future research may likely follow this path, as the Sarmatians of the region were closely intertwined with the cultural and economic context of their Germanic neighbors.

Recently, Lavinia Grumeza has sought new approaches to partially synchronize the Hungarian archaeological chronology with the northern and central European Germanic material. Additionally, Norbert Kapcsos has applied mathematical methods to develop a new chronological framework for sites along the lower Maros (Mureș) River, contributing further to the study of the Sarmatian period in the region [23,64].

#### DESCRIPTION OF THE STUDIED SARMATIAN PERIOD CEMETERIES AND GRAVES

##### Apátfalva – Nagyút dűlő M43 site 43 (AND; Csongrád-Csanád County, Hungary) [65–68]

The site represents a late Sarmatian cemetery consisting of 47 graves, fully excavated by Tibor Paluch between 2010 and 2011. The burial ground was encircled by a combination of artificially created and naturally formed ditches likely connected to the river Maros. The majority of the graves exhibited a south-north (SN) orientation, with slight deviations noted in some towards south-southwest (SSW) to north-northeast (NNE), and in initial cases, north-northeast to south-southwest (NNE–SSW) and north to south (N–S) orientations. These simple pit graves, larger than the average size in other Sarmatian cemeteries, featured five graves at the northern and southern borders encircled by ditches. Some graves revealed traces of coffins or shrouds within the pit. While most of the graves represented single burials, skeletal remains belonging to two individuals were found in three graves. In general, the skeletons were found in a supine position with straighten or bent elbows. Approximately 43.18% of the burials were disturbed or robbed, limiting the interpretative scope of archaeological and anthropological findings. The archaeological assemblage included clothing elements (e.g., brooches, belt buckles), jewelry (e.g., torques, amulets, bracelets), ceramic vessels, and tools (e.g., iron knives, bone needle case). Iron fragments, potentially from a spear, were found in one grave. The cemetery was dated to the mid-4$^{th}$ century and the first third of the 5$^{th}$ century CE. Certain artifacts, such as the bone needle case and vessels suggested eastern influences, while burial customs and grave goods exhibited parallels with contemporaneous local sites, such as the cemetery of Óföldeák–Ürmös. The anthropological analysis of the human bones was conducted by Antónia Marcsik and Yvett Kujáni. The preservation of the skeletal remains, in general, was found to be low. The examination of bones from the 47 graves revealed 43 individuals, consisting of 13 sub-adults and 30 adults. In terms of biological sex, 10 males and 23 females were identified. Notably, there were no significant metric or morphological differences observed in the skulls between the sexes. Predominantly Europid characteristics, particularly brachycran and Cromagnoid types, were noted, although minor Mongoloid cranial features were observed in several cases. We could involve five samples in our analysis.

***AND-128: Grave No. 128:***

The robbed grave contained the skeleton of a middle adult male, accompanied by an iron knife and two ceramic vessels. The skull, of Cromagnoid type, displayed Europid characteristics with minor Mongoloid features.

***AND-131: Grave No. 131:***

This robbed burial of a middle adult male, characterized by Europid (Pamyrian) features on the skull, was oriented in the S–N direction and included an iron belt buckle. The grave was encircled by a ditch.

***AND-138: Grave No. 138:***

The skeleton of a middle adult male with Europid (Cromagnoid–Pamyrian) features on the skull was found in the robbed grave, containing an iron knife, a vessel, a bronze belt buckle, and traces of a wooden coffin. Radiocarbon dating has placed the burial earlier, from the second half of the 1$^{st}$ century CE to the beginning of the 3$^{rd}$ century CE.

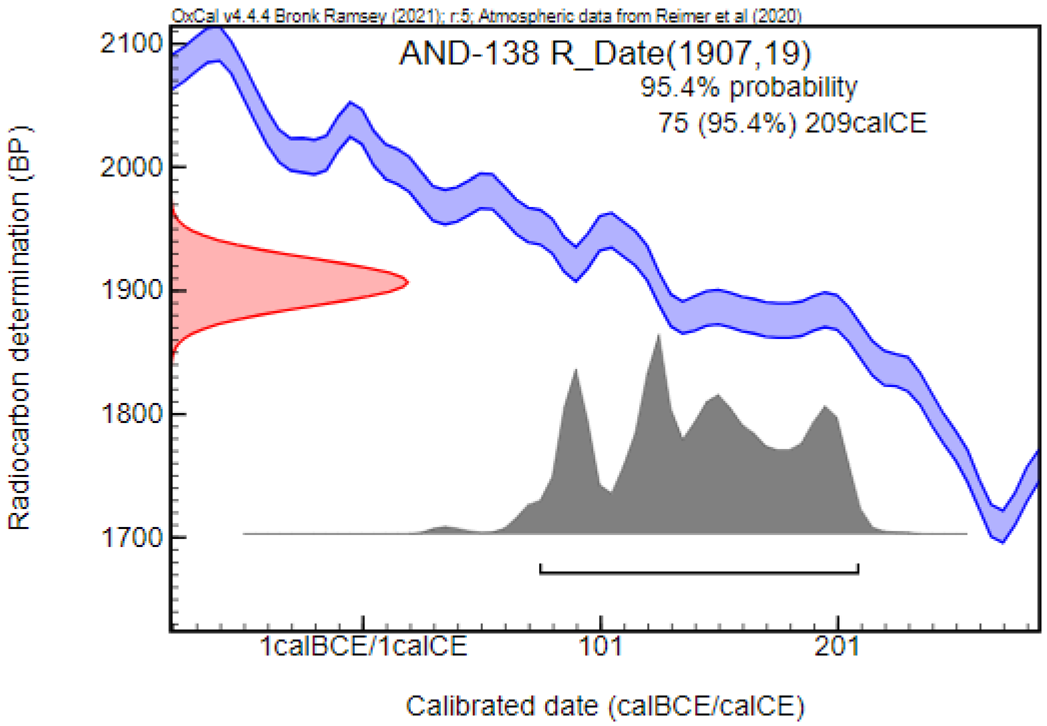


***AND-145: Grave No. 145:***

The robbed burial contained the skeletal remains of a young adult female with Mongolid characteristics on the skull. Traces of a wooden coffin, two bronze brooches, and a bronze earring with hook and loop ends were found in the grave.

***AND-177: Grave No. 177:***

This burial of a young adult female with Europid characteristics and Mongoloid features on the skull, found in a robbed grave encircled by a ditch, contained traces of a wooden coffin and an iron knife.

#### Apc – Farkas-major (AFM; Heves County, Hungary) [69]

During excavations led by Mónika Gutay in 2014, associated with the extension of main road No. 21, settlement objects and a burial ground dating to the Sarmatian and Hun periods were discovered. While the archeological analysis is still in progress, preliminary data revealed layers showing traces of burning in some of the settlement objects, indicative of the settlement's destruction. Additionally, human skeletons were found in some of the settlement objects, an uncommon but characteristic phenomenon of the Late Sarmatian and Hun periods.

The main burial area consisted of a central group of 15–20 graves, along with several smaller groups of 2–3 graves distributed throughout different parts of the burial ground. Grave goods in male burials included iron knives, belt buckles, and occasionally spearheads. The female burials were richer in inventory, including jewelry (e.g., earrings, strings of beads, and bracelets) and ceramic vessels. Roman coins and brooches were commonly found in both the settlement objects and burials. These findings, along with the terra sigillata fragments, indicating the Roman connections of this community. The cemetery was associated with the Sarmatians and dated to the 3rd–5th centuries CE. Anthropological analysis conducted by Antónia Marcsik revealed that the series consisted of the skeletal remains of at least 80 individuals, with generally poor preservation. While the detailed evaluation and publication of the results is still in progress, 65 adults and 15 sub-adults were identified, with 31 females and 35 males. The skulls available for taxonomical analysis mostly showed Europid characteristics, with several belonging to Mongolid groups. We could include seven samples in our analysis.

***AFM-213: Object No. 213:***

The grave, oriented in a SE–NW direction, contained the skeletal remains of an adult female in a supine position, with Mongolid characteristics on the skull. The grave goods included a glass vessel, a buckle near the legs, a spindle whorl, beads near the wrists and neck, and a ceramic vessel shard.

***AFM-273: Object No. 273:***

The grave, oriented in a N–S direction, was highly affected by contemporary robbing, and only fragments of the skeleton, possibly belonging to an adult male, were found in a secondary position. The artifacts recovered from the burial consisted of iron parts possibly belonging to a wooden coffin.

***AFM-590: Object No. 590:***

The burial, oriented in a S–N direction, contained the skeleton of an adult female in a supine position, with Europid characteristics and minor Mongolid features on the skull. The grave goods included coral beads near the neck, a bronze fibula, and a pair of bronze bracelets.

***AFM-940: Object No. 940:***

The fragmented skeletal remains of two adult female individuals in a secondary position were uncovered from the robbed pit, with only some elements of the lower parts preserved in anatomical position. The pit was oriented in a SE–NW direction.

***AFM-1156: Object No. 1156:***

The skeletal remains of an adult female were discovered in the grave, oriented in a S–N direction. Fragment of a buckle, a spindle whorl, and beads near the lower leg bones were found in the grave. Radiocarbon measurement, partially overlapping with the archaeological dating, placed the burial between the second third of the 2nd century CE and the first half of the 3rd century CE.

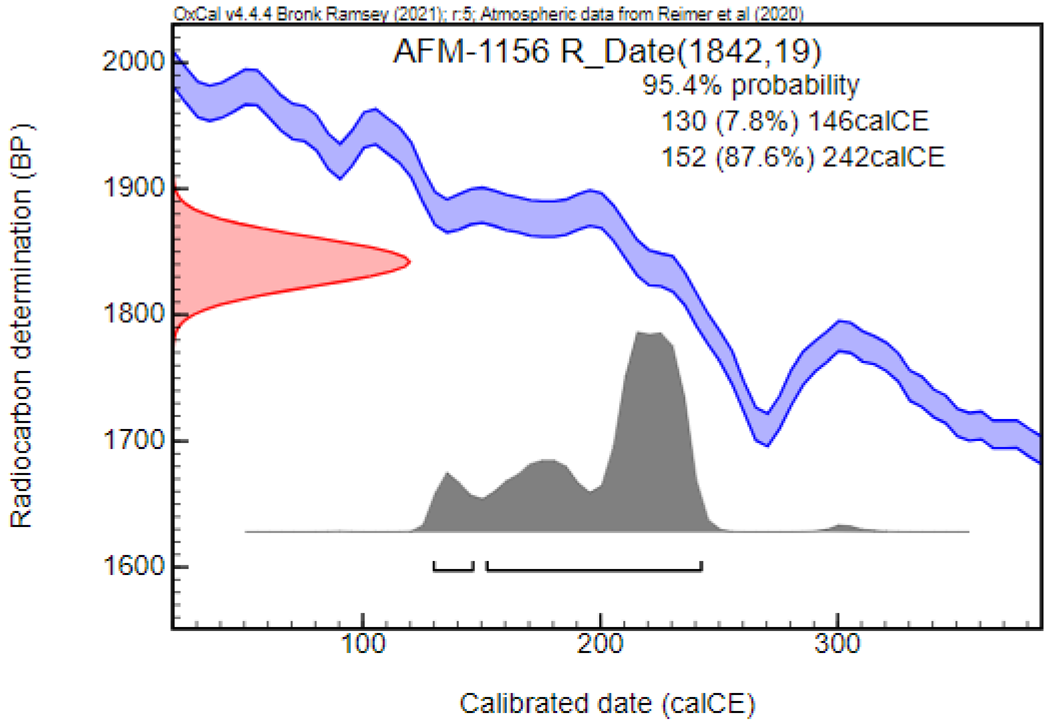


***AFM-1255: Object No. 1255:***

The robbed burial contained the skeletal remains of an adult female, oriented in a SE–NW direction. The lower leg bones were partially missing, and the skeleton was located at the northwestern side of the grave with the knees close to the pit wall. It was assumed that postcranial skeletal remains belonging to two individuals were recovered from the burial. The grave goods included a ceramic vessel.

***AFM-1861: Object No. 1861:***

The bone fragments of an adult female were recovered from the burial. Radiocarbon measurement, partially overlapping with the archaeological dating, placed the burial from the mid-2nd century CE to the mid-3rd century CE.

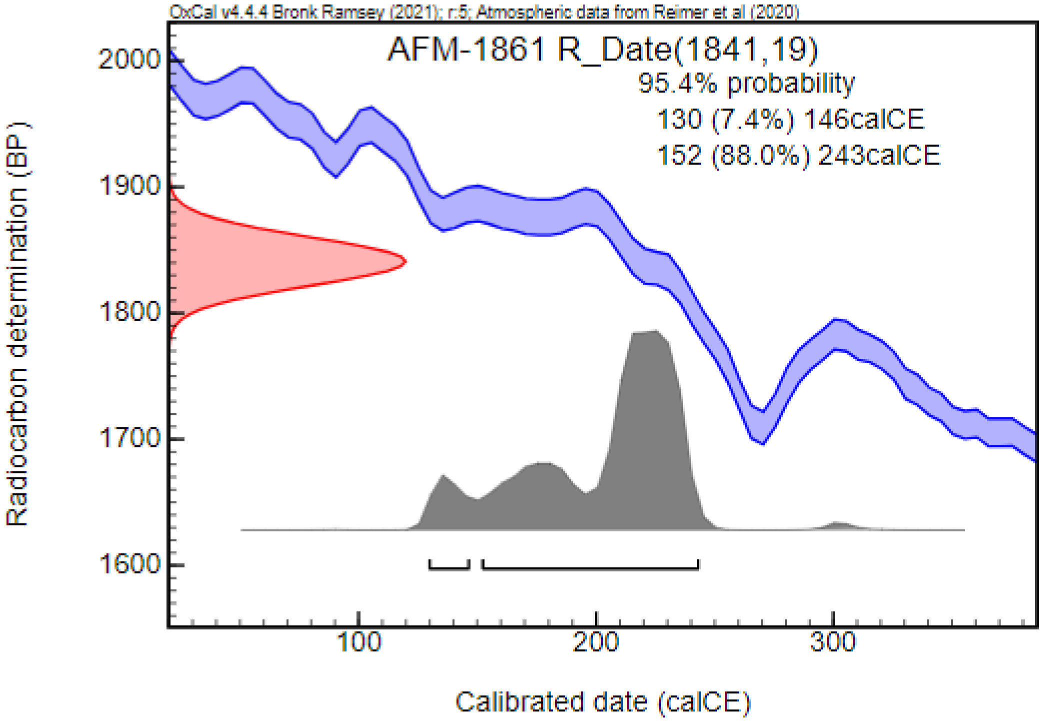


#### Botoșani (BOT; Botoșani County, Romania) [70,71]

In the eastern part of Botoșani, a field survey conducted in 1959 led to the discovery of a settlement, a cemetery, and additional remains of material culture from the Iron Age (Latene III Period) and the beginning of the Migration Period (3rd–5th centuries CE). Subsequent surveys in 1960 and 1961 revealed additional archaeological materials dating to the Iron Age (Latene III Period) and the 4th century CE, along with the identification of a second burial ground. These sites were situated approximately 100 to 500 m from each other. Systematic excavations registered 23 graves in the first cemetery, consisting of 21 single and 2 double burials. In addition, five graves were uncovered in the second cemetery. The archaeological material, not yet detailed in publications, includes jewelry (e.g., bronze earrings, and beads), cloth fittings (e.g., brooches and belt buckles), and mirror fragments. One notable aspect of these cemeteries is the high frequency of skeletons with deformed skulls. The burial customs and grave goods recovered from the sites differ from the characteristics of the Sintana de Mureș-Cerneahov culture, likely containing a mix of elements associated with different populations such as Ostrogoths and Sarmatians. The cemeteries were dated to the 3rd–5th centuries CE, before the Gepid Period. We could involve one sample in our analysis.

***BOT-2: Grave No. 2:***

The skeletal remains of an adult individual with a deformed skull were found in the grave, located in the first cemetery. The grave goods are unknown. Radiocarbon dating has placed the burial primarily from the second half of the 4th century CE to the second third of the 5th century CE, with the possibility of being related to the second half of the 3rd century CE (1.6%) or the first third of the 6th century CE (0.7%).

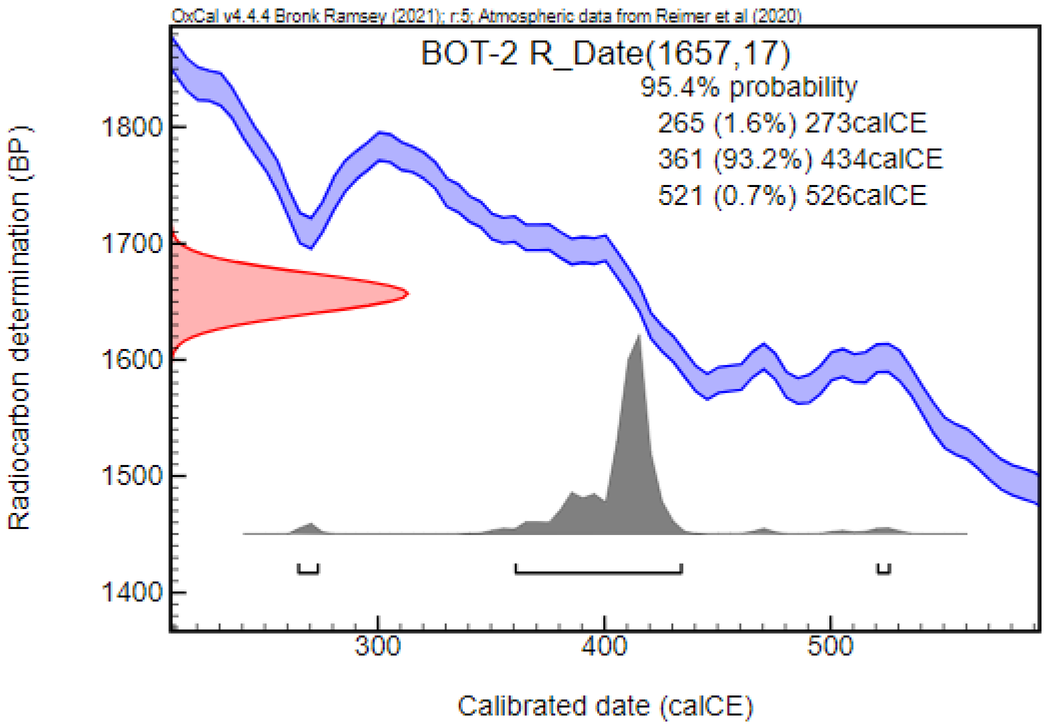


#### Budapest XVII. Rákoscsaba, Péceli út (RAK; Pest County, Hungary) [72]

In 1969, the remains of the first human skeletons and ceramic fragments were uncovered during the excavation of an industrial channel trench. Margit Nagy, representing the Budapest History Museum, identified the site, and under her supervision, 270 burials were excavated between 1971 and 1986 as part of a planned excavation, which was published in 2018.The cemetery consists of two clearly distinguishable groups of graves: in the higher, eastern grave row, burials are placed closer together, while in the western row, they are spaced farther apart and more loosely. Grave surrounded by flat circular ditch and stone-lined burials can be found in both groups. Differences in burial rites are observed between the two groups. On the eastern side, the graves were dug deeper and oriented south-north, while on the western side, the shallower burials were oriented southeast-northwest. Additionally, there are differences between the two groups in terms of certain features and the composition of grave goods (narrower graves in the west, wider graves in the east; in the east, knee brooches that may be associated with Roman military service, while in the west, weapon burials are found, along with different types of amulets used in both groups). The site's uniqueness lies in the stone-lined burials constructed from natural stones. The site is dated between the 2nd and 4th centuries CE. Germanic connections may be suggested by the silver brooch (Straže-Sakrau-type, Almgren VII group) found in Grave 10, certain groups of ceramic vessels, and a belt buckle decorated with a swastika symbol. According to the excavator, Margit Nagy, the population that established the cemetery may have been a mixed ethnic group (Sarmatian and Germanic, possibly Quadi). Nagy suggests that this mixed population could have been a community settled on the Pest Plain by the Romans for primarily military purposes and placed in the vicinity of Aquincum.

***RAK-115: Grave No. 115***

The burial is oriented SSE-NNW, with the upper body of the deceased disturbed and showing moderate preservation. The skeleton's length is 170 cm. The skeletal remains of an adult male were found in the grave. The grave depth is 105 cm, and the dimensions of the burial pit are 234 x 90 x 85 cm. The disturbance likely occurred simultaneously with Grave 109, as a fragment of terra sigillata from the other grave was mixed into the soil of this one. A bronze knee brooch was found beneath the right clavicle and jaw, a limestone bead under the left shoulder, the tip of an iron knife next to the right hand bone, and a knife placed crosswise on the right pelvic bone. Additionally, an iron object and a silex blade were discovered, along with a hand-formed, thick-walled, poorly fired ceramic vessel on the outer side of the right lower leg. This grave can be dated by Margit Nagy to the period C2 (260–330 CE).

***RAK-136: Grave No. 136***

A disturbed burial oriented SSE-NNW was uncovered, containing a deceased individual measuring 150 cm in length. This grave contained the skeleton of a middle adult female. The grave depth is 124 cm, and the dimensions of the burial pit are 192 x 73-69 cm. The grave contained five beads located around the neck area and the outer side of the left femur: a brown twin bead, a yellow and a white, pressed spherical glass bead, a worn barrel-shaped glass bead with a light base and a brown coating, and a black pressed spherical glass bead decorated with a white thread. On the outer side of the right lower leg, an asymmetrical, hand-turned, thick-walled ceramic vessel was found, brownish-black in color. Based on the archaeological material this grave can be dated by Margit Nagy to the period C2 (260–330 CE).

***RAK-211: Grave No. 211***

A burial oriented SSE-NNW was excavated at a depth of 131 cm, with the grave pit measuring 250 x 90 cm. The skeleton of an old adult male was uncovered in the grave. The grave was surrounded by a ditch frame with a diameter of 9 m and a width of 1 m, open on both sides, but disturbed during World War II. The ditch had a depth of 120-143 cm, with a turtle-shaped bottom, from which glass fragments were recovered. Within the grave, two levels of stone rows were found (stone lined burials), but there was no stone packing in the 15-20 cm thick layer above the level of the human skeleton. A long-handled, single-edged iron knife, likely in its original position, was found across the left side of the pelvis, with the blade pointing toward the left forearm. An iron buckle was discovered as a grave good beneath the right pubic bone.

***RAK-251: Grave No. 251***

A disturbed burial oriented SSE-NNW was uncovered at a depth of 120 cm in a pit with a base area of 268 x 80 cm. This burial contained the skeleton of a young adult female. A section of a circular ditch was also observed near the southeastern end of the grave. The spine and right side of the upper body showed signs of disturbance, with the shoulders slightly raised and the skull turned to the right. Three beads were found near the neck: one double truncated cone-shaped white glass bead, one pressed spherical dark blue glass bead decorated with intersecting white threads, and one pressed spherical transparent mosaic glass bead with a green base and yellow flower inlay. An oval iron buckle was located on the left side of the pelvis. About 10 cm from the feet, a well-silted, round-bodied ceramic vessel was discovered, on top of which a double truncated cone-shaped spindle whorl had been placed. Based on the archaeological material this grave can be dated to C3 period (330–400 CE)

#### Csanádpalota – Országhatár M43, 56. lh (CSO; Csongrád County, Hungary) [73]

A preventive excavation led by Dániel Pópity was conducted in relation to the M43 highway section between Makó and the Hungarian border in 2010, 2011, and 2012. The site, situated to the south of the present-day town of Csanádpalota, contained a late Sarmatian cemetery with 53 graves. The burials were generally oriented in the S–N main directions with minor deviations. However, in one case, an opposite orientation toward the NE–SW directions was observed. The majority of the grave pits were regular with rectangular shape and rounded corners. In two cases, the grave pits were constructed in a square shape with large dimensions. Concerning the size of the grave pits, a total of five burials were constructed with significantly larger dimensions compared to the average length and width of the pits in the cemetery. Presumably, these burials belonged to prominent members of the society. Eleven burials were encircled by a ditch. Some of these ditches contained shards of clay vessels, possibly related to the funeral feast. Traces of wooden constructions were detected in four graves, and traces of wooden coffins were found in 12 graves. Sixty-six percent of the graves were affected by robbery, and only 16 graves were left undisturbed. In these cases, the deceased were mostly laid to rest in a supine position. The archaeological material consisted of jewelry (e.g., earrings, necklaces, bracelets, and pendants), cloth fittings (e.g., brooches, belt buckles, and beads), tools (e.g., knives, spindle whorls, and a needle case), and clay vessels. The cemetery was dated from the first half of the 4th century CE to the beginning of the 5th century CE. The anthropological analysis was carried out by Antónia Marcsik. Out of the 51 burials containing skeletal remains, 49 were suitable for anthropological analysis. A total of 37 adults and 12 sub-adults were identified. In terms of biological sex, the series included 17 females and 19 males. We could involve three samples in our analysis.

***CSO-507: Grave No. 40 (OBNR: 266, SNR: 507)***

The grave was oriented in a north-south direction, and a circular ditch was associated with the burial on the southeastern side. Unfortunately, the grave had been robbed or disturbed. Despite this disturbance, traces of a coffin and a burial shroud were still observable. The fragmented skeleton of an adult male was found in this grave accompanied by a single-piece silver brooch with a returned foot and a bronze brooch with a side-turned foot were recovered.

***CSO-526: Grave No. 42 (OBNR: 271, SNR: 526)***

The grave was oriented in a south-north direction (D-N), but based on the available data, the burial had been significantly disturbed. This burial contained the fragmented skeleton of an adult female. From the grave, a bronze brooch with a side-turned foot was recovered.

***CSO-502: Grave No. 50 (OBNR: 297, SNR: 502)***

The grave was oriented in the north-south direction; however, unfortunately, it was disturbed or robbed after the burial. The deceased was interred in a log coffin, and a burial shroud was also used during the interment. From the grave, only a fragment of a brooch and unidentified iron fragments were recovered. The fragmented skeletal remains of an adult male were found in the grave. Radiocarbon measurement, partially overlapping with the archaeological findings, placed the burial between the second third of the 3rd century CE and the first half of the 4th century CE.

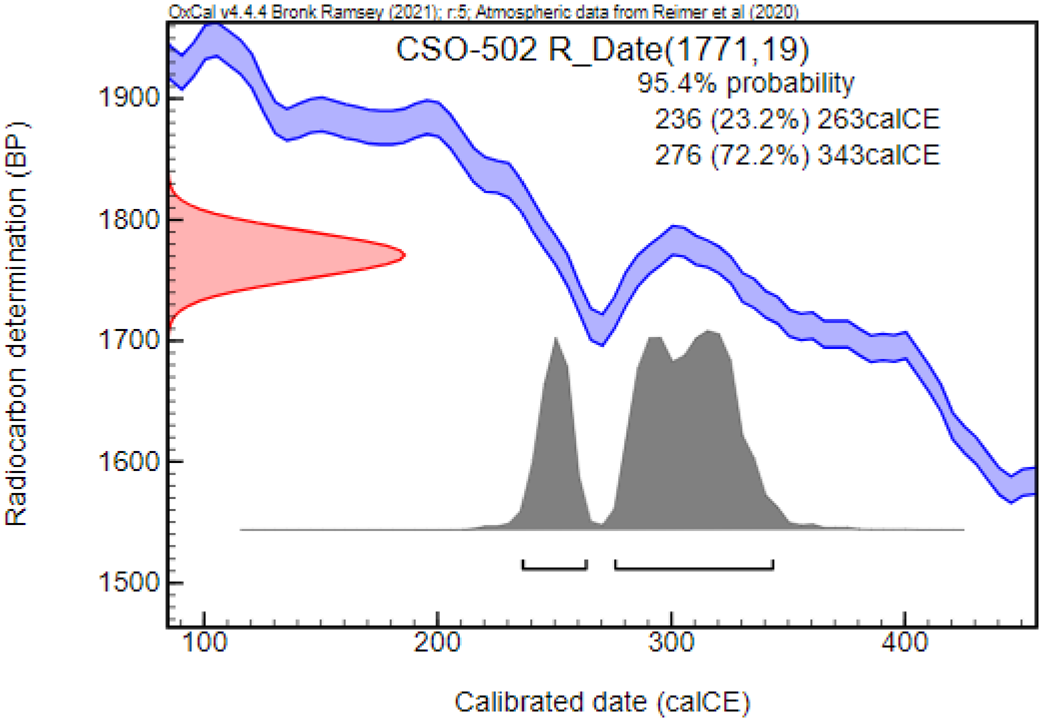


#### Csongrád – Kenderföldek (CSK; Csongrád County, Hungary) [51,52]

Thanks to the excavations conducted by Mihály Párducz in 1949–1950 at Csongrád-Kenderföldek, 122 graves were uncovered and published in 1959. In 1960, during the construction of the

Csongrád military barracks, 18 additional graves were excavated, which were published three years later. The sites, published under different names, are presumably two parts of the same cemetery, a hypothesis supported by the burial rites and the artifacts recovered from the graves. Several factors complicated the research. No comprehensive analysis was made of the first excavation, and aside from basic data and artifact reports, little else is known. The situation is somewhat better for the graves uncovered during the barracks construction, but a thorough analysis was not conducted for these either. Over the past few decades, numerous researchers have studied individual artifact types from the cemetery, providing further data for analysis. Unfortunately, a significant portion of the graves was disturbed, but despite this, a rich assemblage of finds was uncovered. Based on the material culture, the use of the site can be dated to the mid-4th century through the middle third of the 5th century. The main characteristics of the material culture trace back to late Sarmatian traditions, while the influence of the D1 period (new types of belt buckles, silver plate brooches, needle case) and Hunnic-period influences are also clearly visible in the artifacts. The cemetery primarily reflects the burial customs of the earlier Sarmatian period (south-north orientation). Unfortunately, due to contemporary excavation methods, it was not possible to observe the graves surrounded by flat circular ditches. The ceramic material from the graves reflects late Sarmatian traditions. Compared to other sites from this period, glass vessels (cups and jugs) are found in unusually high numbers in the burials. Additionally, weapon grave goods, which are relatively rare in Sarmatian burials, appear more frequently (most of these reflect D1 period and Hunnic-period influences). The closing horizon of the site is difficult to define. Currently, the latest datable objects (Levice–Prša type brooch) place the terminal phase in the middle third of the 5th century, around the middle of the century. The initial anthropological analysis of the human bones was conducted by Antónia Marcsik. However, the low state of preservation of the bones significantly constrained the analysis and subsequent discussion of the results. Specifically, only 59 out of the total 140 graves containing skeletal remains, primarily skulls, were deemed suitable for analysis. Age-at-death analysis revealed 57 adults and only two sub-adult individuals within the series. Among the adults, 24 males and 33 females were identified. The metric and morphological analysis of the skulls indicated potential differences between the sexes, with female skulls predominantly exhibiting gracile features and dolichocrany. The observed cranial characteristics predominantly displayed Europid traits, with Cromagnoid and Nordoid types being the most prevalent. Additionally, minor Mongoloid cranial features were noted in several cases. It was hypothesized that the population represented a mixture of remaining Sarmatians and groups carrying characteristics from the 5th–6th century CE Gepid population. Our analysis included nine samples in this study.

***CSK-9: Grave No. 9***

The skeletal remains of a middle adult male with Europid characteristics on the skull was uncovered in the S–N oriented, robbed grave. Grave goods included an iron knife, a Roman silver coin (Marcus Aurelius denarius), and a glass cup. In addition, iron coffin clamps were found. Radiocarbon dating, confirming the archaeological findings, placed the burial between the mid-4$^{th}$ century CE and the first third of the 5$^{th}$ century CE, with the possibility of being dated to the third quarter of the 3$^{rd}$ century CE (3.8%).

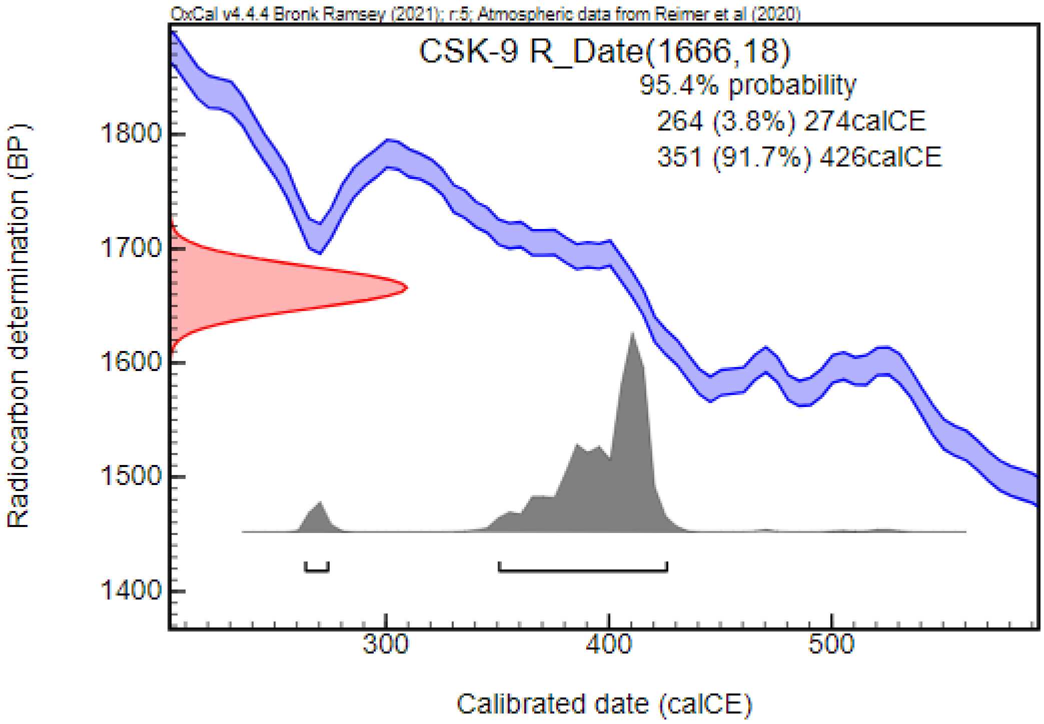


***CSK-18: Grave No. 18***

This grave contained the skeleton of a young adult female with Europid characteristics and Mongoloid features on the skull, accompanied by a biconic ceramic vessel.

***CSK-25: Grave No. 25***

The skeletal remains of a middle adult female in a supine position were recovered from this N–S oriented, robbed grave. Next to the skull, a Murga type jar was found. This grave can be dated to the first half of the 5th century CE.

***CSK-61: Grave No. 61***

This burial contained the fragmented skeleton of a young adult female with Europid characteristics on the skull. Grave goods included an iron spearhead, fittings of a wooden bucket, an iron disc, a silver buckle, two fragments of a glass bead, and fragments of an iron knife (?). Based on the archaeological material this grave can be dated to the last third of the 4th century CE to the beginning of the 5th century CE. Radiocarbon dating, suggesting an earlier date, placed the burial from the second third of the 3rd century CE to the middle of the 4th century CE.

***CSK-101: Grave No. 101***

The skeleton of a young adult male with Europid characteristics and Mongoloid features on the skull was found in the S–N oriented grave, without known grave goods.

***CSK-126: Grave No. 126***

This robbed burial, oriented in an S–N direction, contained the fragmented skeleton of a middle adult female with Europid characteristics on the skull. Grave goods included a glass cup, a ceramic vessel, and two glass beads.

***CSK-127: Grave No. 127***

The fragmented skeleton of an old adult female was found in the robbed burial, near the west wall of the grave, oriented in an S–N direction. The recovered artifacts consisted of an iron brooch, an iron bracelet (?), an iron knife, fragments of an iron awl, two silver brooch (Prague type), a bone needle case, and a cylindric bone item. This grave can be dated to the end of the 4th century CE to the first half of the 5th century CE.

***CSK-135: Grave No. 135***

This S–N oriented robbed burial contained the fragmented skeleton of a middle adult male, with the bones thrown in the southwestern end of the pit. Iron coffin clamps were found in the grave pit. Grave goods included a Roman silver coin (Commodus denarius), a ceramic bowl, and the fragment of an iron spearhead.

***CSK-137: Grave No. 137***

The skeletal remains of an old adult male with Europid characteristics on the skull, thrown together into a pile, were found in the grave. Recovered artifacts included a silver buckle, a small silver crescent-shaped earring, and a fragment of an iron spear. Radiocarbon measurement, supporting an earlier date, placed the burial between the first third of the 3rd century CE and the first third of the 4th century CE. Based on the archaeological material this grave can be dated to the last third of the 4th century CE to the beginning of the 5th century CE.

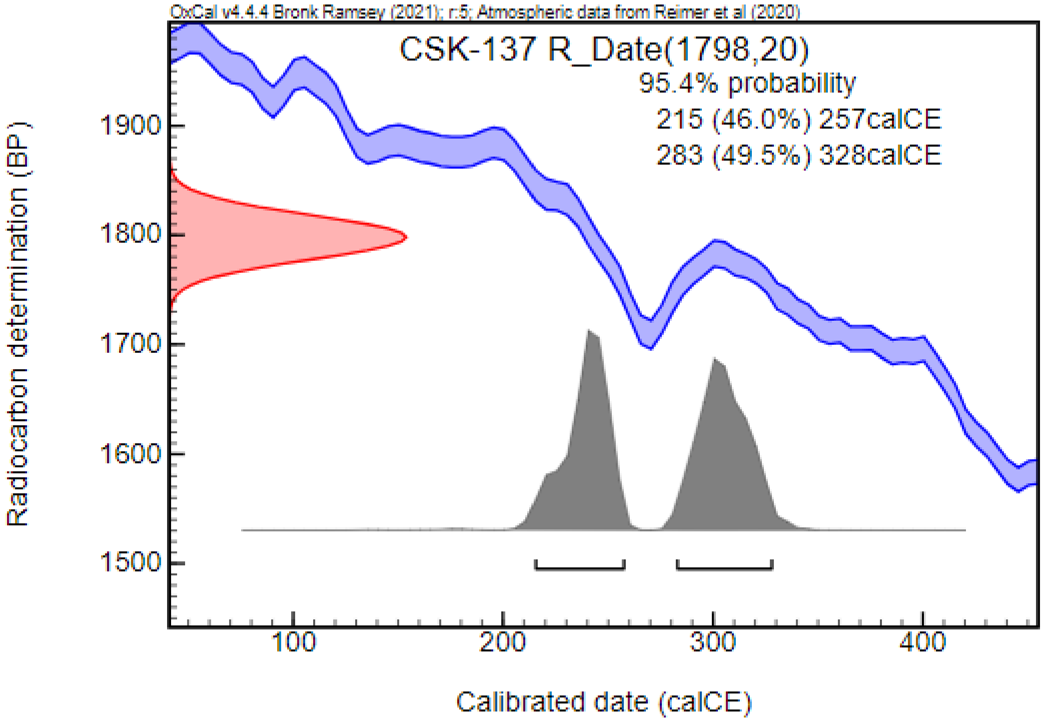


#### Dormánd – Zsidótemető (DZS; Heves County, Hungary) [74]

In 2005 and 2006, preventive excavations led by László Domboróczki were conducted at the western border of the village of Dormánd, on the western bank of the Laskó River backwater. The excavations uncovered settlement objects dating to the Late Bronze Aga and Iron Age, as well as traces of a trench from the Late Medieval or Early Modern Period. In addition, 21 graves from a Sarmatian cemetery were discovered. Although the archaeological and anthropological analyses are still in progress and only preliminary data are available, this cemetery is listed among the earliest Sarmatian-period archaeological sites [75]. We could include 9 samples in our analysis.

***DZS-1: Grave No. 1***

This grave contained the skeletal remains of a sub-adult (juvenile) individual. The grave goods are unknown.

***DZS-3: Grave No. 3***

The skeletal remains of an adult male were uncovered in this grave. The grave goods are unknown.

***DZS-5: Grave No. 5***

This grave contained the burial of a sub-adult (juvenile) individual with female characteristics on the skeleton. The grave goods are unknown.

***DZS-6: Grave No. 6***

Fragmented skeletal remains of an adult individual were discovered in this grave. The grave goods are unknown.

***DZS-41: Grave No. 41***

This was the burial of an adult male. The grave goods are unknown. Radiocarbon dating, consistent with the archaeological findings, placed the burial to the early Sarmatian period, between the first third of the 1st century CE and the first third of the 2nd century CE, with the possibility of dating to the mid-2nd century CE (1.3%).

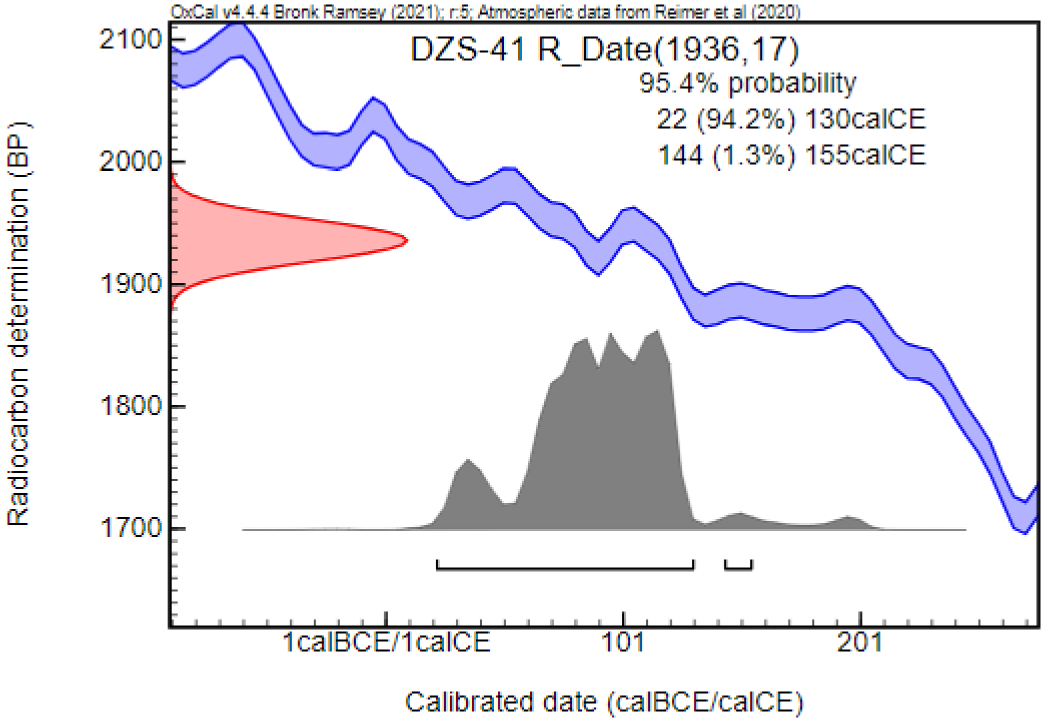


***DZS-42: Grave No. 42***

Only bone fragments of an adult individual were uncovered in this grave. The grave goods are unknown. Radiocarbon dating, supporting the archaeological findings, placed the burial to the early Sarmatian period, between the beginning of the 1st century CE and the first third of the 2nd century CE, with the possibility of being dated to the last quarter of the 1st century BCE (1.2%).

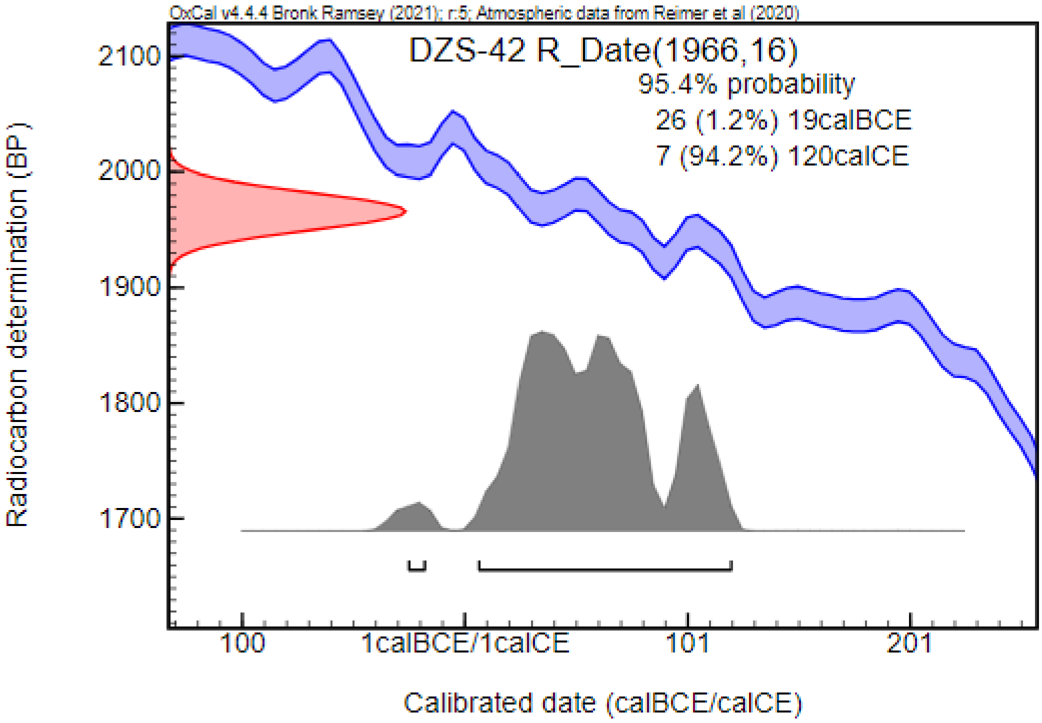


***DZS-43: Grave No. 43***

Fragmented skeletal remains of a sub-adult (*infantia* I or II) individual were uncovered in this grave. The grave goods are unknown.

***DZS-44: Grave No. 44***

This grave contained the burial of a sub-adult individual (*infantia* I). The grave goods are unknown.

***DZS-48: Grave No. 48***

Fragmented remains of a sub-adult (*infantia* I or II) individual were recovered from this grave. The grave goods are unknown.

#### Füzesabony – Kastélydűlő (FKD; Heves County, Hungary) [76]

Adjacent to settlements dating back to the Sarmatian period and the Middle Ages, a Sarmatian-period cemetery was discovered during a rescue excavation led by Csilla Farkas in 1997. Although only seven graves were documented, the burial ground is considered only partially excavated. The graves were situated approximately 10 meters from each other, forming two rows in NW–SE directions. The skeletons were predominantly found in a supine position with straightened extremities, oriented in the SN direction with minor deviations toward S–SW in the eastern part of the cemetery (group I) and S–SE in the western group of graves (group II). Variations were observed in the form and size of the grave pits, which were assumed to be correlated with the use of wooden coffins. The use of coffins was suspected in three graves based on the traces of organic materials and the position of the skeletal remains. Remarkably, the burials, a rare phenomenon in this period, were not affected by contemporary robbery. The archaeological material consisted of jewelry (e.g., bracelets and beads), clothing elements (e.g., brooches), tools (e.g., spindle-whorl), and vessels. The burials in the southern row of graves were wealthier and some contained jewelry and cloth fittings made of gold. Conversely, the burials in the northern row contained fewer artifacts, suggesting social differences between the two groups. The number of mirrors found in the graves is high compared to the low number of burials. The artifacts and burial customs indicate connections with the Sarmatians of the Black Sea (e.g., clothing elements and jewelry), the Roman Empire (e.g., certain vessel types, fibulas, and mirrors), and the native population of the Carpathian Basin, such as Dacians and Celts (e.g., vessels). The cemetery was dated to the turn of the 1st–2nd centuries CE and the beginning of the 2nd century CE, connected to the 2nd or 3rd migrating wave of the Jazygs. Anthropological investigation conducted on the skeletal remains by Antónia Marcsik revealed a female surplus in the cemetery, with six females and only one male identified in the series. Additionally, one juvenile/young adult, one young adult, and five middle adult individuals were identified. However, the low qualitative and quantitative preservation of the bones limited further analysis. Three burials were available for analysis.

***FKD-60: Grave No. 60***

The skeletal remains of a young-adult female (23–40 years) buried in a supine position with straight arms, oriented in SW with minor deviations in SSW, were found in a rectangular grave pit. The burial contained a pair of golden earrings with hook and loop ends, a bronze needle with hooked end, a string of carnelian beads (a total of 41), flower-shaped golden cloth ornaments (a total of 10), a tinned bronze mirror, a spindle whorl, triangle-shaped press golden clothing ornaments (a total of 15), a pair of bracelets made of carnelian beads, and a jug. Radiocarbon dating generally confirmed the archaeological analysis, placing the burial mostly between the $1^{st}$ and $2^{nd}$ centuries CE.

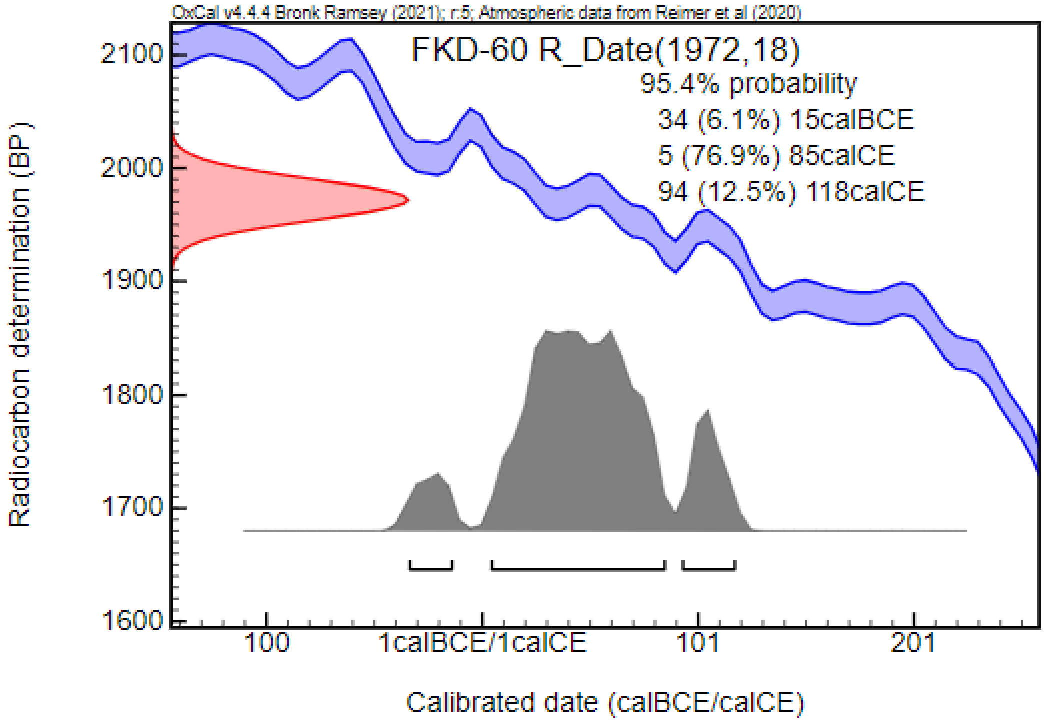


***FKD-140: Grave No. 140***

The oval grave pit contained the fragmented skeleton of a middle adult female in a supine position, oriented to SN with minor deviation to SE. Two spindle whorls, fragments of a tinned bronze mirror, iron plates of a chest (?), a strongly profiled bronze fibula, glass beads, and two bowls were found in the burial.

***FKD-150: Grave No. 150***

This was the burial of a juvenile or young adult female (16–20 years). The skeletal remains were found in the rectangular grave pit in a supine position with straight arms, oriented in SN with minor deviations to SE. The grave goods included a pair of golden earrings with funnel end, a bronze needle with hooked end, a spindle whorl, glass-, amber- and gold beads in the neck region, a gold ornament (pendant?), a tinned bronze mirror, a pair of bronze bracelets, glass beads around the left wrist, unidentified fragmented iron artifacts, a pyxis, and further beads (e.g., glass and carnelian) from the soil. Radiocarbon dating has placed the burial from the end of the $1^{st}$ century CE to the beginning of the $3^{rd}$ century CE.

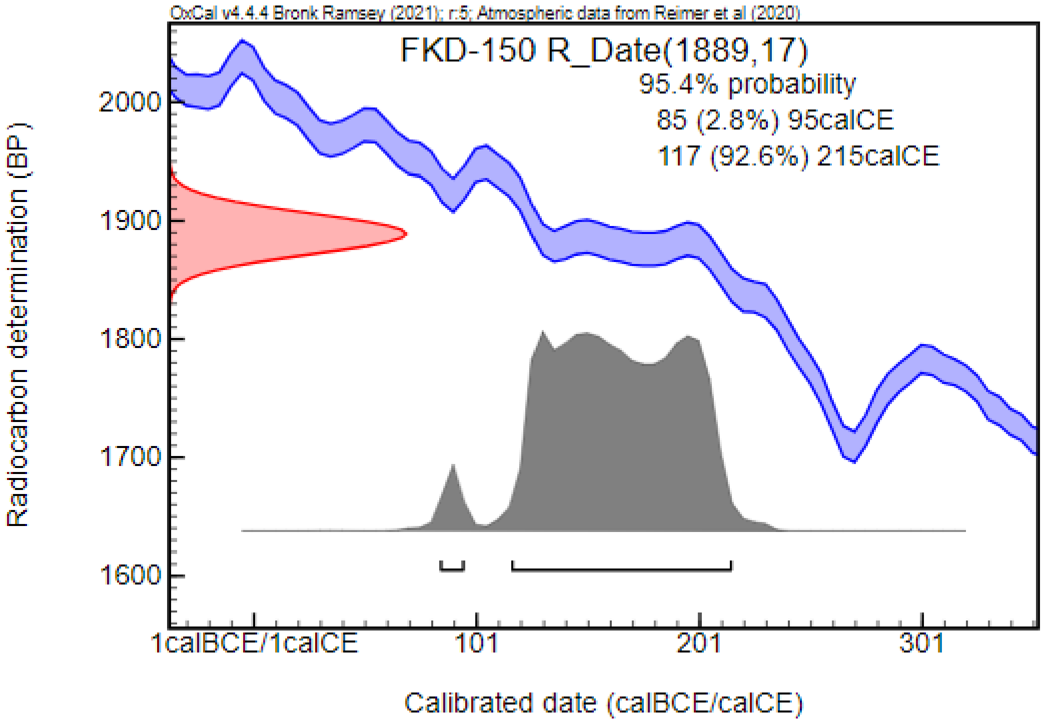


#### Hévízgyörk (HGY; Pest County, Hungary) [77,78]

In the course of preventive archaeological research led by Edit Tari related to the renovation of the medieval church of Hévízgyörk, two Sarmatian graves were discovered in 1985 and 1986. These burials were found in superposition with – and partially destroyed by – the foundation walls of the church and burials dating to the Medieval period. It is presumed that additional graves were destroyed during the construction of the church in the Middle Ages. The skeletons were found in a supine position, oriented to NW–SE, which is considered unusual among the Sarmatians. The archaeological material composed of jewelry and clothing elements (e.g., beads, brooches), tools and other objects (e.g., knife, fire-lighting tools, wooden chest, mirror), vessels made of glass and clay. In addition weapons (e.g., a sword), and horse riding-related deposits (e.g., iron bit, girth buckle) were present. The burial customs and grave goods suggest extensive connections with the Sarmatians of the Pontic region, German tribes of the north (i.e., Przeworsk culture), and the Roman Empire (i.e., Pannonia province). The burials were dated to the second half of the 2nd century CE. The skeletal remains at this site were examined by Márta Ferencz. The qualitative and quantitative preservation of the bones was generally low, marked by fragmented skulls and postcranial elements. The analysis identified the skeletons as a young adult female and a middle adult male. Remarkably, traces of surgical trephination were detected on one of the skulls, a rare phenomenon for this era. Unfortunately, only one sample was deemed suitable for analysis.

##### *HGY-14: Grave No. 14*

The NW–SE oriented grave was overlapped and partially destroyed by the foundation of the church and a medieval grave. The partially disturbed burial contained the skeletal remains of a young adult female in a supine position. The grave-goods included two glass beads at the left side of the neck region, amber beads spanning from the left clavicle to the pelvic region, coral beads, and a chalk bead near the mandible. In addition, further glass and chalcedony beads were discovered around the disturbed chest region. Notably, the grave contained a one-piece bronze fibula with a foot bent under, and a bronze knee fibula. Several additional artifacts were recovered from various parts of the burial. These included a total of 23 beads (made of glass, crystal, and chalcedony) near the left femur, an iron knife and an iron awl near the left femur, approximately 1000 glass beads around the feet, and a purple glass flagon near the right ankle have been found. Moreover, iron plates and bronze nails of a wooden chest that contained a vessel, a bronze mirror, a spindle whorl, and two amber beads were found in the western part of the grave pit. Radiocarbon dating has placed the burial from the first third of the 2nd century to the first third of the 3rd century CE.

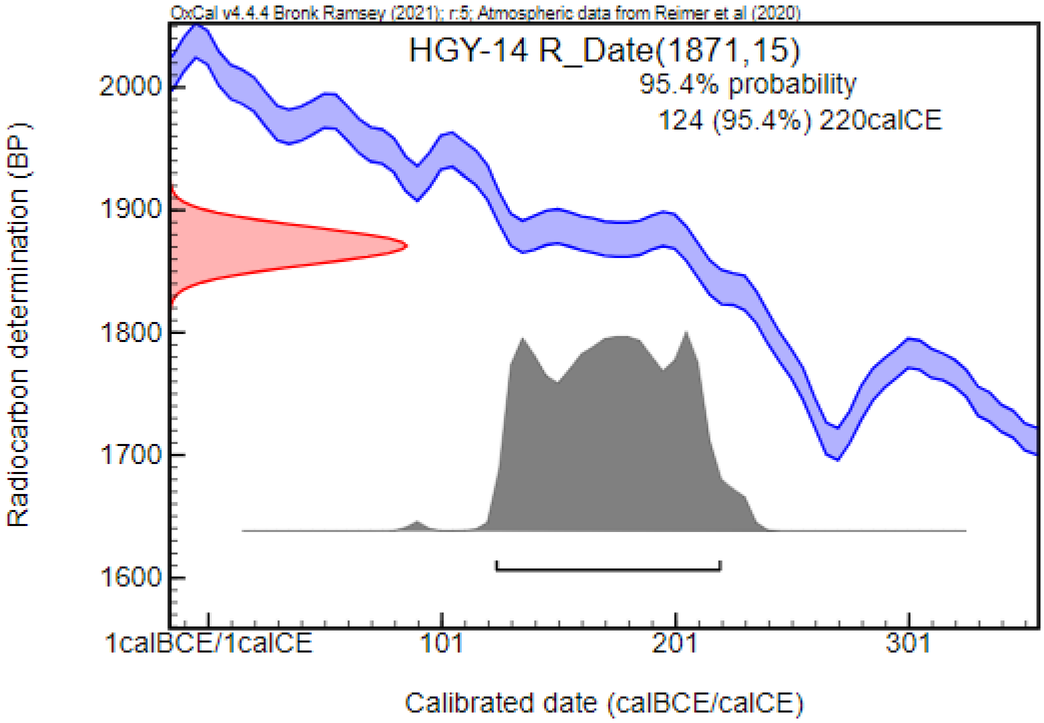


#### Hódmezővásárhely – Fehértó (HVF; Heves County, Hungary) [79,80]

A Sarmatian burial was discovered near Fehértó lake in the close vicinity of Hódmezővásárhely in 1895. Decades later, in 1943, excavations were conducted at the same location, revealing a Sarmatian cemetery with 22 graves. The cemetery was found approximately 200–250 m away from the first grave, and, presumably, they belonged to the same burial ground. However, it was not allowed to conduct field survey in the area between them; thus, the cemetery is considered only partially excavated. Most of the burials were oriented in S–N directions with minor deviations toward SSW–NNE in 8 cases and SW–NE in 2 cases. Some of the graves were affected by contemporary robbery. The archaeological material contained jewelry (e.g., beads and bracelets), cloth fittings (e.g., beads, brooches, belt buckles), tools (e.g., fire-lighting tools, knives, awls, spindle whorls), clay vessels, and Roman coins. The cemetery was dated from the 1st century/beginning of the 2nd century CE to the 3rd century CE. The anthropological analysis, conducted by Lajos Bartucz, revealed that the skeletal remains of 21 individuals was available for analysis. The state of preservation of the bones is generally low with fragmented skulls and postcranial elements. The series is composed of 5 sub-adults and 16 adults. Among the adults, 8 females and 8 males were identified. Europid characteristics were described on the observable skulls with both dolichocran (Nordoid and Mediterranean) and brachycran (Pamyrian, Dinarian, and Turanian) types being present in the population. A total of 8 samples could be involved in the analysis.

##### *HVF-2: Grave No. 2*

The burial, oriented in SW–NW, contained the skeletal remains of a young-adult female with Europid (East European or Turanian features) characteristics on the skull. A string of beads near the neck, a spindle whorl, a clay vessel, approximately 200 glass beads around the ankles, further beads near the hip and thorax, an iron knife, and an iron awl have been found in the grave.

##### *HVF-3: Grave No. 3*

The skeleton of a young adult female with Europid characteristics (Pamyrian features) on the skull was found in the grave oriented in SSW–NNE. The burial contained a string of glass beads near the neck, a ceramic bowl, bronze and silver rings, different types of beads near the upper body with a small bronze bell, a spindle whorl, fragments of an iron knife, an enamelled brooch made of five discs and four rings connected to the axis by spokes, a small silver box, and approximately 600 glass beads around the ankles. This grave can be dated to the second half of the 2nd century – and the first half of mid-3rd century CE.

***HVF-4: Grave No. 4***

The skeletal remains of a middle adult male with Europid characteristics (Mediterranean features) on the skull were found in the grave oriented in S–N directions. The archaeological material composed of an iron knife, an iron awl, further unknown iron shards, and a silver coin emitted by Emperor Marcus Aurelius. Radiocarbon measurement dated the burial from the end of the 1st century CE to the beginning of the 3rd century CE.

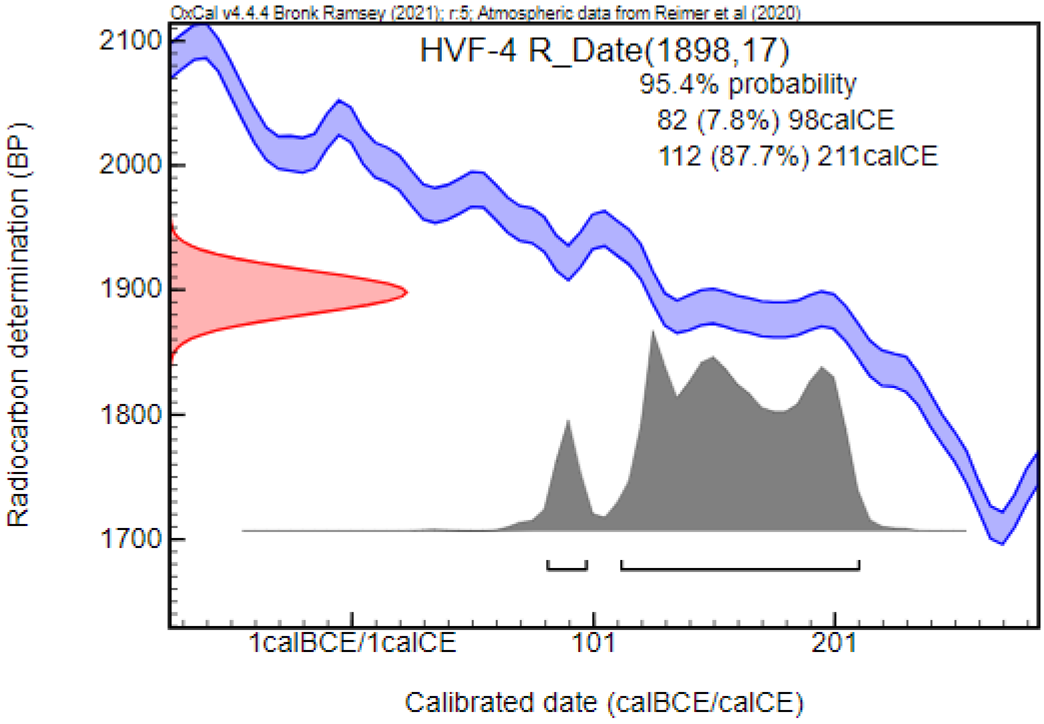


***HVF-8: Grave No. 8***

The SSW–NNE oriented grave contained the skeleton of a young adult female with Europid characteristics on the skull. The grave goods included a hande-made cermaic vessel, approximately 650 glass beads around the ankles, an iron knife, different types of beads near the hip and the upper body, and a bronze brooch with a returned foot. This grave can be dated to the second third and half of the 3rd century CE.

***HVF-10: Grave No. 10***

The burial was severely robbed and disturbed, containing only fragmented human bones. The skeletal remains belonged to a middle adult male with the skull showing Europid characteristics (possibly East European or Turanian features).

***HVF-15: Grave No. 15***

The grave, oriented in S–N directions, contained the skeleton of a middle adult female with Europid characteristics (East European or Turanian features) on the skull. The inventory of the burial consisted of a clay vessel, approximately 800 glass beads around the ankles, an iron knife, a spindle whorl, a string of beads around the neck, a bronze fibula with strong profile, and different types of beads near the upper body. This grave can be dated to the beginning and first half of the 2nd century CE.

***HVF-17: Grave No. 17***

Skeletal remains of a middle adult female with Europid characteristics (East European or Turanian features) on the skull were found in the grave, oriented in S–N directions. A ceramic vessel, a bronze knee brooch, a bronze needle, a spindle whorl, and unknown iron fragments were recovered from the burial. This grave can be dated to the second half of the 2nd century – and the first third of mid-3rd CE.

***HVF-21: Grave No. 21***

The grave, containing the skeleton of a middle adult male with Europid characteristics (Mediterranean, Nordoid features) on the skull, was oriented in S–N, and the bottom of the pit was covered with an unknown white material. The grave goods consisted of a ceramic bowl, a fragmented iron knife, a ‘Sarmatian-type’ belt buckle, an iron awl, fire-lighting tools, and a bronze crossbow Sarmatian type brooch. Radiocarbon measurement dated the burial from the second third of the 2nd century CE to the first third of the 3rd century CE.

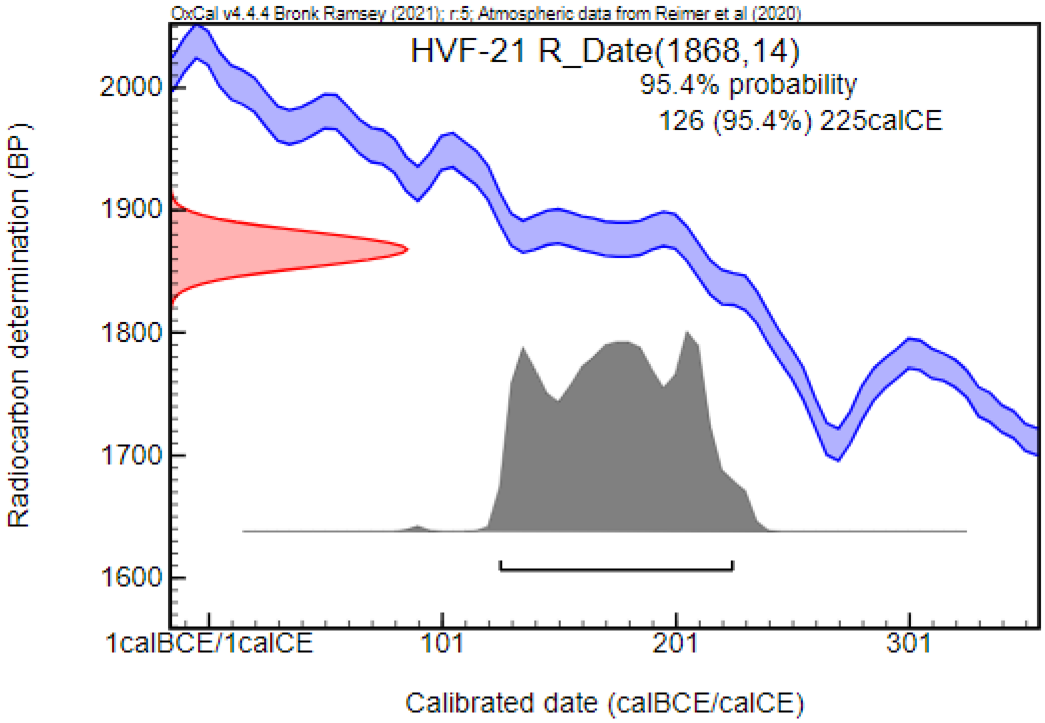


#### Hódmezővásárhely – Kenyere-ér, Bereczki-tanya II. [38,81]

A preventive excavation, lead by Sándor Varga, was conducted in conjunction with the construction works of the Hódmezővásárhely bypass section of main road #47 between 2015 and 2017. While materials from various periods (e.g., Avar Age) were documented, the majority of the findings were related to the Roman Age, including a Sarmatian-period settlement and three cemeteries. The three cemeteries, situated on small ground elevations, were separated from each other; however, their exact extent is unknown as they are only partially excavated. The anthropological analysis was conducted by Olga Spekker and colleagues, and the general description of the series is yet to be published. The state of preservation of the bones was generally low, with mostly fragmented skulls and postcranial bones, greatly limiting the analysis and evaluation of data.

From the summary of the preliminary results, it is known that among the nine graves of the first cemetery, two or three were encircled by a ditch. The grave, pits oriented in an S–N direction, were rectangular with rounded corners. The archaeological material derived from the burials includes jewelry (e.g., earrings, pendants, bracelets, and beads), cloth fittings (e.g., brooches, beads, and belt buckles), tools (e.g., knives), ceramic vessels, and Roman coins. In the anthropological series of the first cemetery, three sub-adults and five adults were identified. Among the adults, two females were determined. Additionally, one fragmented skeleton showed female characteristics, and another fragmented skeleton exhibited male characteristics. To the south of the first cemetery, 18 burials of the second cemetery were excavated. Smaller sections of possible ditches related to burials were also found. Seven or eight of the graves were encircled by ditches. The rectangular grave pits, oriented in an S–N direction, formed approximately four rows in a W–E direction. Most of the burials were affected by contemporary robbery. The archaeological material, characteristic of the Sarmatian period, included jewelry (e.g., earrings, bracelets, and beads), cloth fittings (e.g., belt buckles and beads), mirror fragments, tools (e.g., knives, spindle whorls), ceramic vessels, and Roman coins. In addition, one burial contained weapon equipment and spurs, indicating possible connections with neighboring northern Germanic tribes. The anthropological series of the second cemetery consisted of 4 sub-adults and 11 adults. In the sub-adult group, the skeletal remains of a juvenile individual showed female characteristics. Furthermore, four individuals were described with female characteristics and three individuals were determined as males. Only three skulls were available for morphological analysis, and all of them showed Europid characteristics. In three additional cases, shoveling of the incisor teeth was documented, which is rather associated by scholars with skulls exhibiting Mongoloid morphological characteristics. The anthropological data suggests a complex, heterogenous population with local and eastern origins.

Three graves of the third cemetery were excavated approximately 150 meters west of the second cemetery. One of them was encircled by a ditch, but it was completely robbed. Limited anthropological data is available concerning the series of the third cemetery due to the low number of individuals and the poor preservation of the skeletal remains. Two adult individuals were identified in the series, and among them, one skeleton showed female characteristics. While bone preservation prevented the morphological and metric analysis of the skulls, shoveling of the incisors was detected in one case. We could involve one sample from the second cemetery in our analysis.

***HKB-309: Grave No. 309***

This was the robbed burial of a middle or old adult (>50 years old) male with Europid characteristics on the skull and a robust skeleton. The grave, oriented in an S–N direction, was encircled by a ditch. The archaeological material consisted of a bronze strap distributor, an iron spearhead, iron elements of a shield, a belt buckle, a strap end, a pair of iron spurs, and a silver Roman coin dated to the 2nd century CE. While the observed burial customs are consistent with the other Sarmatian period graves of the cemetery, the grave goods are characteristic of the northern Germanic tribes such as the Quads, Marcomanni, and Vandals. Radiocarbon measurement dated the burial from the second third of the 2nd century CE to the first half of the 3rd century CE.

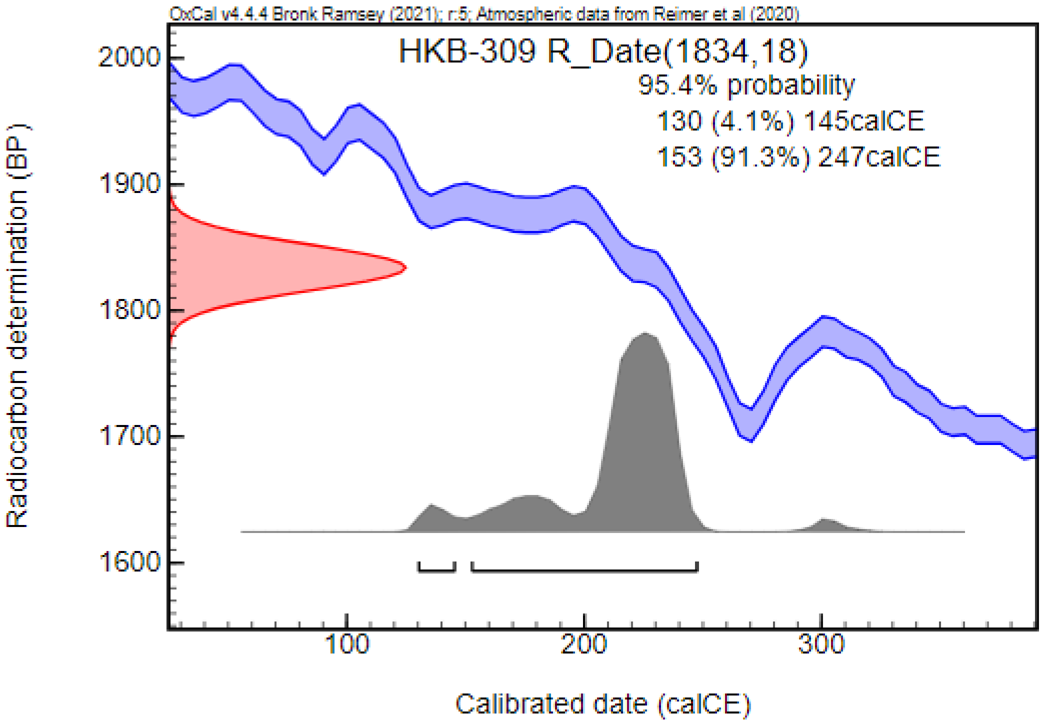


#### Kapolcs (KAP; Veszprém County, Hungary) [82,83]

Two burials, oriented in a W–E direction, were uncovered during a rescue excavation led by Alán Kralovánszky in 1979. Both graves were significantly deep, with the skeletal remains recovered from a depth of more than 2 meters. One of the burials was richly equipped with jewelry (e.g., earrings, necklaces, bracelets, beads, pendants, finger rings, and pins) and cloth fittings (e.g., a belt buckle) made of precious metal, a mirror, a bone comb, and a ceramic vessel, while the other contained silver pins possibly connected to the clothing. The burials were dated to the second half of the 5th century CE and were presumably associated with Eastern German groups. The gilded silver belt buckle from grave No. 1 showed analogies with artifacts found in the territories of Italy, Germany, and Romania. Kinga Éry conducted the anthropological analysis. The state of preservation of the bones is moderate, with fragmented skulls. Both individuals were identified as females, one of them belonged to the sub-adult (juvenile) age group, while the other individual was estimated as an adult. The skulls showed Europid characteristics, and traces of slight artificial deformation were detected on both. We could include one sample in our analysis.

***KAP-2: Grave No. 2***

The burial, oriented in a W–E direction, contained the skeleton of a middle adult female with Europid characteristics and traces of slight artificial deformation on the skull. The grave pit was deep, with the skeletal remains recovered from a depth of 215 cm. Two silver pins were found near the chest.

#### Kecskemét – Mercedes RL 15 (KM; Bács-Kiskun County, Hungary) [84]

Following extensive field research, including area excavations and excavations using the sondage method, in 2008, 2009, and 2013, rescue excavations were conducted by the team of Katona József Museum in collaboration with Ásatárs Ltd. and Salisbury Ltd. in 2017 and 2018 at the construction site of the Mercedes Factory. As a result, a large and contiguous area was uncovered, revealing Bronze Age, several Sarmatian-period, Avar-age, and two Medieval settlements, along with Avar-age and two Sarmatian-period cemeteries dating to the 2nd–4th-centures CE and 4th–5th centuries CE. These findings were distributed across multiple sites in the excavation area. The archaeological and anthropological investigations are ongoing, and only preliminary data are available.

Site RL15 was excavated by Nikoletta Lukács in 2017, revealing a total of 1151 features, including a Bronze Age palisade wall, Sarmatian and Medieval settlements, and four Sarmatian graves. One of the Sarmatian grave was found in the eastern part of the site, in the pit of a house, while the other three burials were discovered in the western part of the site, positioned next to each other. We could include two samples in our analysis.

***KM-1005: Grave with SNR No. 1005***

The fragmented skeletal remains of an adult female were found in a supine position in this grave, oriented in a W–E direction. Traces possibly belonging to a coffin or shroud were discovered in the pit. Grave goods included a coral bead near the neck, a silver brooch, a bronze mirror, a ceramic vessel, and three additional coral beads near the hip. Radiocarbon dating placed the burial in the Sarmatian period, between the middle of the 3rd century CE and the second half of the 4th century CE.

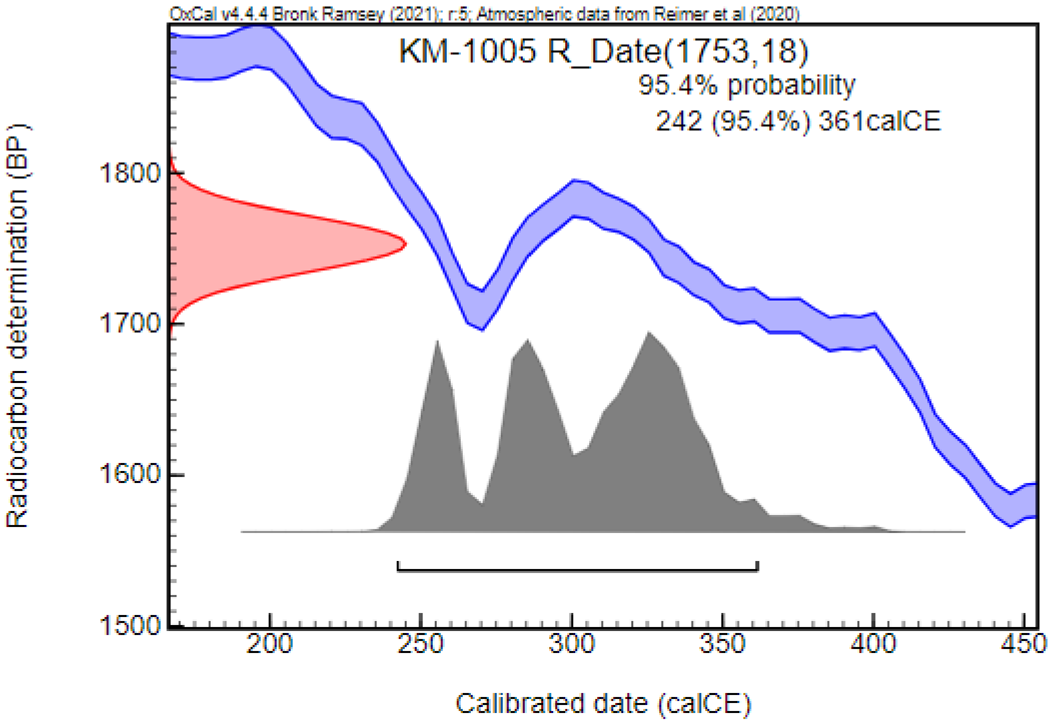


***KM-747: Grave with SNR No. 747***

This SW–NE oriented burial was discovered in the eastern part of the site, in the pit of a house. The fragmented skeleton of an adult female was uncovered in a supine position, accompanied by two silver earrings, a string of coral, amber, glass, and silver beads, a silver box brooch, a silver pendant, and a silver one-piece brooch. In addition, two narrow, unidentified iron objects, an amber bead near the right wrist, a spindle whorl, a ceramic vessel (along with a shard of another vessel found in the soil), an iron knife, and an iron awl were also found.

#### Kecskemét – Mindszenti-dűlő I, Mercedes RL 11 (KM; Bács-Kiskun County, Hungary) [85]

Following extensive archaeological investigation, including field survey and preventive excavations in 2008 and 2009, rescue excavations led by Gábor Wilhelm were conducted in 2017 at site RL11. These excavations uncovered numerous settlement features, mostly dating to the Sarmatian period and the Avar Age. In addition, Sarmatian period graves were discovered. Although the archaeological and anthropological analyses are still in progress, the burials have been preliminary dated to the 3rd–4th centuries CE. We could include one sample in our analysis.

***KM-3679: Grave with SNR No. SNR3679/B***

This NW–SE oriented grave contained a double burial, supposed to be of an adult female and an adult male. The skeletons were found in a supine position, holding each other's hands. Grave goods, including an iron knife and a ceramic bowl, were recovered only in association with Skeleton B, which was located in the southern part of the grave. Radiocarbon measurement, partially overlapping with the archeological findings, placed the burial in the late Sarmatian period, from the second third of the 4th century CE to the first third of the 5th century CE, with the possibility of being dated to the last third of the 3rd century CE (13.1%).

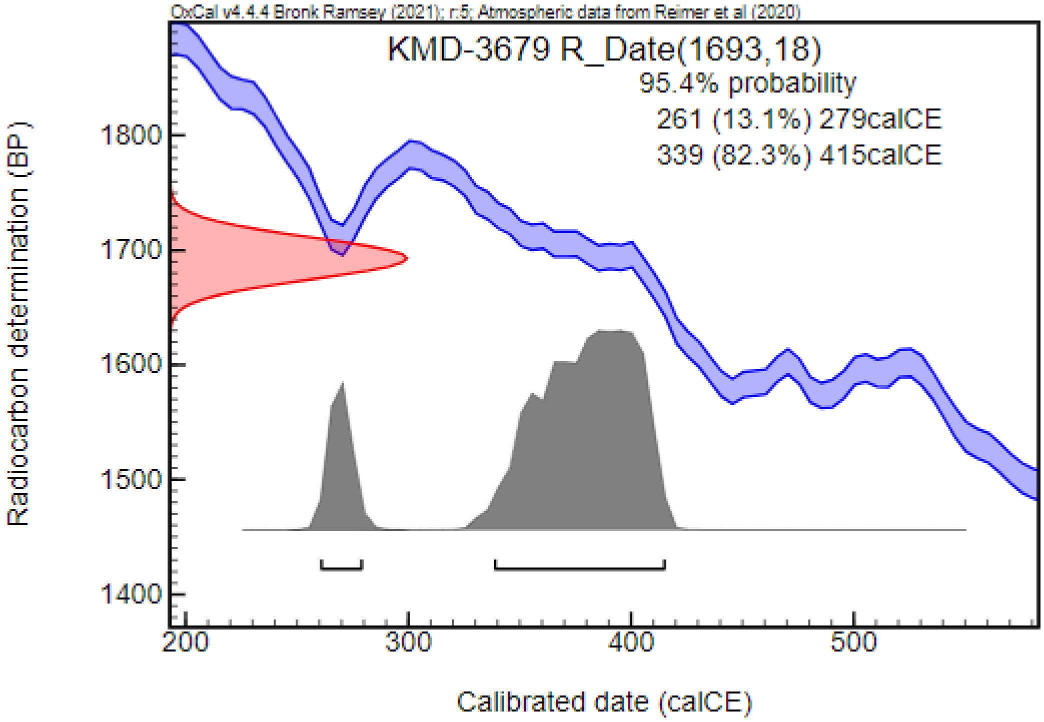


#### Kecskemét – Mindszenti-dűlő II, Mercedes RL12 (RL 17) (KM; Bács-Kiskun County, Hungary) [84]

In 2018, excavations led by Bernadett Ny. Kovacsóczy continued at site RL 17, revealing that it is continuous with RL 12. In addition to Avar Age settlement features, 57 graves of a Sarmatian-period cemetery, including graves surrounded by ditches, were discovered, along with three additional burials possibly dating to later periods, which were positioned in or on the ditches surrounding the Sarmatian burials. The archaeological and anthropological investigations are ongoing, and preliminary data are available. Despite most of the burials being disturbed by contemporary robbing, numerous artifacts were still recovered, including jewelry (e.g., bracelets), clothing elements (e.g., brooches and beads), and a mirror. Furthermore, all burials contained ceramic vessels of various forms and quality, primarily found near the legs. In some graves, traces of a coffin were discovered, and in one grave, a wooden burial chamber was documented. The Sarmatian cemetery has been dated between the 2nd and 4th centuries CE. We could include one sample in our analysis.

***KM-156: Grave with SNR No. 156***

This ENE–WSW oriented grave contained the burial of an adult female. Due to contemporary robbing, the bones and artifacts were mostly found in a secondary position. The recovered grave goods included a spindle whorl, beads made of amber, calcedony, calcite, and coral, as well as a bronze pendant. Additionally, numerous beads, possibly from the clothing, were found near the original location of the lower legs, though they were scattered without any clear pattern.

#### Kiskőrös – Fekete halom (KFH; Bács-Kiskun County, Hungary) [86]

A rescue excavation led by György V. Székely was conducted following the discovery of a Sarmatian period settlement and burial ground during construction works. Six graves were uncovered, two of which were surrounded by a ditch. Grave No. 1 differed from the others in both location (80 meters away from the other burials) and orientation, and no grave goods were recovered from it, raising questions about its association with the Sarmatian cemetery. Most of the burials had been robbed. The grave goods included ceramic vessels, an iron knife, jewelry (e.g., a string of beads), brooches, and a spindle whorl. The burials were dated to the 3rd and 4th centuries CE.

The state of preservation of the bones was generally poor, and skeletal remains from three burials (excluding grave No. 1) were suitable for anthropological analysis, conducted by our colleagues (LK, OS, BT) specifically for this study. In one of these burials (grave No. 4), skeletal remains of two individuals were identified: one sub-adult (infantia I or II) and one older, indeterminate individual. The other two burials contained the skeletons of two adult individuals, and we included these latter two samples in our study.

***KFH-2: Grave No. 2***

The fragmented skeletal remains of an adult male, oriented in a SE–NW direction, were found in this robbed burial. The grave goods included an iron knife, an unidentified iron object, fragments of a stone blade, and a ceramic vessel.

***KFH-3: Grave No. 3***

This robbed grave, oriented in a SE–NW direction, contained the fragmented skeletal remains of an adult, possibly female individual without any known grave goods.

#### Kiskundorozsma – Subasa 26/78 (KDS; Csongrád County, Hungary) [87,88]

Preventive excavation was carried out in conjunction with the construction of the M5 highway section near Kiskundorozsma. Alongside settlement objects and burials spanning various periods (e.g., Iron Age, Hungarian Conquest period, Árpádian-Age), sixty graves of a Sarmatian-period cemetery were uncovered. The graves, of varying sizes, were oriented in a S–N direction with minor deviations to SW–NE in some instances. Sixteen of the graves were surrounded by a ditch, but all had been looted. Approximately half of the graves without ditches were also affected by contemporary robbery. The deceased were laid to rest in a supine position with straightened arms and legs. In a few cases, traces of coffins were detected (i.e., metal parts). Preliminary findings indicate that the archaeological material, characteristic of the Sarmatian period, included jewelry (e.g., torques, pendants, beads, and bracelets), cloth fittings (e.g., brooches, belt buckles, and beads), cereamic vessels, tools (e.g., iron knives, spindle whorls, needles, and awls), and Roman coins dating from the 1st–4th centuries CE. While the partially excavated cemetery was initially dated to the 4th century CE based on a burial in the center, equipped with a coin minted between 293 and 305 CE, reconsideration of archaeological data and radiocarbon data in this paper suggest that the burial ground was already in use in previous centuries. The anthropological analysis was conducted by László Paja and his colleagues. Of the 60 excavated graves, the skeletal remains of 57 individuals were available for analysis due to the poor state of bone preservation. The series included a total of 15 sub-adult and 42 adults, with 17 females and 19 males identified in the population. While the morphological and metric analysis of the skulls was limited, the observable skulls were brachycranic and exhibited Europid characteristics. We could involve 4 samples in our analysis.

***KDS-121: Grave No. 121***

The burial of a young adult female, oriented in a S–N direction, contained a string of carnelian beads, a torque, a pair of bead strings around the wrists with pendants, a pair of bronze bracelets, and shards of a ceramic vessel. In addition, calcite, carnelian, and glass beads, two pieces of a bronze spiral wire, a “Scythian type” arrowhead on a bronze penannular ring, and a bronze pyxis were found near the left forearm. In connection with the clothing of the deceased, a brooch and approximately 800 glass beads around the legs were discovered in the grave.

***KDS-138: Grave No. 138***

The skeletal remains of an adult male were found in the grave, oriented in a SSW–NNE direction. The burial contained a silver torque ornamented with bronze beads, a silver pendant (lunula), a pair of silver bracelets, an iron brooch, a belt buckle, an iron knife, an iron awl, and two Roman coins from the 1st–2nd centuries CE.

***KDS-148: Grave No. 148***

The grave, oriented in a S–N direction, contained the skeleton of an adult male buried with an iron bracelet, a silver double-bracelet, a silver pendant, and a glass bead. Radiocarbon measurement dated the burial from the end of the 1st century CE to the beginning of the 3rd century CE, indicating that the burial ground could have been in use before the 4th century CE.

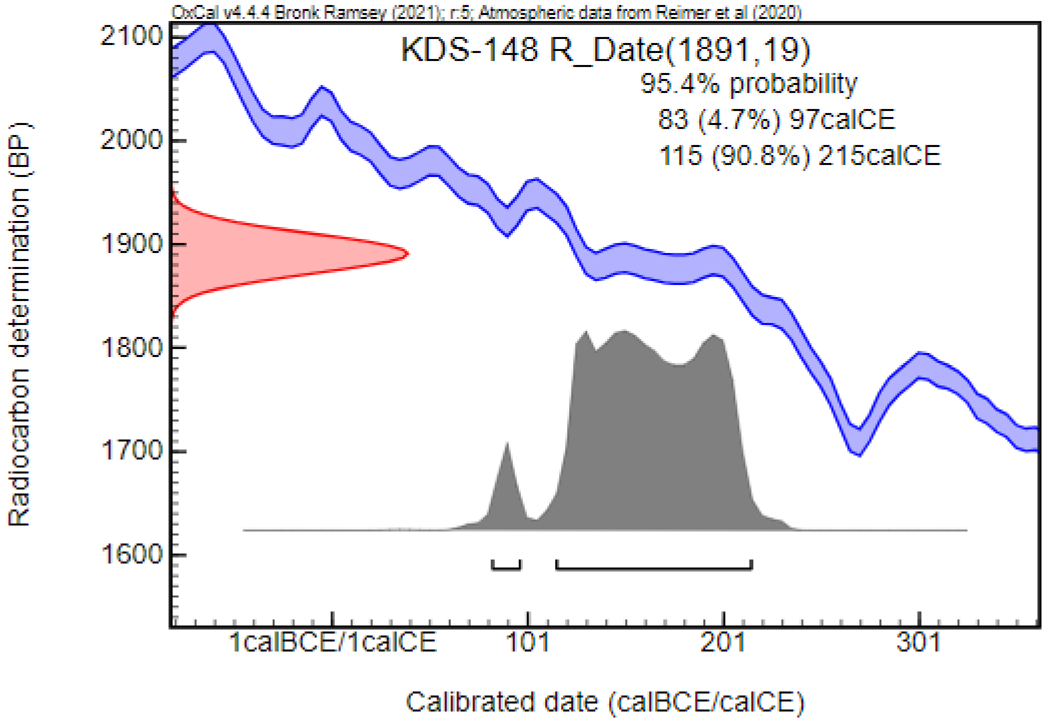


***KDS-260: Grave No. 260***

This was the burial of a young adult male, located in the center of the currently known burial ground. The known grave goods included a bronze and an iron belt buckle, an iron knife, and follis emitted by Constantinus Chlorus between 293 and 305 CE. The burial was dated to the 4th century CE.

#### Kunadacs – Községi temető (KAK; Bács-Kiskun County, Hungary) [86,89]

Two Sarmatian graves were discovered during the rescue excavation of a Bronze Age settlement led by Elvira H. Tóth in 1983. The burials, located on the western part of the site, were oriented in a S–N and SSW–NNE direction and both of them were robbed. The grave goods included animal bones, an arrowhead, a bronze plate, and fragments of iron objects. In 1984, further two Sarmatian graves were uncovered during a preventive excavation led by Valéria Kulcsár. These burials, oriented in a S–N direction, had also been robbed, containing only fragments of the skeletons, a glass bead, fragments of a knife, and a bronze strap end. The burials were dated to the Sarmatian period (possibly to the 3–4th centuries CE); however, a more precise dating is unachievable due to contemporary robbing. The bones were generally poorly preserved, and only skeletal remains of two adult individuals from the two burials found in 1984 were suitable for anthropological analysis, conducted by our colleagues (LK, OS, BT) specifically for this study. We could include one sample in our analysis.

***KAK-3: Grave No. 3***

The fragmented remains of an adult individual were recovered from the robbed grave oriented in a S–N direction. The grave goods included a glass bead and fragments of an iron knife. Radiocarbon measurement, partially confirming the archaeological dating, placed the burial in the Sarmatian period, from the second half of the 1st century CE to the beginning of the 3rd century CE.

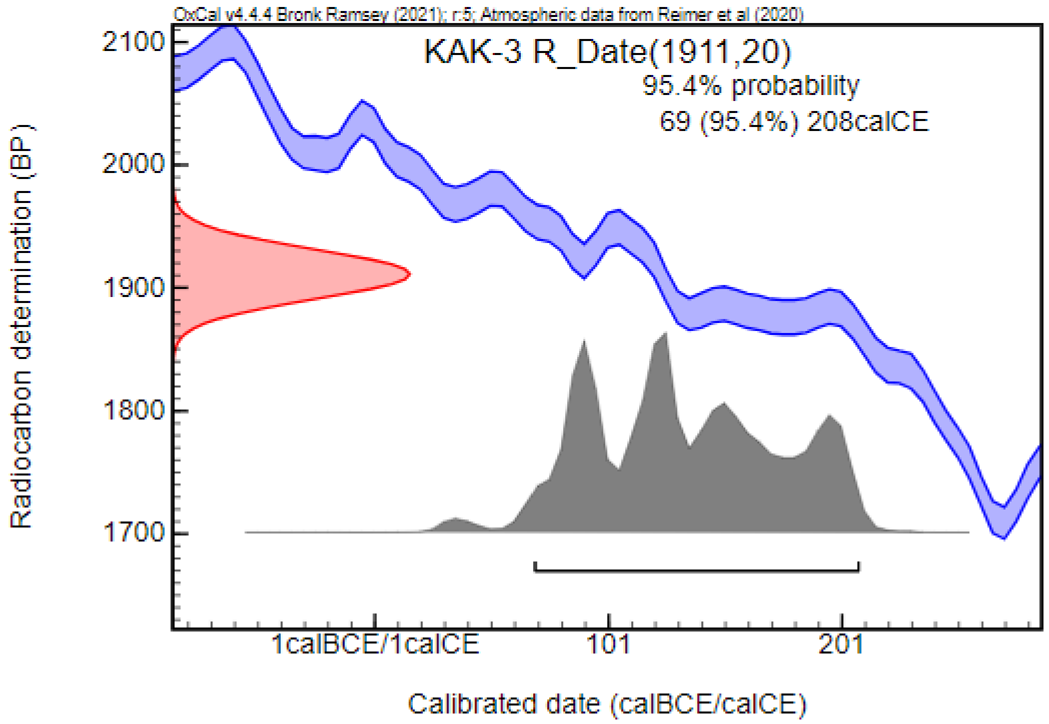


#### Kunszentmiklós – Nyakvágócsárda (KSZN-10321; Bács-Kiskun County, Hungary) [89]

In 1976, a burial oriented in a N–D direction was found during the digging of a pit. The grave contained a ceramic jug. The burial was dated to the Sarmatian period. The anthropological analysis, conducted by our colleagues (LK, OS, BT), in connection with this study, revealed that the skeletal remains belonged to a young adult male individual. Traces of a sharp force trauma with a possible surgical intervention are present on the skull vault being a rare phenomenon among the Sarmatians. Radiocarbon measurement confirmed the archaeological observation and dated the burial to the Sarmatian period, from the second third of the 1st century CE to the beginning of the 3rd century CE.

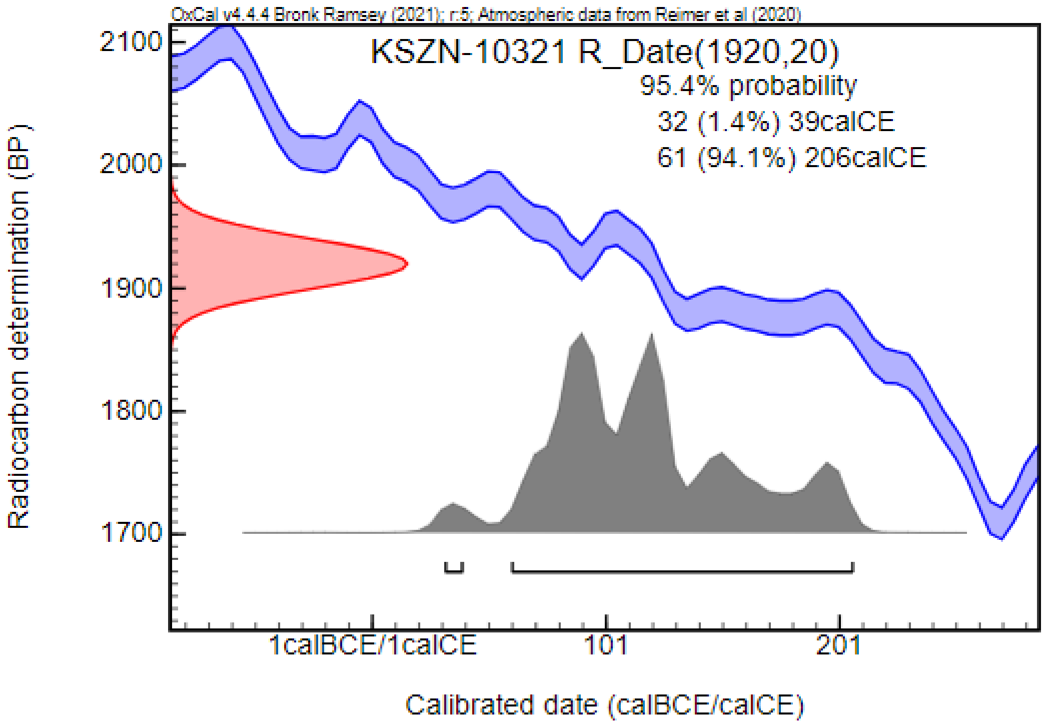


#### Lişcoteanca – Moş Filon (LMF; Brăila County, Romania) [90]

This multi-period site near Lişcoteanca features both settlement objects and graves and is situated on the northern terrace of the Călmăţui River. During excavations conducted between 1971 and 1986, six burials dating back to the Sarmatian period were unearthed from a Neolithic mound. These graves were primarily oriented in a NW–SE direction with occasional minor deviations, and in initial cases in a W–E and an opposite NE–SW direction. The deceased individuals were laid to rest in a supine position with their arms either straightened or bent. Grave goods discovered included only a few ceramic vessels, iron knives, several beads, and a brooch. The Sarmatian burials were mostly dated to the 1st and 2nd centuries CE based on Roman import artifacts. Three adults and two sub-adults were identified in the material. Anthropological investigation was carried out by Andrei Dorian Soficaru and Ana Ştefan. We could involve two samples in our analysis.

***LMF-6: Grave No. 6***

This grave contained the skeletal remains of a female adult individual in a supine position with bent arms, oriented in a W–E direction. Additionally, a bead with "eye" decoration, considered a Roman import artifact and three bone beads were discovered near the neck of the individual. The burial was dated to the 1st and 2nd centuries CE.

***LMF-7: Grave No. 7***

The skeletal remains of a male adult individual were found in this grave in a supine position with straightened extremities, and oriented in a NE–SW direction. The grave goods included a mug, a bronze brooch, and an iron dagger. Radiocarbon measurement dated the burial from the first third of the 1st century CE to the first third of the 2nd century CE, confirming the archaeological dating.

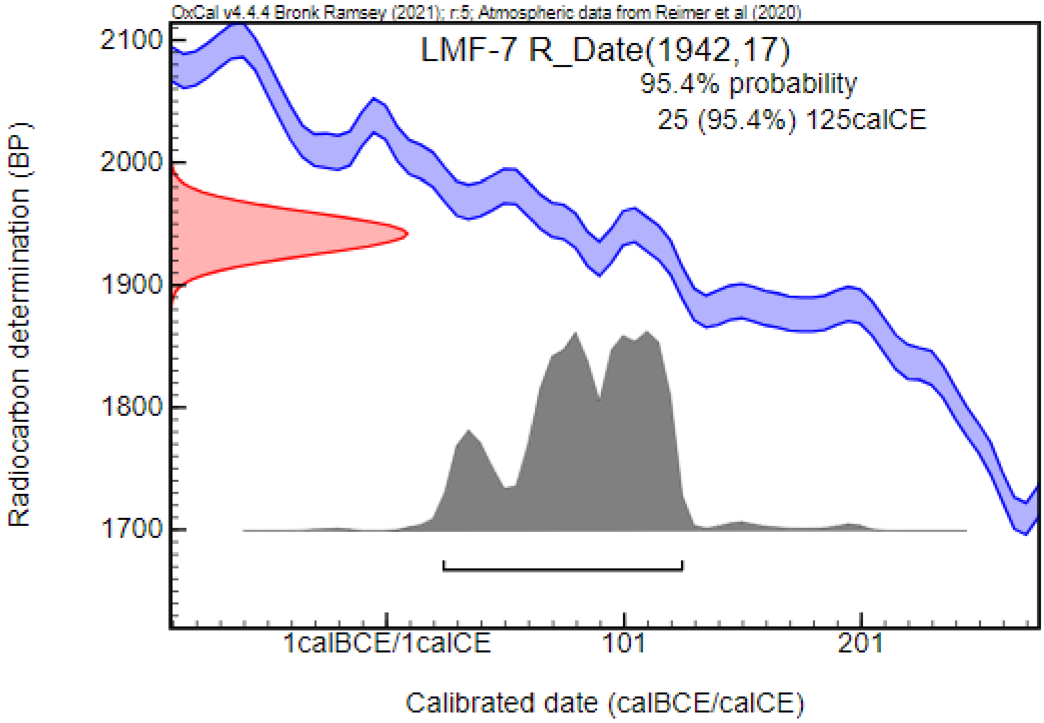


#### Madaras – Halmok (MDH; Bács-Kiskun County, Hungary) [91,92]

Following smaller excavations and fieldwork in 1903, 1952, and 1957, systematic rescue and preventive excavations led by Mihály Kőhegyi began at the border of Madaras village between 1963 and 1975. These excavations uncovered settlement features and burials from the Árpádian-age, as well as a Sarmatian burial ground containing 632 objects, including graves and pits containing animal bones. This cemetery is still considered the largest completely excavated Sarmatian cemetery in the Carpathian Basin that was published both archaeologically and anthropologically. More than half of the burials were partially or fully disturbed and/or robbed, complicating the archaeological and anthropological evaluation of the cemetery.

Three types of burial forms were identified in the cemetery: most were unmarked grave pits (481 cases), but graves with surrounding ditches (102 cases), and ditched barrow burials (14 cases) were also found. While these burial types did not form separate groups, four territorial groups were distinguished, containing a mix of these burial forms.

The size of the pits varied between burial types, with unmarked graves having smaller dimensions. Further differences have been found in age categories, with the graves of adolescences generally being smaller compared to those of adults. The orientation of the graves varied with SE¬–NW, SSE–NNW, and SSW–NNE directions being the most common. In addition, some graves were oriented in W–E and E–W directions, and in one case, an opposite, NE–SW orientation was observed. In a few cases, traces of fire were observed in the grave pits; however, contemporary robbery significantly limited the understanding of this phenomenon. In some cases, calcite layers, 0.3–3 cm thick, were discovered on the bottoms and sidewalls of the grave pits.

Most skeletons were found in a supine position with straightened extremities. Alternate positions, such as lying on their side or in a strongly flexed position, were observed only occasionally. The cemetery contained primarily single burials, with several exceptions where double burials of an adult and a sub-adult were documented. Iron parts of wooden coffins in various forms were found in good preservation in 50 burials, 10 of which were barrow burials. In addition, traces of small wooden chambers were detected in three barrow burials.

A high number of artifacts were recovered from the cemetery. The inventory of sub-adult burials was significantly poorer compared to adult burials, mostly containing unvaried beads and ceramic vessels, with jewelry (e.g., bracelets, earrings) being uncommon. Female graves were the richest in artifacts, featuring a high number of jewelry items (earrings, brooches, strings of beads, torques, pendants, bracelets) and various cloth fittings (e.g., bead ornaments on clothes). In addition, tools (e.g., awls, fire-lighting tools, needles, knives, and spindle whorls), mirrors, ceramic vessels, and, in one case, horse riding equipment were found in these burials. Male burials, on the other hand, contained fewer artifacts, with items traditionally associated with males, such as weapons, being particularly rare. Their inventory mostly included belt buckles, knives, ceramic vessels, with a smaller extent of jewelry (e.g., brooches, torques, pendants, bracelets), and, in one case, a sickle. Certain artifacts, including ceramic vessels, glass products, mirrors, coins, and brooches, indicate close contacts with the Roman Empire.

The Madaras cemetery was dated between the end of the 2nd century CE (i.e., the end of the Marcomannic wars) and the beginning of the 5th century CE. The abovementioned four main groups exhibited different characteristics. The first group, located in the southern part of the site and composed of both barrows and unmarked graves, was dated from the turn of the 2nd and 3rd centuries CE to the beginning of the 5th century CE. The second group, located in the southwestern part of the site, contained the earliest burials, with its use ending in the mid-3rd century CE. The third, central group consisted of a core of burials dating to the earliest period, spreading circularly mainly towards the south and southwest over time. The fourth group, located in the northern part of the cemetery, was dated between the mid-3rd and the beginning of the 5th centuries CE.

The results of the anthropological research were summarized by Antónia Marcsik. The state of preservation of the bones was generally poor, with a total of 463 skeletons available for analysis. Among them, 107 sub-adults and 356 adults were identified, including 205 females and 132 males. Based on morphometric analysis, Europid characteristics were observed in most of the skulls, primarily the Pamyrian type. However, a minor proportion of the individuals exhibited Mongolid and Europo-Mongolid features. Craniometric analysis revealed population heterogeneity, with only 70% of individuals sharing characteristics of this population, while 30% showed analogies with other 4th–6th-century CE series, such as Csongrád–Kenderföldek, from the southern region of the Great Hungarian Plain. In addition, traces of artificial deformation were observed on four skulls.

The Madaras–Halmok series illustrates the challenges researchers face when conducting bioarchaeological investigations on the Sarmatian period population of the Carpathian Basin. Out of 632 burial features, 463 contained skeletal remains available for anthropological analysis. However, only 46 individuals had skull elements (either petrous bone or tooth) preserved and suitable for archaeogenetic sampling. Ultimately, we could include 27 samples from the four groups of the cemetery in our analysis.

***MDH-28: Grave No. 28***

This was the robbed burial of a middle adult male, oriented in a SE–NW direction, which did not contain any known grave goods. The burial was in the 4th group of graves, dated to the 4th–5th centuries CE.

***MDH-48: Grave No. 48***

The grave with a mound contained the robbed burial of an adult male, oriented in a SE–NW direction. Found within were an iron knife – possibly from a modern period, a fragmented bronze needle, a biconic ceramic vessel, fragments of an iron buckle, fragments of an unknown iron artifact, iron parts of a wooden coffin, and animal bones belonging to horses, a cow, a sheep, and a rabbit. The burial was in the 4th group of graves, dated to the 4th–5th centuries CE.

***MDH-82: Grave No. 82***

The undisturbed grave contained the fragmented skeletal remains of a middle adult male, oriented in a SE–NW direction. Fragments of an iron brooch, a Roman coin emitted by Antonius Pius, an iron knife, and an iron buckle were discovered in the grave. The burial was in the 3rd group of graves, dated to the 3rd–4th centuries CE. Radiocarbon measurement confirmed the archaeological dating, placing the burial from the middle of the 3rd century CE to the first half of the 4th century CE.

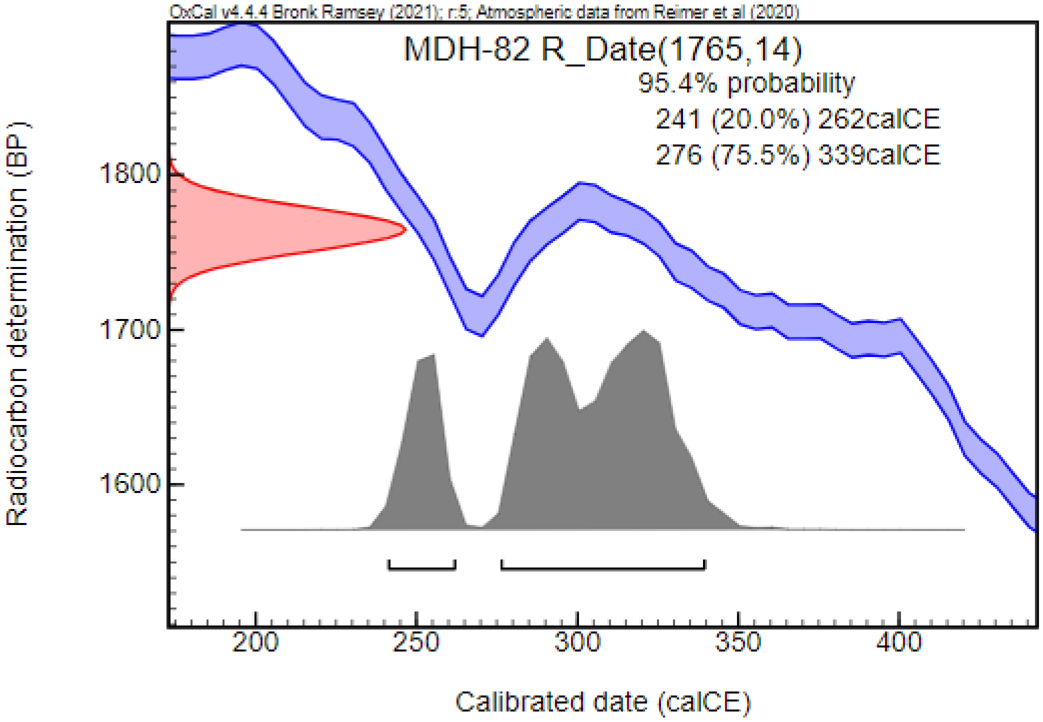


##### *MDH-84: Grave No. 84*

The fragmented skeleton of a sub-adult (12–14 years old) individual was found in the undisturbed grave, oriented in a S–N direction. The burial contained a pair of silver earrings, a string of glass beads around the neck, a two-piece bronze brooch, two bronze bracelets on the right forearm, an iron bracelet near the left forearm, approximately 395 glass beads on the lower legs and feet bones, and a hand made ceramic vessel. The burial was in the 3rd group of graves, dated to the 3rd–4th centuries CE.

##### *MDH-162: Grave No. 162*

The undisturbed burial contained the fragmented skeletal remains of an adult female, oriented in a SE–NW direction. Discovered within the grave were a string of beads near the neck region, an enameled brooch, a disc brooch, a spindle whorl, and a ceramic vessel. The burial was in the 3rd group of graves, dated to the 2nd–3rd centuries CE.

##### *MDH-178: Grave No. 178*

The fragmented skeleton of an old adult male was found in the undisturbed burial situated in the ditch around grave No. 174. Grave goods consisted of an iron knife and an iron belt buckle. The burial was in the 3rd group of graves, dated to the 2nd–3rd centuries CE.

##### *MDH-188: Grave No. 188*

This grave, surrounded by a mound and encircled by a ditch, contained the fragmented skeleton of an adult female. The burial had been robbed, and evidence of fire was found at the bottom of the pit. Among the artifacts recovered were an iron belt buckle, an iron dagger, an iron awl, a ceramic vessel, a bronze brooch with side-turned foot made of two pieces, and iron parts of a wooden coffin. The burial was in the 3rd group of graves, dated to the 4th–5th centuries CE.

***MDH-209: Grave No. 209***

The fragmented skeletal remains of a middle adult female with Mongolid characteristics and an artificially deformed skull were found in the grave, oriented in the opposite N–S direction. No grave goods were recovered from the narrow and short grave pit. Directly under this pit, another burial, grave No. 213 was discovered. The burial was in the 4th group of graves, dated to the 4th–5th centuries CE.

***MDH-221: Grave No. 221***

The robbed burial contained the fragmented skeleton of a middle adult female, oriented in a S–N direction. Discovered within the grave were an iron knife, a total of fifteen beads made of calcite, glass, or amber, a spindle whorl, shards of a ceramic bowl, and a ritual ceramic vessel. The burial, belonging to the 3rd group of graves, was dated to the 2nd–3rd centuries CE.

***MDH-249: Grave No. 249***

This undisturbed grave contained the skeletal remains of a middle adult male, oriented in a SE–NW direction. Grave goods included an iron belt buckle and a globular vessel. The burial belonged to the 3rd group of graves, dated to the 3rd–4th centuries CE. Radiocarbon measurement confirmed the archaeological dating and placed the burial from the second third of the 3rd century CE to the first third of the 4th century CE.

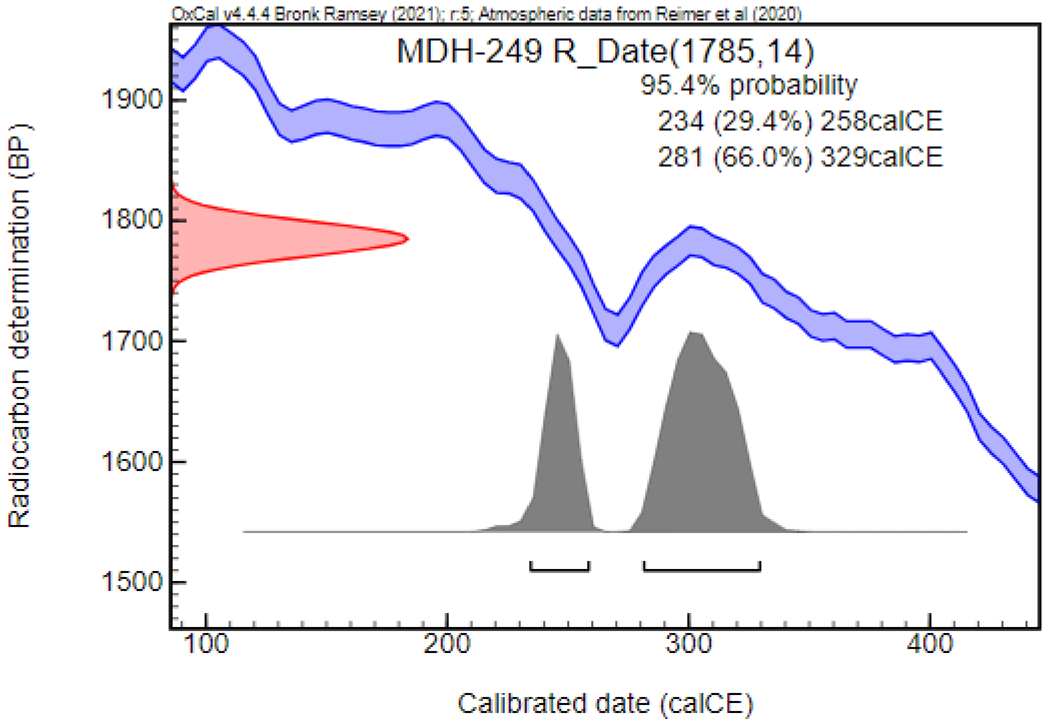


***MDH-263: Grave No. 263***

The undisturbed grave contained the fragmented skeletal remains of a young adult male, oriented in a S–N direction. Grave goods consisted of an iron knife, a hand made ceramic vessel, and fragments of an iron bracelet. The burial, belonging to the 3rd group of graves, was dated to the 2nd–3rd centuries CE.

***MDH-265: Grave No. 265***

This undisturbed burial contained the fragmented skeletal remains of a young adult male, positioned on their right side and oriented in an E–W direction. No grave goods were found. The burial belonged to the 3rd group of graves and was dated to the 2nd–3rd centuries CE. The archaeological dating was partially confirmed by the radiocarbon measurement, placing the burial from the middle of the 3rd century CE to the first half of the 4th century CE.

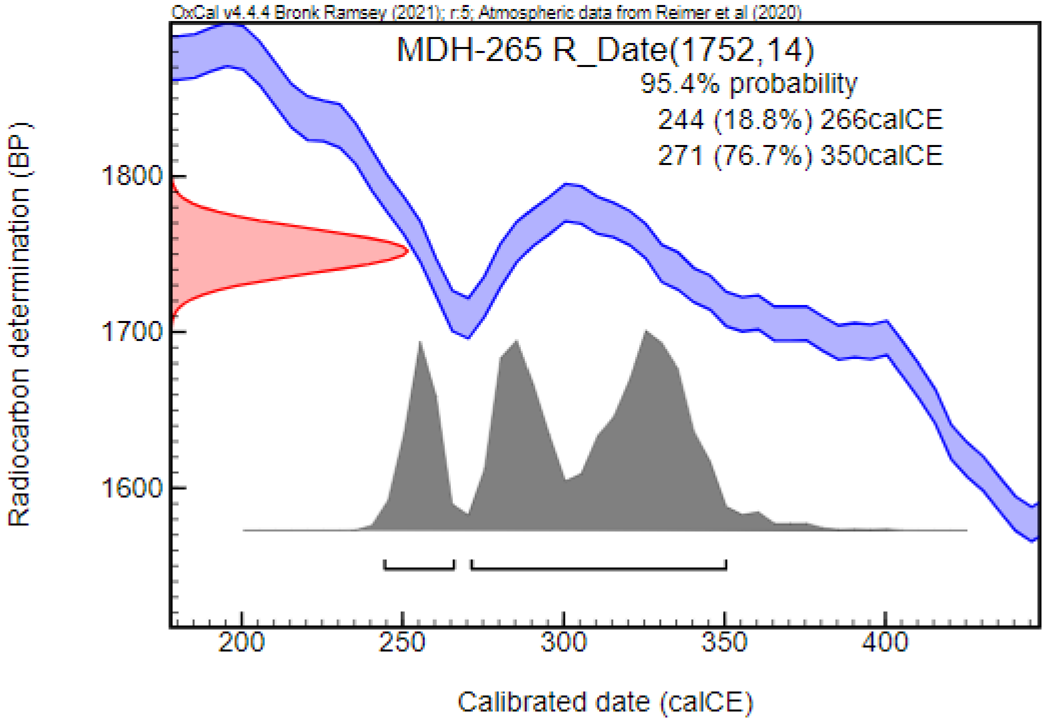


***MDH-291: Grave No. 291***

In this undisturbed grave, the skeleton of a middle adult female was found, oriented in a NE–SW direction. Among the recovered artifacts were a silver torque, a string of calcite and carnelian beads, a silver pendant (lunula), a pair of two-piece silver brooches with side-turned foot, an iron belt buckle, bead ornaments of the belt, and two U-shaped bronze plates. In addition, an iron knife, fragments of two iron bracelets, a string of beads on the left forearm, approximately 670 beads around the bones of the lower leg, and a ceramic vessel were found in the grave. The burial belonged to the 3rd group of graves and was dated to the 4th–5th centuries CE.

***MDH-342: Grave No. 342***

This undisturbed grave surrounded by a mound and encircled by a ditch, contained the skeleton of a young adult female, oriented in a SE–NW direction. Grave goods included a bronze ring near the head (earring?), a copy of a Faustina junior denar, a pair of bronze bracelets, and a fragment of ceramic vessel. The burial belonged to the 4th group of graves and was dated to the 4th–5th centuries CE.

***MDH-344: Grave No. 344***

The skeletal remains of a young adult female were found in the robbed burial, oriented in a S–N direction. The grave, surrounded by a mound and encircled by a ditch, contained only a ceramic vessel as grave goods. The burial belonged to the 4th group of graves and was dated to the 4th–5th centuries CE. Radiocarbon measurement, partially overlapping with the archaeological periodization, dated the burial from the beginning of the 3rd century CE to the first third of the 4th century CE.

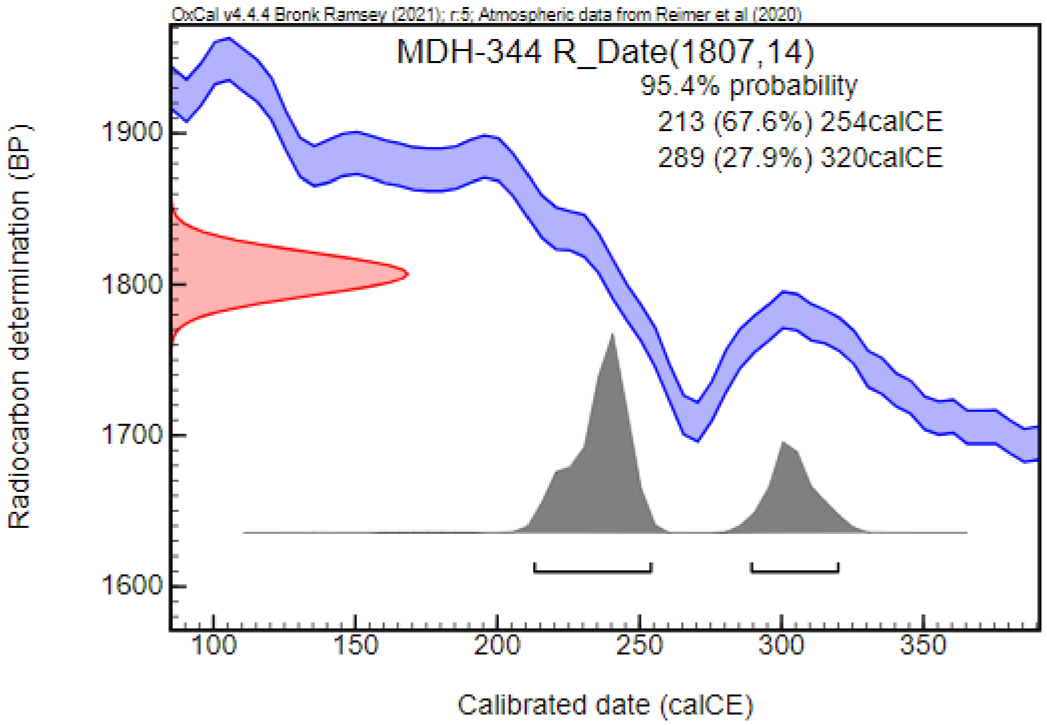


***MDH-357: Grave No. 357***

This robbed burial contained the fragmented skeleton of a young adult male, oriented in a S–N direction. Grave goods contained a ceramic vessel, a Roman coin emitted by Hadrianus, and the fragment of a two-piece iron brooch. The burial belonged to the 4th group of graves, dated to the 4th–5th centuries CE. Radiocarbon measurement, partially overlapping with the archaeological dating, placed the burial from the middle of the 3rd century CE to the middle of the 4th century CE.

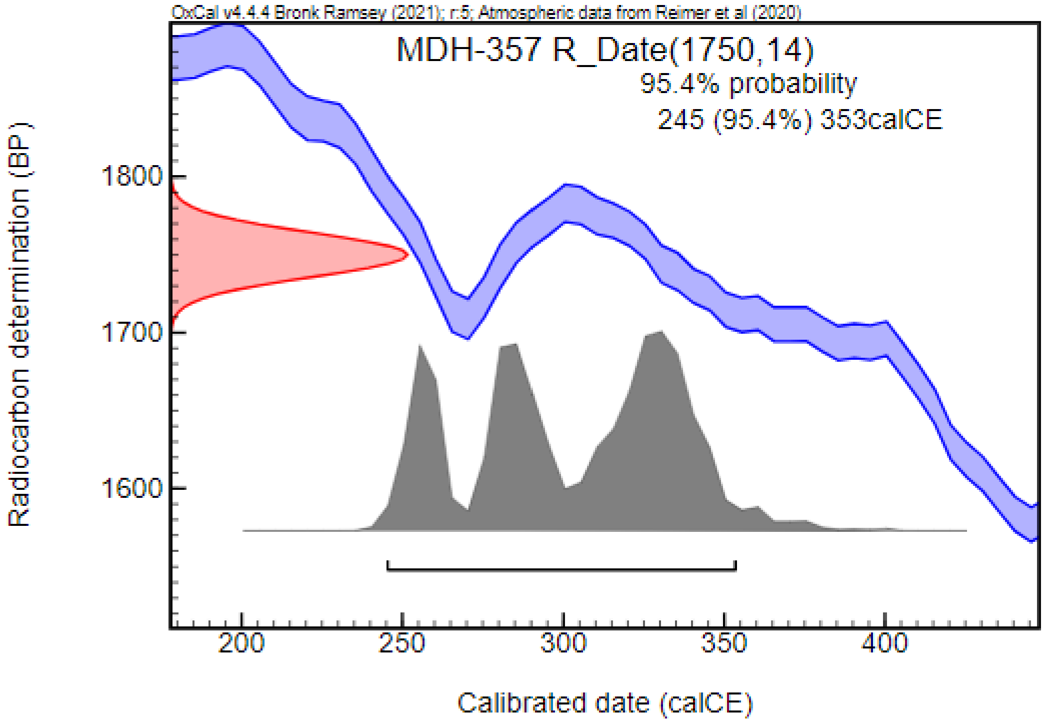


***MDH-405: Grave No. 405***

The skeletal remains of a young adult male were found in an undisturbed grave, oriented in a SE–NW direction. Grave goods included a glass bead under the skull, a bronze bracelet, a bronze torque, an iron bracelet, an iron belt buckle, and a shard of a ceramic vessel. The burial belonged to the 2nd group of graves, dated to the 4th–5th centuries CE.

***MDH-410: Grave No. 410***

In the undisturbed grave, the fragmented skeletal remains of a young adult female were found, oriented in a SE–NW direction. Grave goods included a pair of silver earrings, a silver torque, a string of glass and carnelian beads around the neck, and a one-piece silver brooch. In addition, two bronze bracelets on the left forearm, two bronze bracelets on the right forearm, approximately 754 beads around the lower leg bones, and two iron nails were uncovered. The burial belonged to the 3rd group of graves and was dated to the 3rd–4th centuries CE.

***MDH-444: Grave No. 444***

This robbed grave contained the fragmented skeleton of a middle adult male, oriented in a S–N direction. Artifacts recovered from the burial were an iron two-piece brooch, a piece of iron wire near the right part of the hip, an iron belt buckle, two coins emitted by the Constantinus dynasty, an iron knife, and a ceramic vessel. The burial belonged to the 3rd group of graves and was dated to the 4th–5th centuries CE.

***MDH-462: Grave No. 462***

This was the robbed burial of a young adult male, oriented in a SE–NW direction. Grave goods consisted of a crossbow brooch with onion-shaped knobs, a Roman coin emitted by Diocletianus, a belt buckle, and a ceramic vessel. The burial belonged to the 3rd group of graves and was dated to the 3rd–4th centuries CE

***MDH-483: Grave No. 483***

The fragmented skeleton of a middle adult female, oriented in a SW–NE direction, was found in this undisturbed burial. Grave goods included a pair of bronze earrings, an iron torque, a string of amber and glass beads, a pair of bronze bracelets, and approximately 465 beads around the lower leg bones. The burial belonged to the 3rd group of graves and was dated to the 3rd–4th centuries CE

***MDH-500: Grave No. 500***

This robbed grave, possibly once surrounded by a mound, contained the fragmented skeletal remains of a young adult female, oriented in a S–N direction. Artifacts discovered in the grave contained horse riding equipment including an iron bit, bronze plate of the harness and a girth buckle. In addition, a silver bracelet, fragments of a belt buckle or belt ornaments, fragments of a saw, and animal bones, belonging to at least two cows, and two sheep were uncovered. The burial belonged to the 3rd group of graves and was dated to the 4th–5th centuries CE. At the same time, it is important to note that similar types of horse harnesses can be dated much earlier. The bridles found as analogies are more commonly associated with the material culture of the late 2nd century CE and the first half of the 3rd century CE.

***MDH-514: Grave No. 514***

The fragmented skeleton of an old adult male was discovered in an undisturbed burial, oriented in a SE–NW direction. Grave goods included the fragments of an iron brooch, a dagger, and a ceramic jar. The burial belonged to the 3rd group of graves and was dated to the 3rd–4th centuries CE.

***MDH-592: Grave No. 592***

This robbed burial contained the fragmented skeleton of a middle adult male, oriented in a S–N direction. Artifacts found within the grave included a two-piece iron brooch, an iron knife, a ceramic vessel, and an iron awl. The burial belonged to the 2nd group of graves and was dated to the 3rd–4th centuries CE. Radiocarbon measurement, partially overlapping with the archeological dating, placed the burial from the second third of the 2nd century CE to the second third of the 3rd century CE.

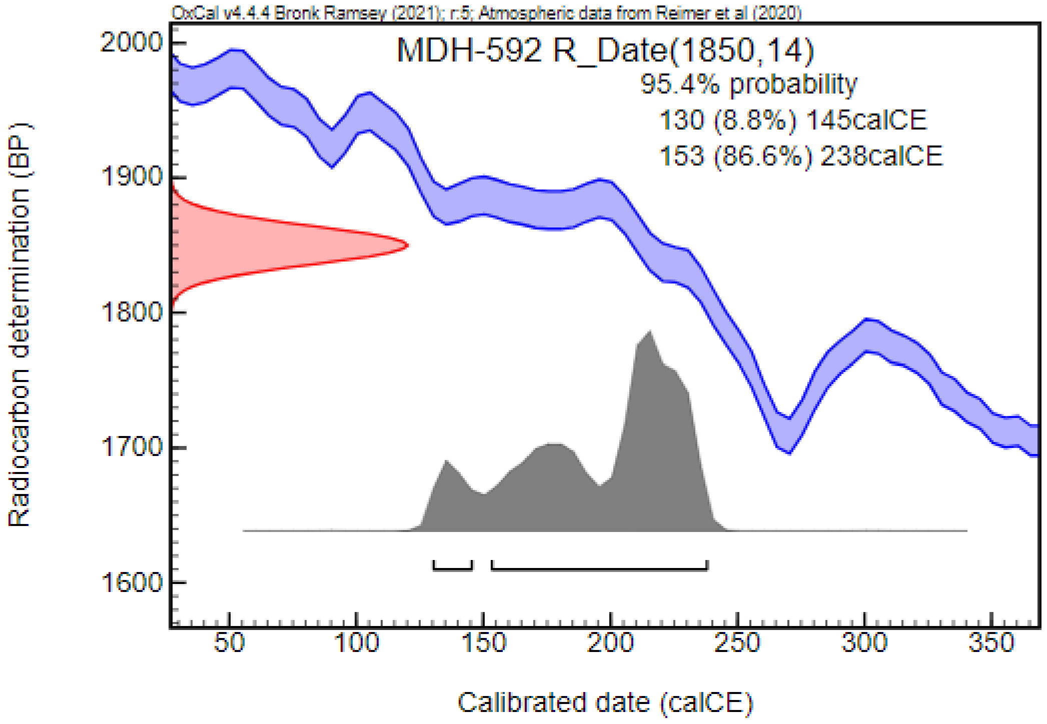


***MDH-630: Grave No. 630***

The fragmented skeleton of a middle adult female, oriented in a SE–NW direction, was found in this undisturbed grave. Fragments of an iron brooch were uncovered from the burial. The burial belonged to the 2nd group of graves and was dated to the 2nd–3rd centuries CE.

***MDH-656: Grave No. 656***

This was the burial of a young adult female, oriented in a SE–NW direction. Assumingly, based on the size of the pit and the composition of the material recovered, the grave was originally constructed with a mound. Artifacts found included four S-shaped coffin clamps, three leaf-shaped pressed golden ornaments, a spindle whorl, approximately 54 beads from the disturbed part of the grave, and a pendant. In addition, bronze plates and nails possibly belonging to a wooden chest, a shard of a terra sigillata, a fragment of an iron plate, and fragments of bronze plates were recovered from the grave pit. The burial belonged to the 1st group of graves and was dated to the 2nd–3rd centuries CE.

***MDH-661: Grave No. 661***

The fragmented skeleton of a middle adult male, oriented in a SE–NW direction, was discovered in the robbed burial. Assumingly, the grave was originally constructed with a mound, shared with grave No. 559. Grave goods included five S-shaped coffin clamps, a Roman coin emitted by Commodus, and a belt buckle. The burial belonged to the 1st group of graves and was dated to the 2nd–3rd centuries CE

#### Makó – Igási Járandó (MIJ; Csongrád-Csanád County, Hungary) [93]

In 2008, preventive excavations led by Csilla Balogh were conducted in connection with the construction of the M43 highway. A multiple-period site with Bronze Age, Iron Age, and Medieval settlement features, as well as a Medieval and a Sarmatian burial grounds, was discovered. The fully excavated Sarmatian cemetery consisted of 16 graves. Most of the graves were oriented in a S–N direction with minor deviations toward SW–NE or SSW–NNE, while in two cases, WNW–ESE and NW–SE orientations were registered. The grave pits were simple and mostly rectangular, with no trenches or barrows observed. Traces of wooden coffins were detected in five graves, while in one case, traces of burned organic remains were documented between the knees of the deceased. Most of the burials were affected by contemporary disturbance and/or robbery, limiting the archaeological and anthropological evaluation of the cemetery. The cemetery consisted only of single burials, with the skeletons found in a supine position. Animal bones were found only in one grave, where horse metatarsal bone fragments were documented. Artifacts were found in 11 graves. The grave goods included jewelry (e.g., earrings, strings of beads, torques, pendants, bracelets, and finger rings), cloth fittings (e.g., brooches, belt elements, and bead ornaments), tools (e.g., knives, spindle whorls, and awls), and ceramic vessels. Some of the artifacts found in the burials, such as brooches and beads, indicate the community's contacts with the Roman Empire and the eastern part of the Black Sea region. The cemetery was dated from the mid-2nd century CE to the mid-3rd century CE.

The state of preservation of the bones was generally poor, limiting the analysis of the skeletons. The anthropological investigation conducted by Antónia Marcsik revealed a disparity between sexes: out of the 16 burials, 13 individuals were determined to be female, and only one skeleton showed male characteristics. There were 12, and only 3 sub-adults were identified in the series. We could include 4 samples in our analysis.

***MIJ-1: Grave No. 1 (SNR 261)***

Traces of organic material, possibly belonging to a wooden coffin, were discovered during the excavation of a medieval house pit. Although the grave pit was undetectable, the skeletal remains of a middle adult female were found in a supine position, oriented in an S–N direction. A total of 15 beads scattered across the upper body, a spindle whorl, a ceramic vessel, and an iron knife were uncovered in the burial. Radiocarbon measurement, confirming the archeological dating, placed the burial from the first third of the 2nd century CE to the first third of the 3rd century CE.

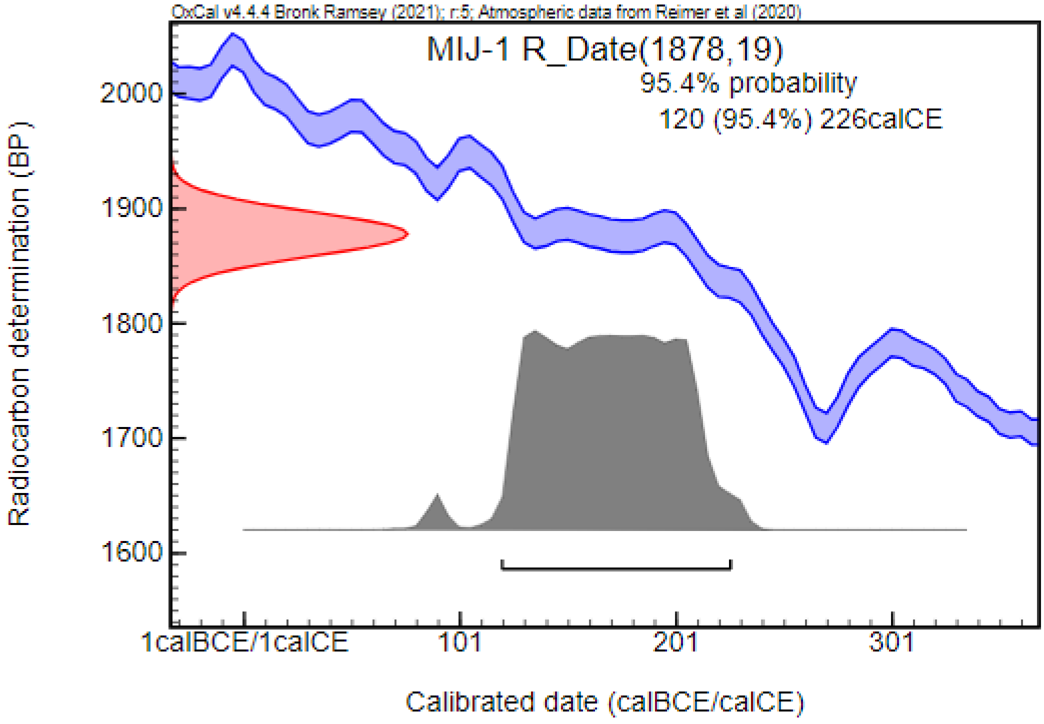


***MIJ-3: Grave No. 3 (SNR 266)***

This burial contained the skeletal remains of a middle adult female, oriented in a SW–NE direction. Traces of organic material were detected under the skeleton. Grave goods included a two-piece bronze brooch, a piece of stone, and horse metatarsal bones. Radiocarbon measurement, partially overlapping with the archeological dating, placed the burial from the beginning of the 1st century CE to the first third of the 2nd century CE, with the possibility of dating to the last third of the 1st century BCE (3.7%).

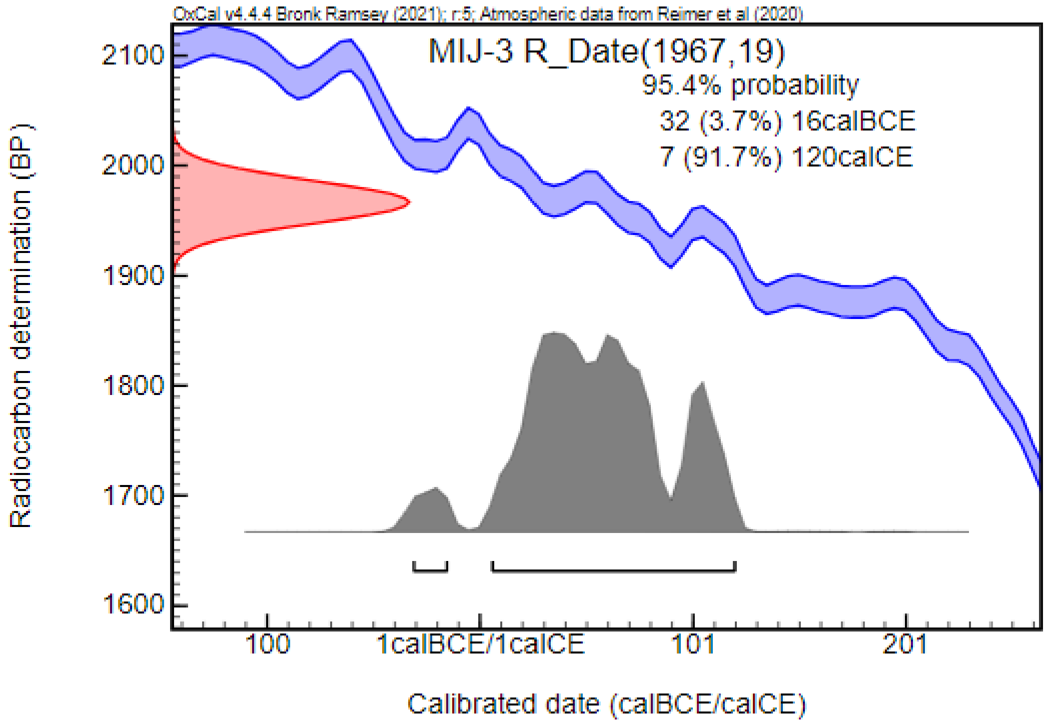


***MIJ-4: Grave No. 4 (SNR 291)***

The skeletal remains of a young adult female, oriented in an SSW–NNE direction, were found. Artifacts in the grave included a bronze earring, bronze plate fragments, a bronze pendant (lunula), a silver torque, four circular bronze pendants, two bronze finger rings, a total of 44 beads across the upper body, a bronze brooch inlaid with enamel, and a bronze two-piece 'T' brooch. In addition, a bronze ring under the left forearm, groups of beads near the hip, a spindle whorl, two glass balls, a cowry shell, a bronze spring-like ring, strings of beads near the wrists, a total of 391 beads near the lower leg and feet bones, fragments of an iron knife, a bone needle case, a bronze needle, and a ceramic mug were found. The re-calibration of the radiocarbon results from the analysis conducted earlier[93], confirmed the archaeological dating, placing the burial from the 2nd century CE to the second third of the 3rd century CE, with a possibility of dating to the end of the 1st century CE (5.1%).

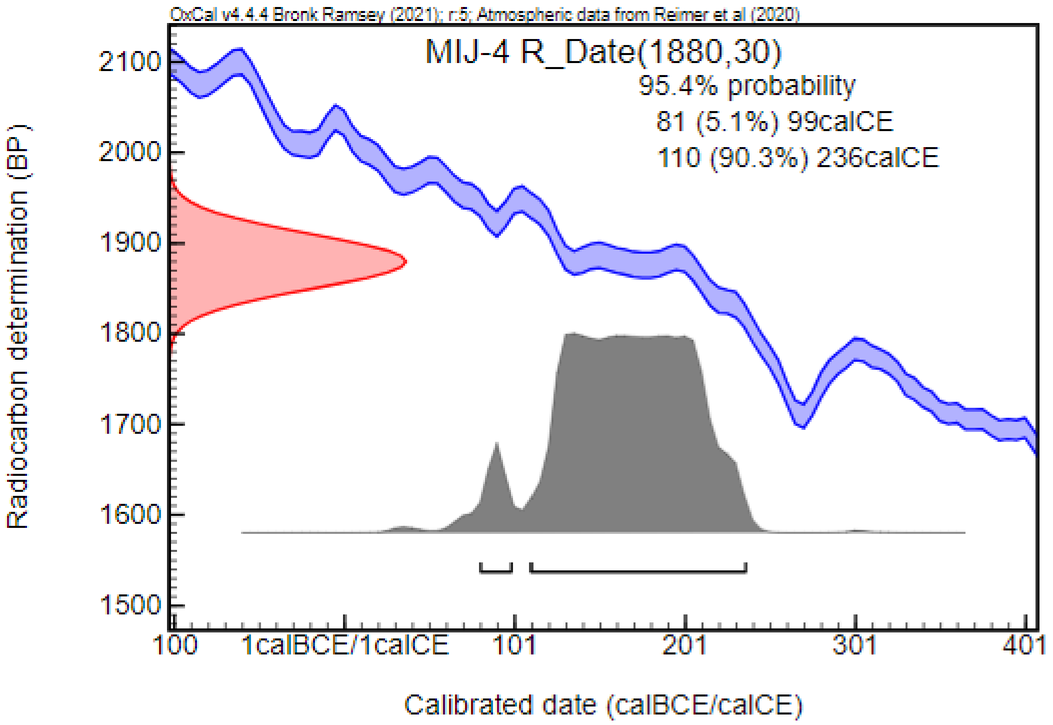


***MIJ-7: Grave No. 7 (SNR 340)***

A rectangular grave pit with traces of a wooden coffin was discovered. This was the burial of a middle adult female, buried in a supine position and oriented in an SSW–NNE direction. Grave goods included fragments of a coiled bronze earring, a total of 17 beads near the chest and shoulders, and three calcite beads near the right elbow, two silver pendants with boxes, a total of 21 beads near the right wrist, a bronze ring near the hip, a total of 130 beads above the lower leg bones, 15 beads near the left and right femur. Additionally, a glass spindle whorl, a cross-shaped two-piece bronze brooch, a silver strongly profiled brooch, a bronze plate fragment, an iron knife, and a ceramic mug were found. The re-calibration of the radiocarbon results from the analysis conducted earlier [93], confirmed the archaeological dating, placing the burial from the last third of the 1st century CE to the second third of the 3rd century CE.

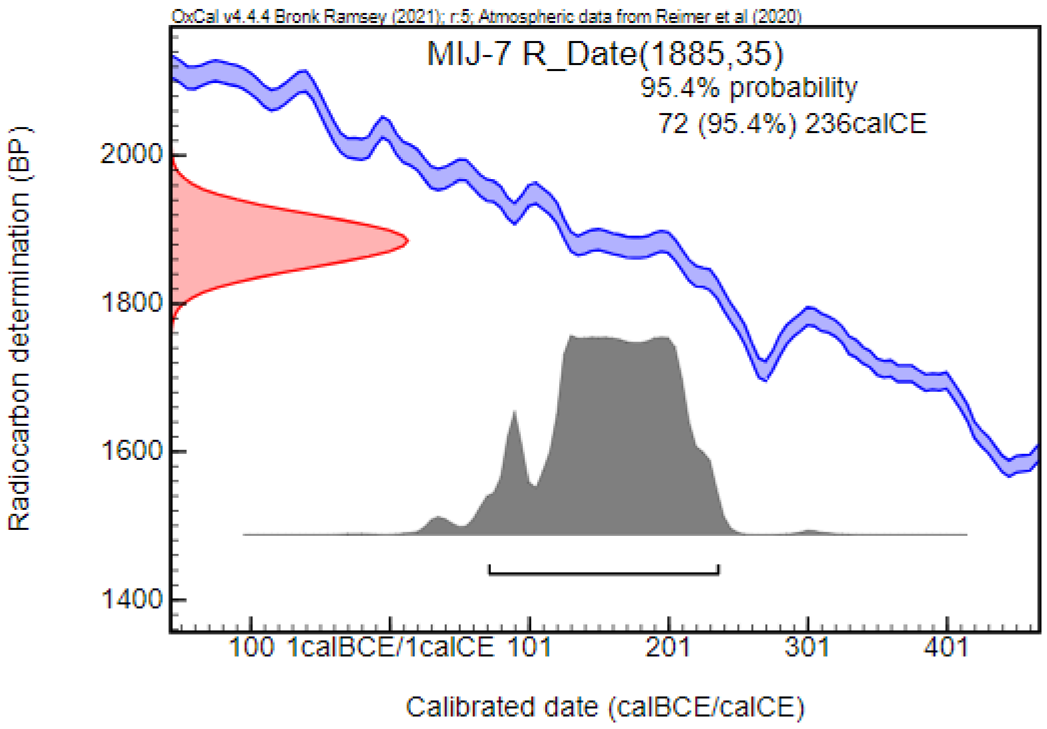


#### Makó – Mikócsa dűlő M43, 31. lh. (MM2; Csongrád-Csanád County, Hungary) [94]

In 2008, preventive excavations led by Dániel Pópity were carried out in connection with the construction of the M43 highway in the close vicinity of Makó. Settlement objects dated to the Sarmatian and Avar periods, a burial from an unknown period, and two burials dated to the Sarmatian period were discovered. The two Sarmatian-period burials were oriented in a SW–NE direction, and contained various grave goods, including jewelry, cloth fittings, tools and ceramic vessels. Some of the artifacts, such as the mirror, chest ornaments, and amber beads, suggest connections with the Roman Empire. The burials were dated to the late Sarmatian period, between the end of the 4th and beginning of the 5th centuries CE. The human bones were analyzed by Antónia Marcsik. Both graves contained the skeletal remains of adult female individuals. We included both Sarmatian-period graves in our analysis.

***MM2-215: Grave No. 215 (SNR 289)***

The grave, oriented in a SW–NE direction, contained the skeleton of an adult female in a supine position. Traces of a wooden coffin were discovered in the grave. Grave goods included a bone needle case with an iron needle, a pair of bronze earrings, a two-piece bronze brooch, a pair of bronze bracelets, a total of 85 beads near the feet, a ceramic jar, a ceramic vessel, a spindle whorl, and an iron knife. A further 31 beads of various types were found throughout the grave.

***MM2-245: Grave No. 245 (SNR 290)***

This burial contained the skeletal remains of a young adult female in a supine position, oriented in an SW–NE direction. A bronze earring, a string of beads with a bronze pendant (lunula), four small semi-spherical bronze ornaments, a pair of tear-shaped bronze pendants, two small bronze tubes, a two-piece silver brooch, and an iron hook were found in the grave. Further grave goods, including bronze chest ornaments, a bronze mirror, two silver bracelets, and a total of 449 beads near the lower leg bones, were also discovered. Radiocarbon measurement supported an earlier dating, placing the burial from the beginning of the 3rd century CE to the first third of the 4th century CE.

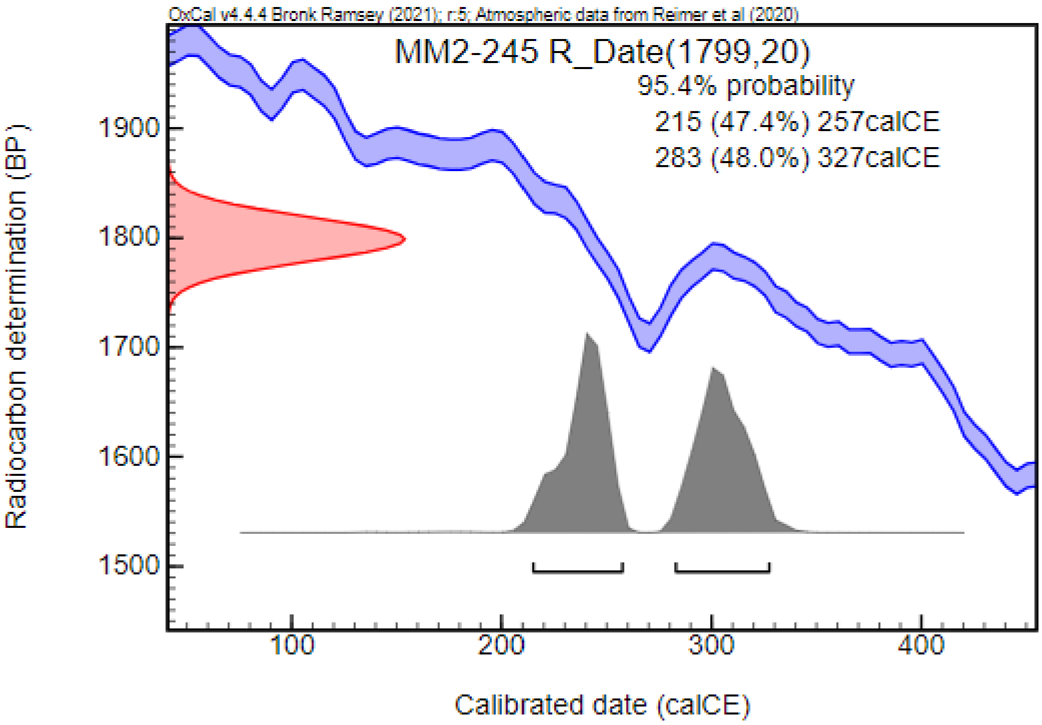


#### Mezőcsát–Hörcsögös (MEH; Borsod-Abaúj-Zemplén County, Hungary) [95]

During the excavations led by Erzsébet Patak and Nándor Kalicz at a multi-period site, six Sarmatian-period graves were uncovered in two burial groups. The first group was richer in artifacts. The graves, oriented mostly in an S–N direction, were unmarked and contained skeletal remains in a supine position. The archaeological and initial anthropological analyses indicated a demographic shift, identifying four females and a sub-adult, accompanied by artifacts traditionally associated with females. The grave goods included ceramic vessels, tools (e.g., spindle whorl, knife), cloth fittings (e.g., belt elements, brooch), and jewelry (e.g., pendant, beads, bracelets), including golden artifacts (e.g., beads, pendant). The cemetery was associated with the early Sarmatians and dated to the 1st and 2nd centuries CE. Due to the poor preservation of the skeletal remains, only one sample was included in our analysis.

##### *MEH-2p: Pit Grave No. 2*

As part of the second burial group, this settlement grave No. 2, oriented in an S–N direction, contained the fragmented skeleton of an adult female. The grave goods included a ceramic cup, an iron bracelet, a bronze bracelet, and a total of 120 beads around the lower leg bones. Radiocarbon measurement, partially overlapping with the archaeological dating, placed the burial from the second third of the 2nd century CE to the beginning of the 4th century CE.

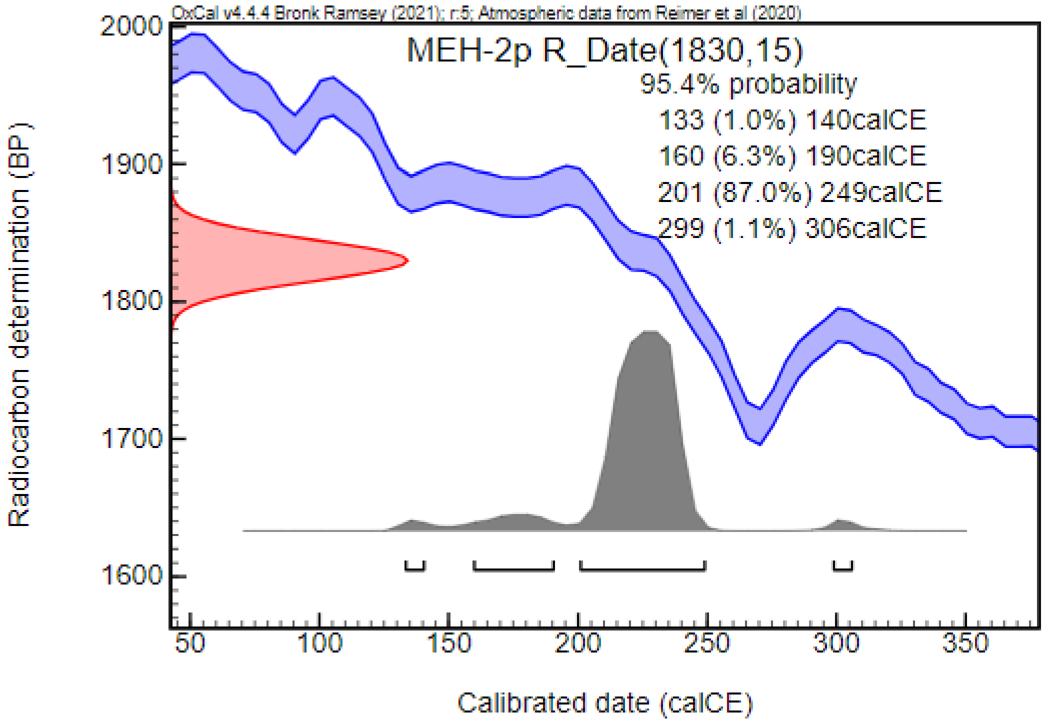


#### Mezőkövesd – Mocsolyás (MEM; Borsod-Abaúj-Zemplén County, Hungary) [96]

During the excavation of a Neolith settlement at the M3 Highway section near Mezőkövesd, three burials dating to the 5th century CE were uncovered. The burials, oriented in a NE–SW direction, contained the skeletal remains of an adult female with signs of artificial cranial deformation and two sub-adult individuals. The graves were richly furnished with jewelry (e.g., hair ring, earrings, bracelets, and finger rings) and cloth fittings (brooches, belt buckles), with artifacts made of silver and gilded bronze, as well as an iron knife, a mirror, and ceramic vessels. Although the orientation of the graves is reminiscent of Sarmatian-period burial customs, the style of the jewelry and cloth fittings, notably the large silver brooch found in Grave No. 2, along with the cranial deformation, indicate cultural influences from the Hun period and the subsequent decades We could include one sample in our analysis.

***MEM-2: Grave No. 2 (object No. 367)***

The large grave, oriented in a NE–SW direction, contained the skeleton of an adult female with an artificially deformed skull, placed in a supine position. The grave goods included a pair of gilded bronze earrings with polyhedron pendants, two large silver plate brooches (approximately 19.7 cm in length), glass and amber beads near the rib area, an iron knife, a white metal mirror, two silver bracelets, a bronze plate finger ring, a silver plate finger ring, a silver belt buckle, another silver buckle, and a Murga-type ceramic jug. Radiocarbon measurement, confirming the archaeological dating, placed the burial in the 5th and early 6th centuries CE.

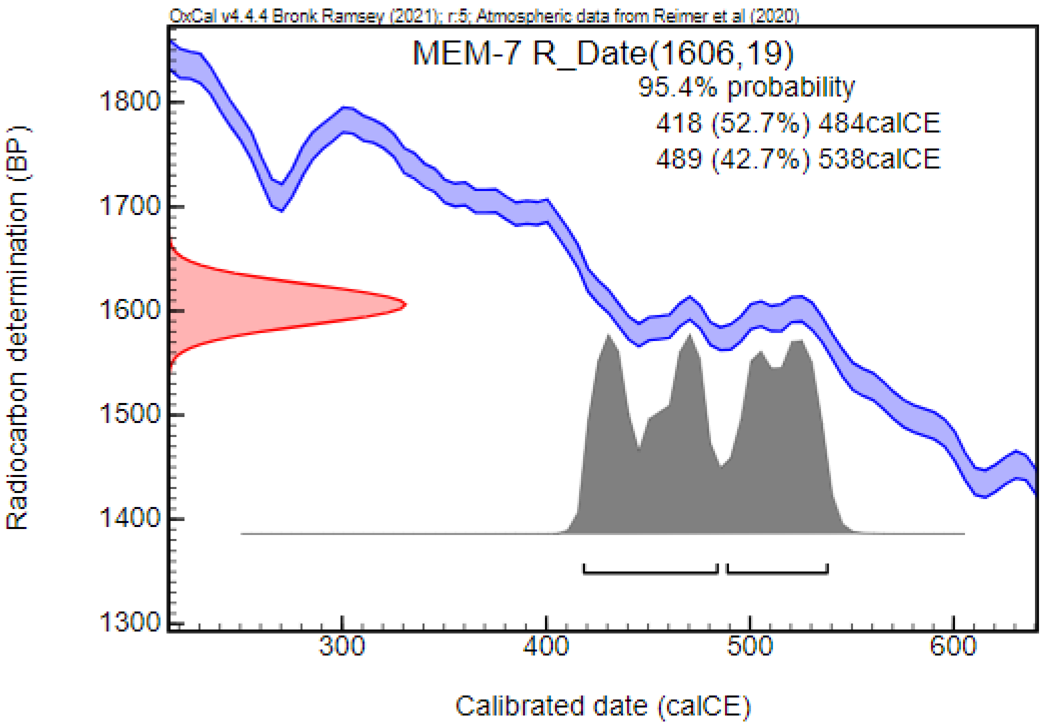


#### Nagykálló – Kis Ludas-tó dűlő (NKL; Szabolcs-Szatmár-Bereg County, Hungary) [97]

Preventive excavations, led by Gábor Pintye, were conducted in 2008 and 2009 in the area of an industrial park near Kisludas-tó dűlő. These excavations uncovered settlement parts from the Scythian, Sarmatian, and Avar periods, as well as two Scythian-period graves, two Hun-period burials, and a Sarmatian burial ground with 57 graves. The cemetery is considered partially excavated and the archaeological and anthropological analysis are still in progress. The graves were severly disturbed, with three exceptions. The deceased were commonly interred in a supine upright position, with the exception of one skeleton lying on its sides. The orientation of the graves predominantly followed a south-north direction, with slight variations toward the east and west. In one instance, there is evidence of a diagonal positioning of a coffin. Of the 67 burials, 20 were graves surrounded by flat circular ditches. The use of coffins was noted in several cases. The Sarmatian artifacts date from the early to mid-3rd century CE to the mid-5th century CE. We could include 4 samples in our analysis.

##### *NKL-7: Object No. 7*

A rectangular bermed burial pit with rounded corners has been constructed. The deceased was oriented in a south-north direction and was placed in a coffin. The burial also featured a circular ditch. This burial contained the skeletal remains of an adult female. The grave itself has been disturbed or looted, with the robber's pit excavated on the northwest side. All that remains of the skeleton is the skull, which faces east, with the jaw oriented to the west. Height: 342 cm; Width: 160 cm; Depth: 104 cm. Grave Goods included six flat carnelian beads shaped like cuboctahedra (one with iron residue), seven pressed spherical blue glass beads with encaustic decoration, a biconical gray clay spindle whorl, a bronze plate bead (?), folded back at both ends, and a biconical pressed silver bead (?). The burial can be dated to the 3rd–4th centuries CE. Radiocarbon dating places the burial in the Sarmatian period, from the end of the 1st century CE to the early-3rd century CE.

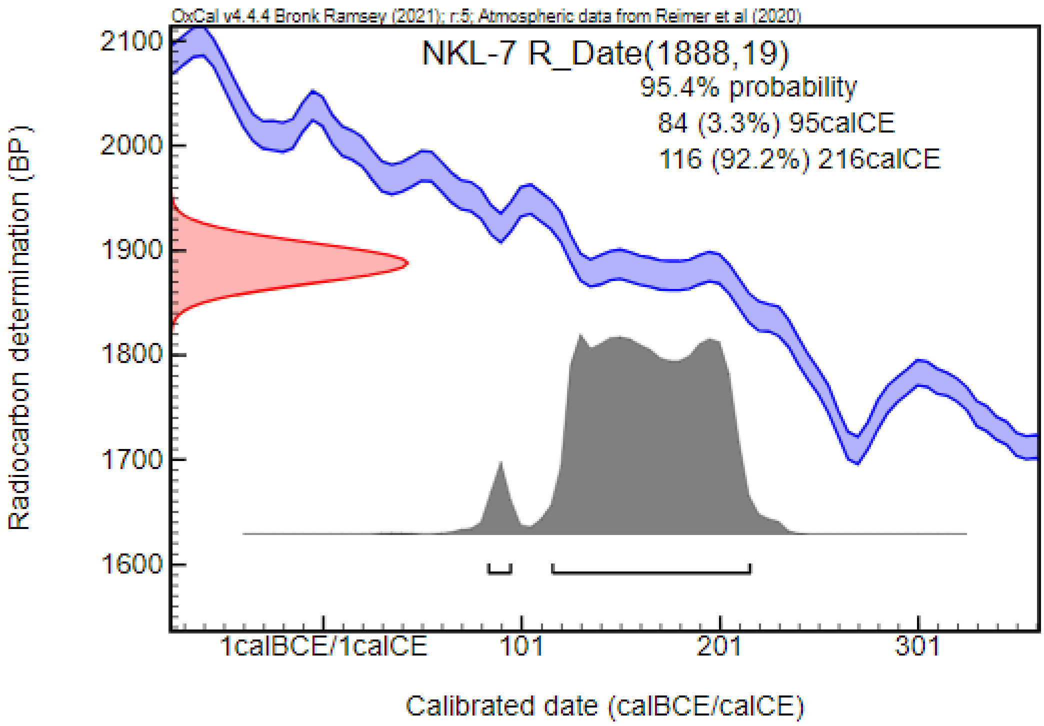


***NKL-112: Object No. 112***

A rounded-heeled grave pit with a southeast-northeast (SE-NE) orientation was found. The skeletal remains of an adult female were recovered from this burial. The burial was robbed, with the bulk of the bones concentrated in the middle third of the grave. Height: 240 cm; Width: 90 cm; Depth: 40 cm. The grave goods included one rectangular green glass bead, one green glass bead shaped like a pentagon, and one fragment of a grey wheel--made ceramic vessel. The finds did not allow for a more precise dating within the period of the 2nd to 5th centuries CE.

***NKL-135: Object No. 135***

A rectangular grave pit was discovered, oriented southeast-northeast (SE-NE). This burial contained the skeletal remains of an adult male. The grave had been robbed, and the skeleton was found in a supine position, originally stretched, but the middle third had been heavily disturbed. Height: 264 cm; Width: 113 cm; Depth: 57 cm. Grave goods included only a D-shaped iron buckle. The burial can be dated to the 4th century CE based on the archaeological finds. Radiocarbon dating places the burial in the Sarmatian period, between the first third of the 1st century CE and the beginning of the 3rd century CE.

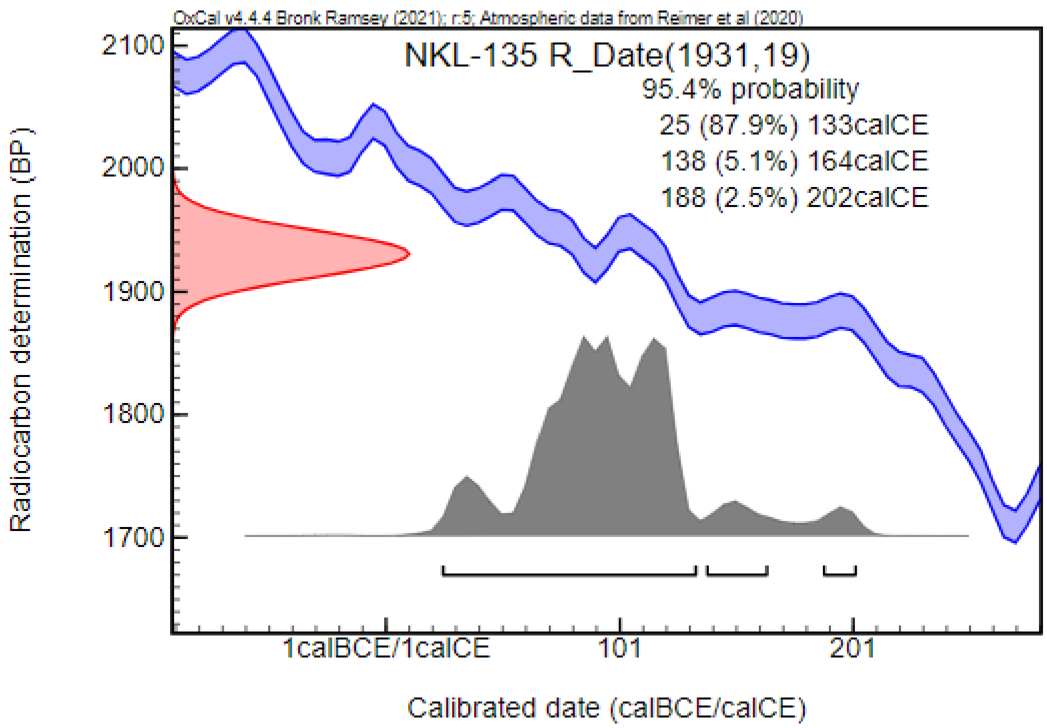


***NKL-157: Object No. 157***

A rectangular tomb, oriented south-north, has been robbed or disturbed. The skeleton of the deceased lies on its back in a stretched position, with the pelvis on the right side. The skeleton of an adult female was found in this grave. Height: 244 cm; Width: 108 cm; Depth: 20 cm. Grave goods included fourteen cuboctahedron-shaped carnelian beads, one cuboctahedron-shaped amber bead, one cylindrical amber bead, a bronze brooch with a side-folded foot, a grey wheel-thrown bowl, and two bronze earrings with closing system made of a hook and a loot-bent extremity. The burial can be dated to the 4th and early 5th centuries CE based on the archaeological finds. Radiocarbon dating places the burial in the Sarmatian period, between the last third of the 2nd century CE and the first third of the 4th century CE.

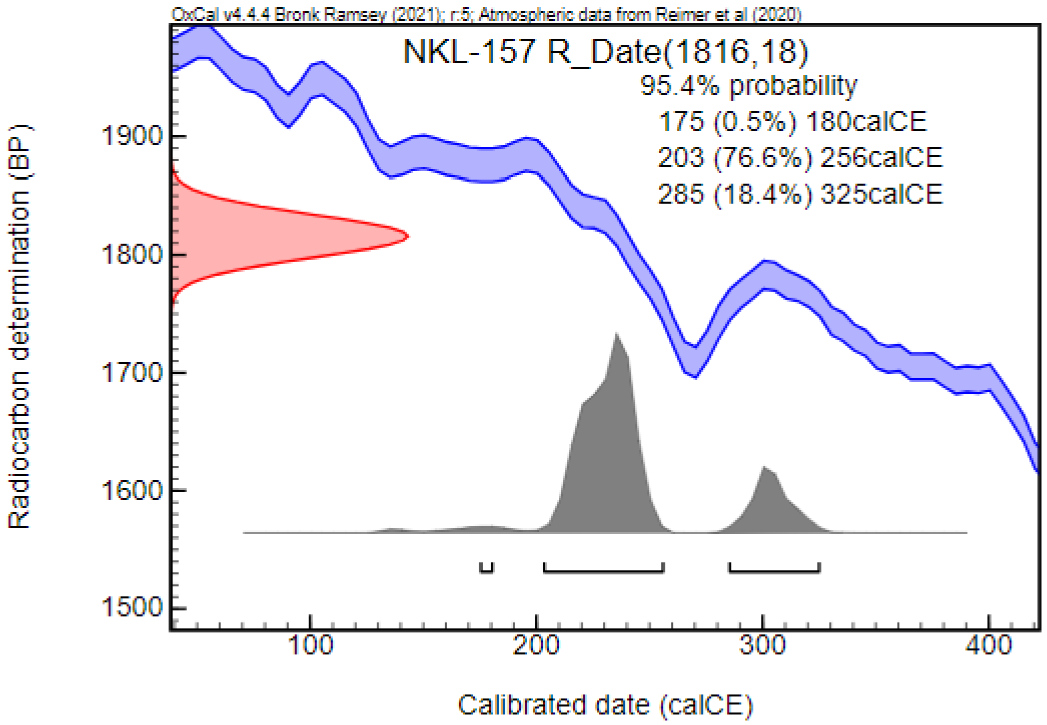


#### Nagykálló – Kis Ludas-tó dűlő (NKL; Szabolcs-Szatmár-Bereg County, Hungary)[97]

During the same excavation described above, apart from the Sarmatian-period cemetery, two graves dating to the Hun period were discovered. The orientation of these graves differed from the Sarmatians burials and from each other, suggesting cultural differences. The graves contained the skeletal remains of a female and male, with the female grave being richer in artifacts, such as jewelry, compared to the male counterpart. One of the graves was superimposed with a settlement object, indicating that the Sarmatian settlement was still in use when these burials were made. The artifacts found in the graves have parallels in the 4th and 5th century CE cemeteries of the Carpathian Basin, such as those at Csongrád–Kenderföldek and Óföldeák–Ürmös II. The two burials were dated to the end of the 4th and 5th centuries CE. Connections with the Sarmatian-period population as well as Eastern Germanic relations were suggested, but the ethnicity of the two individuals remains a subject of debate.

***NKL-899: Object No. 899***

This grave, oriented in a W–E direction, contained the skeleton of an adult female in a supine position. Grave goods included a pair of one-piece iron brooch, an iron knife, two earrings with polyhedron pendants, and a string of glass beads near the neck. Radiocarbon dating, suggesting an earlier date, placed the burial from the second half of the 3rd century CE to the early 5th century CE.

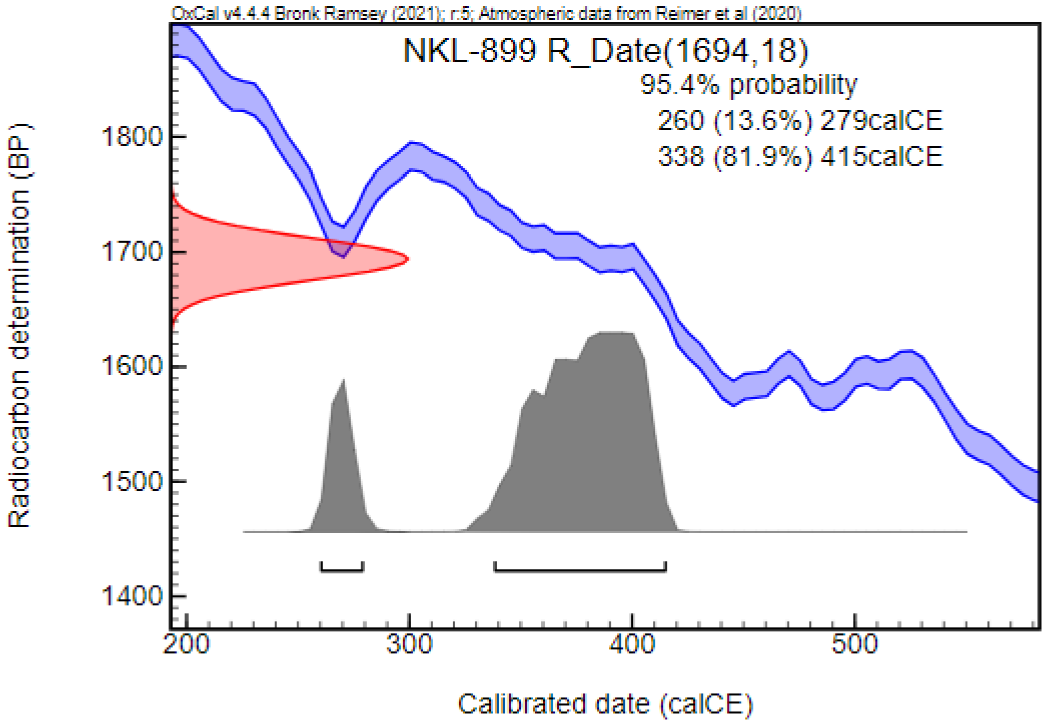


##### *NKL-1038: Object No. 1038*

The grave pit, oriented in a N–S direction, was superimposed with a settlement object. The skeletal remains of an adult male were found in a supine position, accompanied with a Murga-type ceramic jug and a kidney-shaped iron belt buckle.

#### Nemesnádudvar – Külbogyiszló, Túzok-hát (NKT; Bács-Kiskun County, Hungary)

In the spring of 2021, a rescue excavation led by Réka Cs. Andrási near Nemesnádudvar, on a field affected by agricultural works, resulted in the discovery of three burials from the 10th century CE. The excavations continued in the autumn, uncovering an additional nine 10th-century-CE burials, a Bronze Age burial, ten Sarmatian graves –including a ditched burial– and a grave from an unknown period. The site is considered only partially excavated, and further graves are expected in the unexplored areas. The site is unpublished, and the archeological and anthropological investigation of the recovered material is still in progress. Most of the Sarmatian graves were robbed, leaving few artifacts or human bones for analysis. Grave goods were recovered from only one burial, while other graves contained ceramic shards, stray beads, and spindle whorl fragments found in the grave soil. The preservation of the bones was poor, with only two graves containing more than a few small bone fragments. Anthropological analysis, conducted by Balázs Tihanyi and Orsolya Anna Váradi, identified the skeletal remains of two adult individuals. These two cases were included in our analysis.

##### *NKT-20: Grave No. 20*

Located in the southeastern corner of the excavated area, this grave, oriented in an S–N direction, contained the fragmented skeletal remains of an adult, possibly female individual, in a supine position with straightened extremities. The grave goods included a ceramic vessel, a string of glass and carnelian beads near the neck, and beads recovered from the lower leg area. Radiocarban dating, confirming the archaeological dating, positioned the burial to the Sarmatian period, between the last third of the 1st century CE and the beginning of the 3rd century CE.

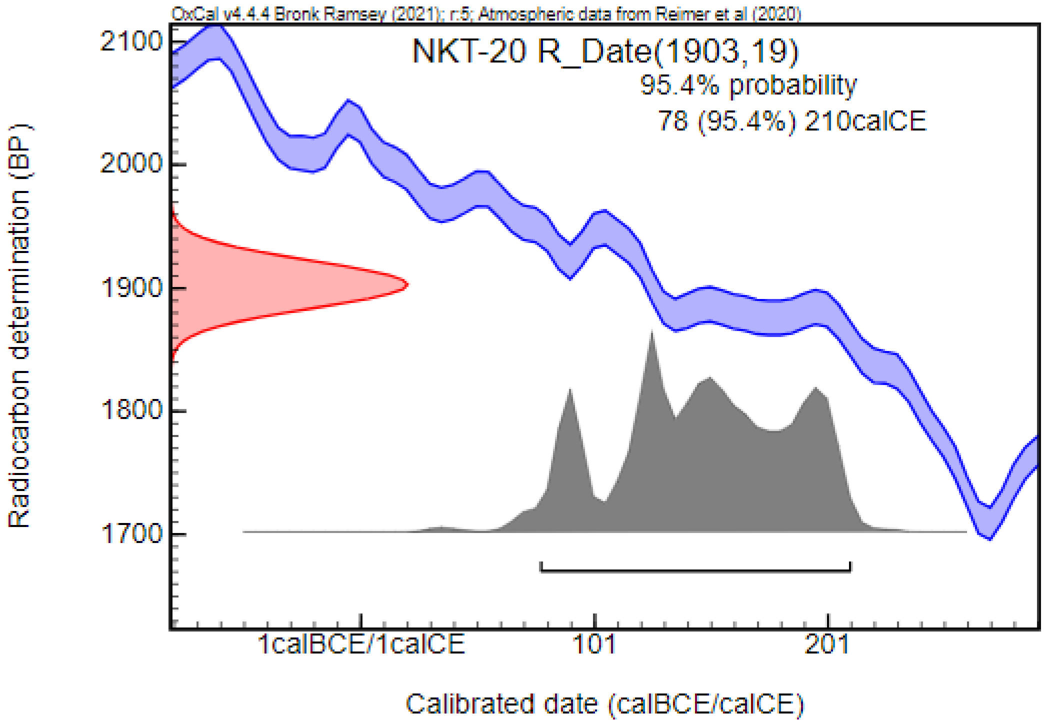


***NKT-32: Grave No. 32***

This robbed burial, oriented along an S–N axis, contained only the fragmented remains of an adult individual, with ceramic shards in the soil of the grave.

#### Óföldeák – Ürmös M43 9–10. lh. (OFU; Csongrád-Csanád County, Hungary) [98]

In 2008 and 2009, preventive excavations were conducted near Óföldeák in conjunction with the construction of the M43 highway. Sites numbered 9, 10, and 11, though separated by a road and a canal, appear to constitute a single archaeological site. Six grave groups were discovered, with group 5, located in the southwestern part of site No. 9, comprising three graves oriented in a N–S direction. All three graves featured a sidewall niche, and one was superimposed with a Sarmatian-period settlement object. The grave goods were sparse, including clothing elements (e.g., belt buckle and brooch) and tools (e.g., knife, comb, spindle whorl), resembling those found in other 'non-elite' cemeteries of the Great Hungarian Plain, such as Hajdúnánás–Fürj-halom-járás (M3 41/A). The burials contained the skeletal remains of two adults and one sub-adult. The skulls exhibited Europid characteristics, but Mongolid features, particularly in dental morphology, were also observed. Artificial cranial deformation was noted in one instance.

Grave No. 422, located 60 m southeast of the other three graves, constituted a separate group (group 6). Unlike the graves in group 5, it was oriented in a W–E direction with a minor deviation and was superimposed with a Sarmatian-period settlement object. Only an unidentified bronze artifact was recovered from this grave. The skeletal remains belonged to an adult, exhibiting Europid characteristics with Mongolid dental morphological features, similar to the individuals from group 5.

The graves were dated from the end of the 4th century CE to the mid-5th century CE. The burial customs suggested possible connections with the local Sarmatian-period population, yet specific features, such as the sidewall niches and Mongolid cranial characteristics point to a newly arrived Sarmatian group with Eastern European origins. We could include 3 samples in our analysis.

***OFU-190: Grave No. 190 (SNR 290)***

This grave, featuring a sidewall niche and oriented in a N–S direction, contained the skeletal remains of an adult female in a supine position. The skull showed Europid characteristics with Mongolid dental morphological features. Grave goods included two spindle whorls and an iron knife. Radiocarbon measurement, partially confirming the archaeological dating, placed the burial in the 5th century CE to the first third of the 6th centuries CE.

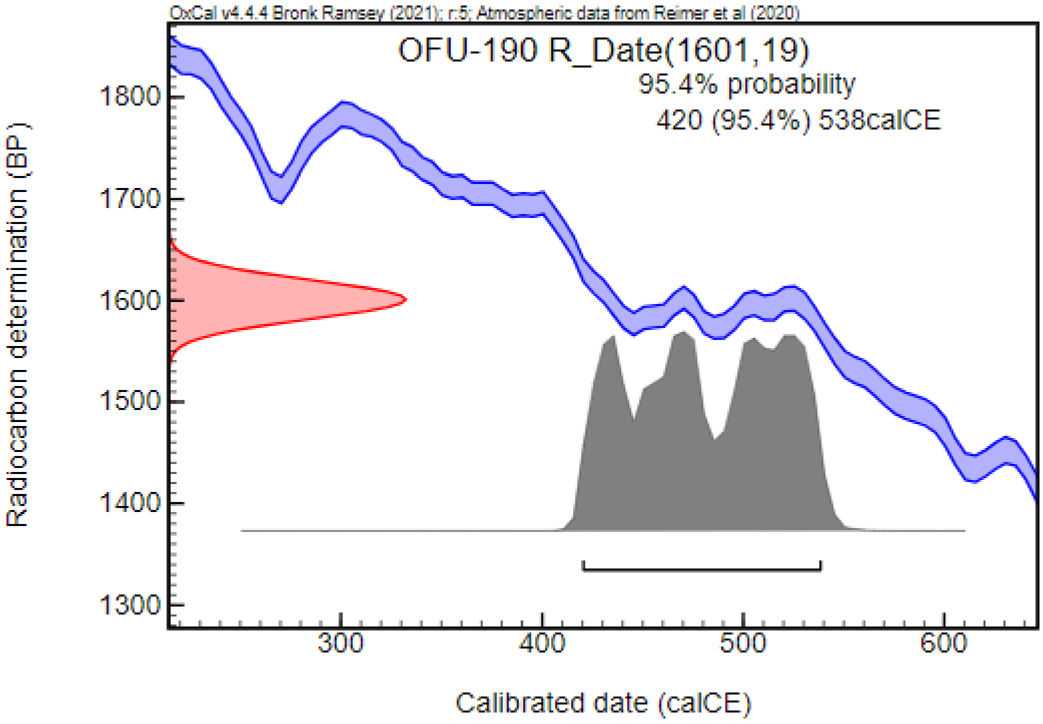


***OFU-191: Grave No. 191 (SNR291)***

The grave, oriented in a N–S direction, contained the skeleton of a sub-adult with possible Mongolid dental features, positioned supine. Recovered artifacts included a two-sided bone comb, a spindle whorl, and glass shards.

***OFU-422: Grave No. 422 (SNR 176)***

Oriented in a WNW–ESE direction and superimposed with a Sarmatian-period house pit, this grave contained the skeletal remains of an adult male, found in a supine position. The skull showed Europid characteristics with Mongolid dental features. The fragment of a bronze item was found in the grave. Radiocarbon measurement supported an earlier dating, placing the burial from the mid-4th century CE to the early 5th century CE, with a possibility dating to the second half of the 3rd century CE (12.4%).

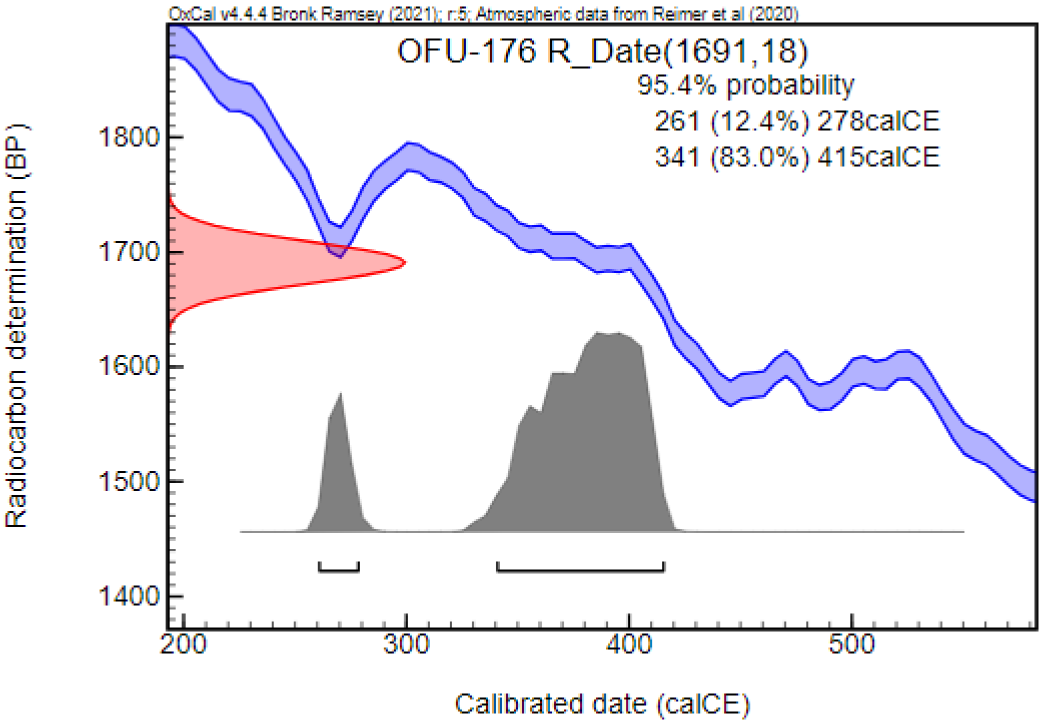


#### Óföldeák – Ürmös II. M43 10. lh. (OFU; Csongrád-Csanád County, Hungary) [99]

During the above-described excavations related to the construction of the M43 highway, settlement objects and 48 Sarmatian-period graves were discovered at site No. 10, ten of which were ditched. These Sarmatian graves were organized into three groups situated in different parts of the site. Group 1, composed of 18 graves, was located at the eastern edge of the Sarmatian settlement. Group 2, consisting of three burials, was located 180 m west of group 1. Group 3, with 27 burials, was a further 25 m west of group 2. The burials were primarily oriented in a SE–NW direction except for group 2, where a NW–SE orientation was observed. A significant number of the graves, 77%, had been partially or fully robbed, limiting the analysis. In one-third of the graves, traces of wooden coffins were detected, with only one burial containing iron coffin clamps. Artifacts recovered from the burials included jewelry, clothing fittings, tools (e.g., iron knives, spindle whorls), Roman coins, and vessels (ceramic, wooden, and glass). Jewelry and clothing elements found in female and sub-adult burials included earrings, neckrings, strings of beads, brooches, bracelets, belt buckles, and bead ornaments. Male burials, typically simpler in their inventory, contained fewer jewelry and cloth fittings, mostly brooches and belt buckles. The cemetery was dated from the second half of the 4th century CE to the first half of the 5th century CE. Artifacts indicated connections with the Roman Empire, the Black Sea region, as well as the Marosszentanna/Sântana de Mureş–Cherniakhov and Przeworsk Cultures. The archaeological materials of the cemetery have parallels with Hun-period burial grounds in the southern Great Hungarian Plane, such as Apátfalva–Nagyút-dűlő, Csongrád–Kenderföldek, Sándorfalva–Eperjes, and Tápé–Malajdok. Of the 48 graves, 45 contained human skeletal remains. Anthropological analysis conducted by Antónia Marcsik identified 16 sub-adults, 17 adult males and 16 adult females in the series. We could include 6 samples in our analysis from the three burial groups.

***OFU-15: Grave No. 15 (SNR17)***

This robbed grave, oriented in a NW–SE direction, contained the skeletal remains of an adult male in a supine position. Grave goods included an iron belt buckle, fragments of a bronze belt ornament, a ceramic vessel, and shards of other ceramic vessels.

***OFU-26: Grave No. 26 (SNR35)***

This robbed burial contained the skeleton of an adult male, in a supine position and oriented in a NW–SE direction. Artifacts recovered included a two-piece iron belt buckle, a fragment of an iron belt buckle, and shards from multiple ceramic vessels. Traces of a wooden coffin were also documented.

***OFU-27: Grave No. 27 (SNR36)***

The skeletal remains of an adult male, placed in a supine position, were uncovered in this grave, oriented in a NW–SE direction. Traces of a wooden coffin were discovered in the grave pit. In addition, a ceramic jug and shards belonging to a mug and other vessels were found in the burial.

***OFU-93: Grave No. 93 (SNR131)***

This robbed grave, oriented in a SE–NW direction, contained the skeleton of a sub-adult (6–7 years old) in a supine position, surrounded by traces of organic material, possibly from a shroud. Grave goods included two bronze earrings, a silver torc, coral and carnelian beads under the jaw, coral beads around the wrists, and shards of a ceramic vessel.

***OFU-138: Grave No. 138 (SNR182)***

The skeletal remains of a sub-adult (juvenile), in a supine position, were discovered in this grave, oriented in a SE–NW direction. While no traces of robbery were documented, the position of a coffin clamp found in the soil and the preservation of some artifacts suggest partial robbing. Artifacts recovered from the grave included a bronze earring, a string of glass and amber beads around the neck with iron and bronze pendants, glass and ceramic pendants, an iron brooch, three bronze bracelets, an iron knife, a ceramic bowl, and bead ornaments around the lower legs, including more than 305 glass beads and 59 amber beads. An iron coffin clamp and an unidentified deformed iron artifact were also found in the grave soil. Radiocarbon measurement, supporting an earlier date, placed the burial from the second third of the 3rd century CE to the second third of the 4th century CE.

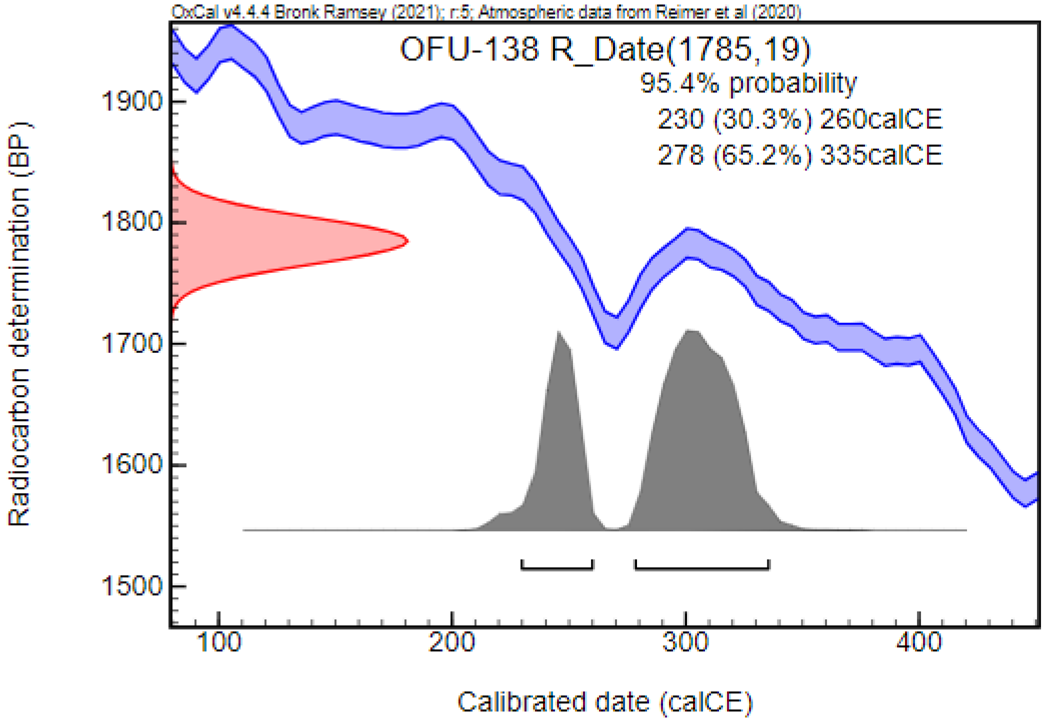


***OFU-168: Grave 168 (SNR221)***

The skeleton of a young-adult female in a secondary position was found in this robbed grave, oriented in a SE–NV direction. Grave goods included a tube-shaped gold plate near the skull, a string of coral beads near the neck, a two-piece bronze brooch, a bronze box brooch with a golden plate, a two-piece bronze belt buckle, two amber beads associated with the belt, a fragment of a bronze wire belt ornament, two cowries, and further amber and glass beds near the hip region. A Kowalk-type glass, a small paint pot, and a bronze mirror were also recovered. Further bronze fragments, possibly belt ornaments were found in a secondary position in the SE corner of the grave. Radiocarbon measurement, supporting an earlier date, placed the burial from the end of the first third of the 2nd century CE to the beginning of the second third of the 3rd century CE. Based on the currently established chronology of Kowalk-type glass cups, a significant discrepancy can be observed between radiocarbon dating and archaeological dating. this case, the archaeological dating appears to be more reliable.

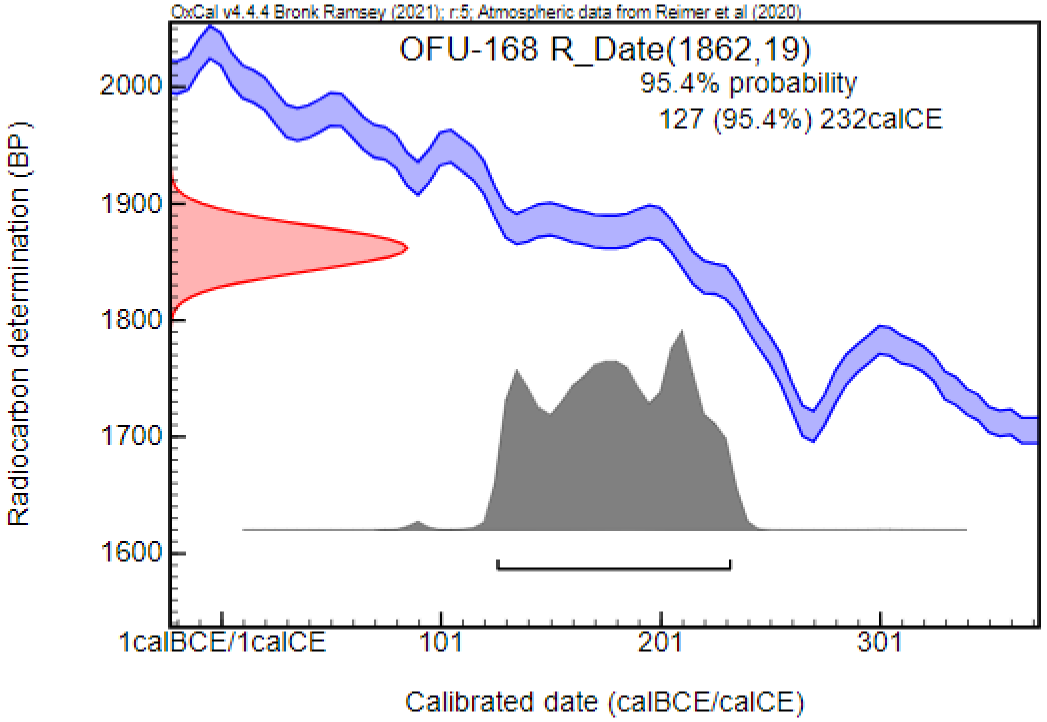


#### Oltenița – Renie (1968) (OLR; Călărași County, Romania) [100]

The archaeological site is situated on the edge of a low terrace in the close vicinity of Oltenita. In addition to findings dated to the Eneolithic and Medieval periods, 16 Sarmatian burials were uncovered between 1960 and 1967. However, the cemetery is considered only partially excavated, and some of the graves were destroyed by construction works in 1966–1967. The Sarmatian graves were in groups of 4–6 graves, but archaeological data is available only for nine graves in the literature. These graves were oriented in a N–S direction with minor variations toward NNE–SSW. The artifacts recovered from the burials included various types of ceramic vessels, beads, a knife, and a bronze mirror. The burials were dated to the 2nd–3rd centuries CE. A total of two sub-adults and possibly five adults are suspected in the series, but no anthropological analysis was conducted on the material. Only one sample could be involved in our analysis.

##### *OLR-2: Grave No. 2*

The skeletal remains of a sub-adult individual (possibly female) were found in the grave, oriented in a N–S direction. The grave goods included a ceramic jar. Radiocarbon measurement supports an earlier date, positioning the burial from the second half of the 4th century BCE to the middle of the 1st century BCE.

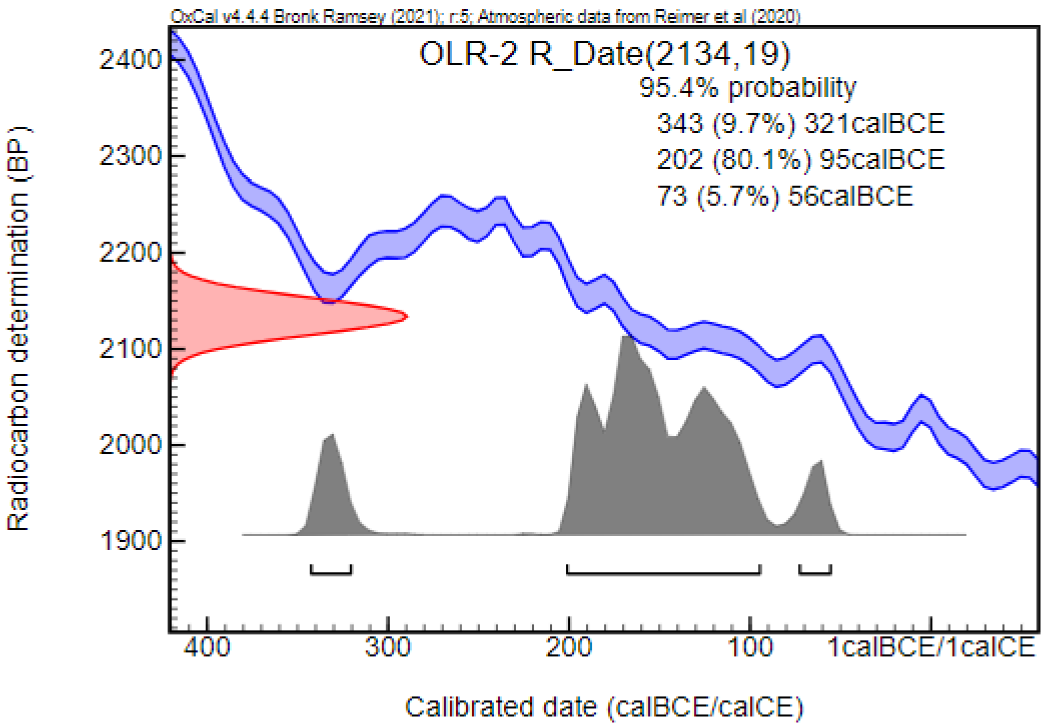


#### Oltenița – Ulmeni (1960) (OSU; Călărași County, Romania) [100,101]

The archaeological site is located 1 kilometer from the village of Ulmeni and 5 kilometers from Oltenița. The burial ground was accidentally discovered in 1957 and 1960 during the construction of an irrigation system, which led to the destruction of several inhumation graves. During these incidents, two Sarmatian graves were uncovered. Following these accidental finds, systematic excavations began in 1960, resulting in the discovery of five inhumation graves, three of which were identified as Sarmatian. Thus, a total of five Sarmatian graves were documented at the burial ground. The burials were mostly oriented along a N–S axis, with occasional deviations towards NNE–SSW or WNW–ESE directions. The position of the deceased in the grave pit was observed in only one case, as most of the burials had been affected by contemporary or recent disturbance. In this case, the body was found in a supine position with straightened arms. Two sub-adults were identified based on the descriptions of the archaeologist leading the excavation. However, no anthropological analysis was conducted on the skeletal remains. The artifacts recovered from the graves included ceramic vessels, (among them Roman mugs), multiple glass, amber, and metal beads near the neck and feet, a bronze brooch, bronze earrings, a bronze mirror, and a bronze bell. Ceramic vessels, displaying characteristics of both Sarmatian, Dacian, and Roman traditions, were the most common objects in the cemetery, found in four out of five graves. The dating of the site is complex, as the material contained artifacts from both an earlier (end of the 1st century CE) and a later period (2-3rd century CE). The fragmentary data did not allow researchers to draw final conclusions, leading to two hypotheses: either this was the cemetery of a Sarmatian community that appeared early and continuously occupied the territory until the end of the 2nd century CE, or, considering the limited number of graves, it could have been used by different communities in two separate chronological phases.

##### *OSU-1: Grave No. 1*

The grave containing an adult female, oriented in a N–S direction, was discovered during the construction of the irrigation system. The burial contained two ceramic jars, two ceramic jar lids, multiple glass beads, and a Roman bronze brooch, dating the grave to the late 1st and early 2nd centuries CE. Radiocarbon measurement supports an earlier date, placing the burial from the middle third of the 2nd century BCE to the middle of the 1st century BCE.

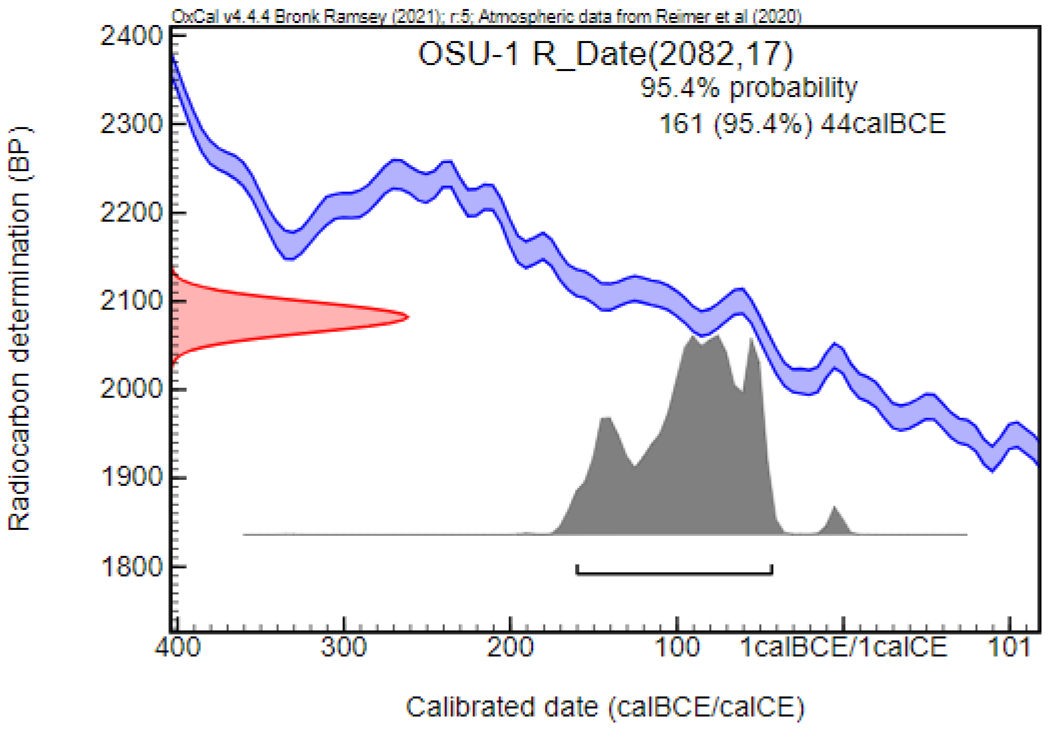


#### Páty – Malom-dűlő, 9. lh. (PMD; Pest County, Hungary) [102]

In 1997–1998, preventive excavations during the construction of the M1 highway section near Páty uncovered a multi-period cemetery with approximately 910 graves. The earliest burials date to the Bronze Age, while the latest are from the Árpádian-Age, with continuous use of the burial ground beginning in the 1st–2nd centuries CE. A distinct group of 12 graves was discovered at the NW border of the Late Roman cemetery. These burials differed from the Late Roman graves in both burial customs and grave goods, suggesting the presence of a different population. The graves were oriented in a W–E direction, with the deceased placed in a supine position. The cemetery predominantly featured single burials, except for one grave that contained bone fragments of a second individual, likely due to contemporary grave robbing. Of the 12 graves, 7 were affected by such robbing, and an additional two grave pits were empty, lacking both human remains and artifacts. Traces of wooden coffins were found in two cases. The number of the grave goods was low, likely due to the contemporary robbing, with artifacts recovered from only four graves. These included jewelry (bronze beads), cloth fittings (e.g., belt and shoe buckles, brooches), bone combs, tools (iron knives, needle case, and needle), and glass vessels. The archaeological characteristics of this cemetery, including grave goods and burial customs, closely parallel those of the Late Roman and early Migration Period cemeteries in the Transdanubia region, such as Csákvár, Visegrád–Gizellamajor, Lébény, Lengyeltóti, and Intercisa. The cemetery is dated to the first half of the 5th century CE and is associated with Eastern Germanic groups. The preservation of human skeletal remains was generally poor. In total, six adults and three sub-adults were identified, including three females and three males. We could include one sample in our analysis.

***PMD-564: Grave No. 564***

This grave, oriented in a W–E direction, contained the skeleton of an adult female in a supine position. Grave goods included bronze spiral beads near the neck and two silver brooches. In addition, traces of a wooden coffin were documented.

#### Pécs – Málom (PM; Baranya County, Hungary) [103,104]

In the spring of 1992, a burial was destroyed during construction work related to a Leisure Centre. Only a few human bones, an iron knife, and an iron belt buckle were saved. Following this discovery, a rescue excavation led by Erzsébet Nagy and Gábor Kárpáti uncovered four graves. However, the cemetery is considered only partially excavated. The graves were primarily oriented in a W–E direction, with the skeletons found in a supine position. In some instances, slight color changes in the soil suggested the presence of wooden coffins. The burial inventory was rather sparse, consisting of jewelry (an earring and a hair ring), clothing elements (belt buckles), iron knives, and bone combs. The cemetery is dated to the first half of the 5th century CE and was associated with the population of the Alanian, Eastern Gothic, and Hunnic Alliance. Anthropological analysis conducted by Ferenc Szalai identified one sub-adult and four adult skeletons, including two females and two males. In two cases, traces of artificial cranial deformation were detected. We could include one sample in our analysis.

***PM-5: Grave No. 5***

This grave, oriented in a W–E direction, contained the skeleton of an adult female in a supine position. Grave goods included a bronze earring with polyhedron pendant and a two-sided bone comb.

#### Pogorăști (POG; Botoșani County, Romania) [105–107]

In the northwestern sector of Pogorăști village, 28 burials from a Sarmatian cemetery were uncovered during rescue excavations between 1960 and 1962, following accidental discoveries in 1958. The burials were mostly oriented in a north-south direction, with skeletons found in a supine position with straightened extremities. The archaeological materials recovered included ceramic vessels (among them a Roman bowl), beads, belt buckles, and iron knives. However, only the archaeological data for the first five graves were published. Additionally, anthropological data for three individuals (Graves No. 2–4) is available, all of whom were determined to be adult males. The cemetery is generally dated to the 2nd–3rd centuries CE. Only one sample could be included in our analysis.

***POG-10: Grave No. 10***

The skeletal remains of an adult male exhibiting signs of artificial cranial deformation were discovered in Grave No. 10. The specifics of the burial customs and grave goods associated with this individual are unknown. Radiocarbon dating suggests an earlier period, placing the burial from the mid-1st century BCE to the last third of the 1st century CE.

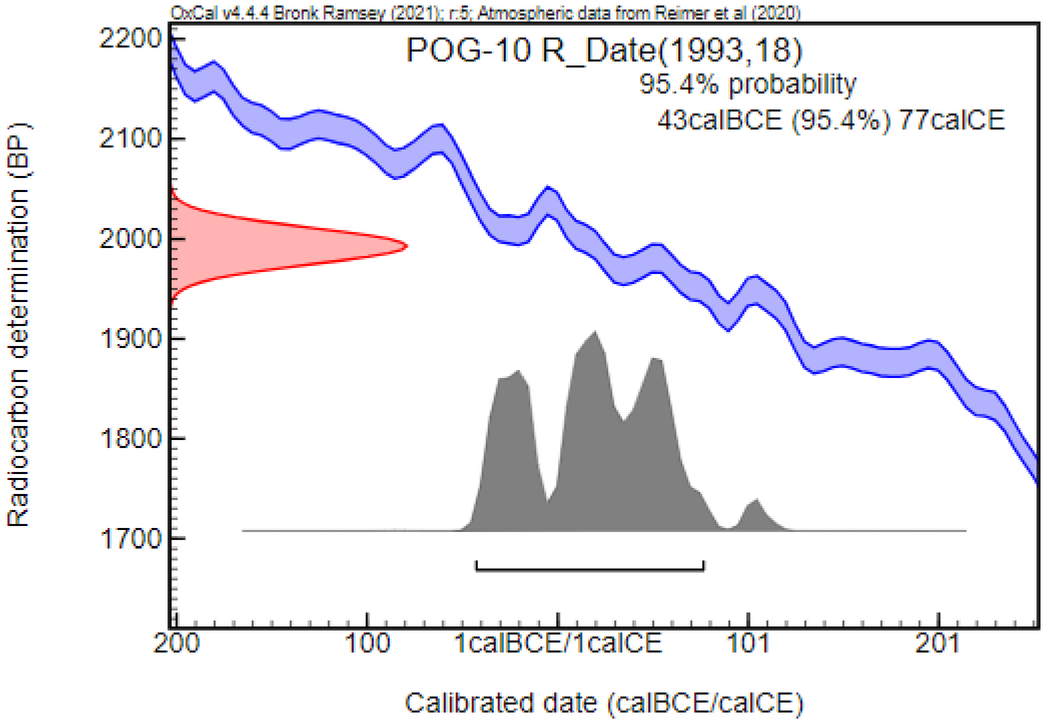

#### **Probota (PRO; Iași County, Romania)** [108,109]

Archaeological investigations were conducted by E. Zaharia and N. Zaharia between 1959 and 1961 at the northwestern edge of the village of Probota. The findings included burials from the Bronze Age, Iron Age, and Sarmatian period. Unfortunately, the site remains unpublished, and contradictory information exists in the literature regarding the total number of Sarmatian graves, with estimates ranging from 12 to 25 (summarized in [110]). Many graves were previously disturbed due to clay or sand mining activities. While the orientation of the burials varied, the skeletons were mostly found in a supine position with straightened extremities, except for one instance where the arms were placed on the stomach.

Based on the known data, the archaeological materials recovered included ceramic vessels, earrings, beads (used both as jewelry and clothing ornaments), belt buckles, and mirrors. One grave contained animal bones, possibly as food offerings. The cemetery is dated to the 2nd and 3rd centuries CE. The anthropological material recovered from the site has not been subjected to a detailed study. Two samples could be involved in our study.

***PRO-37: Grave No. 37/1960***

The skeletal remains of a sub-adult (*Infantia* I) with an artificially deformed skull were found in the grave. The grave goods are unknown. Radiocarbon dating confirmed the archaeological dating of the site, placing the burial in the 2nd and 3rd centuries CE.

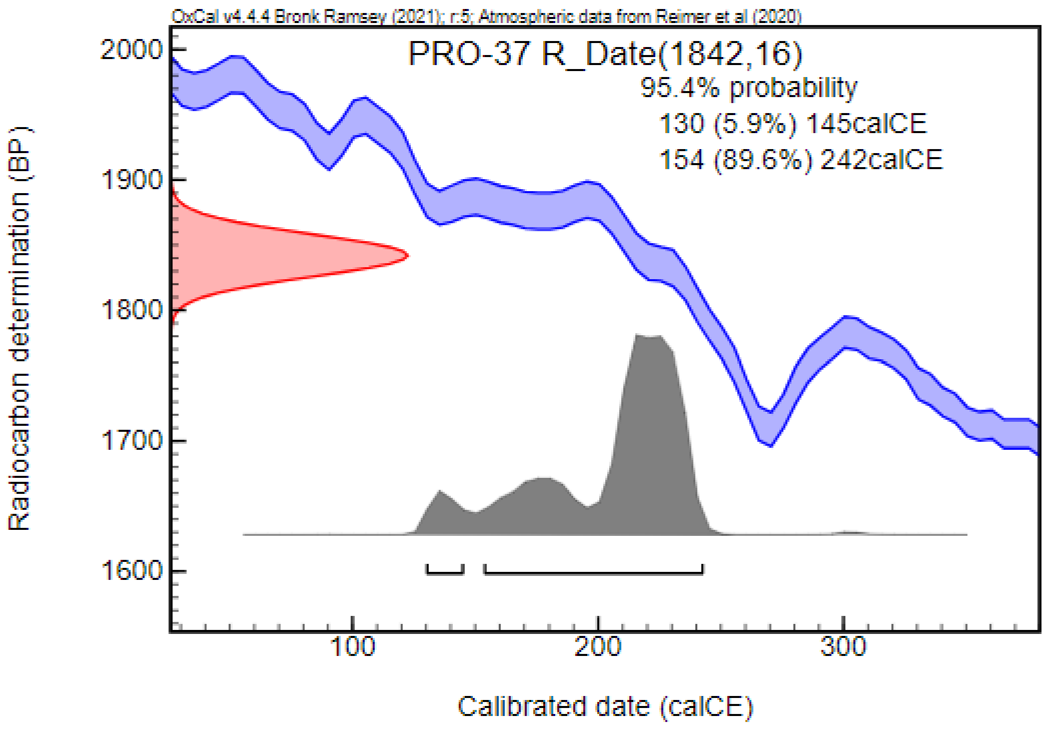


***PRO-47: Grave No. 47/1961***

This grave contained a skeleton with an artificially deformed skull, beads, and a mirror. However, detailed archaeological and anthropological data are unknown. Radiocarbon dating confirmed the archaeological dating of the site, placing the burial in the 2nd and 3rd centuries CE.

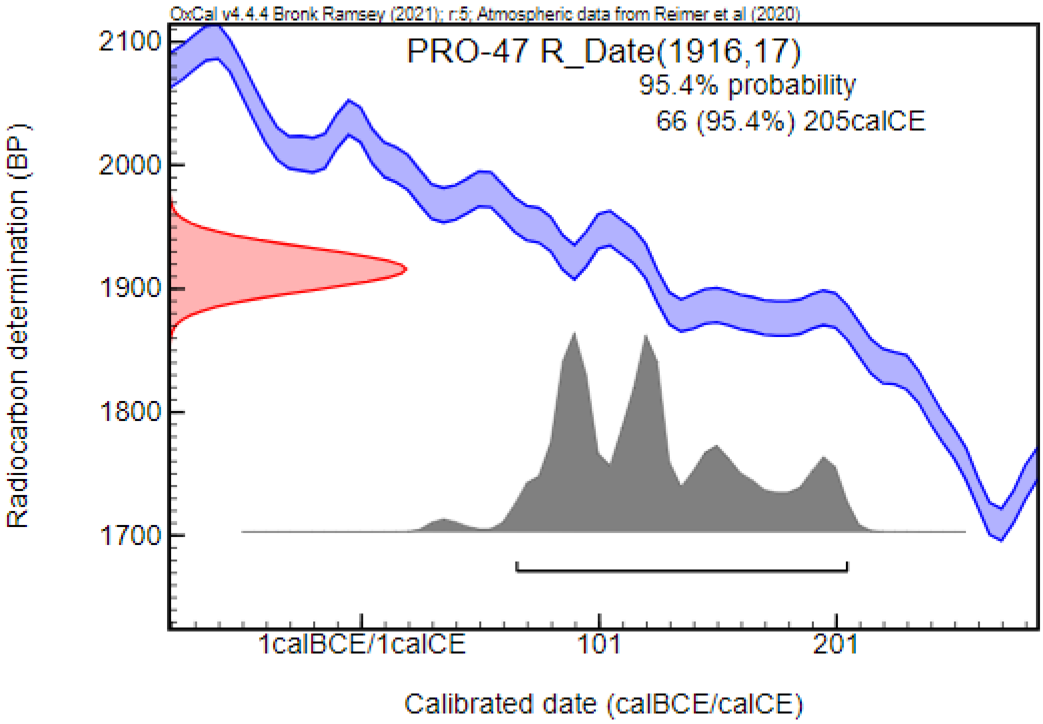


#### Püspökladány – Görepart (PLG; Hajdú-Bihar County, Hungary) [111,112]

A total of nine graves were documented during a rescue excavation led by Ibolya M. Nepper in 1973. Unfortunately, most of the graves were destroyed due to previous soil work, and the cemetery was only partially excavated. The archaeological findings comprised various artifacts such as jewels (e.g., beads and brooches), ceramic vessels, Roman coins, and a spindle-whorl. Notably, one of the graves was encircled by a ditch. The cemetery was dated to the 2nd and 3rd centuries CE. The anthropological analysis of skeletal remains was carried out by Antónia Marcsik, revealing that bone preservation was generally low. Of the six graves containing human remains, one sub-adult and four adults were identified—two females and two males. However, the age-at-death and biological sex of one individual remained uncertain due to poor bone preservation. Only one skull was suitable for craniometric and morphological analysis, exhibiting similarities with individuals from the Sarmatian period in the Lower Volga region, particularly Western Kazakhstan. We could involve three samples in our analysis.

***PLG-1: Grave No. 1***

The grave, oriented in the NE–SW direction, contained the skeletal remains of a juvenile/ young adult female interred in a supine position with almandine and glass beads.

***PLG-4: Grave No. 4***

The skeletal remains of a middle adult male were found in the robbed burial. The grave, oriented in the N–S directions did not contain any known grave goods. The postcranial bones displayed overall robustness, and the skull was brachycranic, suggesting possible Europid (Cromagnoid) features. Radiocarbon dating placed the burial between the second third of the 3rd century CE and the first third of the 4th century CE.

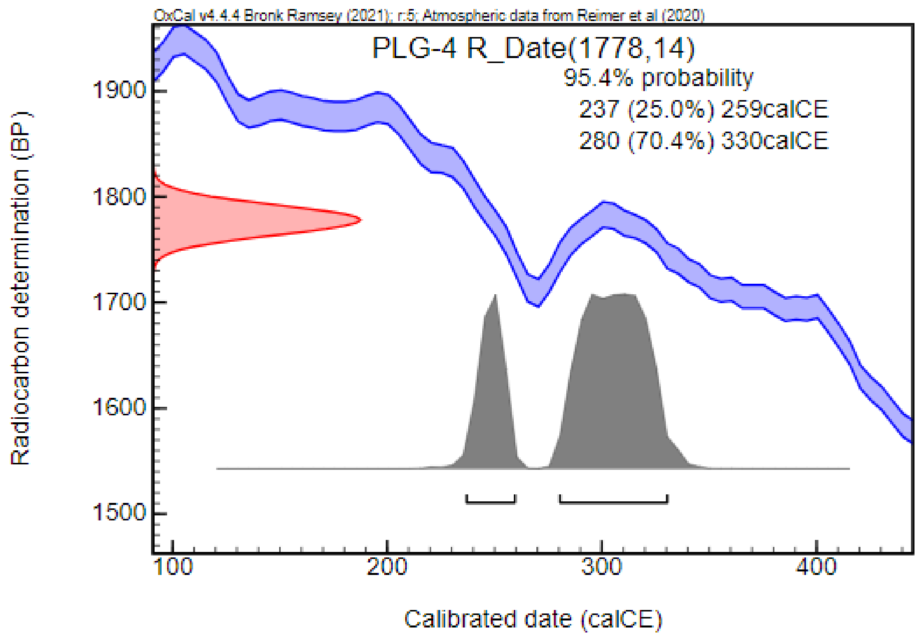


***PLG-5: Grave No. 5***

The skeletal remains of a middle adult male were discovered in the NE–SW oriented grave. Only an iron fragment was documented in the previously robbed burial.

#### Rákóczifalva – Bivaly-tó, Rokkant-föld II. 4. lh (RBR; Jász-Nagykun-Szolnok County, Hungary) [113]

In 2006, preventive excavations were conducted by Marietta Csányi and Judit Tárnoki in the area of Rákóczifalva–Bivalytó. At the 4th site of Rokkant-föld II, nine Sarmatian-period burials were discovered. The archaeological analysis is still in progress, and only preliminary data are available. Most of the burials were robbed, limiting the evaluation of the finds. Anthropological analysis was conducted by Tamás Hajdu and Zsolt Bernert. The preservation of the bones was generally poor, which limited the analysis and evaluation of the findings. Of the nine Sarmatian burial containing human remains, eight were adults and one was sub-adult, including three females and five males. We could include one sample in our analysis.

***RBR-118: Object No. 118***

This grave contained the fragmented skeletal remains of an adult male. Radiocarbon measurement, consistent with the archaeological findings, dated the burial to the Sarmatian period, ranging from the mid-3rd century CE to the last third of the 4th century CE.

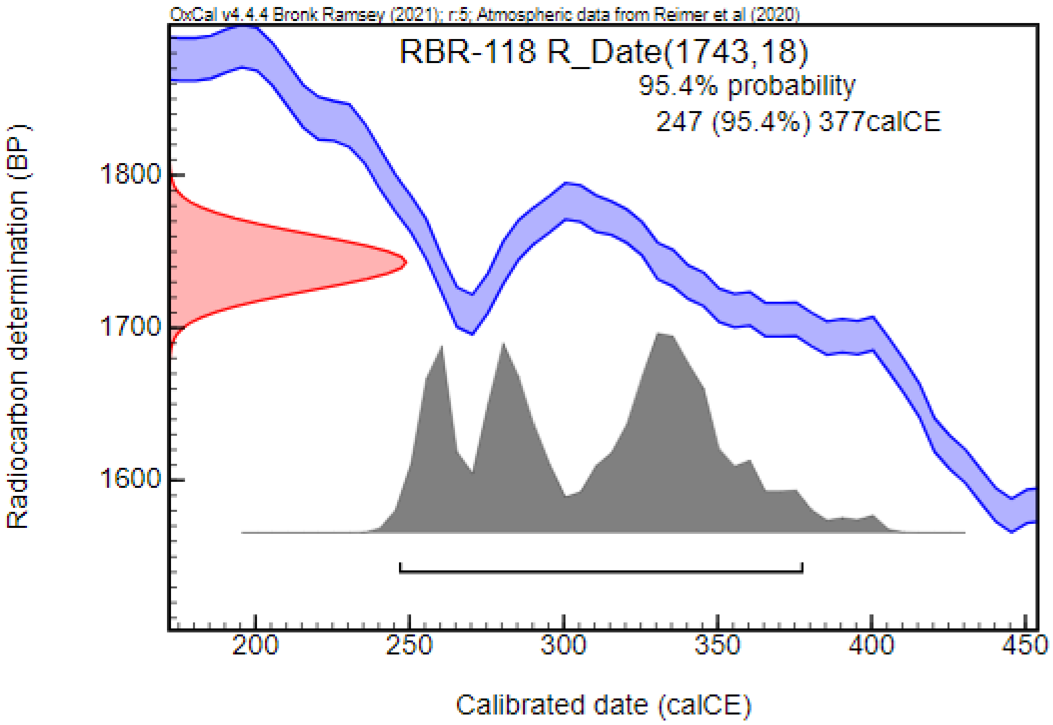


#### Râmnicelu (1969) (RAM; Brăila County, Romania) [90,114]

Systematic excavations were conducted between 1968 and 1970, near Râmnicelu village, located 34 km west of Brăila, on the upper terrace of the right bank of the Buzău River. These excavations uncovered a Neolithic settlement and 20 graves – dated to the Bronze Age, Iron Age, Sarmatian period, and Medieval period. Initially, 18 burials were classified as Sarmatian; however, two of these were later identified as belonging to the Scythian period. The Sarmatian graves, forming two groups in the northern half of the settlement, were mostly oriented in a W–E direction, with occasional deviations toward SW–NE and N–S. The skeletons were found in a supine position, primarily with straightened arms. The archaeological material included ceramic vessels (among them a Roman import bowl), beads, belt buckles, metal ornaments, a spindle whorl, and a spearhead. This cemetery was categorized among the earlier Sarmatian groups in the area, dated to the late 1st and 2nd centuries CE. Based on the archaeologists' field notes, a total of 7 adults and 8 sub-adults were identified. However, no anthropological analysis was conducted on the skeletal remains. Initially, we could include two samples in our analysis. Radiocarbon dating, however, indicated that grave No. 7 dates to the Iron Age, leading us to exclude this case from the Sarmatian sample list.

##### *RAM-13: Grave No. 13*

The skeletal remains of an adult female were found in a supine position with straightened arms in a grave oriented in a W–E direction. The grave goods included fragments of a bronze object (in the form of a plate, fixed with nails), wood and leather fragments, another bronze plate (fixed with three bronze nails), and five fragments made of bronze wire (shaped like links).

#### Ripiceni – La Stâncă (RLS; Botoșani County, Romania) [115]

Two graves and additional stray finds from the Sarmatian period were discovered in Ripiceni commune, on the right bank of the Prut River, in 1979 and 1983. One of the graves contained the skeleton of an adult female, while a 2–3 years old sub-adult was found in the other. Traces of artificial cranial deformation were discovered on the skulls of both individuals. The burials were found by chance, so no information is available concerning the burial customs or the position of grave goods inside the grave pits.

The site was dated from the second half of the 2nd century CE to the 3rd century CE. We could include one sample in our analysis.

***RLS-1: Grave No. 1***

The skeleton of an adult female with traces of artificial cranial deformation was found in the grave. The grave goods included fragments of one or several brooches possibly with returned foot, fragments of two bronze earrings (?), a spindle whorl, shards of a ceramic vessel, and a total of 197 glass beads. Radiocarbon dating, partially overlapping with the archaeological dating, placed the burial between the 1st century CE and the first third of the 2nd century CE.

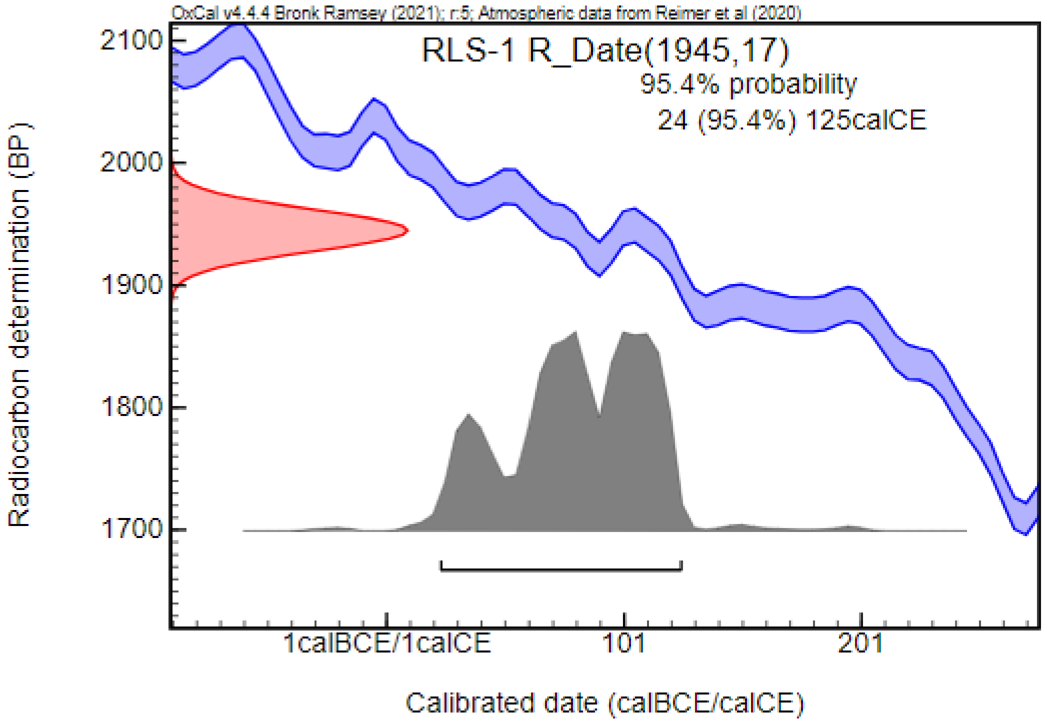


#### Solt – Polya-fok (SPF; Bács-Kiskun County, Hungary) [116]

***SPF-1: Object No. 1 (SNR15)***

In the summer of 2020, a solitary grave was uncovered during the excavation led by Nikoletta Lukács and Eszter Patyi, prior to the construction of a sludge tank for a wastewater treatment plant. A detailed archaeological analysis is still in progress, and only preliminary data are available regarding the burial customs and recovered grave goods. The simple grave pit, oriented in a NW–SE direction, contained fragmented skeletal remains mostly in a secondary position due to contemporary robbery. Only the two femurs were assumed to be found in situ, suggesting the deceased was placed in a supine position into the grave. Grave goods included a bronze earring with spherical or polyhedral pendant, a bronze belt buckle, a string of amber, chalcedony, and glass beads, and a ceramic vessel. The burial was dated to the 5th century CE, showing parallels with sites and artifacts from the Hun period and the following decades. Anthropological analysis conducted by Olga Spekker and colleagues revealed that the fragmented skeletal remains belonged to an adult female. Radiocarbon measurement, supporting the archaeological dating, placed the burial between the first third of the 5th century CE and the second third of the 6th century CE.

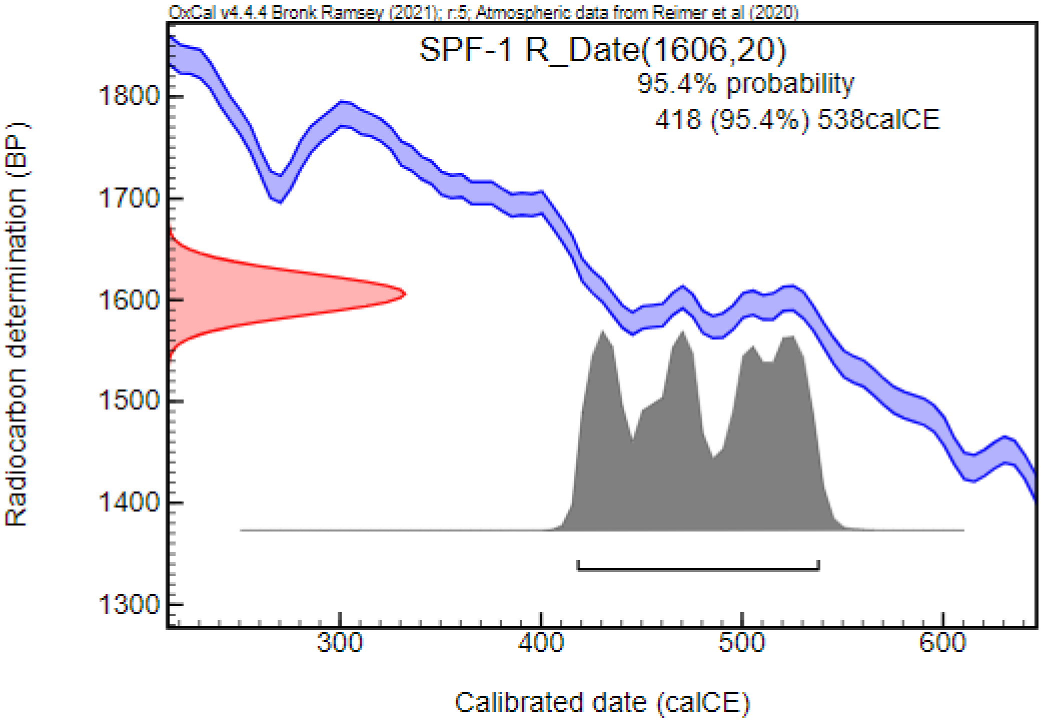


#### Szabadszállás – Boczka tanya (SZB; Bács-Kiskun County, Hungary) [117–119]

##### *SZB-1: Grave No. 1*

During peat extraction along the main channel of the Danube Valley in the early 1960s, prehistoric finds were discovered. Rescue excavations led by Elvira H. Tóth uncovered the remnants of a Bronze Age settlement and a solitary grave. Initially measuring 150 by 285 centimeters, the large grave pit gradually narrowed and was only 65 centimeters wide at a depth of 170–175 centimeters. At the bottom of the pit, in a trough-like dip, the skeletal remains of an adult male with Europid (Mediterranean) characteristics on the skull were found in a supine position, oriented in a W–E direction. The skull and cervical vertebrae were in an unusual position, with the skull cap facing toward the body. No traces of decapitation were observed on the bones, suggesting the head was carefully severed post mortem and placed upside-down in the grave, as described by the excavating archaeologist. Artifacts found in the burial included an iron hoe, an iron spade head, and a richly ornamented, cone-shaped glass made of gilded purple-violet glass. The ornamentation on this unique glass has parallels in the early Christian Egyptian art, particularly textiles. Based on this artifact, the burial was dated to the end of the 4th and the first half of the 5th centuries CE. Radiocarbon measurement, partially overlapping with the archaeological dating, placed the burial from the middle of the 3rd century CE to the middle of the 4th century CE.

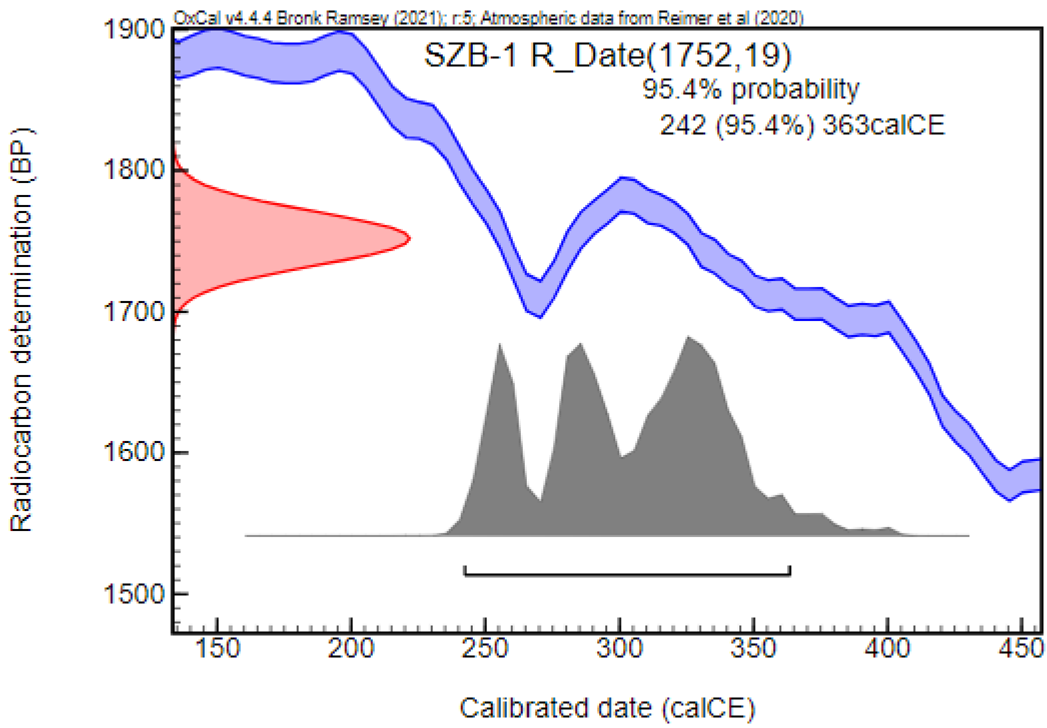


#### Szeged – Csongrádi út (CSU2; Csongrád-Csanád County, Hungary) [120–123]

Thirteen 10th-century CE burials and thirty-one Sarmatian graves were rescued during excavations led by Béla Kürti at Szeged, near the road to Csongrád, in the gardens of the Kiss Ferenc Forestry Vocational High School between 1974 and 1987. The archaeological and anthropological investigation of the Sarmatian burials is still in progress, and only brief summaries and preliminary data are available. Generally, the graves were oriented in an S–N direction with rectangular pits of large dimensions. Traces of wooden coffins were documented in several cases. The inventory of the male graves was rather poor compared to their female counterparts, with ceramic vessels and iron knives being the most characteristic artifacts. Female burials included jewelry and cloth fittings too such as brooches and beads in various types, colors, and functions. The burial ground was dated to the 2nd and 3rd centuries CE, with the possibility that some graves belonged to a somewhat earlier or later phase. The preservation of the skeletal remains was generally poor, with 22 adults and 3 sub-adults identified in the series. Unfortunately, the skeletal remains from burials with a known archaeological background (i.e., burial customs and grave goods) were not fit for archaeogenetic analysis, and we could include only two samples in our analysis.

***CSU2-39: Grave No. 39***

The fragmented skeletal remains of an adult, possibly male individual, were found in this grave. The grave goods are unknown.

***CSU2-43: Grave No. 43***

The burial contained the skeleton of an adult male individual. The grave goods are unknown. Radiocarbon measurement confirmed that the burial dates to the Sarmatian period, placing it from the beginning of the 1st century CE to the first third of the 2nd century CE.

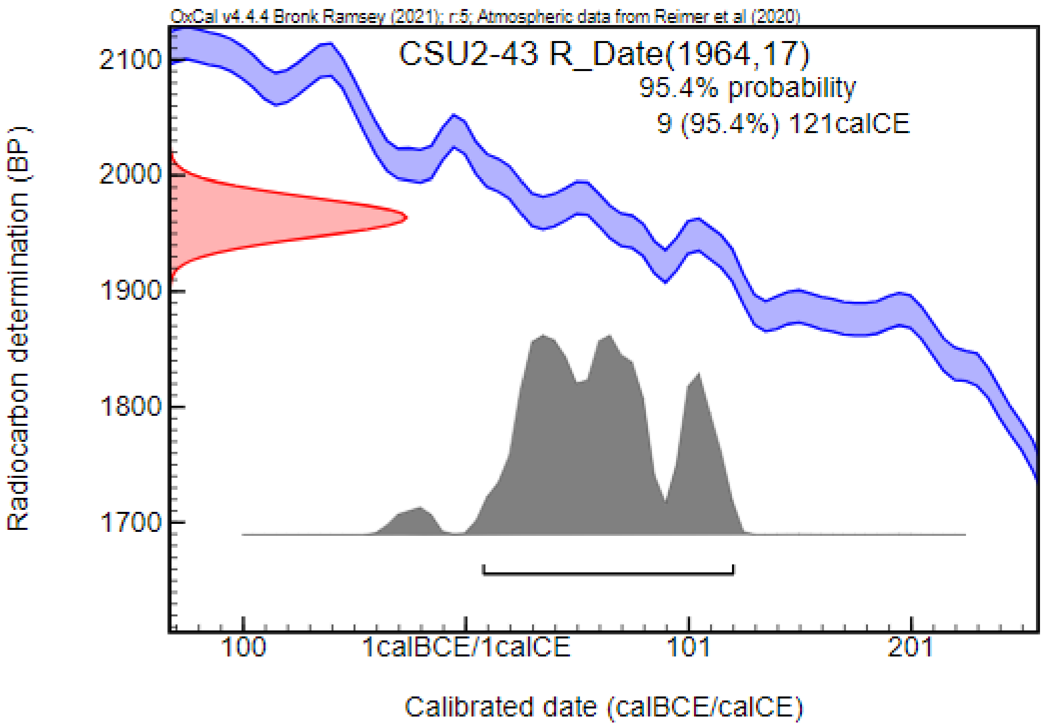


#### Szihalom – Pamlényi tábla (SPT; Heves County, Hungary) [124,125]

A multi period site with more than 1500 settlement objects and 31 burials was excavated between 1995 and 1997, in connection with the construction of the M3 highway near Szihalom. The archaeological and anthropological analysis are still in progress, and only preliminary data are available. Twenty-four graves were dated to the early Migration Period. Most of the burials had been robbed, but the burial customs (such as the presence of a ditched grave) and the grave goods were associated with the Sarmatians. Additional artifacts suggested connections with Germanic groups. The cemetery was dated to the 4th and 5th centuries CE. We could include one sample in our analysis.

***SPT-23: Grave No. 23***

The Sarmatian-period burial was superimposed onto a Bronze Age feature, oriented south to north. The grave pit was rounded and slightly rectangular in shape. This burial contained the skeletal remains of an adult female with Europid characteristics on the skull. The grave had been looted, likely during the decomposition of the body, as suggested by the greater distance between the upper body and the pelvis, along with the unnatural positioning of the upper body. Most of the grave goods, except for the brooch, were found between the upper body and the pelvis. The grave contained a bronze brooch, 15 polyhedral-shaped carnelian beads, one white cylindrical glass bead, a folded cylindrical metal object, an iron knife, an iron buckle, an iron object, and a bronze coin. During the excavation of the grave fill, an additional 16 glass beads were recovered from the chest area. Radiocarbon measurement, partially supporting the archaeological findings, placed the burial to the 4th–5th centuries CE, with a possibility being dated to the second half of the 3rd century CE.

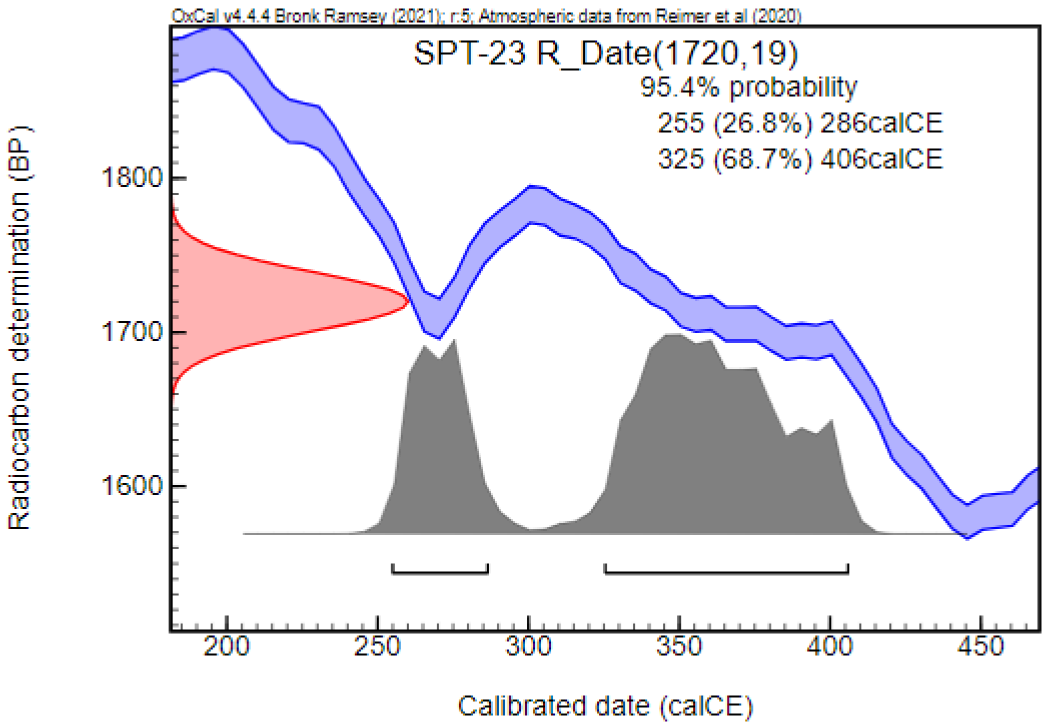


#### Tápé – Malajdok A (TMA; Csongrád-Csanád County, Hungary) [126]

In the vicinity of Tápé, in the area of Malajdok, excavations led by Móra Ferenc uncovered 17 Sarmatian-period graves in 1931. In 1943, Dezső Csallány conducted further excavations next to the site, uncovering an additional 36 burials. Due to incomplete documentation from these early excavations, the structure and relationship of these two sites—whether they belong to the same burial ground or not—remain unknown [86]. The burials were primarily oriented in a S–N direction, with deviations toward SE–NW, SSE–NNW, and SW–NE. The positions of the skeletons in the graves are unknown. Iron coffin clamps were found in several graves, and it was sometimes assumed that the deceased were wrapped in reed or bulrush. Artifacts included jewelry (e.g., strings of beads and torcs), clothing elements (e.g., brooches, belt buckles, beads), mirrors, ceramic vessels, Roman coins, copies of Roman coins, tools (e.g., iron knives, spindle whorls. fire-lighting tools), and weapons (e.g., spearheads). The cemetery was dated to the 4th and 5th centuries CE. Although Mihály Párducz initially proposed that the Tápé–Malajdok site and similar cemeteries in this region were associated with Germanic groups, particularly Gepids, subsequent archaeological research revealed that the burial customs and artifacts were primarily associated with the Sarmatians, with close parallels found in the Tiszadob-type cemeteries of the Upper-Tisza region [75]. It is becoming increasingly evident that, as a result of the great migrations and population movements, a wider spectrum of archaeological materials characteristic of the Maroszentanna-Chernyakhov culture appears in the cemetery's assemblage. The anthropological investigation conducted by Antónia Marcsik is still in progress. Most of the human remains were not recovered from the graves or were lost over the decades, leaving bones –mostly skulls–from only 24 burials available for analysis. All were identified as adults, with 10 males and 14 females. Primarily Europid characteristics were described on the skulls, with Mongolid features present in a few cases. We could include five samples in our analysis.

***TMA-2: Grave No.2***

This completely robbed burial, found during the excavation led by Ferenc Móra, contained the skeletal remains of an adult female accompanied by a ceramic vessel.

***TMA-8: Grave No. 8***

This grave, oriented in a SE–NW direction, was discovered during the excavations led by Ferenc Móra. The skeleton of an adult male with Europid characteristics on the skull was recovered, assumably wrapped in reed or bulrush. Grave goods included a bronze ring under the hip, a bronze belt buckle, a fragment of an iron brooch, fragments of a short iron sword or dagger, an iron knife, iron fragments, and a copy of a Roman coin (possibly copied from a Marcus Aurelius or Lucis Verus denarius).

***TMA-9: Grave No. 9***

The skeleton of an adult individual was found in this S–N oriented grave, discovered during the excavation led by Ferenc Móra. Although archaeological data suggest it was a female burial, the human remains were identified as male. Grave goods included a string of amber, glass, and stone beads, a spindle whorl, a bronze ring, a fragment of an iron knife, and a ceramic vessel. Radiocarbon dating, partially overlapping with the archaeological findings, placed the burial between the second half of the 3rd century CE and the early 5th century CE.

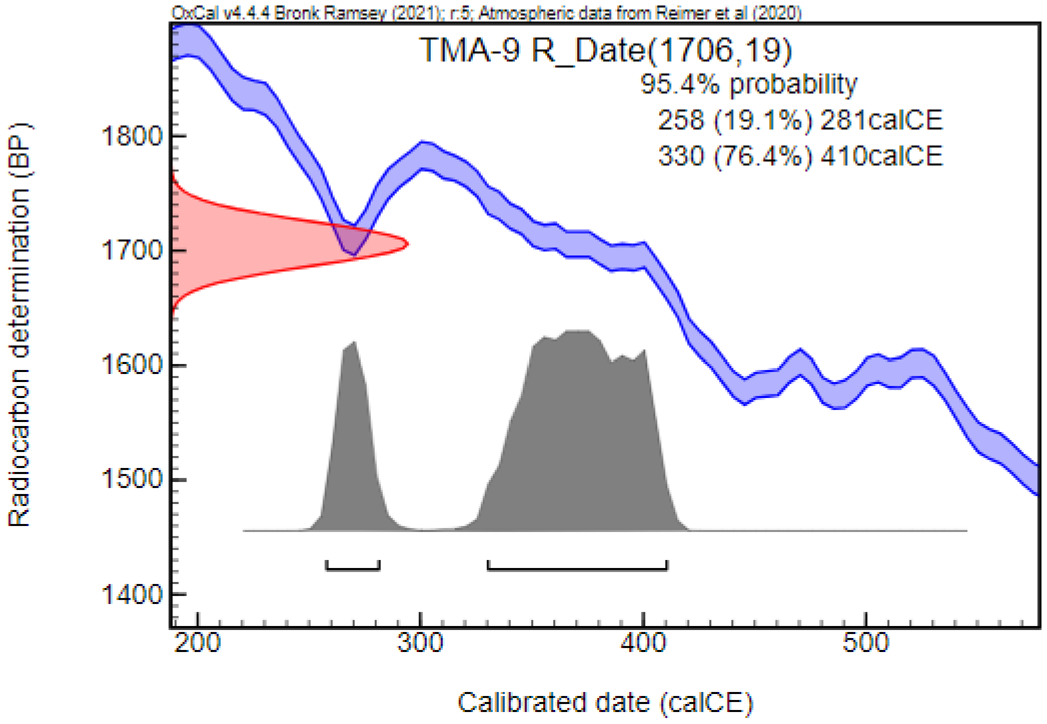


***TMA-10: Grave No. 10***

This S–N oriented grave was found during the excavation led by Ferenc Móra. The skeleton of an adult female with Europid characteristics on the skull was uncovered without known grave goods.

***TMA-36: Grave No. 36***

This burial was discovered during the excavation led by Dezső Csallány. The skeleton of an adult female with Europid characteristics in the skull, oriented in a S–N direction, was uncovered from the grave. Grave goods included a bronze ring, a bronze belt buckle, a fragment of an iron knife or awl, and a gold ring. Radiocarbon dating, partially overlapping with the archaeological findings, placed the burial between the second half of the 3rd century CE and the early-5th century CE.

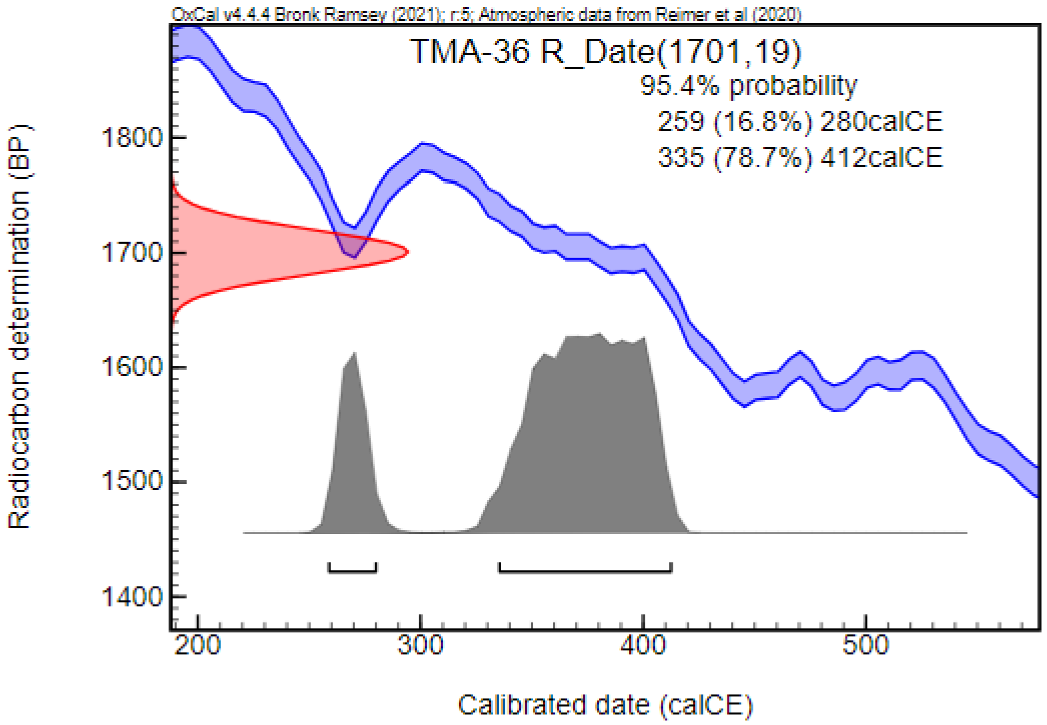


#### Târgşor (1960) (TAR; Prahova County, Romania) [127]

Archaeological excavations began in 1956 near the present-day city of Ploiești, at the medieval settlement of Tîrgşor. Between 1959 and 1961, a burial ground with multiple layers was uncovered belonging to the Bronze Age, Iron Age, Sarmatian period, Migration period, Middle Age, and the Modern Age (extending to the 18th century CE). A total of 286 graves were documented, of which 20 were from Phase I – the Sarmatian period. These graves, located in the eastern and northern parts of the site, were oriented in a N–S direction with occasional deviations toward NE or NW. In one instance, the grave was covered with flat stones. With one exception, the skeletons were found in a supine position with straightened extremities. However, the evaluation of the burial customs and archaeological material was limited as some burials had been disturbed by robbing. Grave goods recovered included ceramic vessels, spindle whorls, mirrors, brooches, earrings, glass beads, and belt buckles. The inventory was primarily composed of ceramic vessels. Their shapes and ornaments show analogies with traditions of the local population, the Roman Empire, and the northern Black Sea region. The Sarmatian cemetery was dated to the 3rd century CE based on archaeological material and stratigraphic superpositions, in some cases with burials and other features dated to the previous centuries or the following centuries, associated with the Cerneahov–Sîntana de Mureș culture. Anthropological analysis by Dardu Nicolăescu-Plopşor revealed that the series included 8 adults and 2 sub-adults, with 1 female and 7 males identified. It was also noted that traces of artificial cranial deformations were observed in more than 50% of the skulls.

Phase III (A, B, and C), containing 101 cremation and 158 inhumation burials dated to the 4th century CE, is the largest group at the site. Identifying burials belonging to this phase was problematic due to similarities with other phases in burial customs and recovered artifacts. Nevertheless, in addition to both inhumation and cremation burials, this phase is characterized by graves oriented in a N–S direction, with some oriented in a E–W, W–E or S–N. Smaller or lager stones covering the grave were described in some cases. The most intriguing characteristic is the high number of ceramic vessels – 443 vessels were uncovered, 211 from cremation burials and 232 from inhumation graves. Other grave goods included spindle whorls, iron and bronze knives, belt buckles (iron, bronze, and in two cases low-quality silver), brooches (bronze, iron, in two cases silver, and in one case white metal), jewelry (pendants, finger rings, earrings), Roman coins, bone needle cases, bone combs,  glass cups (and other glass fragments), beads made of various materials (e.g., glass, amber), and animal bones. This phase was associated with the Cerneahov–Sîntana de Mureș culture. In addition, close analogies were discovered with the Roman Empire, the local populations, and the northern Black Sea region in both burial customs and recovered artifacts. Anthropological analysis of the inhumation burials identified 28 adults and 59 sub-adults. In addition, 7 females and 16 males were described, though many skeletons were unfit for analysis due to poor bone preservation.

We could include 2 samples from Phase I and 1 sample from Phase III in our analysis.

***TAR-118: Grave No. 118***

The skeleton of an adult male in a supine position with straightened extremities was found in the grave. Four larger stones were found above the deceased, one above the skull, one above the chest covering the entire left humerus, one covering the pelvis, and one covering the tibiae and brooches. The burial was dated to the 4th century CE and was associated with the Cerneahov–Sîntana de Mureș culture. Radiocarbon measurement supported an earlier dating, positioning the burial from the 2nd half of the 1st century CE to the beginning of the 3rd century CE.

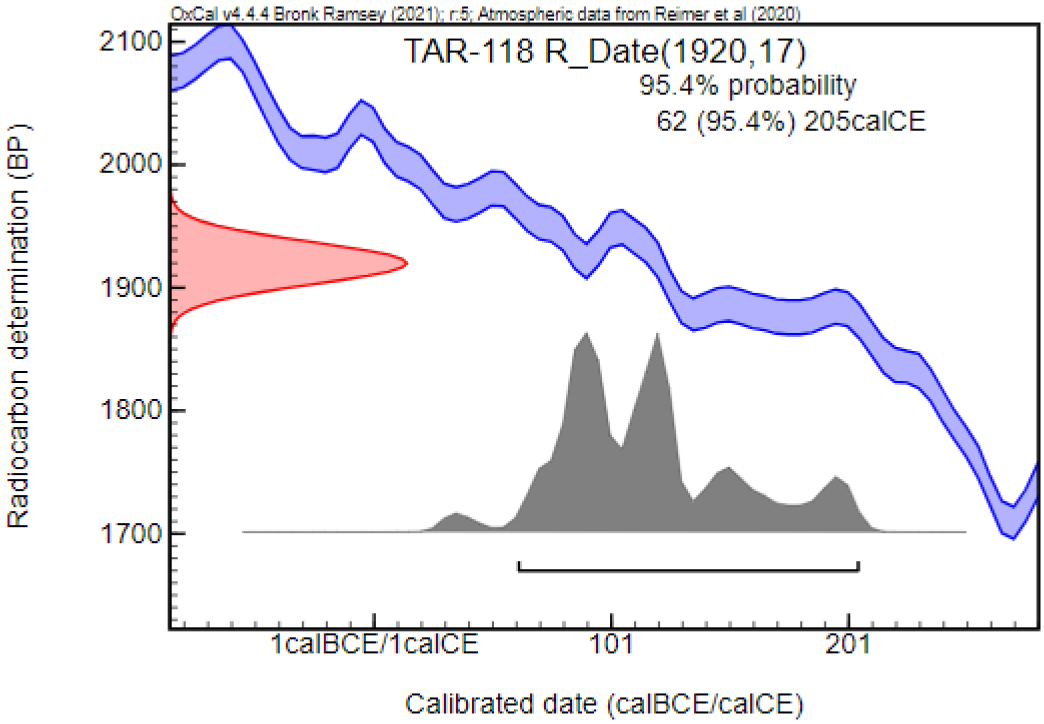


***TAR-184: Grave No. 184***

The skeletal remains of an adult female with traces of artificial cranial deformation were found in the grave. The grave goods included ceramic vessel, a spindle whorl, a mirror, a bronze earring, a string of beads (including glass, carnelian, and coral beads), and a bronze pendant. The burial was associated with the Sarmatians, i.e., Phase I of the cemetery.

***TAR-196: Grave No. 196***

This was a disturbed burial of an adult male oriented in a N–S direction. The skeleton was in a supine position with straightened extremities and traces of artificial cranial deformation were detected. The grave goods included a hand-made ceramic vessel, a wheel made ceramic jar, and two astragali of a lamb. The skeleton was partially in superposition with a pit dated to the 2nd–3rd centuries CE. The burial was associated with the Sarmatians, i.e., Phase I of the cemetery. Radiocarbon measurement supported an earlier dating, positioning the burial from the 1st century BCE to the 1st century CE.

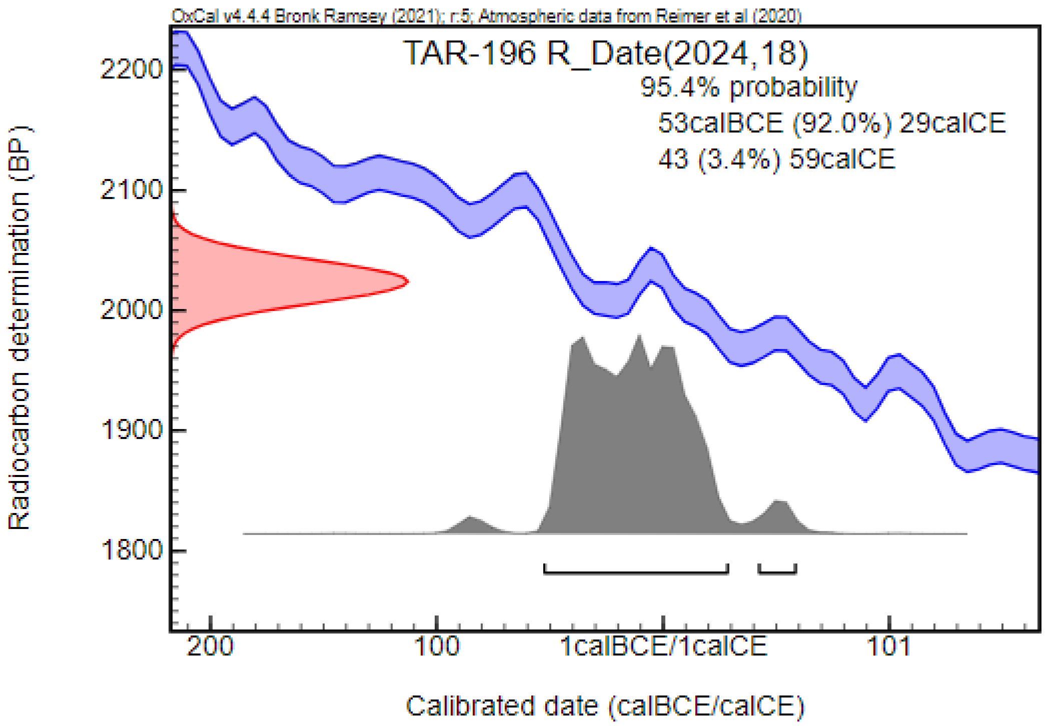


#### Târgu Frumos (TAF; Iași County, Romania) [128]

Following a field assessment in 2018, a barrow (Tumulus No. 1 or T1) near the current city of Târgu Frumos was thoroughly investigated in 2019 and 2020. Twelve burials were uncovered in the mound: ten were dated to the Bronze Age, one was from an unknown period due to significant destruction, and one was associated with the Sarmatians, dated to the 2nd century CE. To the best of our knowledge, the skeletons have not yet undergone anthropological analysis. We included the Sarmatian sample in our analysis.

##### *TAF-11: Tumulus No. 1, Grave No. 11*

This burial was discovered in the southern sector of T1. The rectangular grave pit was oriented in a NNE–SSW direction. The skeleton was in a supine position with straightened extremities. The grave goods included an iron knife, a rectangular bone plate, four beads, and a ceramic vessel. Radiocarbon measurement, supporting the archaeological dating, positioned the burial mostly to the 2nd century CE and the beginning of the 3rd century CE, with a possibility of it being related to the end of the 1st century CE.

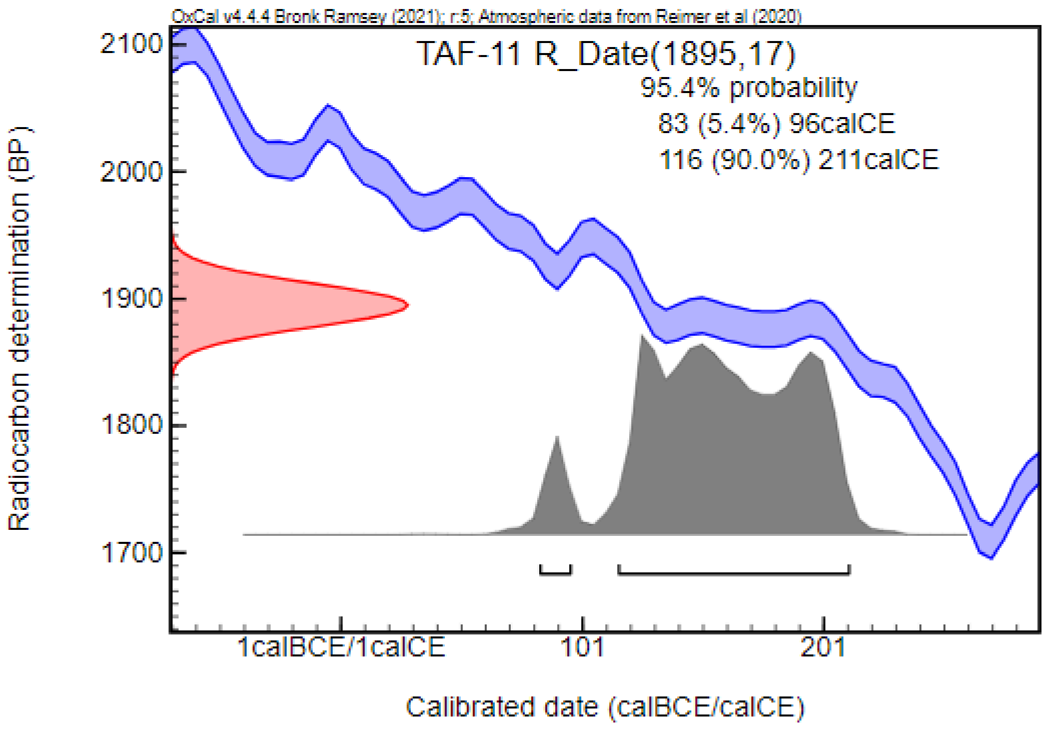


#### Tiszadob – Sziget (TD; Szabolcs-Szatmár-Bereg County, Hungary) [92,129]

During soil works related to the construction of the defensive system against the flooding of the Tisza River, burials were destroyed in 1964. András Gombás, a teacher from the neighboring town of Tiszavasvári, recorded 12 burials and additional stray finds during that period. Given the numerous archeological finds known from this area, systematic excavations were conducted by Eszter Istvánovits between 1983 and 1990, revealing the remaining parts of the cemetery. A total of 38 graves, including three cremation burials, were documented. Among the inhumation burials, two main groups were distinguished based on grave orientation. The first group consists of 23 S–N and SW–NE oriented graves, while the second group contains six graves oriented in a W–E and NW–SE direction. Most of the burials were affected by contemporary robbing. In some instances, remains of coffins, including iron coffin clamps, were found. Artifacts recovered from the burials included jewelry (e.g., earrings, strings of beads, rings, and pendants), clothing elements (e.g., belt buckles, brooches, and beads), tools (iron knives, needle cases), coins, ceramic and glass vessels, and weapons (e.g., sword, spearhead, and shield buckle). The cemetery was dated to the last third of the 4th century CE and first third of the 5th century CE. This burial ground illustrates the population and cultural changes that occurred in the late Sarmatian period. While some of the burial customs and grave goods observed in the cemetery are associated with the Sarmatians, connections with Germanic groups, the Roman Empire, the Marosszentanna/Sântana de Mureş–Cherniakhov, and the Przeworsk Cultures are also indicated. The results of the anthropological analysis are not yet published, and only preliminary reports are available. The series show parallels with the population of the Madaras–Halmok cemetery, including the morphological characteristics of the skulls and the presence of artificial cranial deformation in the series. We could include only one sample in our analysis.

***TD-SZ: Stray find (accessory number: 68.47.11)***

During the discoveries by András Gombás, stray finds, including skeletal remains belonging to an adult male, were recovered. Radiocarbon dating, partially overlapping with the archaeological findings, dates the bones from the mid-3rd century CE to the last third of the 4th century CE.

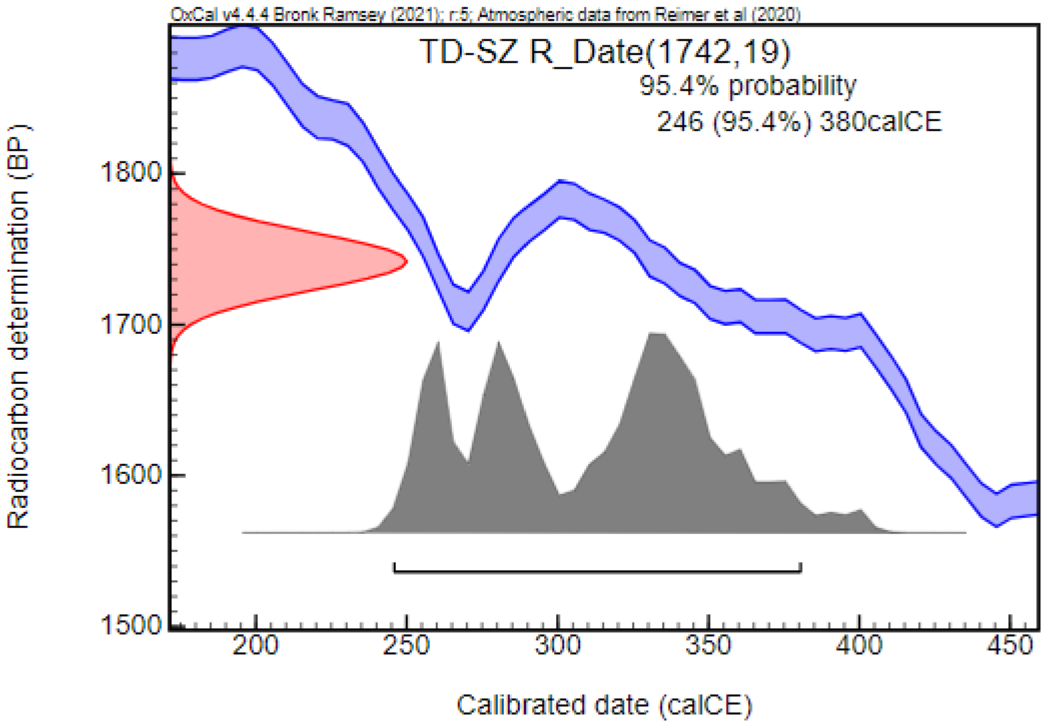


#### Tiszavalk (TIV; Borsod-Abaúj-Zemplén County, Hungary) [130]

The archaeological site is located approximately 1 km southwest of Tiszavalk, near the Tisza dam, in an area called Kenderföld. The first excavation, led by Ervin Mérey-Kádár, was conducted in 1954, uncovering features from the Copper Age and the Sarmatian period. Rescue excavations led by Pál Patay were conducted in 1966, 1967, and 1975, following the destruction of a Copper Age burial in 1966 during the construction of the Tisza dam. A total of 57 Copper Age graves, 21 late Sarmatian-period burials, additional burials from other periods, as well as three houses from the Árpádian-Age, were uncovered. The cemetery is considered partially excavated, with possible graves remaining outside the explored area. Most of the Sarmatian graves were found in one group, except for two graves in the southern part and one grave in the northern part of the site. The graves were secondarily dug into pits from either the Copper Age or the Sarmatian period. The burials were primarily oriented in an S–N direction, with occasional deviations toward the west. Several graves were affected by contemporary robbing. In some instances, traces of wooden coffins and shrouds were detected. The deceased were primarily placed in a supine position, with a few exceptions where the skeletons were found with bent knees. Artifacts recovered from the burials included jewelry (e.g., strings of beads, a bracelet), clothing elements (e.g., brooches, belt buckles, and beads), tools (e.g., iron knives, a needle case, spindle whorls), weapons (e.g., spearheads, a sword, and shield buckles), Roman coins, and ceramic vessels. The cemetery was dated between the last third of the 4th century CE and first third of the 5th century CE. Burial customs and grave goods show parallels with Tiszadob-type cemeteries, indicating a multi-cultural population. Anthropological data on the series has not yet been published, and the identification of biological sex and age-at-death estimation was based on archaeological records. We could include five samples in our analysis.

##### *TIV-1: Grave No. 1*

This S–N oriented grave contained the skeleton of an adult male in a supine position. Grave goods included an iron brooch with returned foot, an iron belt buckle, fragments of an iron knife, a silver Roman coin (Antonius Pius denarius), an iron spearhead, and a ceramic vessel.

##### *TIV-6: Grave No. 6*

The skeleton of an adult male was found in a supine position in this S–N oriented grave, accompanied by a two-piece silver brooch with returned foot, an iron belt buckle and bronze belt fittings, chalcedony beads, an iron knife, a ceramic vessel, a conical shield buckle (type Dobrodzień), an iron shield handle, a fragments of an iron sword, an iron spur, and an iron spearhead.

***TIV-9: Grave No. 9***

This burial, oriented in an S–N direction, contained the skeleton in a supine position. Grave goods included a two-piece iron brooch, an iron belt buckle, an iron knife, a Roman silver coin (Lucius Verus denarius), and a ceramic vessel

***TIV-13: Grave No. 13***

A skeleton in a supine position and a two-piece bronze brooch were discovered in the S–N oriented grave. Radiocarbon dating, partially overlapping with the archaeological findings, placed the burial between the mid-3rd century CE and the beginning of the 5th century CE.

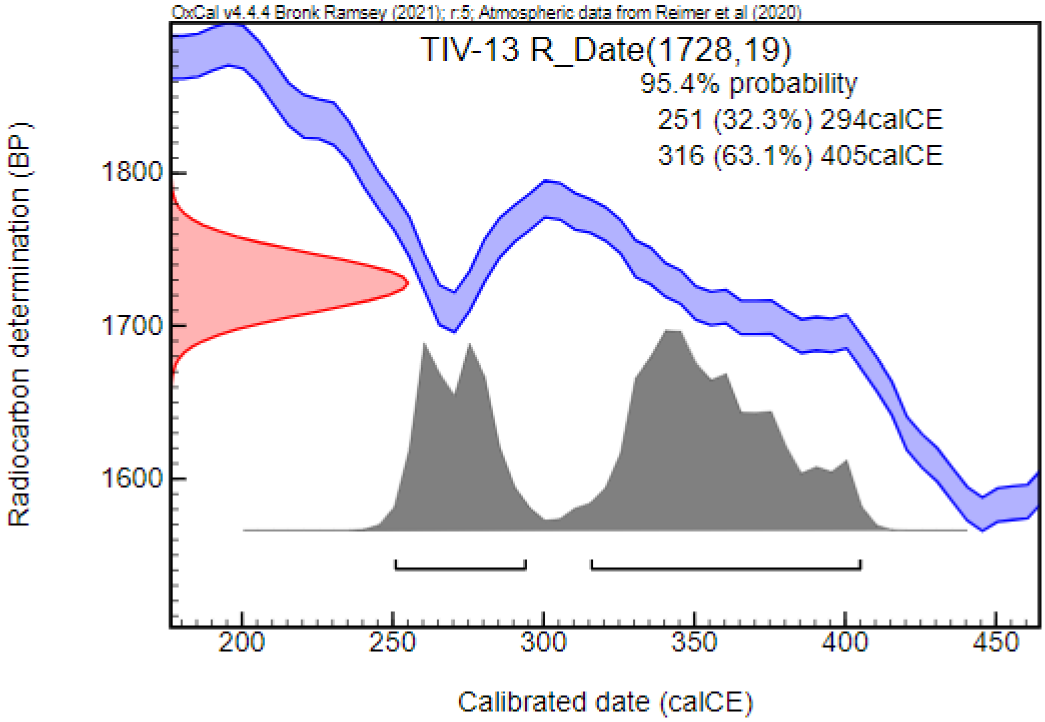


***TIV-17: Grave No. 17***

This S–N oriented burial contained the skeletal remains of an adult male in a supine position. Grave goods consisted of a fragment of a two-piece iron brooch with returned foot, a Roman silver coin, a fire lighting tool, two iron knives, an iron tool (possibly chisel), iron fragments of unknown artifacts, an iron spearhead, an iron shield buckle, an iron shield handle, and a ceramic vessel.

#### Trestiana (TRE; Vaslui County, Romania) [131,132]

A Neolithic settlement was discovered near the present-day town of Trestiana during the construction of a duck lake in 1960. Archeological excavations, conducted from 1964–1993, 1996, and 2001, uncovered a Neolithic settlement and a total of 42 burials. Of these burials, 11 were identified as Neolithic, 5 were dated to the Bronze Age, 3 to the Iron Age, 1 to the Migration period, and 21 had unknown dates. We included the burial dated to the Migration period in our analysis.

***TRE-33: Grave No. 33/1972***

This was the burial of a sub-adult individual (2–3 years old) oriented in an E–W direction. The skeleton was found in a supine position with straightened extremities, and traces of artificial cranial deformations were detected. The burial did not contain any known grave goods, and the archaeological dating was based on the anthropological characteristics, specifically the deformation of the skull. Radiocarbon dating, supporting the archaeological assumptions, placed the burial from the second third of the 5th century CE to the mid-6th century CE.

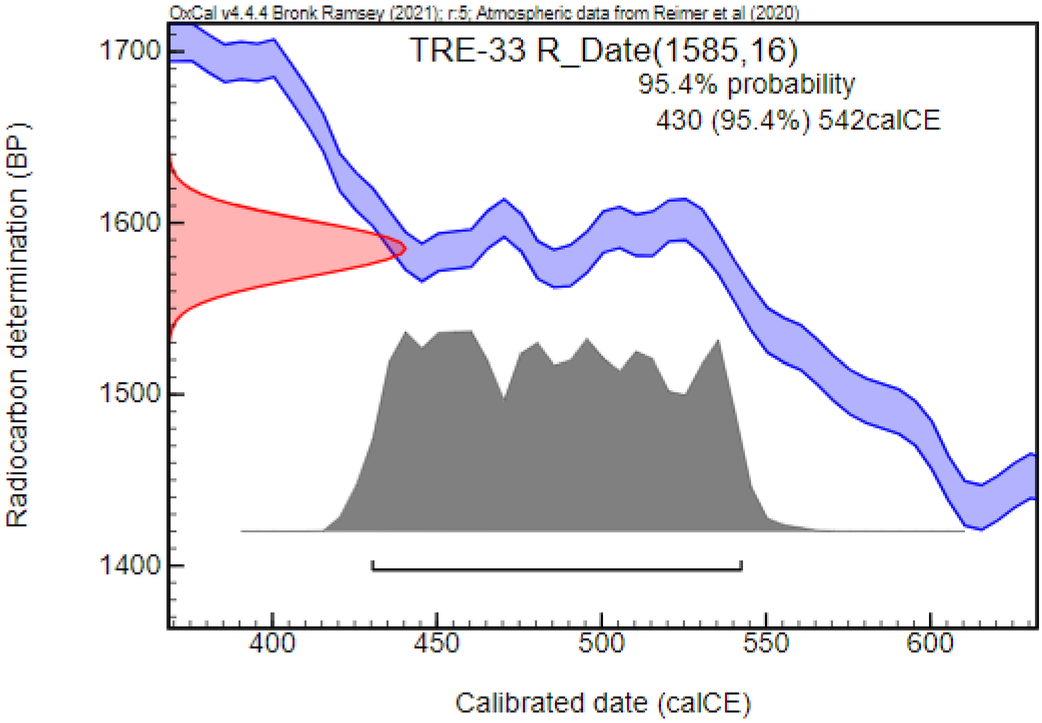


#### Visegrád – Gizellamajor (VIG; Pest County, Hungary) [133,134]

A late Roman fortification, settlement objects, and burials from the Late Roman and Hun periods were discovered during preventive excavations related to the construction of the Bős–Nagymaros hydropower plant. The fortress was built during the rule of Emperor Constantine II (337–340 CE) and remained in use into the first third of the 5th century CE. Additionally, features dated to the middle third of the 5th century CE, distinct from the original function of the fortress, have also been evidenced. Among them, two Hun period burials were discovered, secondarily dug into and next to buildings inside the fortress. The archaeological and anthropological investigation is still in progress. The skeletons of adult females with traces of artificial cranial deformation were found in both burials, accompanied by a rather poor inventory that included jewelry and cloth fittings. We could include one sample in our analysis.

##### *VIG-94/1: Grave No. 94/1*

The skeleton of an adult female with traces of artificial cranial deformation was uncovered inside the northern wing of a building located in the western part of the fortress. Grave goods included bell buttons found near the skeleton, ranging from the feet to the neck. Radiocarbon dating, partially overlapping with the archaeological findings, placed the burial between the end of the first third of the 5th century CE and the second third of the 6th century CE.

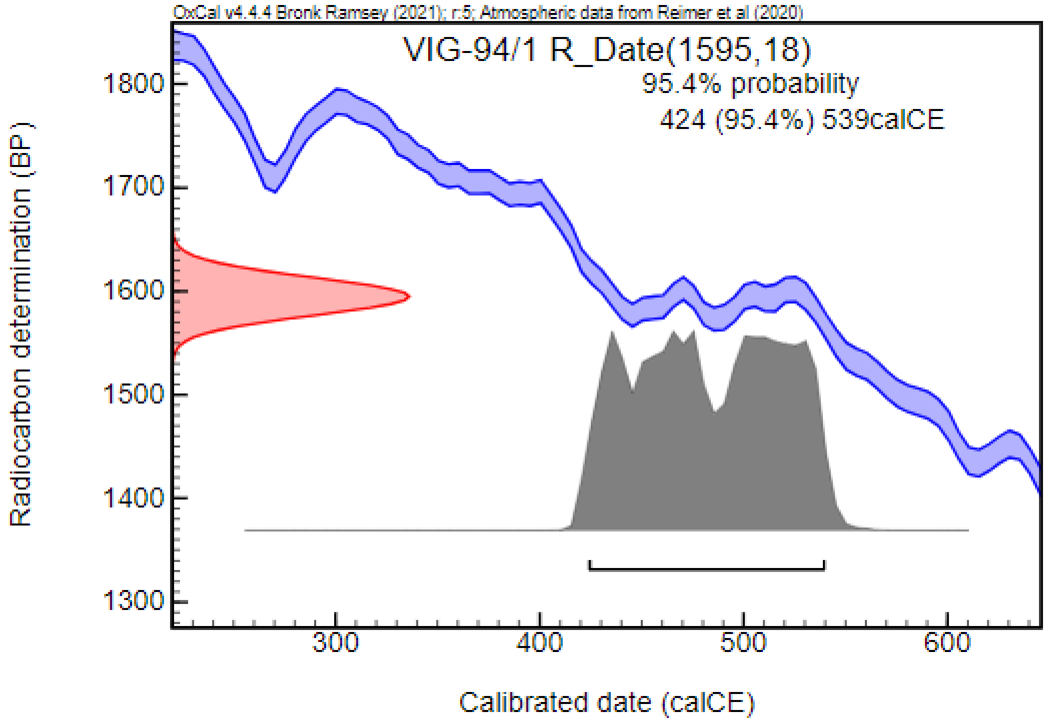


#### Zákányszék – Zákánydűlő NY/69 (ZZ; Csongrád-Csanád County, Hungary) [135,136]

##### *ZZ-1: Grave No. 1*

In 2004, three burials dated to the Sarmatian period were excavated by Gábor Sánta. The anthropological remains were analyzed by Zsolt Bereczki and his colleagues. The graves were disturbed and robbed, two of them contained only some human bone fragments. Solely the burial No. 1 was fit for archaeological and anthropological analysis. A spindle-whorl, an iron knife, and 20–30 beads were found in the grave. The skeletal remains belonged to a middle-adult female, and traces of surgical trepanation were observed on the skull. Radiocarbon measurement dated the burial from the first third of the 2nd century CE to the first third of the 3rd century CE.

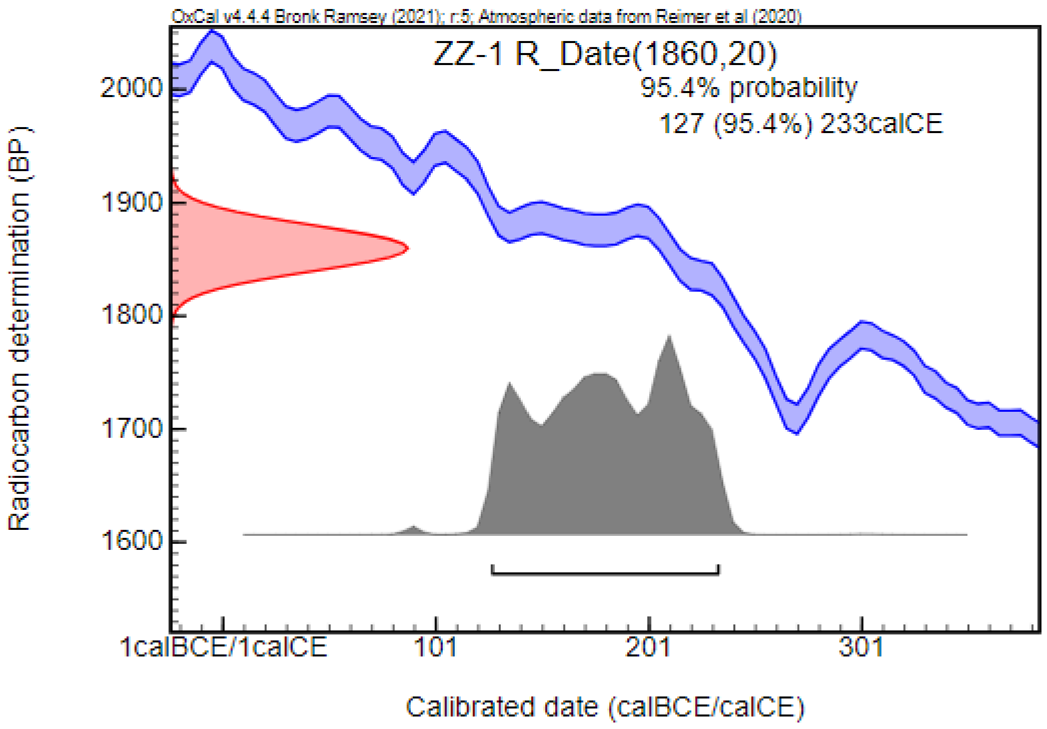


#### DESCRIPTION OF THE STUDIED IRON AGE SAMPLES

##### Lișcoteanca – Movila Olarului (LMO; Brăila County, Romania) [90]

Between 1970 and 1976, twelve burials from the Sarmatian period, along with ten graves from the Bronze Age and two Petcheneg graves, were discovered during the excavation of a Neolithic mound. Originally, the burial involved in our study was associated with the Sarmatians. However, radiocarbon analysis revealed that it dates to the Iron Age.

***LMO-8: Grave No. 8***

The skeleton of an adult male in a supine position was found in the grave, oriented in a W–E direction. A bronze arrowhead was found between the ribs, in the area of the left thoracic cavity. Radiocarbon dating placed the burial to the Iron Age, between the end of the 9th century BCE and mid-8th century BCE, with the possibility of being dated to the first third of the 7th century BCE (4.7%), the last third of the 7th century BCE (0.9%), or the end of the 7th and beginning of 6th centuries BCE (5.3%).

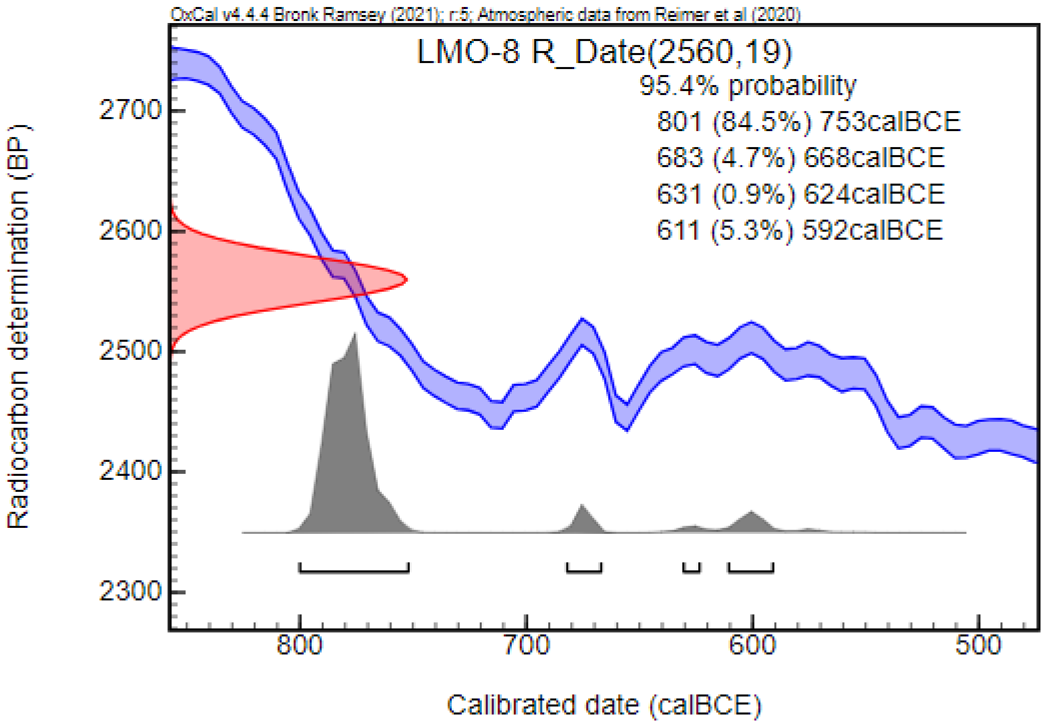


**Râmnicelu (1969) (RAM; Brăila County, Romania)** [90,114]

This burial ground, described in the context of Sarmatian period samples, includes a burial originally listed among the Sarmatian samples by archaeologists. However, radiocarbon analysis revealed that it dates to the Iron Age.

***RAM-7: Grave No. 7***

This burial, oriented in an N–S direction, contained the skeleton of an adult male in a supine position. Grave goods included a destroyed iron buckle (?). Radiocarbon dating placed the burial between the mid-8th century BCE and the last third of the 5th century BCE.

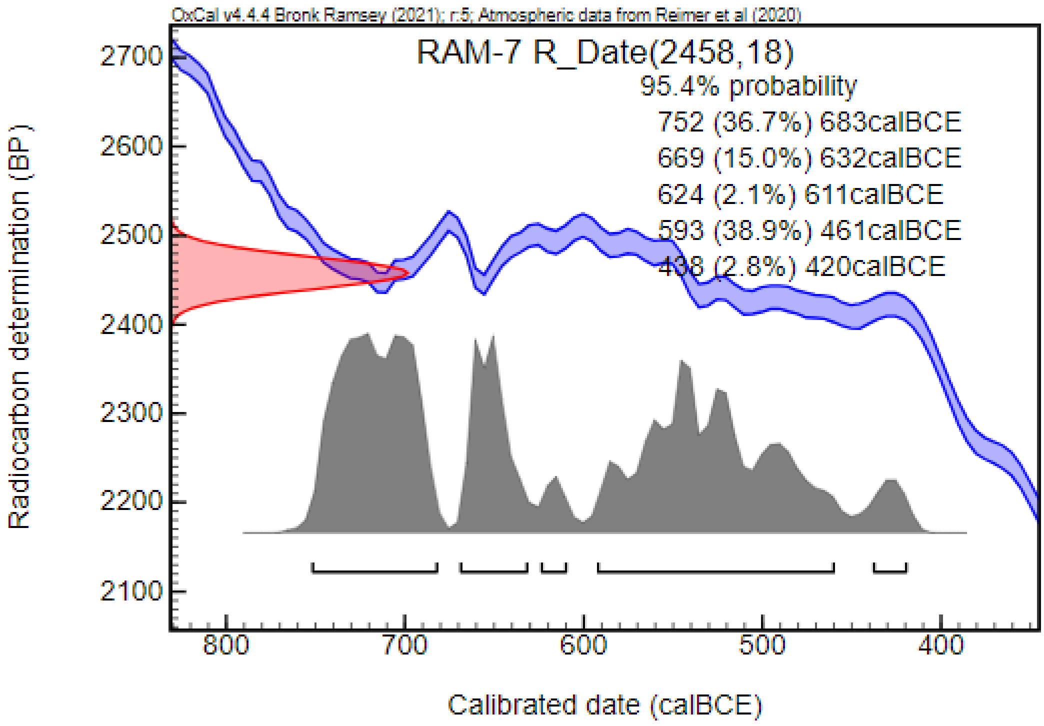
